# Characterizing the Small Non-Coding RNA Pathways in the Invasive Zebra Mussel (*Dreissena polymorpha*)

**DOI:** 10.64898/2026.07.10.737777

**Authors:** Víctor H. Hernández Elizárraga, Lindsey G. O’Brien, Scott Ballantyne, Daryl M. Gohl

## Abstract

The zebra mussel (*Dreissena polymorpha*) is an invasive species that causes extensive economic and ecological damage. Here, we identify and characterize the key components of the small RNA (sRNA) and RNA interference (RNAi) pathways in zebra mussels. Like other mollusks, zebra mussels have extensive microRNA (miRNA) and Piwi-interacting RNA (piRNA) machinery but lack or have modified canonical factors needed to produce small interfering RNA (siRNA). Specifically, the zebra mussel Dicer sequence displays substitutions in the conserved DEAD box motif that is required for substrate processivity, and this organism also lacks some attendant accessory factors such as R2D2. We sequenced the small RNA found in both isolated somatic tissue (adductor muscle) and whole animals (including germline), and identified both conserved and novel miRNA and diverse piRNA sequences, but few endogenous siRNAs. To determine whether their remaining sRNA machinery could still be co-opted to initiate gene silencing, we injected dsRNA targeting several genes into zebra mussel adductor muscle. The injected *rpn8*-targeting dsRNA reduced *rpn8* mRNA levels and was processed into sRNA that resemble endogenous miRNAs and piRNAs. The levels of both sRNA types correlated with mRNA knockdown, suggesting that they may act together to initiate RNAi as seen elsewhere. dsRNA targeting other genes produced variable results suggesting that particular criteria may be needed to trigger an RNAi response in this assay. Our results characterize endogenous sRNA pathways in zebra mussels, establish that dsRNA can induce RNAi, and lay the groundwork for further optimizations to establish RNAi-based genetic manipulation tools for this damaging invasive species.

## Introduction

The zebra mussel, *Dreissena polymorpha (D. polymorpha),* is believed to be native to the Caspian and Aral Seas, as well as the low-salinity lagoons of the Black Sea and adjacent rivers (Ludyanskiy et al. 1993). This invasive species has impacted many waterbodies in North America and Europe. In North America, zebra mussels were first reported in the Great Lakes in 1988 (Hebert et al. 1989) and have since spread to at least 32 states in the USA. Zebra mussels can quickly attain large population sizes, leading to significant economic and ecological impacts. Zebra mussels attach to water intake pipes in power stations, nuclear reactors, and drinking water utilities, disrupting essential infrastructure. In addition, their presence impacts recreation, water-based transport, and property values by attaching to boats, docks, and shorelines. Ecologically, zebra mussel impacts include alteration of food webs and decreasing the abundance and diversity of other macrobenthic species (Karatayev et al. 2002). They harm ecosystems by competing with native species for food and space (Baker and Levinton 2003) while also accumulating pathogenic bacteria and parasites that increase disease risk (Mosteo et al. 2016). It is estimated that annual costs for mitigating damage caused by zebra mussels exceeds $1 billion in the USA alone (Haubrock et al. 2022; Pimentel et al. 2005; Warziniack et al. 2021). Despite the zebra mussel’s significant economic and ecological impact, there is little information on molecular pathways like RNAi, which could potentially be harnessed for genetically-targeted biocontrol.

Non-coding RNAs have diverse functions in developmental gene regulation, pathogen defense, and the promotion of genome stability through the repression of mobile genetic elements. Based on their size, non-coding RNA can be classified into small and long non-coding RNA (sncRNA and lncRNA, respectively). sncRNA, also known as small RNA (sRNA), refers to short ribonucleic acid molecules (typically in the range of ∼20-35 nucleotides long). Small regulatory RNAs control gene expression by guiding effector proteins to target nucleic acid sequences. The three most studied examples of small regulatory RNAs are: small interfering RNA (siRNA), microRNA (miRNA), and Piwi-interacting RNA (piRNA) (Xiong and Zhang 2023).

siRNA and miRNA inhibit mRNA expression through RNA interference (RNAi). The siRNA or miRNA binds to Argonaute (AGO) proteins and their protein partners, and guides the RNA-induced silencing complexes (RISC) to target mRNA. miRNA-RISC complexes inhibit mRNA translation (Wightman et al. 1993) and mRNA stability (Eichhorn et al. 2014), while siRNA-RISC complexes induce mRNA degradation (Elbashir et al. 2001). siRNAs have more stringent base-pairing requirements with their target mRNA than miRNAs. Both siRNA and miRNA are derived from double-stranded RNA (dsRNA). siRNAs are made from two chains of perfectly complementary RNA, while miRNAs originate from a single RNA chain that is self-complementary but contains mismatches and bulges. Another key difference between siRNAs and miRNAs lies in their biogenesis machinery (Svoboda 2020). miRNA production is typically initiated inside the nucleus. First, the RNase III enzyme, Drosha, cleaves primary miRNA (pri-miRNA) stem-loops into ∼70-nucleotide precursor miRNAs (pre-miRNAs). This precursor is recognized by the nuclear export factor, Exportin-5 (via a Ran–GTP-dependent mechanism), and transported into the cytoplasm for final processing (Davis and Hata 2009; Murchison and Hannon 2004). Conversely, siRNA biogenesis bypasses the nucleus entirely. It occurs in the cytoplasm, typically originating from long, double-stranded RNA (dsRNA) introduced from outside the cell, such as from viral infections or experimental injections (Snead and Rossi 2010). Because the starting material is already in the cytoplasm and possesses a fully double-stranded structure, it requires no prior processing or export by Exportin-5.

piRNAs get their name from their association with Piwi-like Argonaute proteins. Piwi was first identified in *Drosophila* and is short for *P-element-induced wimpy testes*, reflecting the male sterility phenotype caused by mutation of the *piwi* gene (Lin and Spradling 1997). The Piwi/piRNA system protects the animal germline genome by inhibiting transposons, and this function is preserved throughout all metazoa. piRNAs have also been described in the soma, and this is particularly common in bivalves (Jehn et al. 2018). The function of somatic piRNAs is less well understood, but growing evidence points to a role in inhibiting foreign RNA replicators (Morazzani et al. 2012; Rafanel et al. 2025; T. Yu et al. 2025) and acting as master suppressors of transposable elements (TEs) (Rivera et al. 2025; Signor et al. 2023). piRNAs are made from single-stranded RNA that is encoded by the cell’s genome. piRNA loci can be clustered or single, with the latter still capable of eliciting a strong response. They are found in intergenic and genic regions, particularly in 3’UTRs (Robine et al. 2009). piRNAs guide Piwi complexes to target RNA and DNA via base pairing, inhibiting gene expression transcriptionally and post-transcriptionally. Some species exhibit “ping-pong amplification,” where Piwi RNA products feed back to amplify piRNA production (Czech and Hannon 2016). piRNAs are ∼28-34 nucleotides long and can be easily distinguished from miRNAs and siRNAs, which are ∼18-21 nucleotides in length. In addition to size, piRNAs have other hallmarks such as 3’ 2-O-methylation (Tian et al. 2011) and Piwi association (Tosar et al. 2018). piRNA can also regulate cytoplasmic mRNA, and this has been exploited in some systems to provide a programmable method for selective gene inactivation, known as piRNAi (Mondal et al. 2020; Priyadarshini et al. 2022).

Although RNAi is broadly conserved among eukaryotes, there is considerable mechanistic variation between species. For instance, RNAi is systemic, amplified, and sometimes heritable in *Caenorhabditis elegans (C. elegans)*, but not in *Drosophila melanogaster (D. melanogaster)*, where it’s restricted to cells producing the RNAi trigger. A diverse array of small regulatory RNAs can initiate or perpetuate RNAi. While RNAi has been reported in some bivalves (Han et al. 2023; R. Zhao et al. 2026), the repertoire of small regulatory RNAs, as well as sRNA biogenesis and effector proteins, differs from model systems (S. Huang et al. 2024). Here, we deliver RNAi constructs via injection and use sRNA sequencing and comparative genomics to characterize the biogenesis and processing of small regulatory RNAs in zebra mussels. Understanding the small noncoding RNA and RNAi-based regulatory systems in zebra mussels will enable the specific knock-down of target mRNAs, facilitating both basic research and potentially applied methods for controlling this damaging invasive species.

## Methods

### Analysis of key sRNA biogenesis and RNAi elements in zebra mussel

A manual search was conducted for key sRNA biogenesis and RNAi machinery annotations in the zebra mussel genome (McCartney et al. 2022) (NCBI accession number: GCA_020536995.1) (Supplementary Table 1). A presence/absence comparison was made against RNAi factors from model species (*Homo sapiens (H. sapiens), D. melanogaster, C. elegans,* and *Arabidopsis thaliana (A. thaliana)*) to determine the conservation of the zebra mussel RNAi machinery using Jaccard similarity. The overlap in gene factor presence across different species was calculated from a binary matrix to compute the Jaccard distance, and hierarchical clustering was used to group species with similar gene factors. In addition, two custom databases were created for the relevant RNAi protein partners, Dicer (47 entries) and Argonaute (87 entries) proteins. These databases were used for multiple sequence analysis with Clustal Omega to infer phylogenetic relationships of *D. polymorpha* Dicer-like (NCBI accession number XP_052272928.1), Argonaute 2-like (NCBI accession number XP_052234982.1), and Piwi-like protein 1 (NCBI accession number XP_052263121.1) proteins. The domains of Dicer 1, Argonaute 2, and Piwi 1 proteins from *H. sapiens, D. melanogaster, C. elegans*, and *A. thaliana* were compared with their *D. polymorpha* homologs using the Simple Modular Architecture Research Tool (SMART) (Schultz et al. 1998).

### Mussel collection

We sampled zebra mussel (*D. polymorpha*) adults ranging from 2 to 3 cm in length at Grays Bay, Minnetonka Public Landing, Minnesota, USA (44° 57’ 03.4“ N, 93° 29’ 54.8” W) by snorkeling or SCUBA diving (Prohibited Invasive Species Permit #826, valid from 10/24/2023 through 12/31/2028; Minnesota Department of Natural Resources (DNR) Division of Ecological and Water Resources). We kept the mussels at the Minnesota Aquatic Invasive Species Research Center (MAISRC) Containment Lab in 50-gallon fresh water tanks and fed them a diet of *Chlamydomonas reinhardtii (C. reinhardtii) ad libitum*.

### RNA extraction and small RNA library preparation

Total RNA was extracted from adult mussels (whole animal, n=9) and adductor muscle tissues (n=6) using the TRIzol protocol (Thermo Fisher Scientific, Cat. No. 15596026) following the manufacturer’s instructions (Thermo Fisher Scientific document #MAN0016385 Rev. A.0). Samples were DNase-treated on-column using the RNase-Free DNase Set (Qiagen, Cat. No. 79254). For small RNA library preparation, 1 ug of total RNA in 5 uL of nuclease-free water (per sample) was processed using the TruSeq Small RNA Library Preparation Kit (Illumina, Cat. No. RS-200-0012) following the manufacturer’s protocol. The TruSeq Small RNA kit uses engineered ligase enzymes and specialized adapters that only bind to RNA molecules possessing the specific 5′-PO_4_ / 3′-OH signature. This naturally filters out degraded mRNA or ribosomal RNA fragments, which typically lack these specific end modifications. Library quality and size distribution were evaluated using an Agilent 2100 Bioanalyzer with the High Sensitivity DNA Kit (Agilent, Cat. No. 5067-4626). Target size selection (∼160 bp, corresponding to small RNA inserts of ∼20–30 nt plus adapter sequences) was performed using a 3% PippinHT agarose cassette (Sage Science) to isolate small non-coding RNAs and deplete larger RNA species, including intact ribosomal and messenger RNAs. Final library size and purity were verified post-selection on a Bioanalyzer High Sensitivity DNA chip.

### Library conversion and sequencing

For sequencing, Illumina indexed libraries were equimolal pooled (1.5 nM) and sequenced on an Element Biosciences’ AVITI instrument using a CloudBreak Freestyle sequencing kit (Element Biosciences, Cat. No. 860-00015) with a 5% PhiX spike-in.

### Data preprocessing

After sequencing, samples were demultiplexed and sequencing adapters were removed using Cutadapt software (Martin 2011). The sequence TGGAATTCTCGGGTGCCAAGG was used for adapter trimming (TruSeq small RNA kit). The quality of the adapter-trimmed sequences was assessed using FastQC software (“Babraham Bioinformatics - FastQC A Quality Control Tool for High Throughput Sequence Data,” n.d.). In order to remove potential long non-coding RNA (lncRNA) sequence contaminants remaining after physical size selection, bioinformatic filtering of tRNA and rRNA sequences from Rfam (Ontiveros-Palacios et al. 2025) was carried out using Bowtie2 (Langmead and Salzberg 2012). The remaining sRNA reads were split into two sets (1-24 nt and > 25 nt) for further analysis.

### miRNA identifications and mRNA target prediction

MirDeep2 software (Friedländer et al. 2012) was employed to identify known and unknown miRNAs in zebra mussel datasets using default parameters. Briefly, the mapper.pl resource was used to map reads (1-24 nucleotides long dataset) against the *D. polymorpha* reference genome (NCBI accession number: GCA_020536995.1). Read counts were normalized using the quantifier.pl module. For known miRNA identification, the mature.fa dataset from miRBase (https://www.mirbase.org/, accessed January 2025) was used as an input file for mature miRNAs in other species. Based on miRDeep2 predictions, novel and known miRNA sequences were recovered.

Predicted mature miRNA sequences from whole-animal (n=9) and adductor muscle (n=6) samples were filtered as follows: sequences with fewer than 5 CPM and a miRDeep2 score < 10 were removed. Clustering of miRNA isoforms was performed using CD-HIT-EST (Li et al., 2006) with the following parameters:-c 0.85-n 6-T 0-M 0-G 0-aS 0.9-g 1. Shared miRNA clusters were visualized with an UpSet plot using custom R scripts. The whole-animal and adductor muscle samples with higher sequencing depth were selected for downstream bioinformatic analysis.

Precursor and mature sequences were used to classify miRNAs into 5’ and 3’ miRNAs using a custom R script and the probability of base occurrence was calculated for seed sequences (≤8 nucleotides) for both 5’ and 3’ miRNAs using the seqLogo (Bembom 2007) package from R. Loop lengths and distances between mature and star sequences were calculated for all miRNA sequences using custom R scripts. Putative miRNA target sequences were predicted using miRanda software (Enright et al. 2003) employing default parameters. Mature miRNA and zebra mussel transcriptome sequences (NCBI accession number: GCF_020536995.1) were used as input to carry out the miRNA target prediction. Sequences predicted to be targets were filtered based on miRanda scores, and the top 50,000 hits were used to perform KEGG enrichment analysis.

### Trans-species miRNAs

To identify potential trans-species miRNAs derived from the diet, small RNA reads that remained unmapped to the zebra mussel genome were screened against the *C. reinhardtii* reference genome (NCBI accession: GCF_000002595.2). miRNA identification was performed using miRDeep2 with default parameters, utilizing *C. reinhardtii* miRNA sequences from miRBase (v22.1, accessed March 2025) as sRNA references.

### Piwi-interacting (piRNA) prediction and clustering

The ping-pong signature 10 nucleotide overlap analysis was performed using the ‘‘Small RNA signatures’’ toolkit from the ARTbio project (Antoniewski 2014). Sequences > 25 nucleotides long were used as input and the resulting Z-scores per overlap position were visualized using the ggplot2 (Wickham 2011) package in R. The nucleotide preferences characteristic of ping-pong signatures were visualized with custom R scripts. piRNA prediction and clustering was made with proTRAC (Rosenkranz and Zischler 2012) and the chromosomal distribution was plotted with custom R scripts assisted by the ggplot2 package. Putative piRNA cluster sequences were retrieved from the zebra mussel genome using samtools (Li et al. 2009), and their repetitive elements were annotated via CENSOR(Kohany et al. 2006) against the Eukaryota Repbase reference collection.

### Endogenous small interference RNAs (siRNAs) duplex prediction

Clean mapped reads of 18-34 nucleotides in length were used to uncover Dicer processing signatures with the “Small RNA signatures” toolkit from ARTbio (Antoniewski 2014). This file was used as a reference input to predict all potential perfectly matched duplexes within the 18-34 nucleotide length range. Read counts for potential duplexes, duplex size, and Z-scores were visualized using the Plotly (Sievert et al. 2021) package in R. In addition, stepRNA (Murcott et al. 2022) software was utilized to calculate the overhang distance which is the number of nucleotides that a reference sRNA read extends beyond a query read.

### Double-stranded RNA (dsRNA) *in vitro* synthesis

Five genes from the *D. polymorpha* genome (RNA polymerase II subunit (*rpabc5)*, proteasome 26S subunit (*rpn8)*, ubiquitin-activating enzyme E1 (*e1ubi)*, cytoplasmic polyadenylation element binding protein (*cpeb)*, and ribonucleoside diphosphate reductase (*rnrl*)) were selected to obtain double-stranded RNA (dsRNA). PCR product templates were generated from zebra mussel gDNA, with the T7 promoter sequence (5’-TAATACGACTCACTATAGGG-3’) extensions on both ends (Supplementary Table 2). We carried out *in vitro* RNA synthesis reactions using the HiScribe T7 High Yield RNA Synthesis Kit (New England Biolabs, Cat. Number E2040S) according to the manufacturer’s instructions. Finally, the dsRNAs were cleaned using lithium chloride precipitation, confirmed using agarose gel electrophoresis, and quantified with Qubit™ RNA High Sensitivity (HS) (Thermo Fisher Scientific, Cat. Number Q32852).

### Double-stranded RNA (dsRNA) injection

To establish a protocol for injecting nucleic acids into zebra mussel tissue, we evaluated the ability of Mg^2+^ and acetylsalicylic acid to induce shell gaping, thereby allowing unobstructed access to the adductor muscle. Briefly, mussels were exposed to 0.05 g/mL of various magnesium salts (magnesium chloride, magnesium acetate, and magnesium sulfate) or 0.026 g/mL of acetylsalicylic acid over a 2-hour period, and the percentage of gaped mussels was recorded (Supplementary Figure 1). Additionally, concentrations of 1, 10, 100, and 1000 mg/L of the fish anesthetic MS-222 (ethyl 3-aminobenzoate methanesulfonate; MilliporeSigma, Cat. No. E10521-10G) buffered with sodium bicarbonate to neutral pH were tested to induce sedation. Post-treatment survival was monitored for 4 days following sedation (Supplementary Figure 1). We determined that MS-222 successfully induced sedation in the mussels without causing the mortality observed with the Mg^2+^ salts. For the experiments performed in this study, mussels were anesthetized in a solution of 0.96 mM MS-222 and 5.95 mM sodium bicarbonate prior to the injection of dsRNA into the adductor muscle; 10 μM calcein was included as a tracer to confirm successful delivery (Supplementary Figure 1). To evaluate dsRNA-mediated gene silencing in zebra mussels, we conducted a series of injection experiments. Initial validation involved injecting dsRNA targeting five distinct genes across 131 mussels. Focusing specifically on the rpn8 gene, we subsequently tested the dose-dependent effects of varying dsRNA concentrations (n = 20), and the temporal duration of the silencing effect (n = 22). To investigate the molecular mechanisms underlying this response, we performed small RNA (sRNA) sequencing on 20 rpn8 dsRNA-injected specimens, which allowed us to correlate sRNA read abundance with RT-qPCR knockdown levels (n = 20). Finally, to assess region-specific targeting efficacy, we injected an additional 31 mussels with rpn8 dsRNA designed to specifically target exon 2, exon 3, or the junction between these two exons.

Following the incubation period, total RNA was isolated using the TRIzol method as described above. Gene silencing efficiency was evaluated by measuring both mRNA and unspliced mRNA levels via one-step RT-qPCR, in which integrated cDNA synthesis and PCR amplification is carried out in a single reaction. RT-qPCR was performed in a final reaction volume of 10.0 μL using the 1-step RT-qPCR kit (Promega, Cat. Number A6121). Each reaction consisted of 5.0 μL of GoTaq master mix, 0.2 μL of GoScript RT mix, 0.01 μL of CRX reference dye, 0.5 μL each of the forward and reverse primers (10 μM stock), 0.5 μL of probe, 2.0 μL of RNA template, and 1.29 μL of nuclease-free water. The thermal cycling conditions consisted of an initial reverse transcription step at 45.0 °C for 15 minutes and polymerase activation at 95.0 °C for 2 minutes, followed by 40 cycles of denaturation at 95.0 °C for 15 seconds and extension at 60.0 °C for 1 minute. A ramp rate of 1.6 °C/s was used for all steps, with fluorescence data collected during the 60.0 °C extension phase. To establish the robust linear dynamic range for each target, standard curves were generated using serial dilutions of total RNA (Supplementary Figure 2 and Supplementary Table 3). These curves were used to identify the optimal RNA input and to account for potential template inhibition. Relative gene expression was quantified using the ΔΔCt method (Schmittgen and Livak 2008), with all samples falling within the established linear, non-inhibitory range for each primer set.

### Analysis of double-stranded RNA (dsRNA)-derived small RNAs

Small RNA libraries from control mussels and *rpn8* dsRNA-injected mussels were sequenced using the Element AVITI platform as described above. The resulting reads were aligned to the *rpn8* locus using Bowtie2 (Langmead and Salzberg 2012), allowing only perfect matches. Size and mapping distribution of aligned reads were subsequently recovered and visualized using custom R scripts.

## Results

### Small noncoding RNA biogenesis and RNAi machinery in zebra mussels

We identified known sRNA biogenesis and processing factors from a number of organisms with well-characterized RNAi systems (*C. elegans*, *D. melanogaster*, *A. thaliana*, and *H. sapiens*) and compared them against their annotated homologs, in the *D. polymorpha* genome. A summary of *D. polymorpha* sRNA biogenesis and processing genes is shown in Figure 1A and Supplementary Table 1. Annotations of a subset of nuclear effectors, sRNA biosynthesis, RISC components, RNAi inhibitors, and dsRNA uptake and spreading factors were found in zebra mussels. Among these comparisons, the repertoire of zebra mussel small RNA and RNAi components was most similar to that found in humans (Jaccard similarity score of 0.59, Figure 1B). Multiple sequence alignment of 47 Dicer 1 and 2 entries revealed that the *D. polymorpha* Dicer-like protein clustered with type 1 Dicer, together with human and bivalve Dicer 1 (Figure 1D). *D. polymorpha* Dicer-like domains were also specifically compared against their homologs for the four model species (Figure 1C). *D. polymorpha* has a single Dicer, and it most resembles human Dicer 1 (43.69% identity). HELIC, Dicer_dimer, PAZ, RIBOc, and DSRM domains were detected in the zebra mussel Dicer-like protein, which also contains mutations in the ATP-binding domain of the helicase motifs I (SSGRM), II (HNCH), and VI (YSHSKGRAR) (Figure 1C). The RNAi proteins Argonaute 2-like and Piwi-like from *D. polymorpha* were also compared against 87 homologs from the Argonaute family. *D. polymorpha* Argonaute 2-like protein indeed clustered together with AGO2 entries, whereas Piwi-like protein from *D. polymorpha* clustered with Piwi-1 proteins (Figure 1D).

**Figure 1.**
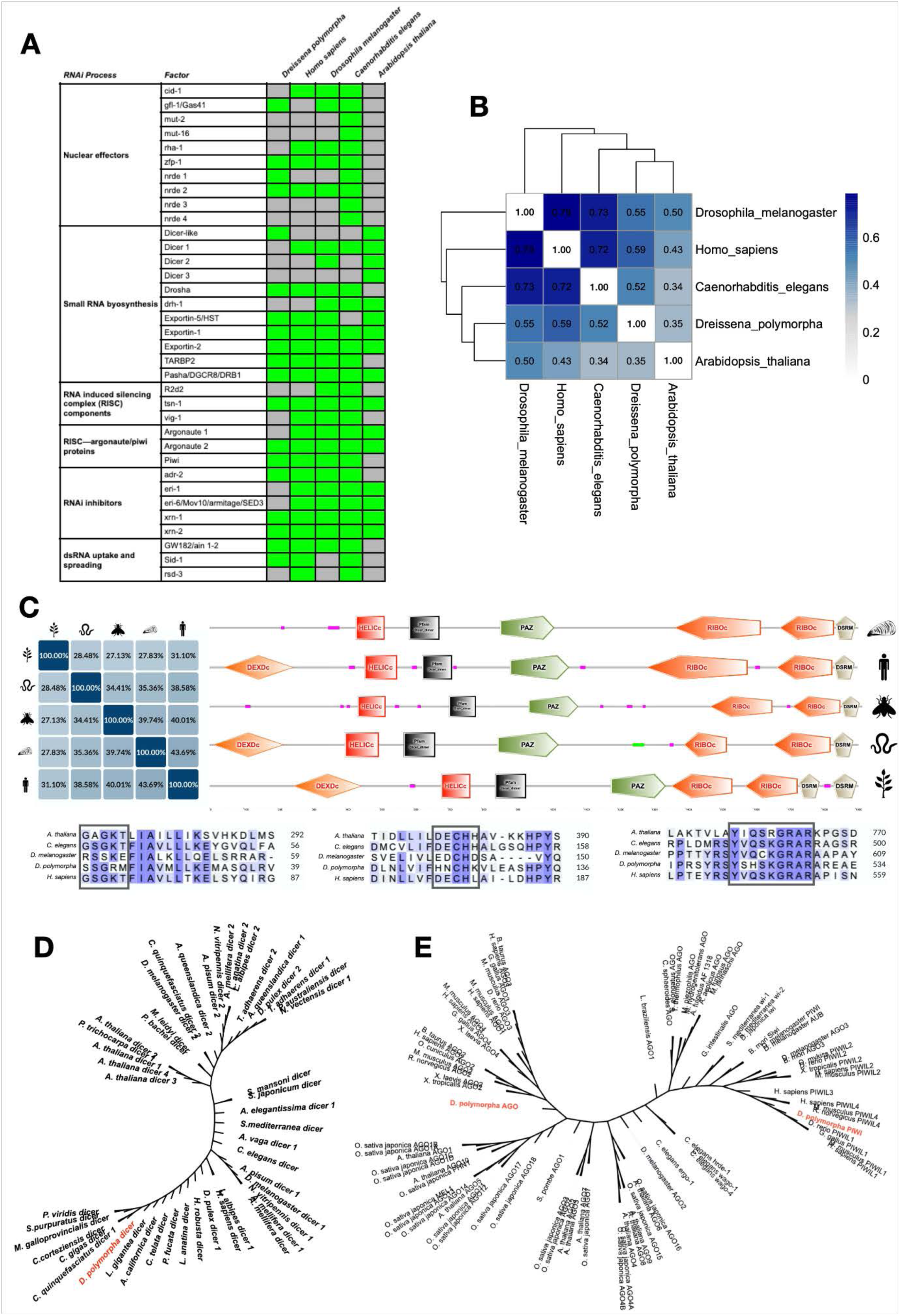
Key RNA biogenesis and RNA interference (RNAi) machinery in zebra mussels. (A) Relevant known sRNA biogenesis and RNAi machinery genes from model species compared with their hypothetical *D. polymorpha* homologs. (B) Jaccard similarity plot for similarities between model species (*Homo sapiens, Drosophila melanogaster, Caenorhabditis elegans*, and *Arabidopsis thaliana*) factors and *D. polymorpha* homologs. (C) Comparison of structural similarity and domain annotations of *D. polymorpha* Dicer-like protein with model species (pink indicates areas of low compositional complexity). Boxed regions for *D. polymorpha* Dicer-like mutations in the ATP-binding domain of the helicase motifs I (SSGRM), II (HNCH), and VI (YSHSKGRAR). (D) Phylogenetic relationship of *D. polymorpha* Dicer-like protein and 47 Dicer 1 and 2 entries from different species. (E) Phylogenetic relationship of *D. polymorpha* Argonaute 2-like protein and Piwi-like protein with 87 Argonaute family entries.

### Size distribution of zebra mussel sncRNA

We sequenced the sRNA from whole animal (n=9) and adductor muscle (n=6) samples. After adapter and quality trimming, and the removal of unmapped reads as well as rRNA and tRNA using the Rfam database, both datasets showed a bimodal read distribution (Figure 2A). Two abundant sRNA size ranges, 19-25 and 26-34 nucleotides in length, were observed for adductor muscle and whole animal samples. The 19–25 nt peak is characteristic of the typical size of miRNAs, while the 26–36 nt peak likely corresponded to piRNA reads. Reads corresponding to these two peaks were isolated for further analysis.

**Figure 2.**
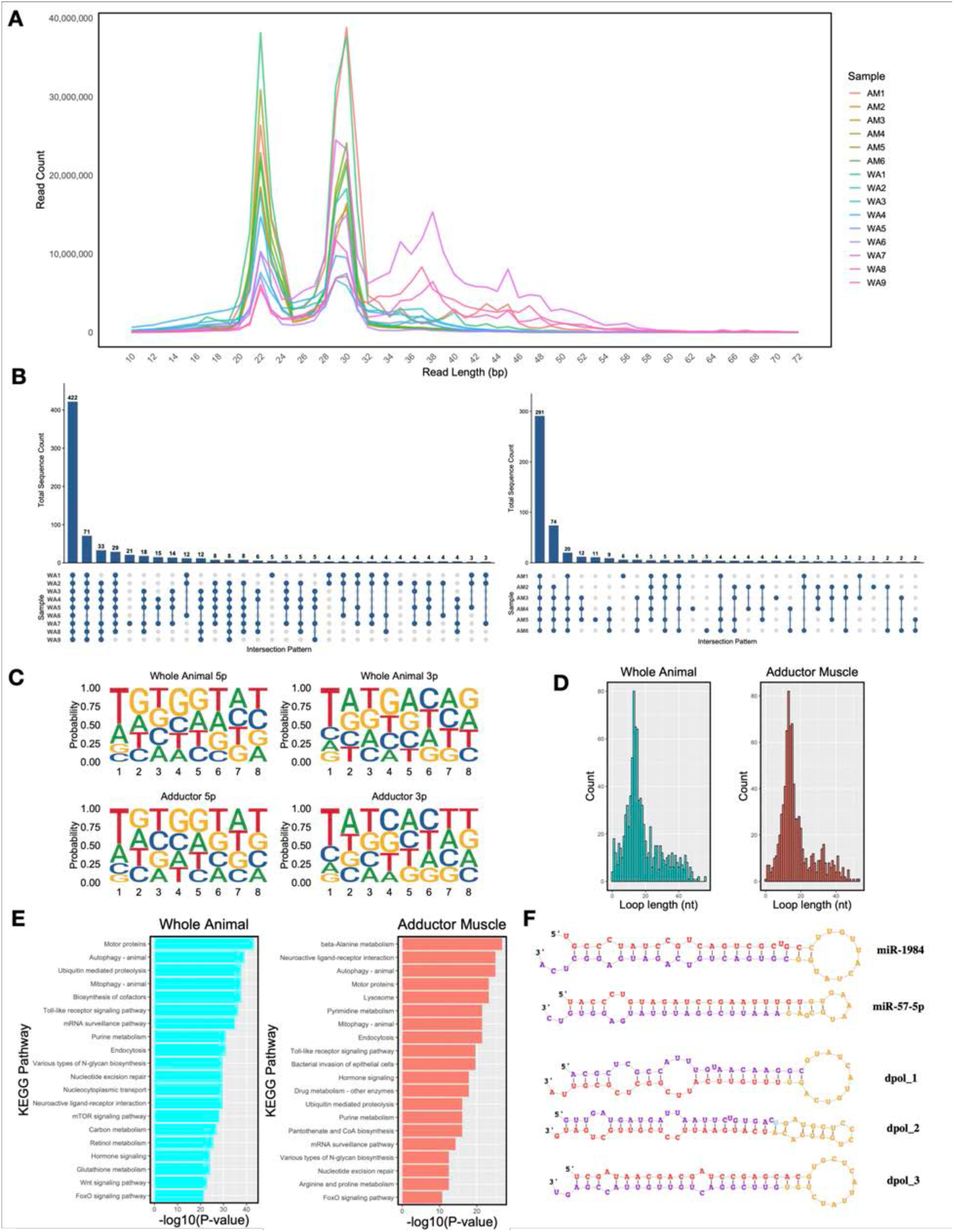
MicroRNAs (miRNAs) in zebra mussels. (A) Bimodal read-size distribution of sRNAs in zebra mussel whole-animal (n=9) and adductor muscle (n=6) samples. (B) Reproducibility of sRNA characterization and conservation of miRNAs across analyzed samples, showing shared mature miRNA sequences after sequence-similarity clustering. (C) Base probability distribution of classified 5′ and 3′ miRNAs across seed sequence positions. (D) Loop-length distribution of miRNA precursor sequences. (E) KEGG pathway enrichment analysis of predicted miRNA targets. (F) Representative structures of detected known and novel miRNAs.

### Zebra mussel microRNAs (miRNAs)

Mature and precursor miRNA sequences were identified using the miRDeep2 algorithm (Friedländer et al. 2012) (Supplementary File 1). miRNA homology sequence annotation using the mature sequence miRBase database (https://www.mirbase.org/, accessed January 2025) were defined as mature sequences with the exact same seed sequence as those from other species. Overall, 422 and 291 putative miRNAs (mirDeep2 Score >10; CPM > 5) were identified in the whole animal and adductor muscle datasets, respectively (Figure 2B). Top five miRNA sequences are displayed in Supplementary Figure 3A. miR-57-5p, miR-25-3p, miR-1984, were among the top fifteen miRNAs identified based on normalized read counts. The miR-57-p sequence has been found to be highly expressed in *C. elegans* subjected to excessive exercise and has been correlated with higher survival and fertility (Xia et al., 2026). In colorectal cancer models, miR-25-3p has been identified as a metastasis promoter that increases vascular permeability and angiogenesis, and it is highly expressed in colorectal cancer patients during the metastatic stage (Z. Zeng et al. 2018). Lastly, evidence suggests that miR-1984 plays a role in the muscle growth of abalone and other mollusks (J. Huang et al. 2018), while also participating in physiological processes like metabolism and the osmotic stress response in *Crassostrea* oysters (X. Zhao et al. 2016).

Examination of miRNA base composition revealed a moderate nucleotide bias for uracil/thymine (U/T) at position 1 in both adductor muscle and whole-animal samples (Supplementary Figure 3B). Representative seed sequences for 3′ and 5′ miRNAs are depicted in Figure 2C. A sequence length range of 13–16 nucleotides was detected as the most abundant loop size in precursor sequences (Figure 2D). In humans, loop size is critical for miRNA processing by Drosha, where the terminal loop required for selective cleavage has been shown to be >10 nucleotides (Zeng et al., 2005).

Putative miRNA targets (Figure 2E; Supplementary File 2) and both novel and known miRNA structures (Figure 2F) were detected. Figure 2E highlights the overrepresented pathways identified via KEGG enrichment analysis of the top 50,000 hits. While thousands of targets were identified, these predictions offer a high-level overview of miRNA-target interactions. The pathways such as “motor proteins”, “beta-Alanine metabolism”, “autophagy”, and “neuroactive ligand-receptor interaction” were among the most represented in this analysis. Finally, we examined miRNA sequences that did not map to the *D. polymorpha* genome. Analysis of these remaining reads against the genome of the mussel’s laboratory diet (*C. reinhardtii*) enabled the detection of putative dietary-derived miRNA sequences (Supplementary File 3). Because these reads were more abundant in the whole-animal samples than in the isolated adductor muscle, it suggests a potential biological uptake from the diet rather than simple surface contamination. However, we acknowledge that we cannot rule out whether these sequences represent true systemic, trans-species sRNAs or localized environmental remnants.

### Zebra mussel Piwi-interacting RNAs (piRNAs)

Abundant piRNAs of characteristic size were detected in both adductor muscle and whole animal samples (Figure 2A). Analysis using the Small RNA Signatures toolkit (Antoniewski 2014) revealed a high Z-score (a measure of standard deviations from the mean) at a 10-base overlap (Figure 3A). piRNA sequences were predicted and clustered using ProTract software (Supplementary File 4). Using the *D. polymorpha* reference genome, a total of 516 and 402 predicted piRNA clusters were defined for the adductor muscle and whole animal sample, respectively. Mono-and bi-directional piRNAs were detected in both datasets (Figure 3B) and piRNA clusters were observed across the genome (Figure 3C). Lollipop plots displaying normalized counts and nucleotide preferences (1T and 10A) for each piRNA cluster are shown in Figure 3D (whole animal) and 3E (adductor muscle). For both samples, a clear, heavy bias for T (U) at position 1 was observed, alongside a variable 10A bias. Lastly, repetitive sequence annotation against the Eukaryota Repbase collection revealed that transposable elements, specifically DNA transposons and LTR retrotransposons, were the most highly represented repeats within the piRNA clusters (Figure 3F and 3G).

**Figure 3.**
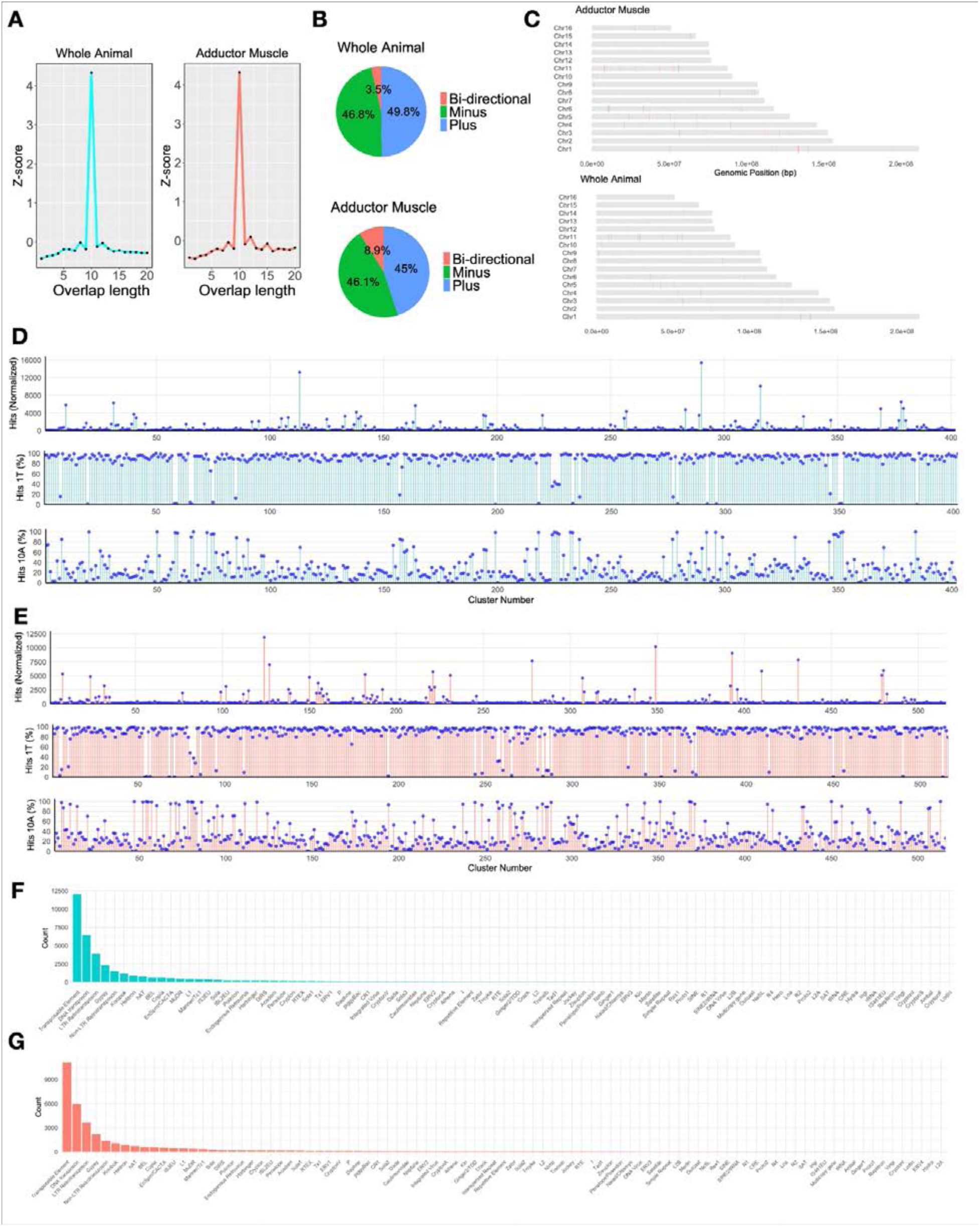
Piwi-interacting RNAs (piRNAs) in zebra mussels. (A) The ping-pong signature showing overlap position. (B) Cluster directionality. (C) Chromosomal distribution of piRNA clusters. (D-E) Lollipop plots for normalized counts and nucleotide preferences (1T and 10A) for each piRNAcluster. Identity of repetitive fragments from whole animal (F) and adductor muscle (G) piRNA cluster sequences.

### Zebra mussel small interfering RNAs (siRNAs)

Cleavage of dsRNA by the Dicer enzyme during endogenous siRNA biogenesis typically produces two structural signatures: 1) duplexes ranging from 21 to 25 nucleotides in length due to RNase III domain activity (Hohjoh 2004), and 2) a 2-3 nucleotide overhang at the 3’ end of the reference strand and a blunt or up to 3-nucleotide overhang at the 5’ end of the reference strand (Murcott et al. 2022). We identified all potential perfectly matched duplexes within the 18-34 nucleotide length range using the “Small RNA signatures” toolkit from ARTbio (Antoniewski 2014). Figure 4A and 4B show the Z-scores of predicted duplexes for whole animal and adductor mussel reads, respectively. For the whole animal, sequence reads of 22 and 26 nucleotides had the highest Z-scores, and those of 22 and 25 had the highest Z-scores in the adductor mussel sample. Overall, the Z-scores in both samples are close to the average (Z-score of 0) or only one standard deviation above (Z-score of 1), indicating no strong abundance of characteristic endogenous siRNA duplex length. Analysis of the predicted siRNA structures (Figure 4C) revealed that a very low percentage of the duplexes exhibited a 3’ overhang combined with a 5’ blunt end (0.85% and 0.78% for the whole animal and adductor muscle, respectively). In addition, a higher percentage of sequences displayed 3’ overhangs on both ends (∼40% of the predicted duplexes for both datasets). Although these two characteristic Dicer processing marks were detectable, the absence of abundant typical siRNA duplex length suggests that while this pathway is present, it may be underrepresented.

**Figure 4.**
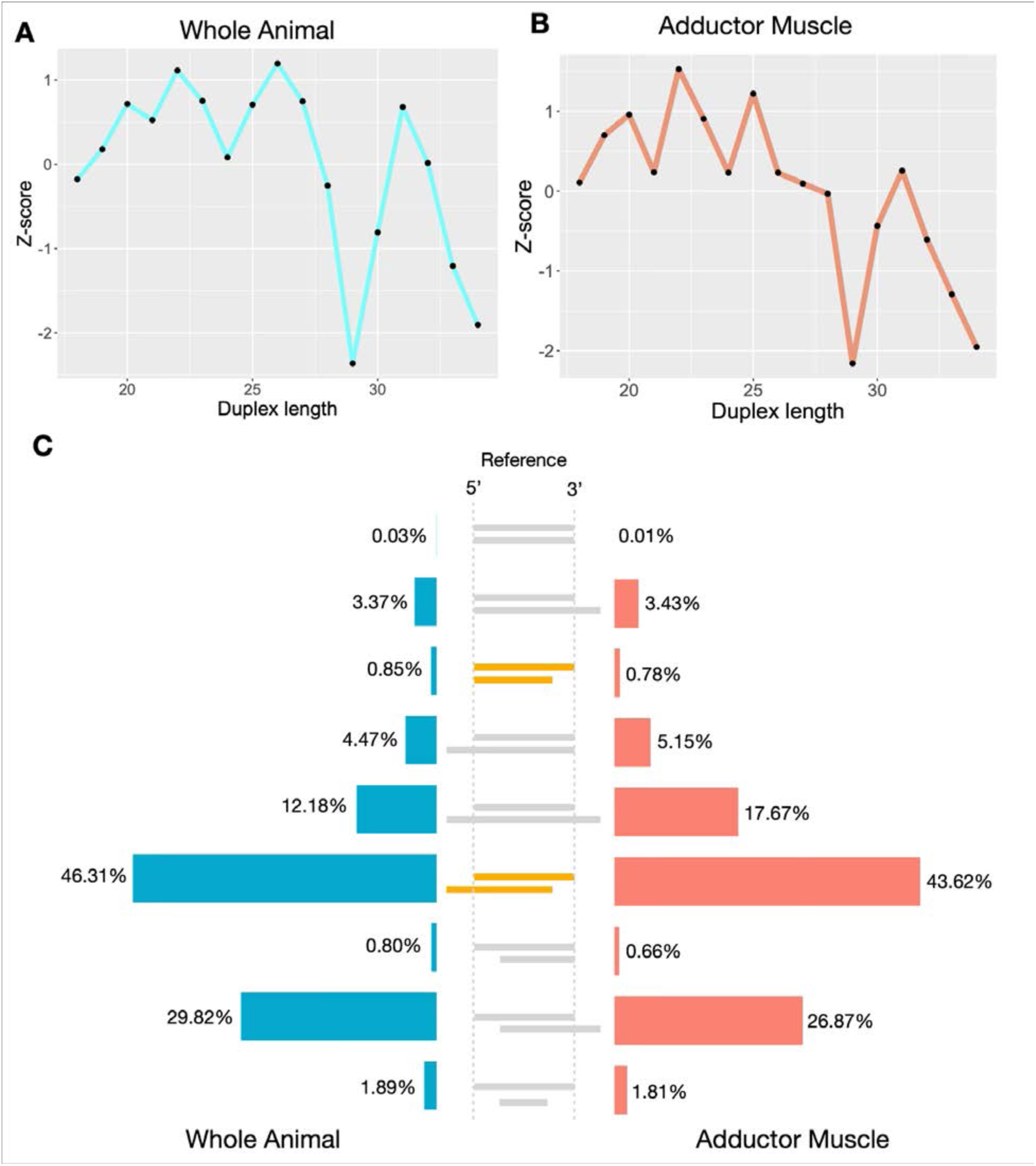
Small interfering RNA (siRNA) profiles in the zebra mussel. Distribution of potential perfectly matched duplexes within the 18–34 nucleotide length range for the (A) whole-animal and (B) adductor muscle datasets. (C) Overhang type classification of predicted siRNAs, with the canonical Dicer signature highlighted in orange.

### RNAi-mediated gene silencing and processing of exogenous dsRNA

We designed five dsRNA constructs targeting putative essential genes from the *D. polymorpha* genome (Figure 5A). 0.1 μg of dsRNA was injected into the adductor muscle, and the degree of gene silencing was assessed 24 h post-injection for mRNA and unspliced pre-mRNA. Two of the five tested constructs, dsRNA targeting *rpabc5* and *rpn8* genes, showed significant knockdown of both mRNA and unspliced pre-mRNA (p < 0.05) (Figure 5B). Additional injection experiments were carried out with 0.1, 1.0, and 2.0 ug of *rpn8* dsRNA. There was a significant decrease (p < 0.05) in *rpn8* gene expression across the three different dsRNA concentrations, with no clear dose-dependent effect observed within the concentration range tested (Figure 5C). In addition, we tested the silencing effect of 2.0 μg of *rpn8* dsRNA at 24, 48, and 72 h post-injection. Knockdown of *rpn8* was observed at 24 h (p < 0.05) and gradually recovered towards baseline during the 48 h and 72 h post-injection timepoints (Figure 5C). Quantification of *rpn8* and three additional target genes (*rpabc5, cpeb,* and *e1ubi*) in rpn8 dsRNA-injected samples confirmed that gene silencing was specific to the construct targeting *rpn8* mRNA (Supplementary Figure 4).

**Figure 5.**
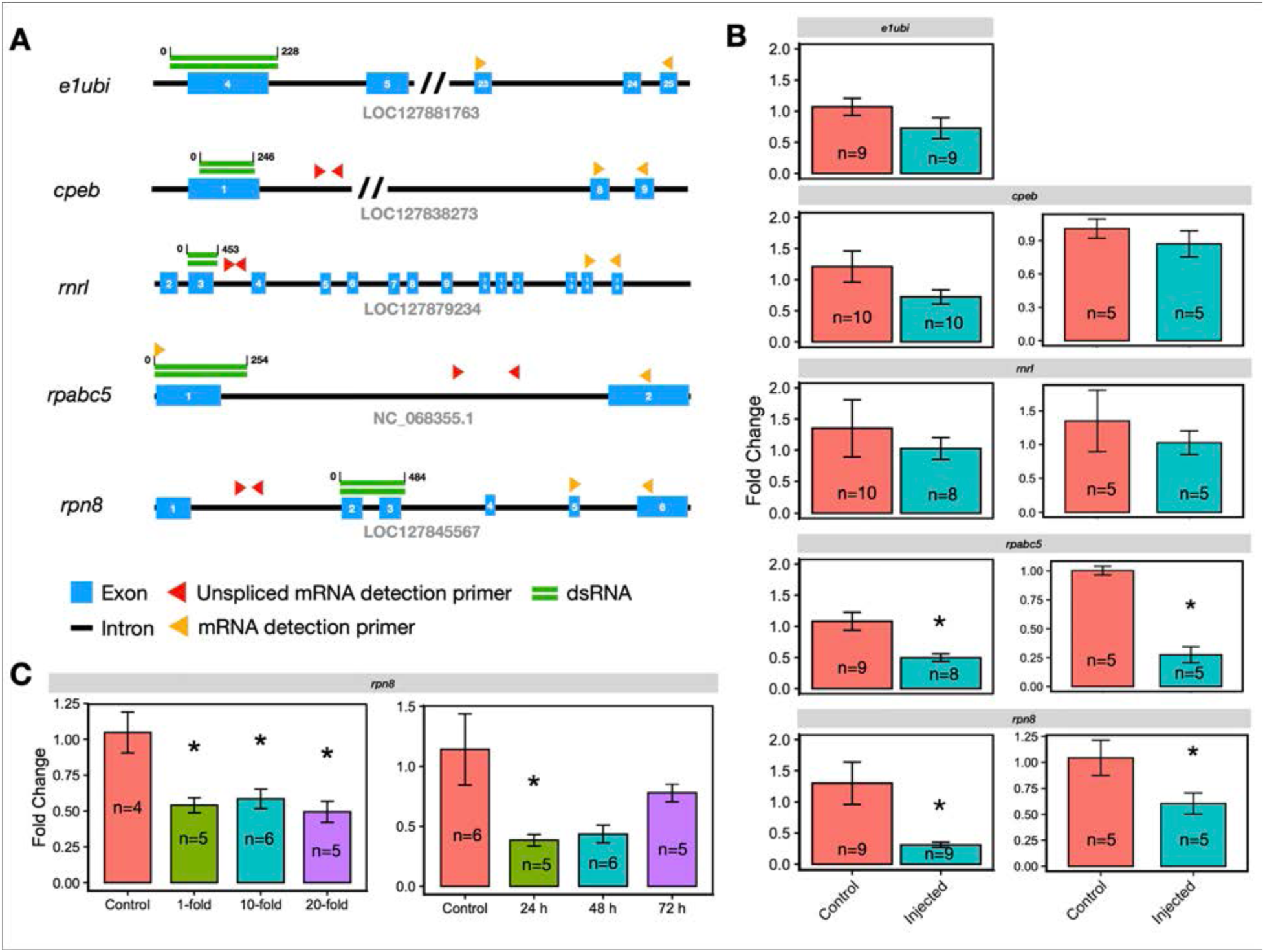
RNA interference (RNAi) knockdown efficiency. (A) dsRNA constructs designs and detection regions for mRNA and unspliced pre-mRNA. (B) Screening of gene expression levels after injection with five different dsRNA constructs (0.1 μg) targeting mRNA (left column) and unspliced pre-mRNA (right column). Knockdown was assessed at 24 hours post-injection (n=131). Data are presented as mean ± SEM. Asterisks indicate a significant difference compared to the control group (p<0.05). (C) Effect of construct amount on knocking down of the *rpn8* gene. Gene expression was measured 24 hours post-injection (left) across three different dsRNA concentrations (0.1 μg, 1.0 μg, and 2.0 μg; n=20) and duration of *rpn8* gene silencing (right) following a single injection of 2.0 μg of the *rpn8* dsRNA construct (n=22). Gene expression was assessed at 24 h, 48 h, and 72 h post-injection.

Next, we utilized sRNA sequencing to analyze the processing of injected *rpn8* dsRNA. sRNA reads from injected samples were mapped against the *rpn8* reference sequence corresponding to the dsRNA construct. Based on the size distribution of mapped reads, we determined that this construct is being processed, generating ≥ 19-nucleotide-long sRNAs (Supplementary Figure 5). A dose-dependent increase in sRNA mapping to this region was observed in injected samples (Figure 6A). Mirroring the kinetics of *rpn8* knockdown, a dose of 2.0 μg leads to a dramatic increase in mapped reads at 24 h and recovers to near baseline levels at 72 h. We observed a correlation between the extent of *rpn8* knockdown and the quantity of normalized sRNA reads mapping to the *rpn8* dsRNA region (Figure 6B).

**Figure 6.**
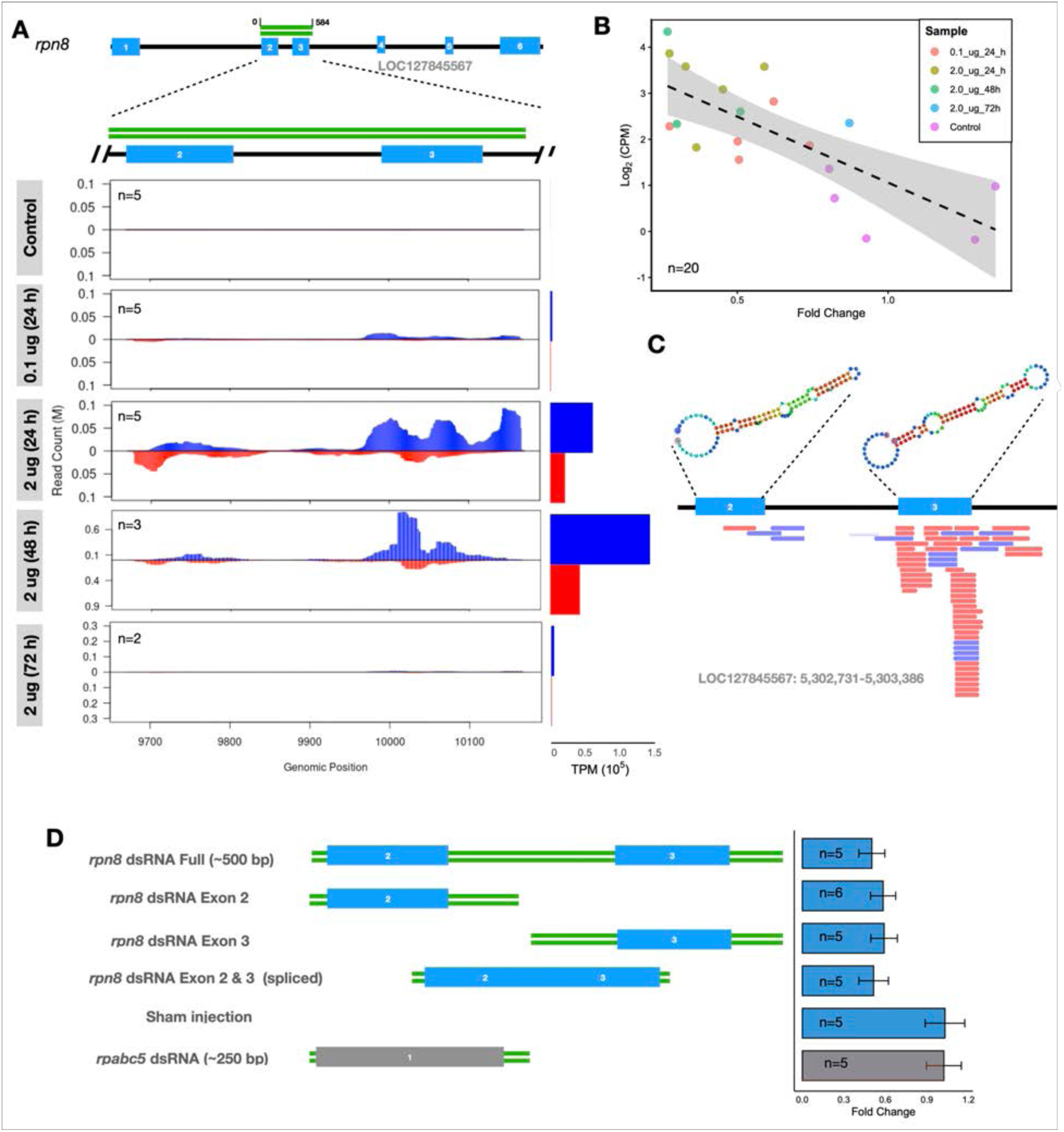
Correlation of sRNA abundance and extent of knockdown. (A) Coverage plot of sRNA reads across the *rpn8* target locus (n=20). (B) Correlation between *rpn8* knockdown and the abundance of sRNA reads mapped to the *rpn8* dsRNA reference (R^2^ = 0.548; n=20). (C) *rpn8* structures which correspond to sites of endogenous sRNA production. (D) Indicated dsRNA constructs were injected into the adductor muscle and mRNA levels for RPN8 gene were measured 24 h post injection (n=31).

We found that *rpn8* pre-mRNA contains secondary structural motifs that also align with sites of endogenous sRNA production, suggesting that dsRNA silencing of this gene may be at least partially tapping into endogenous miRNA or piRNA-based gene regulatory mechanisms (Figure 6C). To further define the dsRNA regions responsible for silencing the *rpn8* gene, we tested portions of the dsRNA construct targeting exon 2, exon 3, and the junction of these two exons. Each of these sub-regions of the *rpn8* dsRNA sequence was sufficient to reduce *rpn8* expression to levels comparable to the full length *rpn8* dsRNA (Figure 6D). No knockdown of *rpn8* expression was observed in a sham injection or with injection of dsRNA targeting a different gene (*rpabc5*).

## Discussion

The zebra mussel is a highly damaging invasive species throughout Europe and North America. While there has been substantial work to understand zebra mussel biology and ecology (Nalepa and Schloesser 1992; Ram and McMahon 1996; Mackie 1992; Mackie and Schloesser 1996; Vanderbush et al. 2021), and efforts to address its impact and control are ongoing (Hernández Elizárraga et al. 2023), development of molecular and genetic tools and a fundamental understanding of the zebra mussel’s molecular regulatory mechanisms are needed. Here we characterize the sRNA and RNAi pathways in zebra mussels. We analyzed homologs of sRNA biogenesis and processing genes in the zebra mussel genome and utilized small RNA sequencing to gain insights into these gene regulatory systems. We also demonstrated that knockdown of gene expression can be achieved through injection of dsRNA into zebra mussel tissues, establishing a methodology for targeted genetic manipulation in zebra mussels.

Since its discovery in *C. elegans* (Fire et al. 1998), RNAi has provided an approach for modulating gene expression in a wide range of organisms without the need for transgenesis. This is especially advantageous for organisms, such as zebra mussels, that remain difficult or impossible to culture over multiple generations in a laboratory setting. Induction of techniques such as RNAi through - feeding of microbes producing dsRNA have proven highly effective in other systems (Kaur et al. 2020; Kunte et al. 2020; Kwon et al. 2013; N. Yu et al. 2013). Our tentative observation that miRNAs derived from dietary C. reinhardtii can be detected within the mussel provides also a proof of concept for using algae as a delivery vehicle for RNAi in zebra mussels.

RNAi pathways have been extensively characterized in many species and also used for applications in areas such as targeted medicine (Barik 2005) and pest species biocontrol (De Schutter et al. 2022). We identified a number of key sRNA biogenesis elements and RNAi machinery components in the zebra mussel genome (Figure 1A). Among the selected genes screened, the zebra mussel RNAi factors most closely resemble the human RNAi system (Figure 1B). Phylogenetic analysis of 47 Dicer enzymes clustered the *D. polymorpha* Dicer within the type 1 group. Similar to humans, zebra mussels do not possess a Dicer 2 ortholog (Figure 1C). In humans, the canonical processing of primary miRNAs is initiated by DGCR8 and the RNase III DROSHA in the nucleus in a process resulting in ∼ 70 nt precursor miRNAs (stem-loop), which are subsequently transported to the cytoplasm by Exportin-5. In the cytoplasm, DICER cleaves the pre-miRNA loop hairpin, generating miRNA duplexes of about 22 nt with a 2 nt overhang at the 3′ end (Vergani-Junior et al. 2021). We found zebra mussel sequence homologs to DGCR8, DROSHA, and Exportin-5 (Figure 1A). In addition, sequencing of zebra mussel sRNA revealed typical miRNA-size (19-25 nt) and piRNA-size (26-36 nt) reads (Figure 2A). We examined the protein sequence of the zebra mussel Dicer 1 and Argonaute protein homologs. Interestingly, *D. polymorpha* has most of the human Dicer domains (e.g., HELIC, Dicer_dimer, PAZ, RIBOc), but differences were also identified, such as the mutations in the ATP-binding domain of the helicase motifs I (SSGRM), II (HNCH), and VI (YSHSKGRAR) (Figure 1D and 1E). Helicase domain mutations can affect Dicer activity. For instance, Drosophila Dicer-1 has comparable helicase domain sequence to zebra mussel Dicer 1 (Figure 1E) and it only processes pre-miRNA (Lee et al. 2004). Introducing similar mutations in Drosophila Dicer-2 can disrupt siRNA processing (Welker et al. 2011). It has been shown in Drosophila that helicase mutants favor terminal siRNA generation (Lee et al. 2004), so it is possible that *D. polymorpha* Dicer 1 is non-processive and thus preferentially generates terminal siRNAs.

The RNA-Induced Silencing Complex (RISC) Argonaute (AGOs) proteins were also investigated. The two *D. polymorpha* AGO-like sequences, Argonaute2 and Piwi1, clustered with their respective homologs in humans and other species. Specifically, *D. polymorpha* Argonaute2 showed the highest similarity to *Xenopus* AGO2, a miRNA-specific AGO, whereas Piwi1 most closely resembled *Danio* PIWIL1 (Figure 1F). The presence of miRNA-specific AGO homologs in the zebra mussel (miAGO) aligns with broader findings in mollusks; all species studied thus far possess at least one AGO copy, with some showing group-specific expansions, though they generally lack siRNA-specific AGOs (siAGOs) (Von Eiff et al. 2025).

Based on sRNA sequencing analysis, we were able to identify miRNA and piRNA-sized sRNA in whole animal and adductor muscle samples (Figure 2A). The presence of abundant piRNA-sized sequences in adductor muscle tissue supports the hypothesis that somatic piRNAs are ubiquitously expressed in mollusks (Jehn et al. 2018).

*D. polymorpha* miRNAs were found to be 19–25 nucleotides long (Figure 2A) and did not display nucleotide bias, except for U/T at position 1 (Supplementary Figure 3B). Additionally, we found that a number of identical seed sequences (around 80%) from position 2 to 8 have been previously reported in other species (Supplementary File 1). Consequently, the vast majority of these putative miRNAs were classified as “known” miRNAs that have homologs previously identified in other species, substantiating that what we are seeing are likely true regulatory sRNAs as opposed to random fragments. The nucleotide range of 13-16 was detected as the most abundant loop size for precursors (Figure 2D). Understanding the structure composition of these endogenous miRNA molecules may permit the rational design of improved synthetic RNAi triggers (Fowler et al. 2016). We identified enriched pathways for potential miRNA targets, encompassing a range of overrepresented cellular processes (Figure 2E). The structural features of miRNA hairpins (Figure 2F) in zebra mussels, along with miRNA target prediction and enrichment analysis provide a global snapshot of miRNA function in zebra mussels and highlight specific genes and pathways that could be exploited to develop tools for genetic manipulation or biocontrol.

Likewise, piRNAs between 26–34 nucleotides in length were identified in zebra mussels. These sequences exhibited the same 5’ terminal uridine (1U) preference (Figure 3D-E) characteristic of other mollusk piRNAs (Jehn et al. 2018), as well as the 10-nucleotide overlap signature characteristic of ping-pong amplification (Figure 3A). Model organisms display a heavy bias toward adenosine at the 10th position (10A) (Stein et al. 2019) of germ cell but not somatic piRNA (Malone et al. 2009). Such nucleotide preferences emerge from the specific interactions between piRNAs and their Piwi-clade partners: while Piwi and Aub typically load sequences defined by a 5’ uracil (1U), Ago3-associated piRNAs generally lack this 5’ signature, instead exhibiting a distinct 10A enrichment (Czech and Hannon 2016). Consistent with their canonical role in transposon silencing, genomic piRNA-producing loci, or piRNA clusters, were detected in the zebra mussel genome. Repetitive sequence annotation against the Eukaryota Repbase collection revealed that transposable elements, specifically DNA transposons and LTR retrotransposons, were the most highly represented repeats within these clusters (Figures 3F and 3G). Most of the detected zebra mussel piRNA clusters were monodirectional (Figure 3B) and scattered across the genome (Figure 3C). These clustered sequences are highly divergent across taxa and are responsible for modulating piRNA populations in different evolutionary lineages (Rosenkranz et al. 2021). While no strong nucleotide preference was observed beyond the first position in zebra mussels, a subset of individual piRNA clusters were enriched for either 10A or both 1U and 10A (Figure 3 D-E).

We also identified perfectly matched duplex sequences that produced sRNA reads, consistent with endogenous siRNA (endo-siRNA). Generation of siRNA duplexes by Dicer enzyme is expected to have a strong 19 nt-overlap signature (Antoniewski 2014). Although endo-siRNA duplex-like structures can be predicted from the analyzed datasets, no strong abundance of a dominating duplex length was found for the whole animal and adductor muscle samples (Figure 4A and 4B), where potential duplexes in the 18-34 nucleotide range displayed mostly Z-score values close to the average or only one standard deviation above. Given our inability to detect strong endo-siRNA-like small RNA signatures and the absence of Dicer 2 proteins in the zebra mussel genome, this may preclude a functional endo-siRNA pathway for this organism, as observed in other bivalve species (Von Eiff et al. 2025). These results also align with previous analyses of small RNA (sRNA) biogenesis pathways in other mollusks, which have shown that some species lack functional siRNA pathways (Von Eiff et al. 2025).

The results of our initial screening identified dsRNA targeting *rpabc5* and *rpn8* as functional RNAi triggers in *D. polymorpha*, demonstrating that RNAi can be delivered through injection into the adductor muscle tissue (Figure 5B). For these two target genes, lower levels of unspliced and spliced mRNA were detected (Figure 5B). While previous research has described Ago protein/miRNA and PIWI protein/piRNA functions in mRNA degradation or translational inhibition, these pathways can also employ epigenetic mechanisms including DNA and RNA methylation, deadenylation, and phosphorylation (Zhang et al. 2023). If recruitment of epigenetic modification factors is occurring in the zebra mussel, this might explain the decreased abundance of unspliced mRNA for *rpn8* and *rpabc5*, as repression may be occurring at the transcriptional level.

Further experiments using *rpn8* dsRNA confirmed persistent gene knockdown at various concentrations, lacking a strong dose-dependent effect in the tested range (Figure 5C). This suggests that the cellular RNAi machinery may already be saturated at the lowest concentration (0.1 μg). The observed silencing effect was significant at 24 hours, beginning to recover by 48 hours (Figure 5C). Analysis via sRNA sequencing provided molecular validation for the knockdown, showing a direct correlation between the extent of silencing and the abundance of processed small RNA reads (Figure 6B). The observation that the dsRNA-targeted region fortuitously corresponds to putative endogenous sRNA production sites (Figure 6C) may provide insights into the mechanism of silencing, which could be via miRNA or piRNA pathways. Our results demonstrate that dsRNA targeting either half of the *rpn8* dsRNA construct or the exon-exon junction is sufficient to reduce *rpn8* gene expression by approximately 40% (Figure 6D). Furthermore, the absence of an effect on *rpn8* mRNA levels following sham or *rpabc5* dsRNA injections confirms the sequence-specificity of this effect. Further optimization and assessment of additional targets, such as *rpabc5* will shed more light on the mechanism of dsRNA-triggered silencing in zebra mussels.

This study provides an initial characterization of the sRNAbiogenesis machinery and RNAi pathways in the invasive zebra mussel. By integrating bioinformatic analysis, sRNA sequencing, and functional knockdown assays, we describe the features of endogenous sRNA in zebra mussels and demonstrate that despite the lack of a Dicer 2 homolog, they remain susceptible to exogenous dsRNA-mediated silencing. These insights into the species-specific processing of regulatory RNAs provide a critical prerequisite for developing programmable genetic manipulation tools and help lay the groundwork for RNAi-based biocontrol strategies to manage this pervasive invasive species.

## Data availability statement

The datasets generated and analyzed during the current study are available in the NCBI BioProject repository under accession number PRJNA1467755.

## Acknowledgements

We thank Krista Espelien, Benjamin Minerich, and John Gerritsen for collecting the zebra mussel specimens. Thanks also to Michael McCartney, Thea Edwards, Cathy Richter, Katy Klymus, and Edward Large for helpful discussions. We also thank Cori Mattke and Jay Maher from the Minnesota Aquatic Invasive Species Center (MAISRC) for administrative and facility support. We thank our colleagues in the University of Minnesota Genomics Center (RRID:SCR_012413), as well as the entire UMGC Innovation Lab staff. The authors acknowledge the Minnesota Supercomputing Institute (MSI) at the University of Minnesota for providing resources that contributed to the research results reported within this paper. URL: http://www.msi.umn.edu.

## Author contributions

VHHE: Conceptualization; Data curation; Visualization; Writing – original draft

LGO: Writing – reviewing & editing; Investigation

SB: Conceptualization; Writing – reviewing & editing; Investigation

DMG: Conceptualization; Writing – reviewing & editing; Supervision; Project Administration

## Supplementary Files

Supplemental File 1. Mature and precursor miRNA sequences identified using the miRDeep2 algorithm.

Supplementary File 2. Predicted miRNA targets.

Supplementary File 3. Trans-species miRNA detected for both adductor muscle and whole animal samples.

Supplementary File 4. Predicted piRNA sequences with ProTract software.

## Supplementary Figures

**Supplementary Figure 1.**
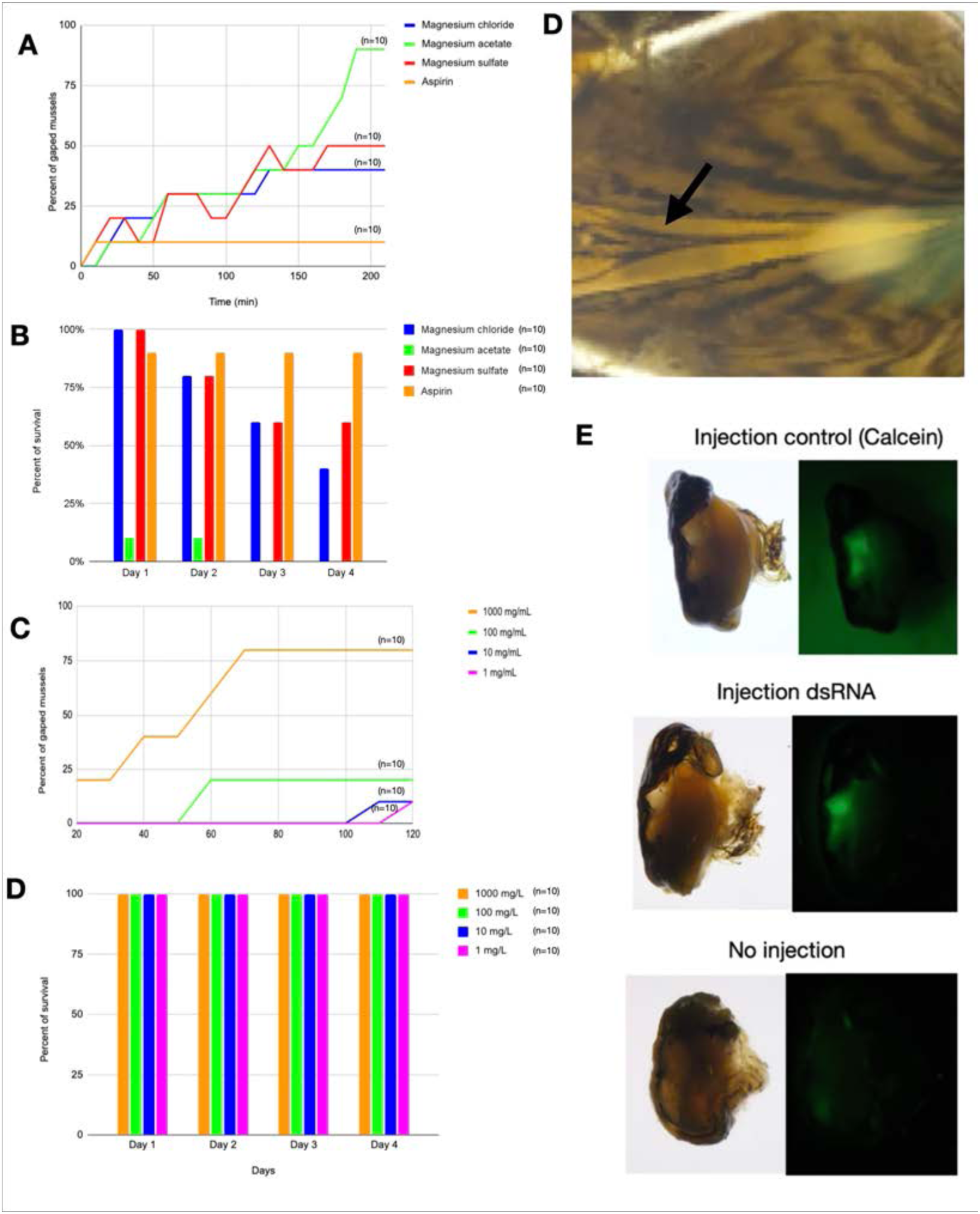
Sedation agents evaluated for zebra mussel nucleic acid injection. A) Percentage of gaped mussels following treatment with Mg^2+^ salts or acetylsalicylic acid. B) Mussel survival rates following sedation with Mg^2+^ salts or acetylsalicylic acid. C) Percentage of anesthetized mussels and (D) percent of survival across different concentrations of MS-222. D) Representative photograph of a relaxed zebra mussel (injection site highlighted with black arrow). E) Calcein fluorescence confirms successful injection into the adductor muscle tissue.

**Supplementary Figure 2.**
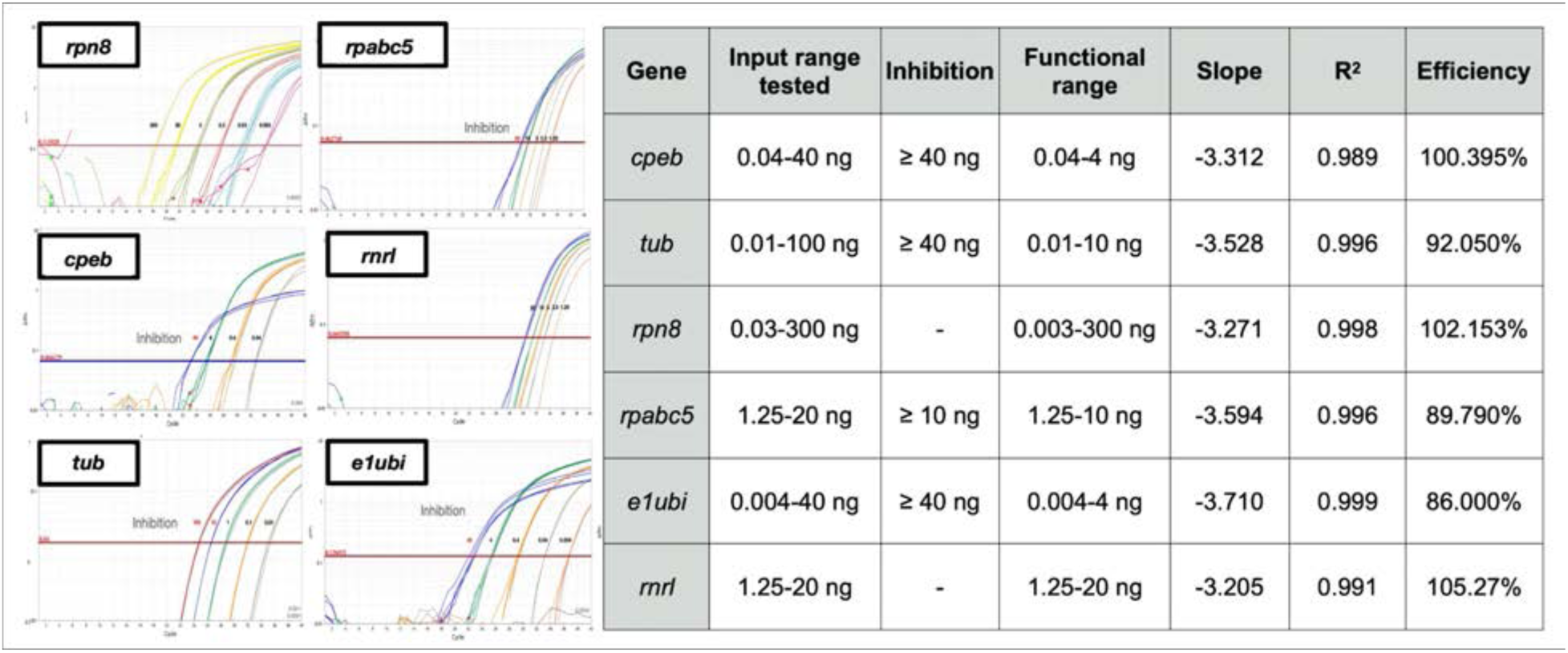
Validation of One-Step RT-qPCR Assays. Representative amplification plots (left) and performance metrics (right) for six target genes (*rpn8, rpabc5, cpeb, rnrl, tub, and e1ubi*). Standard curves were generated using serial dilutions of total RNA to determine the functional range, amplification efficiency, and R^2^ values. “Inhibition” labels indicate the concentration thresholds where template excess resulted in non-linear amplification. These parameters were utilized to standardize RNA input and ensure accurate relative quantification via the ΔΔCt method.

**Supplementary Figure 3.**
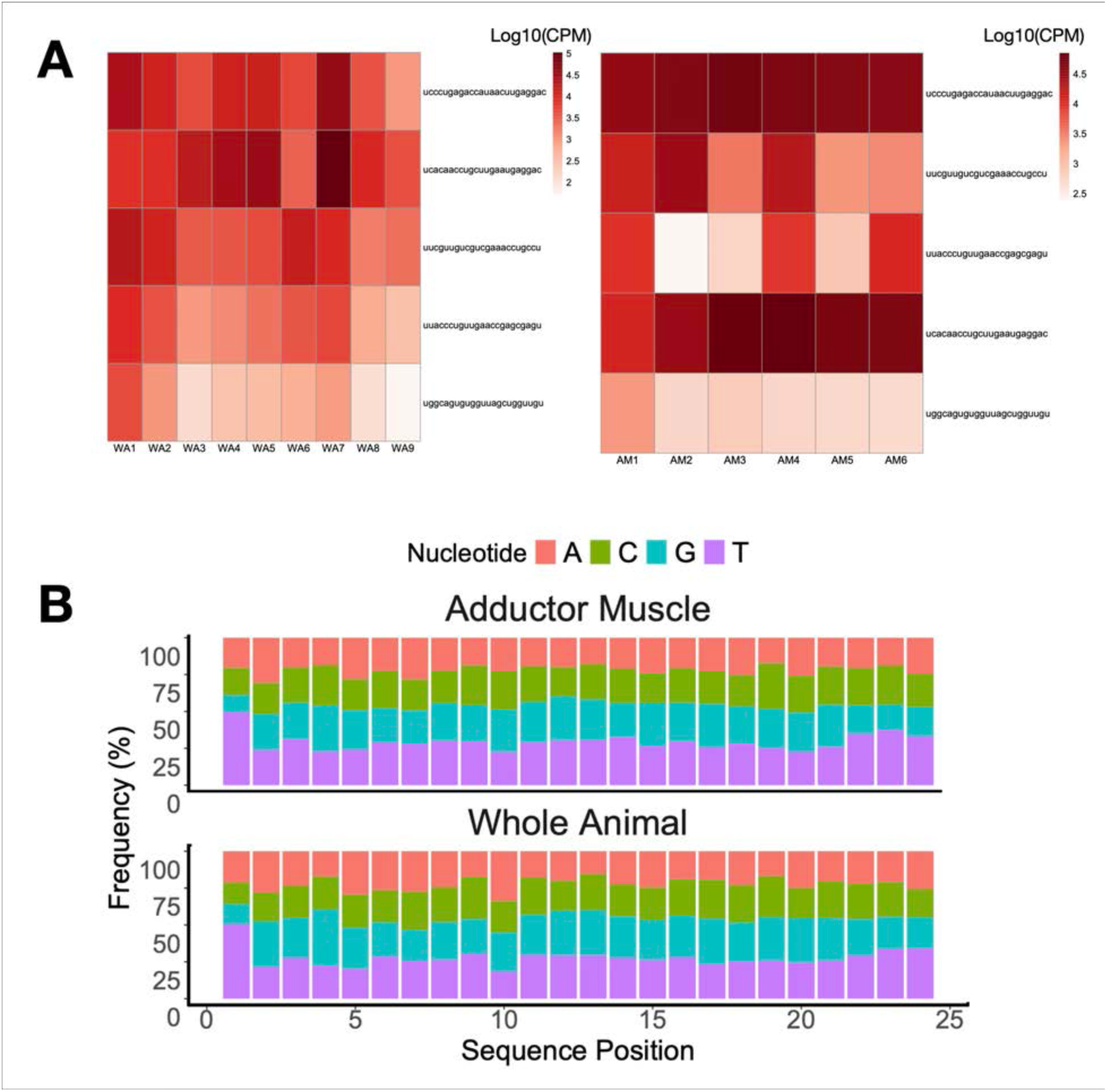
(A) Top5 mature miRNA sequences expressed in whole animal (WA) and adductor muscle (AM) samples. B) Base composition of mature miRNA sequences.

**Supplementary Figure 4.**
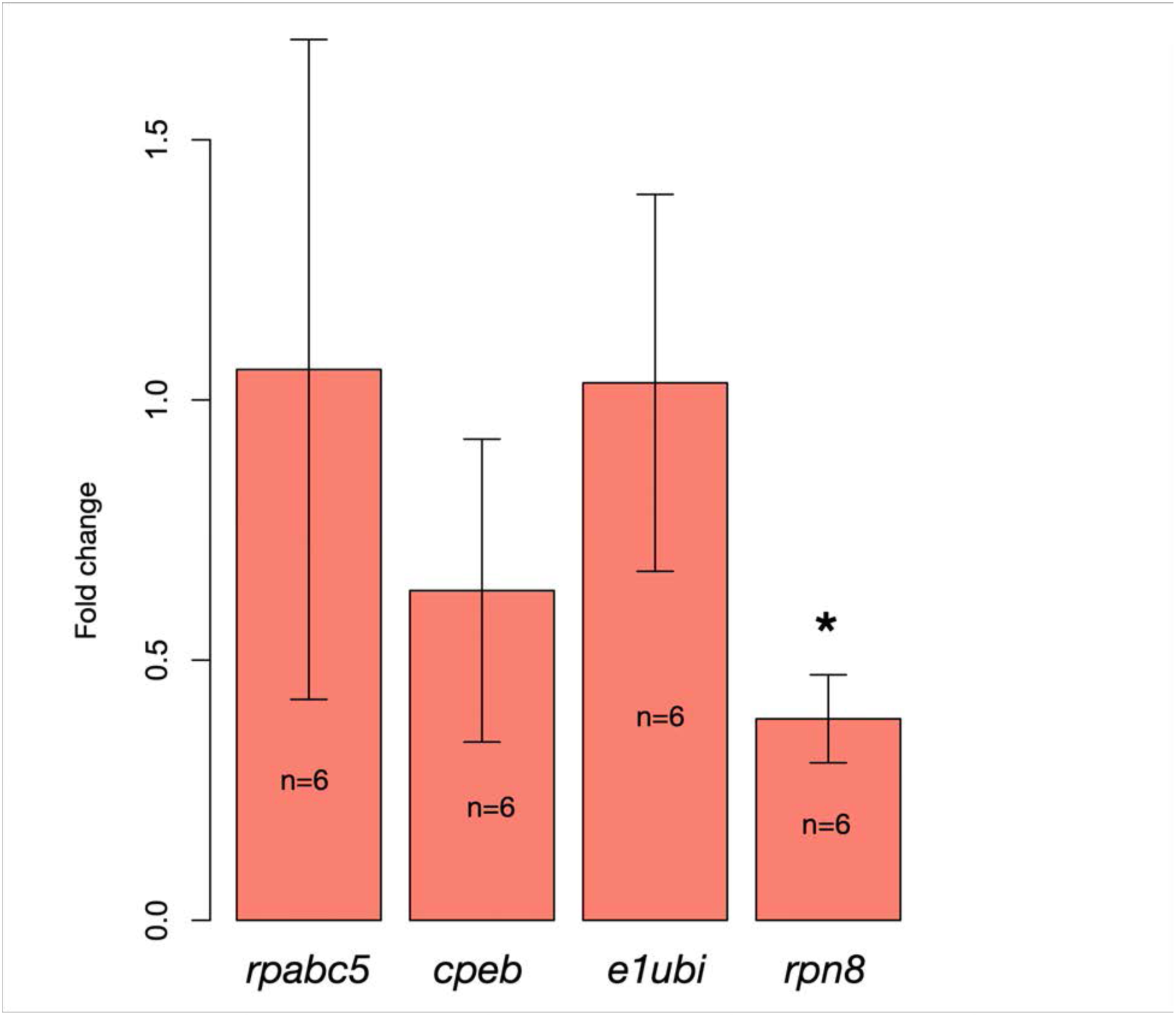
Expression profiles of target genes in *rpn8* dsRNA-injected zebra mussels.

**Supplementary Figure 5.**
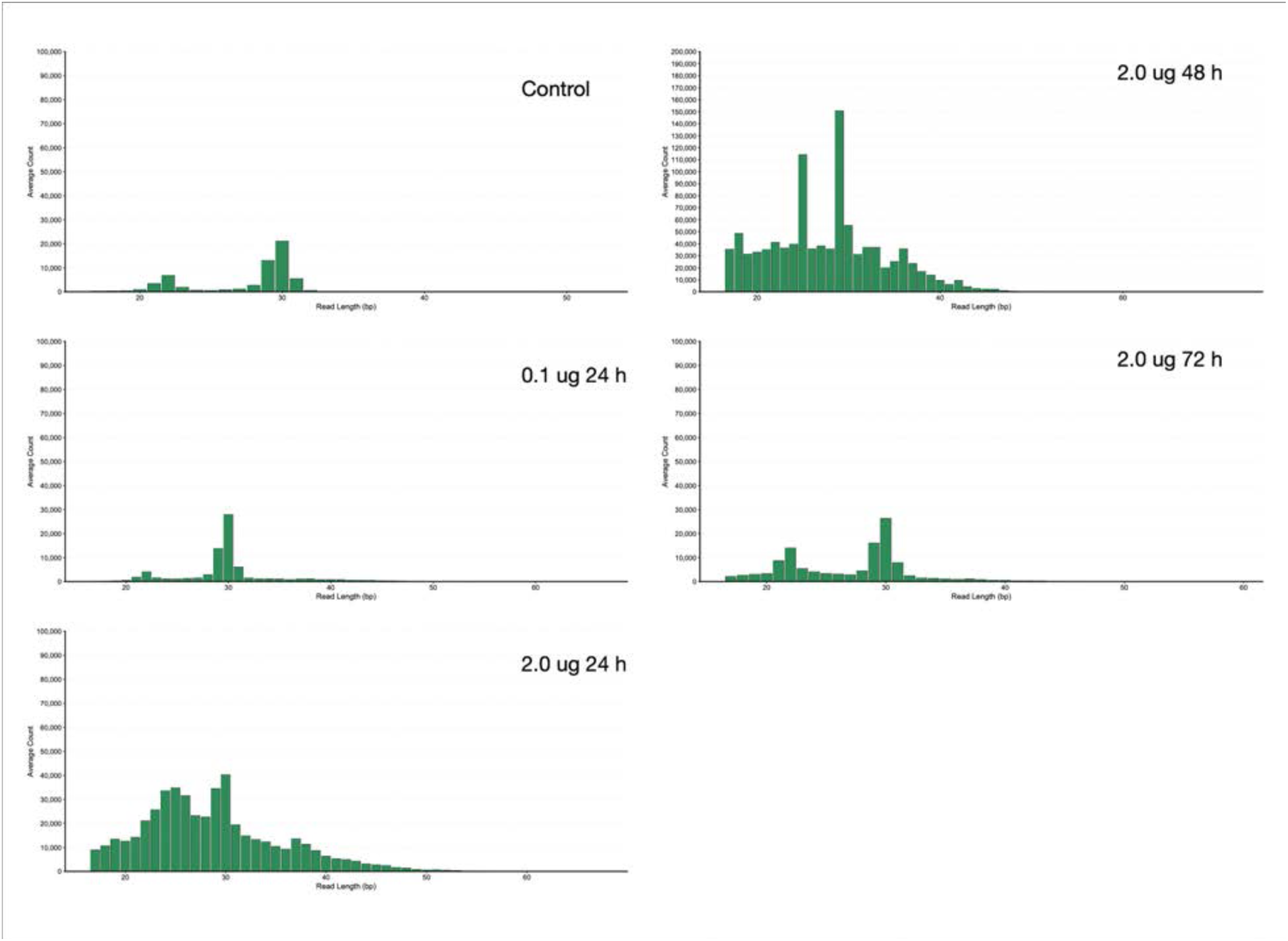
Representative read length distribution of mapped reads across *rpn8* target region.

## Supplementary Tables

**Supplementary Table 1.** Summary of key sRNA biogenesis and processing genes.

|  | *Locus tag/Accession* |  |  |  |  |
| --- | --- | --- | --- | --- | --- |
| **Factor** | *Dreissena polymorpha* | *Homo sapiens* | *Drosophila melanogaster* | *Caenorhabditis elegans* | *Arabidopsis thaliana* |
| cid-1 | -- | CENP-A | Dmel_CG13329 | CELE_K10D2.3 | -- |
| gfl-1/Gas41 | LOC127881330 | -- | Dmel_CG9207 | CELE_H04M03.4 | -- |
| mut-2 | -- | -- | -- | CELE_K04F10.6 | -- |
| mut-16 | -- | -- | -- | CELE_B0379.3 | -- |
| rha-1 | -- | DDX9 | Dmel_CG11908 | CELE_T07D4.3 | -- |
| zfp-1 | LOC127855600 | MLLT10 | Dmel_CG1070 | CELE_F54F2.2 | -- |
| nrde 1 | KAH3844401.1 | -- | -- | CELE_C14B1.6 | -- |
| nrde 2 | XP_052272144.1 | NRDE2 | Dmel_CG5877, | CELE_T01E8.5 | -- |
| nrde 3 | -- | -- | -- | CELE_R04A9.2 | -- |
| nrde 4 | -- | -- | -- | CELE_F45E4.10 | -- |
| Dicer-like | LOC127873210 | -- | -- | -- | AT1G23790 |
| Dicer 1 | -- | DCR1 | Dmel_CG4792 | CELE_K12H4.8 | AT1G01040 |
| Dicer 2 | -- | -- | Dmel_CG6493 | -- | AT3G03300 |
| Dicer 3 | -- | -- | -- | -- | AT3G43920 |
| Drosha | LOC127838009 | DROSHA | Dmel_CG8730 | CELE_F26E4.10 | -- |
| drh-1 | -- | -- | Dmel_CG3697 | CELE_F15B10.2 | AT3G01540 |
| Exportin-5/HST | LOC127872065 | exp5 | Dmel_CG12234 | -- | AT3G05040 |
| Exportin-1 | LOC127857673 | exp1 | Dmel_CG13387 | CELE_ZK742.1 | AT5G17020 |
| Exportin-2 | LOC127862014 | XPOTP2 | Q9XZU1 | CELE_Y48G1A.5 | AT2G46520 |
| TARBP2 | LOC127855042 | TARBP2 | Dmel_CG6866 | CELE_T20G5.11 | -- |
| Pasha/DGCR8/DR B1 | LOC127846964 | DGCR8 | Dmel_CG1800 | CELE_T22A3.5 | AT1G09700 |
| R2d2 | -- | -- | Dmel_CG7138 | CELE_T20G5.11 | -- |
| tsn-1 | LOC127852824 | TSN | Dmel_CG7008 | CELE_F10G7.2 | AT5G07350 |
| vig-1 | -- | SERBP1 | Dmel_CG4170 | CELE_F56D12.5 | -- |
| Argonaute 1 | -- | AGO1 | Dmel_CG6671 | CELE_F48F7.1 | AT1G48410 |
| Argonaute 2 | LOC127847243 | CASC7 | Dmel_CG7439 | CELE_T23D8.7 | AT1G31280 |
| Piwi | LOC127837144 | CT80.1 | Dmel_CG6122 | CELE_K08H10.7 | -- |
| adr-2 | LOC127856227 | ADARB1 | Dmel_CG12598 | CELE_T20H4.4 | -- |
| eri-1 | -- | ERI1 | Dmel_CG44248 | CELE_T07A9.5 | AT3G15140 |
| eri-6/Mov10/armitage/<br>SED3 | -- | MOV10 | Dmel_CG11513 | CELE_C41D11.1 | AT1G05460 |
| xrn-1 | LOC127863549 | XRN1 | Dmel_CG3291 | CELE_Y39G8C.1 | AT1G54490 |
| xrn-2 | LOC127870862 | XRN2 | Dmel_CG10354 | CELE_Y48B6A.3 | AT5G42540 |
| GW182/ain 1-2 | LOC127850135 | GW182 | Dmel_CG31992 | CELE_F48F7.1 | -- |
| Sid-1 | XP_052228431.1 | SIDT | -- | CELE_C04F5.1 | -- |
| rsd-3 | -- | ENTHD1 | -- | CELE_C34E11.1 | -- |

**Supplementary Table 2.**
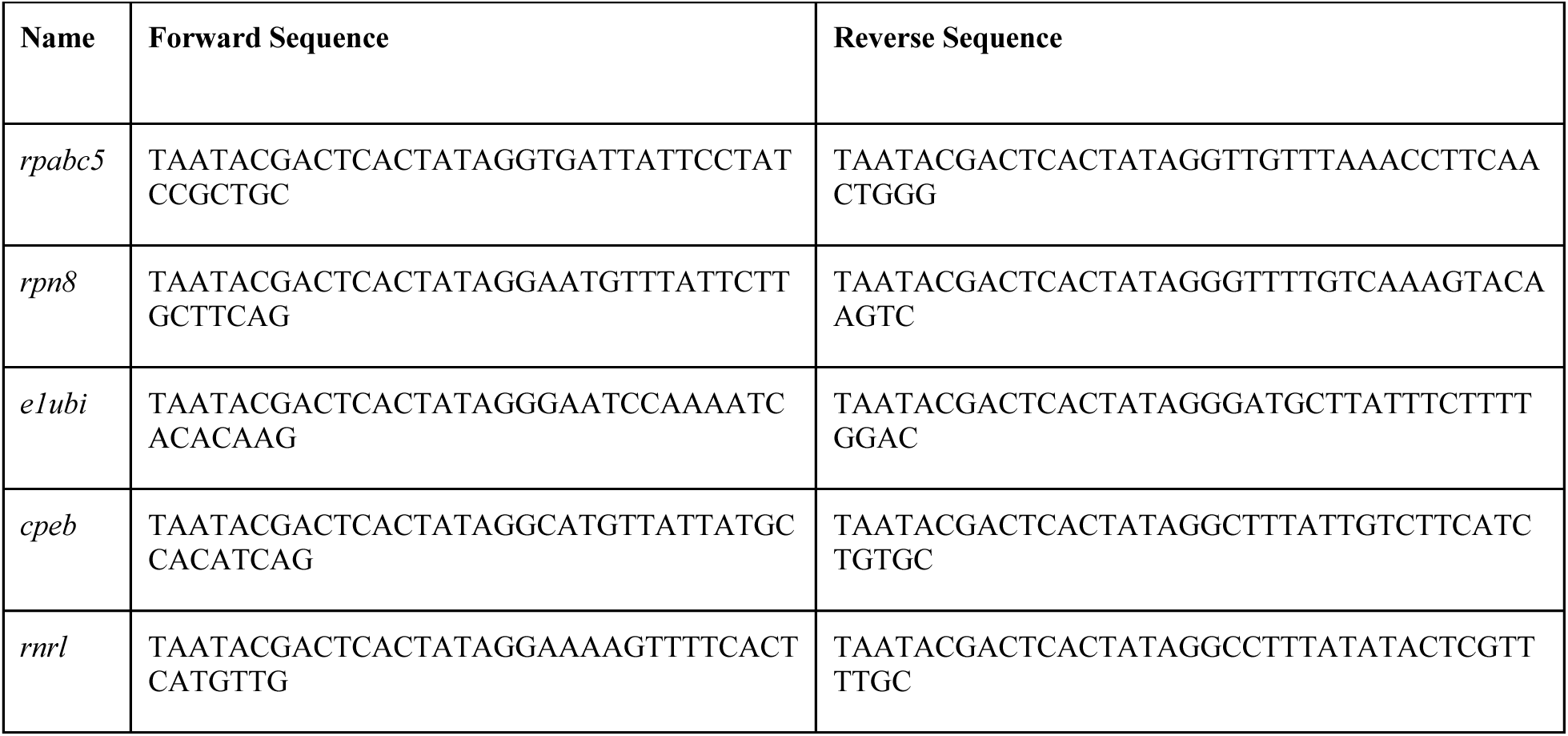
Primer sequences used for the generation of dsRNA templates.

**Supplementary Table 3.**
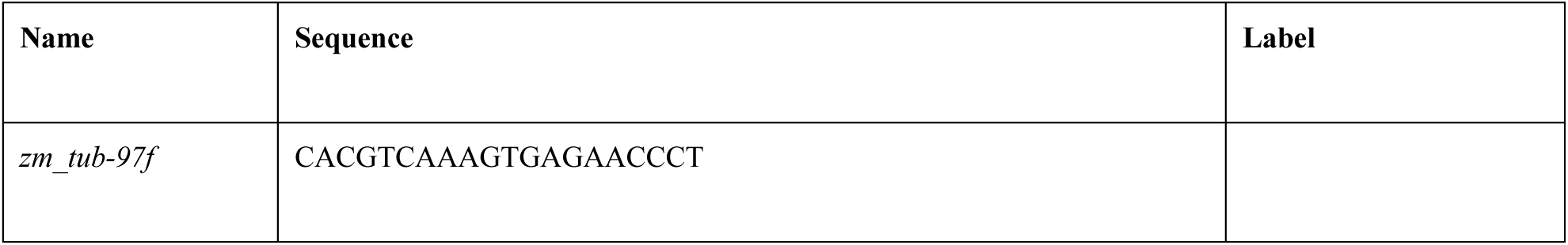
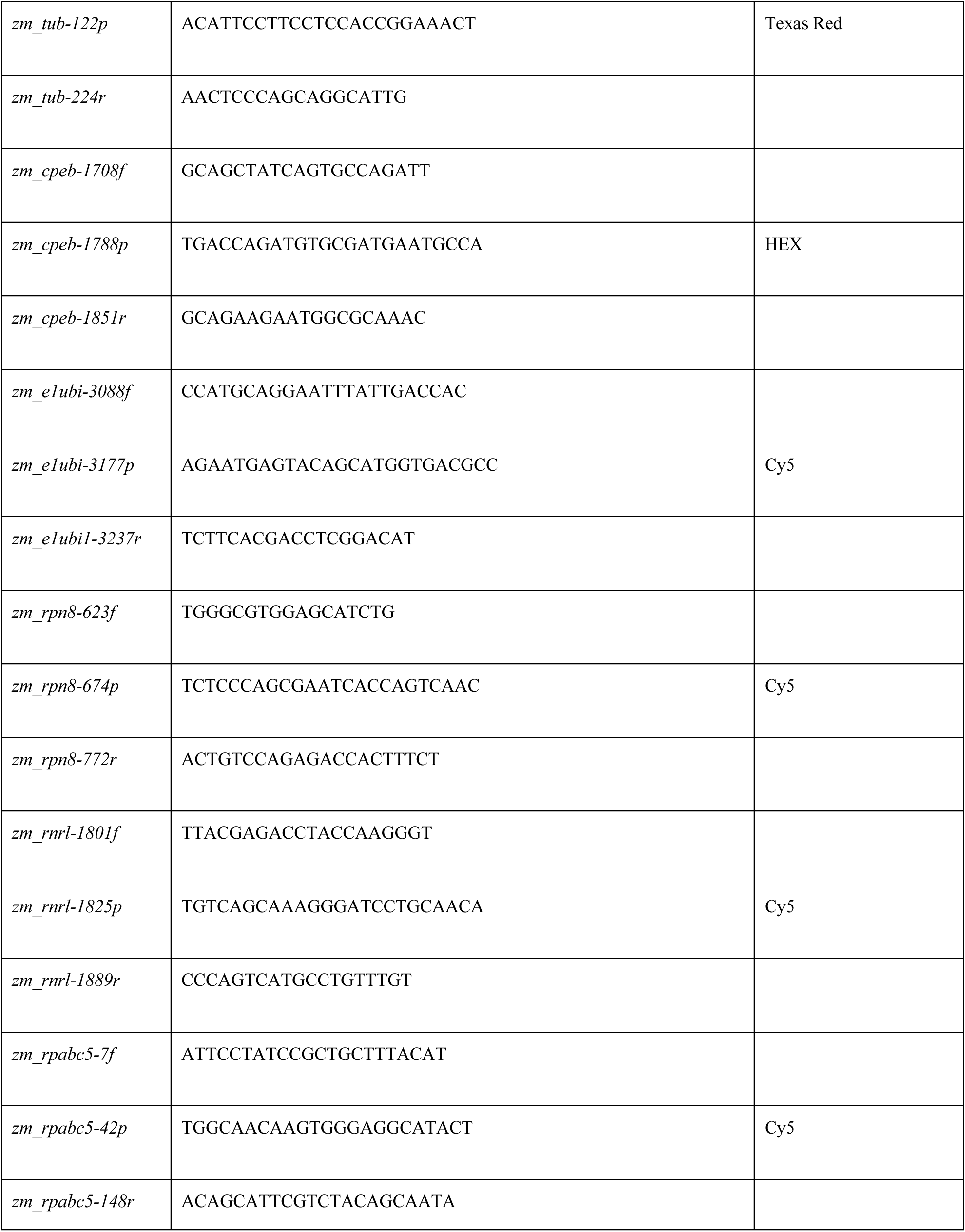
Primer and probe sequences used for RT-qPCR.

