## Supplementary File 4 for "Characterizing the Small Non-Coding RNA Pathways in the Invasive Zebra Mussel (*Dreissena polymorpha*)"

                                              /\
                _______________________/\___ /  \_______
               I                      /  \  /    \      I
               I     pro             /    \/      \     I
               I        TRAC        /               \   I
               I   ________________/_________________\_ I
               I   \              /                     I
               I    \            /                      I
               I     \  /\      /       V.2.4.2         I
               I      \/  \    /                        I
               I___________\  /_________________________I
                            \/


================================= proTRAC ====================================
VERSION: .......... 2.4.2
LAST MODIFIED: .... 11. May 2018

Please cite:
Rosenkranz D, Zischler H. proTRAC - a software for probabilistic piRNA cluster
detection, visualization and analysis. 2012. BMC Bioinformatics 13:5.


Contact:
David Rosenkranz
Institute of Organismic and Molecular Evolutionary Biology
Dept. Anthropology, small RNA group
Johannes Gutenberg University Mainz


You can find the latest proTRAC version at:
http://sourceforge.net/projects/protrac/files
http://www.smallRNAgroup-mainz.de/software
==============================================================================

PARAMETERS:
Map file: .............../Volumes/My_Book/SEQUENCING_DATA/sRNA/sRNA_DETECTION/Adductor_25.sam
Genome file: ............/Volumes/My_Book/SEQUENCING_DATA/sRNA/miRDeep2/Bowtiew_genome_index/GCA_020536995.1_UMN_Dpol_1.0_genomic.fasta
RepeatMasker annotation: n.a.
GeneSet:................./Volumes/My_Book/SEQUENCING_DATA/sRNA/miRDeep2/Bowtiew_genome_index/genomic.gff

Significant (p<=0.01) hit density will be calculated based
on observed hit distribution.

Sliding window size: ........................................ 5000 bp
Sliding window increament: .................................. 1000 bp
Normalize each hit by number of genomic hits: ............... yes
Normalize each hit by number of sequence reads: ............. yes
Normalize values (-> per million mapped reads): ............. yes
Min. fraction of hits with 1T(U) or 10A: .................... 0.75
Alternatively: Min. fraction of hits with 1T(U) and 10A: .... 0.5
Min. fraction of hits with typical piRNA length: ............ 0.75
Typical piRNA length: ....................................... 25-32 nt
Min. size of a piRNA cluster: ............................... 1000 bp.
Min. number of hits (absolute): ............................. 0
Min. number of hits (normalized): ........................... 0
Min. fraction of hits on the mainstrand: .................... 0.75
Top fraction of mapped sequences (in terms of read counts): . 1%
Top fraction accounts for max. n% of sequence reads: ........ 90%
Min. fraction of hits on each arm of a bidirectional cluster: 0.05
Output html file for each cluster: .......................... no
Output a summary table: ..................................... yes
Output a FASTA file for each cluster (piRNA sequences): ..... yes
Output a FASTA file comprising cluster sequences: ........... yes
Output a GTF file for predicted piRNA clusters: ..............yes
Search DNA motifs in clusters: .............................. yes
Output flanking sequences: +/- .............................. 0 bp
Output ~.pTi file: .......................................... no
==============================================================================


Cluster 1	Location: CM035915.1	Coordinates: 6110058-6114292	Size [bp]: 4235	Hits (absolute): 389	Hits (normalized): 90.6758119247916	Hits (normalized) per kb: 21.4111123036848	Normalized hits with 1T: 96.3%	Normalized hits with 10A: 18.5%	Normalized hits 25-32 nt: 99.9%	Normalized hits on the main strand(s): 99.5%	Predicted directionality: mono:minus
Cluster 2	Location: CM035915.1	Coordinates: 20675027-20679683	Size [bp]: 4657	Hits (absolute): 1470	Hits (normalized): 97.1985200390135	Hits (normalized) per kb: 20.8715536714522	Normalized hits with 1T: 86.3%	Normalized hits with 10A: 36.1%	Normalized hits 25-32 nt: 99.2%	Normalized hits on the main strand(s): 83.1%	Predicted directionality: mono:plus
Cluster 3	Location: CM035915.1	Coordinates: 22385016-22391928	Size [bp]: 6913	Hits (absolute): 2526	Hits (normalized): 128.408044879024	Hits (normalized) per kb: 18.5749349337369	Normalized hits with 1T: 95.9%	Normalized hits with 10A: 26%	Normalized hits 25-32 nt: 98.8%	Normalized hits on the main strand(s): 99.8%	Predicted directionality: mono:minus
Cluster 4	Location: CM035915.1	Coordinates: 25451020-25457884	Size [bp]: 6865	Hits (absolute): 940	Hits (normalized): 274.951045531925	Hits (normalized) per kb: 40.051045880942	Normalized hits with 1T: 98.7%	Normalized hits with 10A: 10.1%	Normalized hits 25-32 nt: 99.9%	Normalized hits on the main strand(s): 99.1%	Predicted directionality: mono:minus
Cluster 5	Location: CM035915.1	Coordinates: 28206092-28210703	Size [bp]: 4612	Hits (absolute): 463	Hits (normalized): 110.505149025966	Hits (normalized) per kb: 23.9600376038819	Normalized hits with 1T: 3.5%	Normalized hits with 10A: 97.3%	Normalized hits 25-32 nt: 99.2%	Normalized hits on the main strand(s): 97.9%	Predicted directionality: mono:minus
Cluster 6	Location: CM035915.1	Coordinates: 39074652-39079797	Size [bp]: 5146	Hits (absolute): 855	Hits (normalized): 221.837379346456	Hits (normalized) per kb: 43.108777766975	Normalized hits with 1T: 15.2%	Normalized hits with 10A: 87.1%	Normalized hits 25-32 nt: 99.2%	Normalized hits on the main strand(s): 93.1%	Predicted directionality: bi:plus-minus (split between 39078496 and 39078497)
Cluster 7	Location: CM035915.1	Coordinates: 50015780-50029353	Size [bp]: 13574	Hits (absolute): 2051	Hits (normalized): 806.841969813903	Hits (normalized) per kb: 59.4402111340454	Normalized hits with 1T: 97.2%	Normalized hits with 10A: 42.5%	Normalized hits 25-32 nt: 99.6%	Normalized hits on the main strand(s): 100%	Predicted directionality: mono:minus
Cluster 8	Location: CM035915.1	Coordinates: 50036093-50074946	Size [bp]: 38854	Hits (absolute): 11047	Hits (normalized): 5339.06046836494	Hits (normalized) per kb: 137.413422593105	Normalized hits with 1T: 88.3%	Normalized hits with 10A: 43.8%	Normalized hits 25-32 nt: 99.8%	Normalized hits on the main strand(s): 100%	Predicted directionality: mono:minus
Cluster 9	Location: CM035915.1	Coordinates: 57317195-57324899	Size [bp]: 7705	Hits (absolute): 590	Hits (normalized): 140.69508300238	Hits (normalized) per kb: 18.2604253683164	Normalized hits with 1T: 92.7%	Normalized hits with 10A: 27.5%	Normalized hits 25-32 nt: 98.5%	Normalized hits on the main strand(s): 93.9%	Predicted directionality: bi:minus-plus (split between 57320187 and 57320192)
Cluster 10	Location: CM035915.1	Coordinates: 57671132-57677843	Size [bp]: 6712	Hits (absolute): 685	Hits (normalized): 105.773292404408	Hits (normalized) per kb: 15.759025958005	Normalized hits with 1T: 21%	Normalized hits with 10A: 94.6%	Normalized hits 25-32 nt: 99%	Normalized hits on the main strand(s): 95%	Predicted directionality: mono:plus
Cluster 11	Location: CM035915.1	Coordinates: 79756013-79768025	Size [bp]: 12013	Hits (absolute): 1080	Hits (normalized): 217.509320248842	Hits (normalized) per kb: 18.1059662261876	Normalized hits with 1T: 76.2%	Normalized hits with 10A: 35.5%	Normalized hits 25-32 nt: 99.6%	Normalized hits on the main strand(s): 100%	Predicted directionality: mono:minus
Cluster 12	Location: CM035915.1	Coordinates: 84299195-84313987	Size [bp]: 14793	Hits (absolute): 4327	Hits (normalized): 1027.63375555941	Hits (normalized) per kb: 69.4674749897975	Normalized hits with 1T: 96.6%	Normalized hits with 10A: 14.2%	Normalized hits 25-32 nt: 99.6%	Normalized hits on the main strand(s): 100%	Predicted directionality: mono:plus
Cluster 13	Location: CM035915.1	Coordinates: 91510862-91513965	Size [bp]: 3104	Hits (absolute): 949	Hits (normalized): 442.057638341569	Hits (normalized) per kb: 142.415522503582	Normalized hits with 1T: 84.4%	Normalized hits with 10A: 22.9%	Normalized hits 25-32 nt: 99.7%	Normalized hits on the main strand(s): 88.9%	Predicted directionality: bi:minus-plus (split between 91512060 and 91512063)
Cluster 14	Location: CM035915.1	Coordinates: 99804204-99806930	Size [bp]: 2727	Hits (absolute): 301	Hits (normalized): 140.825942553466	Hits (normalized) per kb: 51.6410728217614	Normalized hits with 1T: 97.8%	Normalized hits with 10A: 89.8%	Normalized hits 25-32 nt: 100%	Normalized hits on the main strand(s): 100%	Predicted directionality: mono:minus
Cluster 15	Location: CM035915.1	Coordinates: 109698761-109703010	Size [bp]: 4250	Hits (absolute): 1577	Hits (normalized): 108.064934976756	Hits (normalized) per kb: 25.4270499990324	Normalized hits with 1T: 85.6%	Normalized hits with 10A: 35.9%	Normalized hits 25-32 nt: 99.2%	Normalized hits on the main strand(s): 85.1%	Predicted directionality: mono:minus
Cluster 16	Location: CM035915.1	Coordinates: 117041064-117049779	Size [bp]: 8716	Hits (absolute): 330	Hits (normalized): 126.99364947613	Hits (normalized) per kb: 14.5701798007154	Normalized hits with 1T: 99.2%	Normalized hits with 10A: 37.4%	Normalized hits 25-32 nt: 99.9%	Normalized hits on the main strand(s): 99.4%	Predicted directionality: mono:minus
Cluster 17	Location: CM035915.1	Coordinates: 119296483-119306840	Size [bp]: 10358	Hits (absolute): 2145	Hits (normalized): 244.247556184193	Hits (normalized) per kb: 23.5805293949411	Normalized hits with 1T: 89.7%	Normalized hits with 10A: 28.1%	Normalized hits 25-32 nt: 99.6%	Normalized hits on the main strand(s): 100%	Predicted directionality: mono:plus
Cluster 18	Location: CM035915.1	Coordinates: 119311212-119327844	Size [bp]: 16633	Hits (absolute): 3309	Hits (normalized): 501.075236315424	Hits (normalized) per kb: 30.1251239962698	Normalized hits with 1T: 93.8%	Normalized hits with 10A: 23.5%	Normalized hits 25-32 nt: 99.8%	Normalized hits on the main strand(s): 100%	Predicted directionality: mono:plus
Cluster 19	Location: CM035915.1	Coordinates: 119340003-119355959	Size [bp]: 15957	Hits (absolute): 2890	Hits (normalized): 385.818793395244	Hits (normalized) per kb: 24.1787964793855	Normalized hits with 1T: 96.8%	Normalized hits with 10A: 23.6%	Normalized hits 25-32 nt: 99.8%	Normalized hits on the main strand(s): 99.9%	Predicted directionality: mono:plus
Cluster 20	Location: CM035915.1	Coordinates: 129807136-129813729	Size [bp]: 6594	Hits (absolute): 1812	Hits (normalized): 235.704507558441	Hits (normalized) per kb: 35.7450604752621	Normalized hits with 1T: 91.9%	Normalized hits with 10A: 70.1%	Normalized hits 25-32 nt: 99.5%	Normalized hits on the main strand(s): 92.1%	Predicted directionality: mono:minus
Cluster 21	Location: CM035915.1	Coordinates: 131087179-131093272	Size [bp]: 6094	Hits (absolute): 405	Hits (normalized): 106.880463510837	Hits (normalized) per kb: 17.5384511881399	Normalized hits with 1T: 98.7%	Normalized hits with 10A: 4.3%	Normalized hits 25-32 nt: 99.9%	Normalized hits on the main strand(s): 98.9%	Predicted directionality: mono:plus
Cluster 22	Location: CM035915.1	Coordinates: 132750070-132758829	Size [bp]: 8760	Hits (absolute): 926	Hits (normalized): 741.336956697421	Hits (normalized) per kb: 84.6275349532127	Normalized hits with 1T: 98.9%	Normalized hits with 10A: 2%	Normalized hits 25-32 nt: 99.9%	Normalized hits on the main strand(s): 99.7%	Predicted directionality: bi:plus-minus (split between 132754410 and 132754431)
Cluster 23	Location: CM035915.1	Coordinates: 133352026-133361027	Size [bp]: 9002	Hits (absolute): 1730	Hits (normalized): 612.107553643979	Hits (normalized) per kb: 67.9969680439201	Normalized hits with 1T: 89.5%	Normalized hits with 10A: 8.2%	Normalized hits 25-32 nt: 99.8%	Normalized hits on the main strand(s): 100%	Predicted directionality: mono:minus
Cluster 24	Location: CM035915.1	Coordinates: 133362002-133450980	Size [bp]: 88979	Hits (absolute): 20797	Hits (normalized): 4903.33048223231	Hits (normalized) per kb: 55.1062693225504	Normalized hits with 1T: 89.3%	Normalized hits with 10A: 23.9%	Normalized hits 25-32 nt: 99.5%	Normalized hits on the main strand(s): 99.7%	Predicted directionality: mono:minus
Cluster 25	Location: CM035915.1	Coordinates: 133712040-133719837	Size [bp]: 7798	Hits (absolute): 1196	Hits (normalized): 159.964157361858	Hits (normalized) per kb: 20.5137116770182	Normalized hits with 1T: 96.7%	Normalized hits with 10A: 22.5%	Normalized hits 25-32 nt: 99.9%	Normalized hits on the main strand(s): 100%	Predicted directionality: mono:plus
Cluster 26	Location: CM035915.1	Coordinates: 133737807-133742918	Size [bp]: 5112	Hits (absolute): 709	Hits (normalized): 90.5960528672128	Hits (normalized) per kb: 17.7222645563746	Normalized hits with 1T: 89.4%	Normalized hits with 10A: 44.1%	Normalized hits 25-32 nt: 99.8%	Normalized hits on the main strand(s): 96.5%	Predicted directionality: mono:plus
Cluster 27	Location: CM035915.1	Coordinates: 133755151-133788018	Size [bp]: 32868	Hits (absolute): 5584	Hits (normalized): 1093.22636148365	Hits (normalized) per kb: 33.2611338185853	Normalized hits with 1T: 89.7%	Normalized hits with 10A: 16.7%	Normalized hits 25-32 nt: 99.7%	Normalized hits on the main strand(s): 100%	Predicted directionality: mono:plus
Cluster 28	Location: CM035915.1	Coordinates: 133792031-133803028	Size [bp]: 10998	Hits (absolute): 1539	Hits (normalized): 318.376984061751	Hits (normalized) per kb: 28.948858221597	Normalized hits with 1T: 93.6%	Normalized hits with 10A: 26.2%	Normalized hits 25-32 nt: 99.6%	Normalized hits on the main strand(s): 99.9%	Predicted directionality: mono:plus
Cluster 29	Location: CM035915.1	Coordinates: 133813067-133846965	Size [bp]: 33899	Hits (absolute): 6193	Hits (normalized): 1309.87192047285	Hits (normalized) per kb: 38.6406452075672	Normalized hits with 1T: 93.5%	Normalized hits with 10A: 22.3%	Normalized hits 25-32 nt: 99.8%	Normalized hits on the main strand(s): 100%	Predicted directionality: mono:plus
Cluster 30	Location: CM035915.1	Coordinates: 135075571-135079998	Size [bp]: 4428	Hits (absolute): 926	Hits (normalized): 92.5630522677932	Hits (normalized) per kb: 20.9037035381397	Normalized hits with 1T: 88.7%	Normalized hits with 10A: 60.9%	Normalized hits 25-32 nt: 99.5%	Normalized hits on the main strand(s): 86.5%	Predicted directionality: bi:plus-minus (split between 135076856 and 135077150)
Cluster 31	Location: CM035915.1	Coordinates: 135524636-135530802	Size [bp]: 6167	Hits (absolute): 1022	Hits (normalized): 164.113931119748	Hits (normalized) per kb: 26.6117026954497	Normalized hits with 1T: 80.7%	Normalized hits with 10A: 26.4%	Normalized hits 25-32 nt: 99.2%	Normalized hits on the main strand(s): 87.6%	Predicted directionality: bi:minus-plus (split between 135528799 and 135528799)
Cluster 32	Location: CM035915.1	Coordinates: 139691221-139740739	Size [bp]: 49519	Hits (absolute): 14160	Hits (normalized): 3257.66433915114	Hits (normalized) per kb: 65.7863152540864	Normalized hits with 1T: 96.8%	Normalized hits with 10A: 16%	Normalized hits 25-32 nt: 99.7%	Normalized hits on the main strand(s): 99.7%	Predicted directionality: mono:minus
Cluster 33	Location: CM035915.1	Coordinates: 139744025-139763243	Size [bp]: 19219	Hits (absolute): 4829	Hits (normalized): 1148.46824112546	Hits (normalized) per kb: 59.7568174299021	Normalized hits with 1T: 96%	Normalized hits with 10A: 19.9%	Normalized hits 25-32 nt: 99.7%	Normalized hits on the main strand(s): 99.9%	Predicted directionality: mono:minus
Cluster 34	Location: CM035915.1	Coordinates: 140104982-140111022	Size [bp]: 6041	Hits (absolute): 1162	Hits (normalized): 115.294064701385	Hits (normalized) per kb: 19.0851393398636	Normalized hits with 1T: 96.9%	Normalized hits with 10A: 17.5%	Normalized hits 25-32 nt: 99.8%	Normalized hits on the main strand(s): 100%	Predicted directionality: mono:minus
Cluster 35	Location: CM035915.1	Coordinates: 140154810-140172001	Size [bp]: 17192	Hits (absolute): 5510	Hits (normalized): 1159.80677457301	Hits (normalized) per kb: 67.4623017827052	Normalized hits with 1T: 92.6%	Normalized hits with 10A: 36.4%	Normalized hits 25-32 nt: 99.6%	Normalized hits on the main strand(s): 100%	Predicted directionality: mono:minus
Cluster 36	Location: CM035915.1	Coordinates: 140178014-140182824	Size [bp]: 4811	Hits (absolute): 860	Hits (normalized): 92.9705166796899	Hits (normalized) per kb: 19.3248655197286	Normalized hits with 1T: 92.2%	Normalized hits with 10A: 35.8%	Normalized hits 25-32 nt: 98.6%	Normalized hits on the main strand(s): 100%	Predicted directionality: mono:minus
Cluster 37	Location: CM035915.1	Coordinates: 155239309-155244968	Size [bp]: 5660	Hits (absolute): 1023	Hits (normalized): 166.959355808791	Hits (normalized) per kb: 29.4982015958649	Normalized hits with 1T: 79.3%	Normalized hits with 10A: 25.9%	Normalized hits 25-32 nt: 99.2%	Normalized hits on the main strand(s): 86.2%	Predicted directionality: bi:minus-plus (split between 155242964 and 155242965)
Cluster 38	Location: CM035915.1	Coordinates: 156283600-156287940	Size [bp]: 4341	Hits (absolute): 671	Hits (normalized): 119.512158174641	Hits (normalized) per kb: 27.5307695366231	Normalized hits with 1T: 98.1%	Normalized hits with 10A: 19.7%	Normalized hits 25-32 nt: 99.7%	Normalized hits on the main strand(s): 100%	Predicted directionality: mono:minus
Cluster 39	Location: CM035915.1	Coordinates: 156303835-156305507	Size [bp]: 1673	Hits (absolute): 400	Hits (normalized): 101.036241056333	Hits (normalized) per kb: 60.3921267520515	Normalized hits with 1T: 97%	Normalized hits with 10A: 12.7%	Normalized hits 25-32 nt: 99.7%	Normalized hits on the main strand(s): 99.8%	Predicted directionality: mono:minus
Cluster 40	Location: CM035915.1	Coordinates: 160827600-160833147	Size [bp]: 5548	Hits (absolute): 727	Hits (normalized): 144.092976383074	Hits (normalized) per kb: 25.972199912428	Normalized hits with 1T: 96.2%	Normalized hits with 10A: 13.8%	Normalized hits 25-32 nt: 97.9%	Normalized hits on the main strand(s): 99.9%	Predicted directionality: mono:minus
Cluster 41	Location: CM035915.1	Coordinates: 174124146-174128697	Size [bp]: 4552	Hits (absolute): 1500	Hits (normalized): 103.191594660016	Hits (normalized) per kb: 22.6698494755123	Normalized hits with 1T: 86.4%	Normalized hits with 10A: 33.8%	Normalized hits 25-32 nt: 99.4%	Normalized hits on the main strand(s): 84.5%	Predicted directionality: mono:plus
Cluster 42	Location: CM035915.1	Coordinates: 179076010-179078647	Size [bp]: 2638	Hits (absolute): 1710	Hits (normalized): 156.379056618468	Hits (normalized) per kb: 59.2794618006082	Normalized hits with 1T: 86.7%	Normalized hits with 10A: 21.7%	Normalized hits 25-32 nt: 99.6%	Normalized hits on the main strand(s): 97.2%	Predicted directionality: mono:minus
Cluster 43	Location: CM035915.1	Coordinates: 192220184-192224879	Size [bp]: 4696	Hits (absolute): 1317	Hits (normalized): 99.2581007836552	Hits (normalized) per kb: 21.1364406165509	Normalized hits with 1T: 86.4%	Normalized hits with 10A: 34.1%	Normalized hits 25-32 nt: 99.6%	Normalized hits on the main strand(s): 87%	Predicted directionality: mono:plus
Cluster 44	Location: CM035915.1	Coordinates: 194367022-194374027	Size [bp]: 7006	Hits (absolute): 1012	Hits (normalized): 130.739786477576	Hits (normalized) per kb: 18.6609008816185	Normalized hits with 1T: 94%	Normalized hits with 10A: 15.3%	Normalized hits 25-32 nt: 99.6%	Normalized hits on the main strand(s): 95.2%	Predicted directionality: bi:plus-minus (split between 194369812 and 194369918)
Cluster 45	Location: CM035915.1	Coordinates: 194386873-194391831	Size [bp]: 4959	Hits (absolute): 774	Hits (normalized): 218.445297818391	Hits (normalized) per kb: 44.0502097328004	Normalized hits with 1T: 97.5%	Normalized hits with 10A: 8.1%	Normalized hits 25-32 nt: 99.9%	Normalized hits on the main strand(s): 100%	Predicted directionality: mono:minus
Cluster 46	Location: CM035915.1	Coordinates: 196437426-196446463	Size [bp]: 9038	Hits (absolute): 426	Hits (normalized): 196.410862855011	Hits (normalized) per kb: 21.7319120604137	Normalized hits with 1T: 98.9%	Normalized hits with 10A: 0.4%	Normalized hits 25-32 nt: 99.9%	Normalized hits on the main strand(s): 99.8%	Predicted directionality: mono:plus
Cluster 47	Location: CM035915.1	Coordinates: 199155040-199159716	Size [bp]: 4677	Hits (absolute): 163	Hits (normalized): 248.710763772016	Hits (normalized) per kb: 53.1772773213044	Normalized hits with 1T: 99.7%	Normalized hits with 10A: 99.7%	Normalized hits 25-32 nt: 99.6%	Normalized hits on the main strand(s): 100%	Predicted directionality: mono:minus
Cluster 48	Location: CM035916.1	Coordinates: 17650807-17655673	Size [bp]: 4867	Hits (absolute): 406	Hits (normalized): 214.285377598935	Hits (normalized) per kb: 44.0285435182937	Normalized hits with 1T: 99.2%	Normalized hits with 10A: 1.1%	Normalized hits 25-32 nt: 99.8%	Normalized hits on the main strand(s): 99.9%	Predicted directionality: mono:minus
Cluster 49	Location: CM035916.1	Coordinates: 42401541-42406764	Size [bp]: 5224	Hits (absolute): 886	Hits (normalized): 107.475829893417	Hits (normalized) per kb: 20.5738179495208	Normalized hits with 1T: 94%	Normalized hits with 10A: 12.2%	Normalized hits 25-32 nt: 99.7%	Normalized hits on the main strand(s): 93.9%	Predicted directionality: mono:plus
Cluster 50	Location: CM035916.1	Coordinates: 44322089-44330981	Size [bp]: 8893	Hits (absolute): 1384	Hits (normalized): 169.302246341239	Hits (normalized) per kb: 19.0376134499778	Normalized hits with 1T: 97.9%	Normalized hits with 10A: 24.6%	Normalized hits 25-32 nt: 99.8%	Normalized hits on the main strand(s): 99.3%	Predicted directionality: mono:minus
Cluster 51	Location: CM035916.1	Coordinates: 46084164-46089850	Size [bp]: 5687	Hits (absolute): 958	Hits (normalized): 466.13028275753	Hits (normalized) per kb: 81.9639883891735	Normalized hits with 1T: 80.1%	Normalized hits with 10A: 22.2%	Normalized hits 25-32 nt: 99.8%	Normalized hits on the main strand(s): 89.4%	Predicted directionality: bi:minus-plus (split between 46085582 and 46085582)
Cluster 52	Location: CM035916.1	Coordinates: 49615434-49619597	Size [bp]: 4164	Hits (absolute): 246	Hits (normalized): 140.434345211481	Hits (normalized) per kb: 33.7259090652623	Normalized hits with 1T: 87.4%	Normalized hits with 10A: 98.6%	Normalized hits 25-32 nt: 99.7%	Normalized hits on the main strand(s): 98.1%	Predicted directionality: mono:plus
Cluster 53	Location: CM035916.1	Coordinates: 52344190-52350940	Size [bp]: 6751	Hits (absolute): 582	Hits (normalized): 114.686694392629	Hits (normalized) per kb: 16.9884089037267	Normalized hits with 1T: 96.9%	Normalized hits with 10A: 16.5%	Normalized hits 25-32 nt: 99.9%	Normalized hits on the main strand(s): 99.7%	Predicted directionality: mono:minus
Cluster 54	Location: CM035916.1	Coordinates: 52658184-52660287	Size [bp]: 2104	Hits (absolute): 244	Hits (normalized): 161.859217211427	Hits (normalized) per kb: 76.9290397018634	Normalized hits with 1T: 0.2%	Normalized hits with 10A: 99.6%	Normalized hits 25-32 nt: 99.7%	Normalized hits on the main strand(s): 100%	Predicted directionality: mono:plus
Cluster 55	Location: CM035916.1	Coordinates: 53501209-53507686	Size [bp]: 6478	Hits (absolute): 321	Hits (normalized): 177.849360322683	Hits (normalized) per kb: 27.4545883307768	Normalized hits with 1T: 1%	Normalized hits with 10A: 98.9%	Normalized hits 25-32 nt: 99.6%	Normalized hits on the main strand(s): 100%	Predicted directionality: mono:plus
Cluster 56	Location: CM035916.1	Coordinates: 53716065-53720962	Size [bp]: 4898	Hits (absolute): 1076	Hits (normalized): 187.863139568033	Hits (normalized) per kb: 38.3547909581072	Normalized hits with 1T: 90%	Normalized hits with 10A: 26.4%	Normalized hits 25-32 nt: 98.2%	Normalized hits on the main strand(s): 94.1%	Predicted directionality: bi:minus-plus (split between 53720311 and 53720312)
Cluster 57	Location: CM035916.1	Coordinates: 55580276-55583723	Size [bp]: 3448	Hits (absolute): 437	Hits (normalized): 185.075625811802	Hits (normalized) per kb: 53.676299165105	Normalized hits with 1T: 0.9%	Normalized hits with 10A: 98.8%	Normalized hits 25-32 nt: 99.4%	Normalized hits on the main strand(s): 100%	Predicted directionality: mono:minus
Cluster 58	Location: CM035916.1	Coordinates: 65386447-65390891	Size [bp]: 4445	Hits (absolute): 173	Hits (normalized): 285.427626258896	Hits (normalized) per kb: 64.2130685168384	Normalized hits with 1T: 97%	Normalized hits with 10A: 0.5%	Normalized hits 25-32 nt: 98.2%	Normalized hits on the main strand(s): 99.8%	Predicted directionality: mono:minus
Cluster 59	Location: CM035916.1	Coordinates: 71627047-71632973	Size [bp]: 5927	Hits (absolute): 1557	Hits (normalized): 187.551855454651	Hits (normalized) per kb: 31.6438557421783	Normalized hits with 1T: 94.7%	Normalized hits with 10A: 12.7%	Normalized hits 25-32 nt: 99.8%	Normalized hits on the main strand(s): 95.2%	Predicted directionality: mono:plus
Cluster 60	Location: CM035916.1	Coordinates: 73466127-73468749	Size [bp]: 2623	Hits (absolute): 381	Hits (normalized): 91.93737203112	Hits (normalized) per kb: 35.0503437907554	Normalized hits with 1T: 96.8%	Normalized hits with 10A: 91.3%	Normalized hits 25-32 nt: 99.6%	Normalized hits on the main strand(s): 96.8%	Predicted directionality: mono:minus
Cluster 61	Location: CM035916.1	Coordinates: 75201335-75220975	Size [bp]: 19641	Hits (absolute): 2984	Hits (normalized): 472.023015616016	Hits (normalized) per kb: 24.0327242590013	Normalized hits with 1T: 83.9%	Normalized hits with 10A: 21.6%	Normalized hits 25-32 nt: 99.7%	Normalized hits on the main strand(s): 99.6%	Predicted directionality: mono:plus
Cluster 62	Location: CM035916.1	Coordinates: 75322047-75331029	Size [bp]: 8983	Hits (absolute): 1091	Hits (normalized): 163.39214566689	Hits (normalized) per kb: 18.1891365334877	Normalized hits with 1T: 89.2%	Normalized hits with 10A: 10.7%	Normalized hits 25-32 nt: 99.7%	Normalized hits on the main strand(s): 100%	Predicted directionality: mono:minus
Cluster 63	Location: CM035916.1	Coordinates: 75866928-75868109	Size [bp]: 1182	Hits (absolute): 404	Hits (normalized): 224.703629464147	Hits (normalized) per kb: 190.104258453226	Normalized hits with 1T: 99%	Normalized hits with 10A: 4.6%	Normalized hits 25-32 nt: 99.9%	Normalized hits on the main strand(s): 99.9%	Predicted directionality: mono:minus
Cluster 64	Location: CM035916.1	Coordinates: 82180437-82188064	Size [bp]: 7628	Hits (absolute): 1505	Hits (normalized): 227.922145979404	Hits (normalized) per kb: 29.8798065352418	Normalized hits with 1T: 96.3%	Normalized hits with 10A: 30.8%	Normalized hits 25-32 nt: 99.6%	Normalized hits on the main strand(s): 99.2%	Predicted directionality: mono:plus
Cluster 65	Location: CM035916.1	Coordinates: 86183183-86189511	Size [bp]: 6329	Hits (absolute): 673	Hits (normalized): 138.644481103753	Hits (normalized) per kb: 21.9059406866131	Normalized hits with 1T: 96%	Normalized hits with 10A: 4.4%	Normalized hits 25-32 nt: 97.8%	Normalized hits on the main strand(s): 99.9%	Predicted directionality: mono:minus
Cluster 66	Location: CM035916.1	Coordinates: 87947717-87954720	Size [bp]: 7004	Hits (absolute): 812	Hits (normalized): 163.64045396217	Hits (normalized) per kb: 23.3638672498736	Normalized hits with 1T: 97.6%	Normalized hits with 10A: 2.6%	Normalized hits 25-32 nt: 98.1%	Normalized hits on the main strand(s): 100%	Predicted directionality: mono:minus
Cluster 67	Location: CM035916.1	Coordinates: 88903933-88908993	Size [bp]: 5061	Hits (absolute): 699	Hits (normalized): 152.959154105439	Hits (normalized) per kb: 30.2229714166229	Normalized hits with 1T: 96.3%	Normalized hits with 10A: 12.6%	Normalized hits 25-32 nt: 98%	Normalized hits on the main strand(s): 99.9%	Predicted directionality: mono:minus
Cluster 68	Location: CM035916.1	Coordinates: 91601220-91602898	Size [bp]: 1679	Hits (absolute): 197	Hits (normalized): 192.55673415336	Hits (normalized) per kb: 114.685563575526	Normalized hits with 1T: 0.3%	Normalized hits with 10A: 95.4%	Normalized hits 25-32 nt: 99.9%	Normalized hits on the main strand(s): 100%	Predicted directionality: mono:plus
Cluster 69	Location: CM035916.1	Coordinates: 98333244-98338809	Size [bp]: 5566	Hits (absolute): 583	Hits (normalized): 115.451225252895	Hits (normalized) per kb: 20.7422552945571	Normalized hits with 1T: 97%	Normalized hits with 10A: 16.4%	Normalized hits 25-32 nt: 99.9%	Normalized hits on the main strand(s): 99.7%	Predicted directionality: mono:minus
Cluster 70	Location: CM035916.1	Coordinates: 101992261-102000633	Size [bp]: 8373	Hits (absolute): 1326	Hits (normalized): 148.810718564366	Hits (normalized) per kb: 17.7725860868419	Normalized hits with 1T: 83.4%	Normalized hits with 10A: 19.9%	Normalized hits 25-32 nt: 99.5%	Normalized hits on the main strand(s): 82.7%	Predicted directionality: mono:plus
Cluster 71	Location: CM035916.1	Coordinates: 102106048-102107332	Size [bp]: 1285	Hits (absolute): 151	Hits (normalized): 151.566486162771	Hits (normalized) per kb: 117.950871774737	Normalized hits with 1T: 0.2%	Normalized hits with 10A: 96.3%	Normalized hits 25-32 nt: 99.7%	Normalized hits on the main strand(s): 100%	Predicted directionality: mono:minus
Cluster 72	Location: CM035916.1	Coordinates: 102789884-102795977	Size [bp]: 6094	Hits (absolute): 1607	Hits (normalized): 107.345922782753	Hits (normalized) per kb: 17.6153313041316	Normalized hits with 1T: 85.5%	Normalized hits with 10A: 35.9%	Normalized hits 25-32 nt: 99.3%	Normalized hits on the main strand(s): 84.7%	Predicted directionality: mono:minus
Cluster 73	Location: CM035916.1	Coordinates: 117906424-117909569	Size [bp]: 3146	Hits (absolute): 431	Hits (normalized): 90.3090494000434	Hits (normalized) per kb: 28.7063364011505	Normalized hits with 1T: 96.7%	Normalized hits with 10A: 93.5%	Normalized hits 25-32 nt: 99.8%	Normalized hits on the main strand(s): 95.1%	Predicted directionality: mono:plus
Cluster 74	Location: CM035916.1	Coordinates: 118291172-118299022	Size [bp]: 7851	Hits (absolute): 2181	Hits (normalized): 148.216476466432	Hits (normalized) per kb: 18.8789608469768	Normalized hits with 1T: 96.4%	Normalized hits with 10A: 21.5%	Normalized hits 25-32 nt: 99%	Normalized hits on the main strand(s): 99.7%	Predicted directionality: mono:plus
Cluster 75	Location: CM035916.1	Coordinates: 121061004-121072969	Size [bp]: 11966	Hits (absolute): 2304	Hits (normalized): 500.186731058752	Hits (normalized) per kb: 41.8004179748255	Normalized hits with 1T: 95.5%	Normalized hits with 10A: 8.8%	Normalized hits 25-32 nt: 99.6%	Normalized hits on the main strand(s): 100%	Predicted directionality: mono:minus
Cluster 76	Location: CM035916.1	Coordinates: 121084092-121090583	Size [bp]: 6492	Hits (absolute): 1088	Hits (normalized): 118.530691873638	Hits (normalized) per kb: 18.2576297277349	Normalized hits with 1T: 96.2%	Normalized hits with 10A: 46.9%	Normalized hits 25-32 nt: 99.3%	Normalized hits on the main strand(s): 100%	Predicted directionality: mono:minus
Cluster 77	Location: CM035916.1	Coordinates: 121097396-121119023	Size [bp]: 21628	Hits (absolute): 8179	Hits (normalized): 1958.12544280399	Hits (normalized) per kb: 90.5368202323919	Normalized hits with 1T: 87.5%	Normalized hits with 10A: 26.5%	Normalized hits 25-32 nt: 99.5%	Normalized hits on the main strand(s): 100%	Predicted directionality: mono:minus
Cluster 78	Location: CM035916.1	Coordinates: 128960295-128963022	Size [bp]: 2728	Hits (absolute): 344	Hits (normalized): 307.579871329105	Hits (normalized) per kb: 112.748883562681	Normalized hits with 1T: 99.3%	Normalized hits with 10A: 99.3%	Normalized hits 25-32 nt: 99%	Normalized hits on the main strand(s): 100%	Predicted directionality: mono:minus
Cluster 79	Location: CM035916.1	Coordinates: 137460790-137466737	Size [bp]: 5948	Hits (absolute): 1040	Hits (normalized): 167.059733588311	Hits (normalized) per kb: 28.0864031021993	Normalized hits with 1T: 79.2%	Normalized hits with 10A: 26%	Normalized hits 25-32 nt: 99.1%	Normalized hits on the main strand(s): 86.2%	Predicted directionality: bi:minus-plus (split between 137462577 and 137462578)
Cluster 80	Location: CM035916.1	Coordinates: 141459821-141461779	Size [bp]: 1959	Hits (absolute): 490	Hits (normalized): 235.165791166389	Hits (normalized) per kb: 120.044107660147	Normalized hits with 1T: 48%	Normalized hits with 10A: 98.9%	Normalized hits 25-32 nt: 98.9%	Normalized hits on the main strand(s): 100%	Predicted directionality: mono:minus
Cluster 81	Location: CM035916.1	Coordinates: 149132323-149136510	Size [bp]: 4188	Hits (absolute): 309	Hits (normalized): 163.491527247312	Hits (normalized) per kb: 39.0383250802878	Normalized hits with 1T: 0.5%	Normalized hits with 10A: 98.7%	Normalized hits 25-32 nt: 99.8%	Normalized hits on the main strand(s): 100%	Predicted directionality: mono:plus
Cluster 82	Location: CM035916.1	Coordinates: 149902876-149904222	Size [bp]: 1347	Hits (absolute): 715	Hits (normalized): 123.663763303064	Hits (normalized) per kb: 91.8067399665455	Normalized hits with 1T: 38.1%	Normalized hits with 10A: 86%	Normalized hits 25-32 nt: 97.6%	Normalized hits on the main strand(s): 99.7%	Predicted directionality: mono:plus
Cluster 83	Location: CM035916.1	Coordinates: 150059620-150060909	Size [bp]: 1290	Hits (absolute): 645	Hits (normalized): 105.540091353194	Hits (normalized) per kb: 81.813722707917	Normalized hits with 1T: 27.9%	Normalized hits with 10A: 75.9%	Normalized hits 25-32 nt: 97.9%	Normalized hits on the main strand(s): 99.6%	Predicted directionality: mono:plus
Cluster 84	Location: CM035916.1	Coordinates: 152408602-152415700	Size [bp]: 7099	Hits (absolute): 1092	Hits (normalized): 188.142534460395	Hits (normalized) per kb: 26.5026727127706	Normalized hits with 1T: 96.4%	Normalized hits with 10A: 12.8%	Normalized hits 25-32 nt: 98.3%	Normalized hits on the main strand(s): 99.4%	Predicted directionality: mono:minus
Cluster 85	Location: CM035916.1	Coordinates: 152420024-152427723	Size [bp]: 7700	Hits (absolute): 662	Hits (normalized): 143.686128850771	Hits (normalized) per kb: 18.6602019714732	Normalized hits with 1T: 97.9%	Normalized hits with 10A: 3.2%	Normalized hits 25-32 nt: 97.9%	Normalized hits on the main strand(s): 99.8%	Predicted directionality: mono:minus
Cluster 86	Location: CM035916.1	Coordinates: 152449506-152450768	Size [bp]: 1263	Hits (absolute): 198	Hits (normalized): 96.2498282457439	Hits (normalized) per kb: 76.2070655216869	Normalized hits with 1T: 4.8%	Normalized hits with 10A: 98.5%	Normalized hits 25-32 nt: 99.5%	Normalized hits on the main strand(s): 100%	Predicted directionality: mono:minus
Cluster 87	Location: CM035917.1	Coordinates: 5362006-5367876	Size [bp]: 5871	Hits (absolute): 2014	Hits (normalized): 101.830125679981	Hits (normalized) per kb: 17.3448530778699	Normalized hits with 1T: 88.2%	Normalized hits with 10A: 29.2%	Normalized hits 25-32 nt: 99.5%	Normalized hits on the main strand(s): 85%	Predicted directionality: mono:minus
Cluster 88	Location: CM035917.1	Coordinates: 7893255-7906640	Size [bp]: 13386	Hits (absolute): 2503	Hits (normalized): 453.318629087264	Hits (normalized) per kb: 33.8649921841927	Normalized hits with 1T: 89.3%	Normalized hits with 10A: 21.8%	Normalized hits 25-32 nt: 99.8%	Normalized hits on the main strand(s): 98.5%	Predicted directionality: mono:plus
Cluster 89	Location: CM035917.1	Coordinates: 7937286-7947969	Size [bp]: 10684	Hits (absolute): 1754	Hits (normalized): 206.219458778126	Hits (normalized) per kb: 19.3018014849311	Normalized hits with 1T: 96%	Normalized hits with 10A: 22.2%	Normalized hits 25-32 nt: 99.2%	Normalized hits on the main strand(s): 99.9%	Predicted directionality: mono:plus
Cluster 90	Location: CM035917.1	Coordinates: 9946416-9954984	Size [bp]: 8569	Hits (absolute): 744	Hits (normalized): 114.158660075659	Hits (normalized) per kb: 13.322625191214	Normalized hits with 1T: 99.1%	Normalized hits with 10A: 60.7%	Normalized hits 25-32 nt: 99.9%	Normalized hits on the main strand(s): 100%	Predicted directionality: mono:minus
Cluster 91	Location: CM035917.1	Coordinates: 9976006-9990997	Size [bp]: 14992	Hits (absolute): 2624	Hits (normalized): 395.62919349572	Hits (normalized) per kb: 26.3894492692192	Normalized hits with 1T: 87.1%	Normalized hits with 10A: 21%	Normalized hits 25-32 nt: 99.9%	Normalized hits on the main strand(s): 100%	Predicted directionality: mono:minus
Cluster 92	Location: CM035917.1	Coordinates: 10484051-10493028	Size [bp]: 8978	Hits (absolute): 1036	Hits (normalized): 166.550004219517	Hits (normalized) per kb: 18.551171988794	Normalized hits with 1T: 82.3%	Normalized hits with 10A: 33.8%	Normalized hits 25-32 nt: 99.9%	Normalized hits on the main strand(s): 99.9%	Predicted directionality: mono:plus
Cluster 93	Location: CM035917.1	Coordinates: 10520026-10525911	Size [bp]: 5886	Hits (absolute): 537	Hits (normalized): 101.474025512874	Hits (normalized) per kb: 17.2400165560631	Normalized hits with 1T: 99.4%	Normalized hits with 10A: 67.5%	Normalized hits 25-32 nt: 99.9%	Normalized hits on the main strand(s): 100%	Predicted directionality: mono:plus
Cluster 94	Location: CM035917.1	Coordinates: 28530949-28537014	Size [bp]: 6066	Hits (absolute): 2131	Hits (normalized): 124.166571061308	Hits (normalized) per kb: 20.4689814277139	Normalized hits with 1T: 94.3%	Normalized hits with 10A: 28.3%	Normalized hits 25-32 nt: 98.9%	Normalized hits on the main strand(s): 99.8%	Predicted directionality: mono:plus
Cluster 95	Location: CM035917.1	Coordinates: 35134386-35140407	Size [bp]: 6022	Hits (absolute): 798	Hits (normalized): 213.865381237517	Hits (normalized) per kb: 35.5137212171417	Normalized hits with 1T: 12.6%	Normalized hits with 10A: 90.5%	Normalized hits 25-32 nt: 99.1%	Normalized hits on the main strand(s): 92.8%	Predicted directionality: bi:plus-minus (split between 35138339 and 35138340)
Cluster 96	Location: CM035917.1	Coordinates: 38349101-38354182	Size [bp]: 5082	Hits (absolute): 1155	Hits (normalized): 116.364793153115	Hits (normalized) per kb: 22.8976941829059	Normalized hits with 1T: 92.2%	Normalized hits with 10A: 32.2%	Normalized hits 25-32 nt: 99.7%	Normalized hits on the main strand(s): 92.5%	Predicted directionality: mono:minus
Cluster 97	Location: CM035917.1	Coordinates: 40563791-40571028	Size [bp]: 7238	Hits (absolute): 2019	Hits (normalized): 149.863857845039	Hits (normalized) per kb: 20.705213056852	Normalized hits with 1T: 89.5%	Normalized hits with 10A: 18.1%	Normalized hits 25-32 nt: 99.3%	Normalized hits on the main strand(s): 92.5%	Predicted directionality: mono:minus
Cluster 98	Location: CM035917.1	Coordinates: 46555002-46560920	Size [bp]: 5919	Hits (absolute): 1815	Hits (normalized): 234.118481911877	Hits (normalized) per kb: 39.5534218574321	Normalized hits with 1T: 91.5%	Normalized hits with 10A: 70.2%	Normalized hits 25-32 nt: 99.5%	Normalized hits on the main strand(s): 91.8%	Predicted directionality: mono:plus
Cluster 99	Location: CM035917.1	Coordinates: 56350242-56383026	Size [bp]: 32785	Hits (absolute): 8865	Hits (normalized): 2158.23163333947	Hits (normalized) per kb: 65.8296476830999	Normalized hits with 1T: 95.2%	Normalized hits with 10A: 22.4%	Normalized hits 25-32 nt: 99.8%	Normalized hits on the main strand(s): 99.9%	Predicted directionality: mono:minus
Cluster 100	Location: CM035917.1	Coordinates: 56420042-56425987	Size [bp]: 5946	Hits (absolute): 702	Hits (normalized): 112.809184820985	Hits (normalized) per kb: 18.9726148064576	Normalized hits with 1T: 95.2%	Normalized hits with 10A: 23.3%	Normalized hits 25-32 nt: 99.6%	Normalized hits on the main strand(s): 99.7%	Predicted directionality: mono:minus
Cluster 101	Location: CM035917.1	Coordinates: 56428195-56435986	Size [bp]: 7792	Hits (absolute): 1120	Hits (normalized): 143.351456740079	Hits (normalized) per kb: 18.3974117568107	Normalized hits with 1T: 98.3%	Normalized hits with 10A: 30%	Normalized hits 25-32 nt: 99.9%	Normalized hits on the main strand(s): 99.9%	Predicted directionality: mono:minus
Cluster 102	Location: CM035917.1	Coordinates: 56445058-56486670	Size [bp]: 41613	Hits (absolute): 12881	Hits (normalized): 3139.56852765734	Hits (normalized) per kb: 75.4466512835146	Normalized hits with 1T: 80.1%	Normalized hits with 10A: 25%	Normalized hits 25-32 nt: 99.7%	Normalized hits on the main strand(s): 99.3%	Predicted directionality: mono:minus
Cluster 103	Location: CM035917.1	Coordinates: 56506005-56515029	Size [bp]: 9025	Hits (absolute): 1046	Hits (normalized): 243.776729583066	Hits (normalized) per kb: 27.0114792986065	Normalized hits with 1T: 95.2%	Normalized hits with 10A: 32.3%	Normalized hits 25-32 nt: 99.5%	Normalized hits on the main strand(s): 99.9%	Predicted directionality: mono:minus
Cluster 104	Location: CM035917.1	Coordinates: 67205525-67210748	Size [bp]: 5224	Hits (absolute): 1742	Hits (normalized): 231.988937696271	Hits (normalized) per kb: 44.4080517272345	Normalized hits with 1T: 92.2%	Normalized hits with 10A: 70.3%	Normalized hits 25-32 nt: 99.5%	Normalized hits on the main strand(s): 92.4%	Predicted directionality: mono:plus
Cluster 105	Location: CM035917.1	Coordinates: 67390484-67395706	Size [bp]: 5223	Hits (absolute): 1658	Hits (normalized): 231.18178026379	Hits (normalized) per kb: 44.2619795068503	Normalized hits with 1T: 92.2%	Normalized hits with 10A: 70.6%	Normalized hits 25-32 nt: 99.6%	Normalized hits on the main strand(s): 92.5%	Predicted directionality: mono:minus
Cluster 106	Location: CM035917.1	Coordinates: 73531012-73538955	Size [bp]: 7944	Hits (absolute): 1173	Hits (normalized): 150.24046229512	Hits (normalized) per kb: 18.912508533955	Normalized hits with 1T: 94.3%	Normalized hits with 10A: 15.8%	Normalized hits 25-32 nt: 99.9%	Normalized hits on the main strand(s): 99.9%	Predicted directionality: mono:minus
Cluster 107	Location: CM035917.1	Coordinates: 73549001-73558984	Size [bp]: 9984	Hits (absolute): 1221	Hits (normalized): 247.943552386314	Hits (normalized) per kb: 24.834374195751	Normalized hits with 1T: 97.2%	Normalized hits with 10A: 16.5%	Normalized hits 25-32 nt: 99.8%	Normalized hits on the main strand(s): 100%	Predicted directionality: mono:minus
Cluster 108	Location: CM035917.1	Coordinates: 73611073-73619016	Size [bp]: 7944	Hits (absolute): 858	Hits (normalized): 156.179030644096	Hits (normalized) per kb: 19.6603423895105	Normalized hits with 1T: 96.8%	Normalized hits with 10A: 41%	Normalized hits 25-32 nt: 99.8%	Normalized hits on the main strand(s): 98.6%	Predicted directionality: mono:minus
Cluster 109	Location: CM035917.1	Coordinates: 73636014-73644028	Size [bp]: 8015	Hits (absolute): 1791	Hits (normalized): 125.028503335102	Hits (normalized) per kb: 15.5989755347133	Normalized hits with 1T: 77.2%	Normalized hits with 10A: 33.9%	Normalized hits 25-32 nt: 99.3%	Normalized hits on the main strand(s): 85.9%	Predicted directionality: mono:minus
Cluster 110	Location: CM035917.1	Coordinates: 74286027-74294955	Size [bp]: 8929	Hits (absolute): 1000	Hits (normalized): 360.692888176293	Hits (normalized) per kb: 40.3956085826138	Normalized hits with 1T: 98.5%	Normalized hits with 10A: 7.5%	Normalized hits 25-32 nt: 99.9%	Normalized hits on the main strand(s): 99.3%	Predicted directionality: mono:minus
Cluster 111	Location: CM035917.1	Coordinates: 76720093-76725410	Size [bp]: 5318	Hits (absolute): 322	Hits (normalized): 166.274215308261	Hits (normalized) per kb: 31.2664442636737	Normalized hits with 1T: 9.4%	Normalized hits with 10A: 95.7%	Normalized hits 25-32 nt: 99.3%	Normalized hits on the main strand(s): 99.3%	Predicted directionality: mono:minus
Cluster 112	Location: CM035917.1	Coordinates: 87979367-87992016	Size [bp]: 12650	Hits (absolute): 4473	Hits (normalized): 1774.40059724306	Hits (normalized) per kb: 140.269169447123	Normalized hits with 1T: 99.1%	Normalized hits with 10A: 16.2%	Normalized hits 25-32 nt: 99.6%	Normalized hits on the main strand(s): 100%	Predicted directionality: mono:plus
Cluster 113	Location: CM035917.1	Coordinates: 94417008-94422160	Size [bp]: 5153	Hits (absolute): 441	Hits (normalized): 91.8649165808859	Hits (normalized) per kb: 17.8277999883268	Normalized hits with 1T: 96.4%	Normalized hits with 10A: 18.4%	Normalized hits 25-32 nt: 99.9%	Normalized hits on the main strand(s): 99.6%	Predicted directionality: mono:minus
Cluster 114	Location: CM035917.1	Coordinates: 111049010-111063020	Size [bp]: 14011	Hits (absolute): 2430	Hits (normalized): 451.465964122355	Hits (normalized) per kb: 32.2225533425521	Normalized hits with 1T: 98.8%	Normalized hits with 10A: 27.9%	Normalized hits 25-32 nt: 99.6%	Normalized hits on the main strand(s): 99.9%	Predicted directionality: mono:minus
Cluster 115	Location: CM035917.1	Coordinates: 111091007-111117903	Size [bp]: 26897	Hits (absolute): 8891	Hits (normalized): 2343.63846730589	Hits (normalized) per kb: 87.1338267345417	Normalized hits with 1T: 94%	Normalized hits with 10A: 23%	Normalized hits 25-32 nt: 99.4%	Normalized hits on the main strand(s): 92.1%	Predicted directionality: mono:minus
Cluster 116	Location: CM035917.1	Coordinates: 111135024-111143619	Size [bp]: 8596	Hits (absolute): 1507	Hits (normalized): 171.36415809634	Hits (normalized) per kb: 19.9350140766444	Normalized hits with 1T: 79.3%	Normalized hits with 10A: 23.1%	Normalized hits 25-32 nt: 99.7%	Normalized hits on the main strand(s): 99%	Predicted directionality: mono:minus
Cluster 117	Location: CM035917.1	Coordinates: 111151085-111156994	Size [bp]: 5910	Hits (absolute): 871	Hits (normalized): 121.366019848427	Hits (normalized) per kb: 20.5353778915249	Normalized hits with 1T: 96.5%	Normalized hits with 10A: 26.1%	Normalized hits 25-32 nt: 99.8%	Normalized hits on the main strand(s): 99.2%	Predicted directionality: mono:minus
Cluster 118	Location: CM035917.1	Coordinates: 111398137-111400023	Size [bp]: 1887	Hits (absolute): 159	Hits (normalized): 104.844048531488	Hits (normalized) per kb: 55.5612598271921	Normalized hits with 1T: 98.4%	Normalized hits with 10A: 98.4%	Normalized hits 25-32 nt: 99.6%	Normalized hits on the main strand(s): 100%	Predicted directionality: mono:plus
Cluster 119	Location: CM035917.1	Coordinates: 111707398-111712940	Size [bp]: 5543	Hits (absolute): 762	Hits (normalized): 107.673337798054	Hits (normalized) per kb: 19.4248096705178	Normalized hits with 1T: 96.9%	Normalized hits with 10A: 26.2%	Normalized hits 25-32 nt: 99.7%	Normalized hits on the main strand(s): 99.4%	Predicted directionality: mono:plus
Cluster 120	Location: CM035917.1	Coordinates: 111717002-111725909	Size [bp]: 8908	Hits (absolute): 1822	Hits (normalized): 232.723165479432	Hits (normalized) per kb: 26.125261234266	Normalized hits with 1T: 83.4%	Normalized hits with 10A: 19.7%	Normalized hits 25-32 nt: 99.3%	Normalized hits on the main strand(s): 99.1%	Predicted directionality: mono:plus
Cluster 121	Location: CM035917.1	Coordinates: 111733032-111741972	Size [bp]: 8941	Hits (absolute): 1529	Hits (normalized): 155.399647441799	Hits (normalized) per kb: 17.3804974952843	Normalized hits with 1T: 95.1%	Normalized hits with 10A: 18.9%	Normalized hits 25-32 nt: 98.2%	Normalized hits on the main strand(s): 99.7%	Predicted directionality: mono:plus
Cluster 122	Location: CM035917.1	Coordinates: 111765006-111778951	Size [bp]: 13946	Hits (absolute): 2413	Hits (normalized): 432.576721034953	Hits (normalized) per kb: 31.0176322519188	Normalized hits with 1T: 97.6%	Normalized hits with 10A: 28.4%	Normalized hits 25-32 nt: 99.6%	Normalized hits on the main strand(s): 99.8%	Predicted directionality: mono:plus
Cluster 123	Location: CM035917.1	Coordinates: 111787435-111796026	Size [bp]: 8592	Hits (absolute): 2010	Hits (normalized): 165.812444481386	Hits (normalized) per kb: 19.2983069342042	Normalized hits with 1T: 93.3%	Normalized hits with 10A: 28.4%	Normalized hits 25-32 nt: 99.9%	Normalized hits on the main strand(s): 93.1%	Predicted directionality: mono:plus
Cluster 124	Location: CM035917.1	Coordinates: 113770060-113785001	Size [bp]: 14942	Hits (absolute): 12000	Hits (normalized): 11856.6122315431	Hits (normalized) per kb: 793.509031376306	Normalized hits with 1T: 78.1%	Normalized hits with 10A: 75.5%	Normalized hits 25-32 nt: 99.8%	Normalized hits on the main strand(s): 100%	Predicted directionality: mono:plus
Cluster 125	Location: CM035917.1	Coordinates: 117908736-117914344	Size [bp]: 5609	Hits (absolute): 768	Hits (normalized): 160.209197525229	Hits (normalized) per kb: 28.5630598213478	Normalized hits with 1T: 96.6%	Normalized hits with 10A: 13.4%	Normalized hits 25-32 nt: 98.2%	Normalized hits on the main strand(s): 99.9%	Predicted directionality: mono:plus
Cluster 126	Location: CM035917.1	Coordinates: 119906338-119914834	Size [bp]: 8497	Hits (absolute): 1264	Hits (normalized): 288.86429462553	Hits (normalized) per kb: 33.9956883813786	Normalized hits with 1T: 97.9%	Normalized hits with 10A: 11.2%	Normalized hits 25-32 nt: 99.8%	Normalized hits on the main strand(s): 98.9%	Predicted directionality: mono:plus
Cluster 127	Location: CM035917.1	Coordinates: 121387005-121462785	Size [bp]: 75781	Hits (absolute): 26670	Hits (normalized): 6991.27020642531	Hits (normalized) per kb: 92.2561391900242	Normalized hits with 1T: 96.3%	Normalized hits with 10A: 27%	Normalized hits 25-32 nt: 99.7%	Normalized hits on the main strand(s): 100%	Predicted directionality: mono:plus
Cluster 128	Location: CM035917.1	Coordinates: 127665105-127677420	Size [bp]: 12316	Hits (absolute): 1691	Hits (normalized): 385.116059528334	Hits (normalized) per kb: 31.2692399042552	Normalized hits with 1T: 90.4%	Normalized hits with 10A: 46.5%	Normalized hits 25-32 nt: 99.7%	Normalized hits on the main strand(s): 94%	Predicted directionality: mono:plus
Cluster 129	Location: CM035917.1	Coordinates: 127679089-127688002	Size [bp]: 8914	Hits (absolute): 1255	Hits (normalized): 614.599310098286	Hits (normalized) per kb: 68.9474858416355	Normalized hits with 1T: 99.3%	Normalized hits with 10A: 9.9%	Normalized hits 25-32 nt: 99.6%	Normalized hits on the main strand(s): 100%	Predicted directionality: mono:plus
Cluster 130	Location: CM035917.1	Coordinates: 127773016-127779889	Size [bp]: 6874	Hits (absolute): 727	Hits (normalized): 226.744247484427	Hits (normalized) per kb: 32.9857632213059	Normalized hits with 1T: 89.5%	Normalized hits with 10A: 57.7%	Normalized hits 25-32 nt: 99.7%	Normalized hits on the main strand(s): 96.5%	Predicted directionality: bi:plus-minus (split between 127777946 and 127777947)
Cluster 131	Location: CM035917.1	Coordinates: 127793032-127801000	Size [bp]: 7969	Hits (absolute): 1037	Hits (normalized): 143.810238473613	Hits (normalized) per kb: 18.045859953685	Normalized hits with 1T: 81.7%	Normalized hits with 10A: 25.7%	Normalized hits 25-32 nt: 99.5%	Normalized hits on the main strand(s): 82.2%	Predicted directionality: mono:plus
Cluster 132	Location: CM035917.1	Coordinates: 127814158-127823011	Size [bp]: 8854	Hits (absolute): 1418	Hits (normalized): 655.68866352728	Hits (normalized) per kb: 74.0558200942104	Normalized hits with 1T: 99.4%	Normalized hits with 10A: 11.1%	Normalized hits 25-32 nt: 99.6%	Normalized hits on the main strand(s): 99.8%	Predicted directionality: mono:plus
Cluster 133	Location: CM035917.1	Coordinates: 132413678-132421024	Size [bp]: 7347	Hits (absolute): 1305	Hits (normalized): 148.080246469685	Hits (normalized) per kb: 20.1551707724388	Normalized hits with 1T: 82.9%	Normalized hits with 10A: 20.3%	Normalized hits 25-32 nt: 99.4%	Normalized hits on the main strand(s): 82.1%	Predicted directionality: mono:plus
Cluster 134	Location: CM035917.1	Coordinates: 138004011-138009509	Size [bp]: 5499	Hits (absolute): 557	Hits (normalized): 114.375249110424	Hits (normalized) per kb: 20.7995659264782	Normalized hits with 1T: 96.8%	Normalized hits with 10A: 16.7%	Normalized hits 25-32 nt: 99.9%	Normalized hits on the main strand(s): 99.6%	Predicted directionality: mono:minus
Cluster 135	Location: CM035917.1	Coordinates: 143548022-143552617	Size [bp]: 4596	Hits (absolute): 1314	Hits (normalized): 97.2041016006253	Hits (normalized) per kb: 21.1497199093131	Normalized hits with 1T: 85.4%	Normalized hits with 10A: 33.1%	Normalized hits 25-32 nt: 99.6%	Normalized hits on the main strand(s): 86.4%	Predicted directionality: mono:plus
Cluster 136	Location: CM035917.1	Coordinates: 151653342-151660924	Size [bp]: 7583	Hits (absolute): 1321	Hits (normalized): 129.511803675565	Hits (normalized) per kb: 17.0792672226259	Normalized hits with 1T: 94.6%	Normalized hits with 10A: 15.8%	Normalized hits 25-32 nt: 99.7%	Normalized hits on the main strand(s): 94.1%	Predicted directionality: bi:plus-minus (split between 151659000 and 151659006)
Cluster 137	Location: CM035918.1	Coordinates: 13557242-13560377	Size [bp]: 3136	Hits (absolute): 259	Hits (normalized): 145.652924219131	Hits (normalized) per kb: 46.4453748010142	Normalized hits with 1T: 95.2%	Normalized hits with 10A: 55.8%	Normalized hits 25-32 nt: 99.5%	Normalized hits on the main strand(s): 99.9%	Predicted directionality: mono:minus
Cluster 138	Location: CM035918.1	Coordinates: 20662002-20710962	Size [bp]: 48961	Hits (absolute): 16181	Hits (normalized): 2601.04055794127	Hits (normalized) per kb: 53.124859060401	Normalized hits with 1T: 95.4%	Normalized hits with 10A: 16%	Normalized hits 25-32 nt: 99.7%	Normalized hits on the main strand(s): 99.6%	Predicted directionality: mono:plus
Cluster 139	Location: CM035918.1	Coordinates: 20720018-20751874	Size [bp]: 31857	Hits (absolute): 7631	Hits (normalized): 1877.45787721996	Hits (normalized) per kb: 58.934200188791	Normalized hits with 1T: 92.9%	Normalized hits with 10A: 28.3%	Normalized hits 25-32 nt: 99.7%	Normalized hits on the main strand(s): 99.2%	Predicted directionality: mono:plus
Cluster 140	Location: CM035918.1	Coordinates: 31709000-31718022	Size [bp]: 9023	Hits (absolute): 1104	Hits (normalized): 173.891284169471	Hits (normalized) per kb: 19.2717483486798	Normalized hits with 1T: 97.2%	Normalized hits with 10A: 14.7%	Normalized hits 25-32 nt: 99.7%	Normalized hits on the main strand(s): 98.9%	Predicted directionality: mono:plus
Cluster 141	Location: CM035918.1	Coordinates: 31720059-31728899	Size [bp]: 8841	Hits (absolute): 1986	Hits (normalized): 318.28798432335	Hits (normalized) per kb: 36.0015604986162	Normalized hits with 1T: 79.6%	Normalized hits with 10A: 22.1%	Normalized hits 25-32 nt: 99.8%	Normalized hits on the main strand(s): 99.9%	Predicted directionality: mono:plus
Cluster 142	Location: CM035918.1	Coordinates: 31734012-31745029	Size [bp]: 11018	Hits (absolute): 3693	Hits (normalized): 529.632698400682	Hits (normalized) per kb: 48.0696419788749	Normalized hits with 1T: 96.1%	Normalized hits with 10A: 8.4%	Normalized hits 25-32 nt: 99.8%	Normalized hits on the main strand(s): 98.4%	Predicted directionality: mono:plus
Cluster 143	Location: CM035918.1	Coordinates: 31765193-31770923	Size [bp]: 5731	Hits (absolute): 1510	Hits (normalized): 137.502333152842	Hits (normalized) per kb: 23.9928863807147	Normalized hits with 1T: 95.9%	Normalized hits with 10A: 22.6%	Normalized hits 25-32 nt: 99.5%	Normalized hits on the main strand(s): 96.3%	Predicted directionality: mono:plus
Cluster 144	Location: CM035918.1	Coordinates: 32458288-32465938	Size [bp]: 7651	Hits (absolute): 2174	Hits (normalized): 158.537960363799	Hits (normalized) per kb: 20.7212879901957	Normalized hits with 1T: 92.1%	Normalized hits with 10A: 32.3%	Normalized hits 25-32 nt: 99.3%	Normalized hits on the main strand(s): 95.2%	Predicted directionality: bi:plus-minus (split between 32460534 and 32460535)
Cluster 145	Location: CM035918.1	Coordinates: 32573188-32581008	Size [bp]: 7821	Hits (absolute): 1682	Hits (normalized): 154.198882777191	Hits (normalized) per kb: 19.7162552011408	Normalized hits with 1T: 96%	Normalized hits with 10A: 21.6%	Normalized hits 25-32 nt: 99.5%	Normalized hits on the main strand(s): 96.6%	Predicted directionality: mono:minus
Cluster 146	Location: CM035918.1	Coordinates: 32613007-32622025	Size [bp]: 9019	Hits (absolute): 1204	Hits (normalized): 182.942125600181	Hits (normalized) per kb: 20.2837702391885	Normalized hits with 1T: 98.4%	Normalized hits with 10A: 13.6%	Normalized hits 25-32 nt: 99.8%	Normalized hits on the main strand(s): 99.9%	Predicted directionality: mono:minus
Cluster 147	Location: CM035918.1	Coordinates: 34382090-34383895	Size [bp]: 1806	Hits (absolute): 138	Hits (normalized): 91.3989060558611	Hits (normalized) per kb: 50.6087825370367	Normalized hits with 1T: 95.1%	Normalized hits with 10A: 0.6%	Normalized hits 25-32 nt: 99.6%	Normalized hits on the main strand(s): 100%	Predicted directionality: mono:minus
Cluster 148	Location: CM035918.1	Coordinates: 53076352-53077940	Size [bp]: 1589	Hits (absolute): 209	Hits (normalized): 137.587451219304	Hits (normalized) per kb: 86.5872790008554	Normalized hits with 1T: 92.7%	Normalized hits with 10A: 0%	Normalized hits 25-32 nt: 99.5%	Normalized hits on the main strand(s): 100%	Predicted directionality: mono:plus
Cluster 149	Location: CM035918.1	Coordinates: 53358518-53376010	Size [bp]: 17493	Hits (absolute): 4237	Hits (normalized): 590.185812107908	Hits (normalized) per kb: 33.7384894478791	Normalized hits with 1T: 94.9%	Normalized hits with 10A: 25.2%	Normalized hits 25-32 nt: 99%	Normalized hits on the main strand(s): 99.9%	Predicted directionality: mono:plus
Cluster 150	Location: CM035918.1	Coordinates: 53377047-53428933	Size [bp]: 51887	Hits (absolute): 15658	Hits (normalized): 4755.33700518895	Hits (normalized) per kb: 91.6480873635445	Normalized hits with 1T: 94.5%	Normalized hits with 10A: 28.3%	Normalized hits 25-32 nt: 99.4%	Normalized hits on the main strand(s): 99.9%	Predicted directionality: mono:plus
Cluster 151	Location: CM035918.1	Coordinates: 53437012-53446014	Size [bp]: 9003	Hits (absolute): 856	Hits (normalized): 112.502131373104	Hits (normalized) per kb: 12.4958144892306	Normalized hits with 1T: 84.9%	Normalized hits with 10A: 43.5%	Normalized hits 25-32 nt: 99.8%	Normalized hits on the main strand(s): 100%	Predicted directionality: mono:plus
Cluster 152	Location: CM035918.1	Coordinates: 67585404-67590589	Size [bp]: 5186	Hits (absolute): 610	Hits (normalized): 132.733702248666	Hits (normalized) per kb: 25.5947884339233	Normalized hits with 1T: 97.6%	Normalized hits with 10A: 3.1%	Normalized hits 25-32 nt: 97.7%	Normalized hits on the main strand(s): 99.6%	Predicted directionality: mono:plus
Cluster 153	Location: CM035918.1	Coordinates: 68113030-68121901	Size [bp]: 8872	Hits (absolute): 1506	Hits (normalized): 147.271925928553	Hits (normalized) per kb: 16.5998148628959	Normalized hits with 1T: 95%	Normalized hits with 10A: 18.3%	Normalized hits 25-32 nt: 99.8%	Normalized hits on the main strand(s): 97.6%	Predicted directionality: mono:minus
Cluster 154	Location: CM035918.1	Coordinates: 68128028-68139026	Size [bp]: 10999	Hits (absolute): 1651	Hits (normalized): 1923.06247064916	Hits (normalized) per kb: 174.840060878149	Normalized hits with 1T: 99.2%	Normalized hits with 10A: 5.5%	Normalized hits 25-32 nt: 99.8%	Normalized hits on the main strand(s): 100%	Predicted directionality: mono:minus
Cluster 155	Location: CM035918.1	Coordinates: 68146136-68161907	Size [bp]: 15772	Hits (absolute): 4396	Hits (normalized): 3733.50078917343	Hits (normalized) per kb: 236.716672778985	Normalized hits with 1T: 98.3%	Normalized hits with 10A: 7.1%	Normalized hits 25-32 nt: 98.6%	Normalized hits on the main strand(s): 100%	Predicted directionality: mono:minus
Cluster 156	Location: CM035918.1	Coordinates: 68163007-68185971	Size [bp]: 22965	Hits (absolute): 3522	Hits (normalized): 1924.17283722678	Hits (normalized) per kb: 83.7874449584672	Normalized hits with 1T: 98.7%	Normalized hits with 10A: 3.3%	Normalized hits 25-32 nt: 99.9%	Normalized hits on the main strand(s): 100%	Predicted directionality: mono:minus
Cluster 157	Location: CM035918.1	Coordinates: 68190158-68210001	Size [bp]: 19844	Hits (absolute): 5891	Hits (normalized): 2704.40487744552	Hits (normalized) per kb: 136.283284888027	Normalized hits with 1T: 96.9%	Normalized hits with 10A: 14.4%	Normalized hits 25-32 nt: 99.7%	Normalized hits on the main strand(s): 100%	Predicted directionality: mono:minus
Cluster 158	Location: CM035918.1	Coordinates: 68218044-68237027	Size [bp]: 18984	Hits (absolute): 4909	Hits (normalized): 1410.93909898994	Hits (normalized) per kb: 74.3228037697452	Normalized hits with 1T: 93.7%	Normalized hits with 10A: 15.3%	Normalized hits 25-32 nt: 99.7%	Normalized hits on the main strand(s): 99.9%	Predicted directionality: mono:minus
Cluster 159	Location: CM035918.1	Coordinates: 84509934-84515010	Size [bp]: 5077	Hits (absolute): 547	Hits (normalized): 136.486909670767	Hits (normalized) per kb: 26.8835787420021	Normalized hits with 1T: 95.2%	Normalized hits with 10A: 13%	Normalized hits 25-32 nt: 99.8%	Normalized hits on the main strand(s): 95.6%	Predicted directionality: mono:minus
Cluster 160	Location: CM035918.1	Coordinates: 89547002-89567855	Size [bp]: 20854	Hits (absolute): 5433	Hits (normalized): 1794.29395741416	Hits (normalized) per kb: 86.040731267169	Normalized hits with 1T: 83.1%	Normalized hits with 10A: 34.9%	Normalized hits 25-32 nt: 99.8%	Normalized hits on the main strand(s): 99.9%	Predicted directionality: mono:plus
Cluster 161	Location: CM035918.1	Coordinates: 89575006-89597019	Size [bp]: 22014	Hits (absolute): 3933	Hits (normalized): 708.737432082872	Hits (normalized) per kb: 32.1945969367369	Normalized hits with 1T: 98.7%	Normalized hits with 10A: 33.4%	Normalized hits 25-32 nt: 99.5%	Normalized hits on the main strand(s): 99.6%	Predicted directionality: mono:plus
Cluster 162	Location: CM035918.1	Coordinates: 92929156-92934847	Size [bp]: 5692	Hits (absolute): 1810	Hits (normalized): 136.841364639112	Hits (normalized) per kb: 24.0411111807459	Normalized hits with 1T: 96.7%	Normalized hits with 10A: 30.9%	Normalized hits 25-32 nt: 99.8%	Normalized hits on the main strand(s): 98.6%	Predicted directionality: mono:plus
Cluster 163	Location: CM035918.1	Coordinates: 103713021-103722027	Size [bp]: 9007	Hits (absolute): 1251	Hits (normalized): 156.000871184526	Hits (normalized) per kb: 17.3196923126363	Normalized hits with 1T: 89.3%	Normalized hits with 10A: 21.9%	Normalized hits 25-32 nt: 99.6%	Normalized hits on the main strand(s): 99.8%	Predicted directionality: mono:plus
Cluster 164	Location: CM035918.1	Coordinates: 111214302-111221593	Size [bp]: 7292	Hits (absolute): 1998	Hits (normalized): 157.178244523834	Hits (normalized) per kb: 21.5550877936329	Normalized hits with 1T: 92.7%	Normalized hits with 10A: 27.7%	Normalized hits 25-32 nt: 99.8%	Normalized hits on the main strand(s): 95.8%	Predicted directionality: bi:plus-minus (split between 111221037 and 111221041)
Cluster 165	Location: CM035918.1	Coordinates: 115281384-115286254	Size [bp]: 4871	Hits (absolute): 647	Hits (normalized): 136.223218083894	Hits (normalized) per kb: 27.9661905571942	Normalized hits with 1T: 91.2%	Normalized hits with 10A: 30.7%	Normalized hits 25-32 nt: 98.4%	Normalized hits on the main strand(s): 91.5%	Predicted directionality: bi:minus-plus (split between 115281795 and 115281796)
Cluster 166	Location: CM035918.1	Coordinates: 116572025-116577993	Size [bp]: 5969	Hits (absolute): 1941	Hits (normalized): 118.877927145884	Hits (normalized) per kb: 19.9161435027192	Normalized hits with 1T: 83.7%	Normalized hits with 10A: 36.8%	Normalized hits 25-32 nt: 99%	Normalized hits on the main strand(s): 80.4%	Predicted directionality: mono:minus
Cluster 167	Location: CM035918.1	Coordinates: 116622093-116628653	Size [bp]: 6561	Hits (absolute): 1828	Hits (normalized): 112.776532115218	Hits (normalized) per kb: 17.1889961154504	Normalized hits with 1T: 83.9%	Normalized hits with 10A: 38.6%	Normalized hits 25-32 nt: 99.2%	Normalized hits on the main strand(s): 78.3%	Predicted directionality: mono:plus
Cluster 168	Location: CM035918.1	Coordinates: 121468723-121473923	Size [bp]: 5201	Hits (absolute): 680	Hits (normalized): 145.709285573744	Hits (normalized) per kb: 28.0158131775161	Normalized hits with 1T: 93.9%	Normalized hits with 10A: 20.5%	Normalized hits 25-32 nt: 99.5%	Normalized hits on the main strand(s): 97.6%	Predicted directionality: mono:minus
Cluster 169	Location: CM035918.1	Coordinates: 121544057-121556008	Size [bp]: 11952	Hits (absolute): 1637	Hits (normalized): 259.92208484789	Hits (normalized) per kb: 21.7472880836121	Normalized hits with 1T: 96.5%	Normalized hits with 10A: 16.6%	Normalized hits 25-32 nt: 99.7%	Normalized hits on the main strand(s): 100%	Predicted directionality: mono:minus
Cluster 170	Location: CM035918.1	Coordinates: 137035719-137043467	Size [bp]: 7749	Hits (absolute): 3000	Hits (normalized): 189.461975954603	Hits (normalized) per kb: 24.4499736157926	Normalized hits with 1T: 92.4%	Normalized hits with 10A: 35%	Normalized hits 25-32 nt: 99.3%	Normalized hits on the main strand(s): 94.6%	Predicted directionality: bi:plus-minus (split between 137037278 and 137037284)
Cluster 171	Location: CM035918.1	Coordinates: 138166007-138170290	Size [bp]: 4284	Hits (absolute): 806	Hits (normalized): 266.950553020067	Hits (normalized) per kb: 62.3134307416983	Normalized hits with 1T: 98.7%	Normalized hits with 10A: 8.9%	Normalized hits 25-32 nt: 99.9%	Normalized hits on the main strand(s): 99.3%	Predicted directionality: mono:minus
Cluster 172	Location: CM035919.1	Coordinates: 12456004-12462173	Size [bp]: 6170	Hits (absolute): 2396	Hits (normalized): 116.578054337193	Hits (normalized) per kb: 18.8943368701751	Normalized hits with 1T: 94.3%	Normalized hits with 10A: 18.8%	Normalized hits 25-32 nt: 98.6%	Normalized hits on the main strand(s): 99.7%	Predicted directionality: mono:minus
Cluster 173	Location: CM035919.1	Coordinates: 15794429-15800217	Size [bp]: 5789	Hits (absolute): 1336	Hits (normalized): 266.350229973005	Hits (normalized) per kb: 46.0099537804431	Normalized hits with 1T: 65.8%	Normalized hits with 10A: 78.5%	Normalized hits 25-32 nt: 96.6%	Normalized hits on the main strand(s): 88.2%	Predicted directionality: mono:plus
Cluster 174	Location: CM035919.1	Coordinates: 16697080-16705627	Size [bp]: 8548	Hits (absolute): 947	Hits (normalized): 340.364837715581	Hits (normalized) per kb: 39.8183088025308	Normalized hits with 1T: 98.7%	Normalized hits with 10A: 51.1%	Normalized hits 25-32 nt: 99.9%	Normalized hits on the main strand(s): 99.9%	Predicted directionality: mono:minus
Cluster 175	Location: CM035919.1	Coordinates: 16709211-16719900	Size [bp]: 10690	Hits (absolute): 2288	Hits (normalized): 977.997723241618	Hits (normalized) per kb: 91.4873380301074	Normalized hits with 1T: 97.4%	Normalized hits with 10A: 60.8%	Normalized hits 25-32 nt: 99.8%	Normalized hits on the main strand(s): 99.4%	Predicted directionality: mono:minus
Cluster 176	Location: CM035919.1	Coordinates: 16736034-16745459	Size [bp]: 9426	Hits (absolute): 1834	Hits (normalized): 259.894420018315	Hits (normalized) per kb: 27.5720052352004	Normalized hits with 1T: 93.3%	Normalized hits with 10A: 11.5%	Normalized hits 25-32 nt: 99.7%	Normalized hits on the main strand(s): 99.8%	Predicted directionality: mono:minus
Cluster 177	Location: CM035919.1	Coordinates: 16771003-16779011	Size [bp]: 8009	Hits (absolute): 1136	Hits (normalized): 132.800791465521	Hits (normalized) per kb: 16.5816431991161	Normalized hits with 1T: 91.9%	Normalized hits with 10A: 27.2%	Normalized hits 25-32 nt: 99.8%	Normalized hits on the main strand(s): 99.8%	Predicted directionality: mono:minus
Cluster 178	Location: CM035919.1	Coordinates: 17369002-17377731	Size [bp]: 8730	Hits (absolute): 1355	Hits (normalized): 163.76009282382	Hits (normalized) per kb: 18.7580493918262	Normalized hits with 1T: 92.1%	Normalized hits with 10A: 22.8%	Normalized hits 25-32 nt: 99.9%	Normalized hits on the main strand(s): 99.9%	Predicted directionality: mono:plus
Cluster 179	Location: CM035919.1	Coordinates: 17416094-17424997	Size [bp]: 8904	Hits (absolute): 2202	Hits (normalized): 255.945649445923	Hits (normalized) per kb: 28.7447764591463	Normalized hits with 1T: 92.4%	Normalized hits with 10A: 13.4%	Normalized hits 25-32 nt: 99.7%	Normalized hits on the main strand(s): 97.5%	Predicted directionality: mono:plus
Cluster 180	Location: CM035919.1	Coordinates: 17426000-17434009	Size [bp]: 8010	Hits (absolute): 1514	Hits (normalized): 121.647569875769	Hits (normalized) per kb: 15.1866185489397	Normalized hits with 1T: 87.8%	Normalized hits with 10A: 31.1%	Normalized hits 25-32 nt: 99.2%	Normalized hits on the main strand(s): 86.9%	Predicted directionality: mono:plus
Cluster 181	Location: CM035919.1	Coordinates: 17457113-17476955	Size [bp]: 19843	Hits (absolute): 3585	Hits (normalized): 1412.03365198398	Hits (normalized) per kb: 71.1602353619054	Normalized hits with 1T: 97.8%	Normalized hits with 10A: 58.3%	Normalized hits 25-32 nt: 99.8%	Normalized hits on the main strand(s): 99.8%	Predicted directionality: mono:plus
Cluster 182	Location: CM035919.1	Coordinates: 23738368-23792023	Size [bp]: 53656	Hits (absolute): 16242	Hits (normalized): 5237.80369909376	Hits (normalized) per kb: 97.6181778253717	Normalized hits with 1T: 93%	Normalized hits with 10A: 31%	Normalized hits 25-32 nt: 99.7%	Normalized hits on the main strand(s): 99.9%	Predicted directionality: mono:minus
Cluster 183	Location: CM035919.1	Coordinates: 23806024-23823020	Size [bp]: 16997	Hits (absolute): 3155	Hits (normalized): 1044.68587535198	Hits (normalized) per kb: 61.4628570947721	Normalized hits with 1T: 94.3%	Normalized hits with 10A: 27.4%	Normalized hits 25-32 nt: 99.8%	Normalized hits on the main strand(s): 100%	Predicted directionality: mono:minus
Cluster 184	Location: CM035919.1	Coordinates: 36831073-36838600	Size [bp]: 7528	Hits (absolute): 1648	Hits (normalized): 148.754990496989	Hits (normalized) per kb: 19.7602865402997	Normalized hits with 1T: 96.6%	Normalized hits with 10A: 20.4%	Normalized hits 25-32 nt: 99.7%	Normalized hits on the main strand(s): 100%	Predicted directionality: mono:plus
Cluster 185	Location: CM035919.1	Coordinates: 36846019-36863805	Size [bp]: 17787	Hits (absolute): 3929	Hits (normalized): 914.646342409248	Hits (normalized) per kb: 51.4223139462578	Normalized hits with 1T: 94.1%	Normalized hits with 10A: 10.7%	Normalized hits 25-32 nt: 99.7%	Normalized hits on the main strand(s): 100%	Predicted directionality: mono:plus
Cluster 186	Location: CM035919.1	Coordinates: 36865012-36929020	Size [bp]: 64009	Hits (absolute): 10413	Hits (normalized): 1556.91548730441	Hits (normalized) per kb: 24.323470879479	Normalized hits with 1T: 91%	Normalized hits with 10A: 27.7%	Normalized hits 25-32 nt: 99.8%	Normalized hits on the main strand(s): 99.9%	Predicted directionality: mono:plus
Cluster 187	Location: CM035919.1	Coordinates: 41910046-41919024	Size [bp]: 8979	Hits (absolute): 1340	Hits (normalized): 264.155278829575	Hits (normalized) per kb: 29.419224749437	Normalized hits with 1T: 94.1%	Normalized hits with 10A: 7.6%	Normalized hits 25-32 nt: 99.6%	Normalized hits on the main strand(s): 99.9%	Predicted directionality: mono:minus
Cluster 188	Location: CM035919.1	Coordinates: 41923022-41936573	Size [bp]: 13552	Hits (absolute): 3537	Hits (normalized): 587.838301167567	Hits (normalized) per kb: 43.3764603526551	Normalized hits with 1T: 93.1%	Normalized hits with 10A: 36.4%	Normalized hits 25-32 nt: 99.8%	Normalized hits on the main strand(s): 100%	Predicted directionality: mono:minus
Cluster 189	Location: CM035919.1	Coordinates: 41942166-41979842	Size [bp]: 37677	Hits (absolute): 7671	Hits (normalized): 1243.02588605428	Hits (normalized) per kb: 32.991354502469	Normalized hits with 1T: 91.7%	Normalized hits with 10A: 17.1%	Normalized hits 25-32 nt: 99.5%	Normalized hits on the main strand(s): 99.9%	Predicted directionality: mono:minus
Cluster 190	Location: CM035919.1	Coordinates: 43759146-43764344	Size [bp]: 5199	Hits (absolute): 1940	Hits (normalized): 115.295019193091	Hits (normalized) per kb: 22.1764189128748	Normalized hits with 1T: 82.2%	Normalized hits with 10A: 36.5%	Normalized hits 25-32 nt: 99%	Normalized hits on the main strand(s): 76.7%	Predicted directionality: mono:minus
Cluster 191	Location: CM035919.1	Coordinates: 48814504-48818482	Size [bp]: 3979	Hits (absolute): 423	Hits (normalized): 126.438266925534	Hits (normalized) per kb: 31.7766486698003	Normalized hits with 1T: 98.8%	Normalized hits with 10A: 49.2%	Normalized hits 25-32 nt: 99.7%	Normalized hits on the main strand(s): 99.4%	Predicted directionality: mono:plus
Cluster 192	Location: CM035919.1	Coordinates: 50601070-50622019	Size [bp]: 20950	Hits (absolute): 7866	Hits (normalized): 2576.28766387942	Hits (normalized) per kb: 122.97324007943	Normalized hits with 1T: 95%	Normalized hits with 10A: 42.9%	Normalized hits 25-32 nt: 99.6%	Normalized hits on the main strand(s): 99.5%	Predicted directionality: mono:minus
Cluster 193	Location: CM035919.1	Coordinates: 50623095-50632028	Size [bp]: 8934	Hits (absolute): 1580	Hits (normalized): 170.799287911664	Hits (normalized) per kb: 19.1179881166964	Normalized hits with 1T: 93.6%	Normalized hits with 10A: 26.7%	Normalized hits 25-32 nt: 99.7%	Normalized hits on the main strand(s): 100%	Predicted directionality: mono:minus
Cluster 194	Location: CM035919.1	Coordinates: 58647799-58650910	Size [bp]: 3112	Hits (absolute): 346	Hits (normalized): 152.717387568134	Hits (normalized) per kb: 49.0739758577845	Normalized hits with 1T: 2.6%	Normalized hits with 10A: 93.3%	Normalized hits 25-32 nt: 99.8%	Normalized hits on the main strand(s): 100%	Predicted directionality: mono:plus
Cluster 195	Location: CM035919.1	Coordinates: 65960788-65963306	Size [bp]: 2519	Hits (absolute): 100	Hits (normalized): 136.239396915052	Hits (normalized) per kb: 54.0844626900063	Normalized hits with 1T: 99.7%	Normalized hits with 10A: 0.1%	Normalized hits 25-32 nt: 97.3%	Normalized hits on the main strand(s): 100%	Predicted directionality: mono:plus
Cluster 196	Location: CM035919.1	Coordinates: 67496046-67503538	Size [bp]: 7493	Hits (absolute): 1377	Hits (normalized): 118.315486760635	Hits (normalized) per kb: 15.7904769145471	Normalized hits with 1T: 92.4%	Normalized hits with 10A: 9.7%	Normalized hits 25-32 nt: 99.6%	Normalized hits on the main strand(s): 99.9%	Predicted directionality: mono:minus
Cluster 197	Location: CM035919.1	Coordinates: 67665009-67691007	Size [bp]: 25999	Hits (absolute): 3362	Hits (normalized): 837.19264906052	Hits (normalized) per kb: 32.2008871280454	Normalized hits with 1T: 95.4%	Normalized hits with 10A: 52.5%	Normalized hits 25-32 nt: 99.9%	Normalized hits on the main strand(s): 99.7%	Predicted directionality: mono:plus
Cluster 198	Location: CM035919.1	Coordinates: 67717013-67726014	Size [bp]: 9002	Hits (absolute): 975	Hits (normalized): 247.901274772654	Hits (normalized) per kb: 27.5384575482222	Normalized hits with 1T: 91.2%	Normalized hits with 10A: 5.3%	Normalized hits 25-32 nt: 97.5%	Normalized hits on the main strand(s): 99.9%	Predicted directionality: mono:plus
Cluster 199	Location: CM035919.1	Coordinates: 68792879-68797469	Size [bp]: 4591	Hits (absolute): 1000	Hits (normalized): 139.276542313149	Hits (normalized) per kb: 30.3368937703196	Normalized hits with 1T: 97.6%	Normalized hits with 10A: 5.9%	Normalized hits 25-32 nt: 99.8%	Normalized hits on the main strand(s): 100%	Predicted directionality: mono:plus
Cluster 200	Location: CM035919.1	Coordinates: 72963100-72968034	Size [bp]: 4935	Hits (absolute): 1049	Hits (normalized): 167.155146719749	Hits (normalized) per kb: 33.8712823755011	Normalized hits with 1T: 79.3%	Normalized hits with 10A: 25.9%	Normalized hits 25-32 nt: 99.2%	Normalized hits on the main strand(s): 86.2%	Predicted directionality: bi:minus-plus (split between 72964881 and 72964881)
Cluster 201	Location: CM035919.1	Coordinates: 73328068-73336993	Size [bp]: 8926	Hits (absolute): 1429	Hits (normalized): 153.60960693823	Hits (normalized) per kb: 17.2092645096664	Normalized hits with 1T: 83.6%	Normalized hits with 10A: 19.6%	Normalized hits 25-32 nt: 99.4%	Normalized hits on the main strand(s): 82.8%	Predicted directionality: mono:plus
Cluster 202	Location: CM035919.1	Coordinates: 82095867-82100983	Size [bp]: 5117	Hits (absolute): 1955	Hits (normalized): 803.930572515913	Hits (normalized) per kb: 157.10940940003	Normalized hits with 1T: 96.2%	Normalized hits with 10A: 17.8%	Normalized hits 25-32 nt: 99.6%	Normalized hits on the main strand(s): 96.5%	Predicted directionality: mono:minus
Cluster 203	Location: CM035919.1	Coordinates: 83820204-83823740	Size [bp]: 3537	Hits (absolute): 547	Hits (normalized): 123.754563379452	Hits (normalized) per kb: 34.9888396979621	Normalized hits with 1T: 99%	Normalized hits with 10A: 4.1%	Normalized hits 25-32 nt: 99.8%	Normalized hits on the main strand(s): 99.9%	Predicted directionality: mono:plus
Cluster 204	Location: CM035919.1	Coordinates: 86727026-86732038	Size [bp]: 5013	Hits (absolute): 1475	Hits (normalized): 105.78998993115	Hits (normalized) per kb: 21.1028929295727	Normalized hits with 1T: 85.8%	Normalized hits with 10A: 35.8%	Normalized hits 25-32 nt: 99.4%	Normalized hits on the main strand(s): 85.2%	Predicted directionality: mono:plus
Cluster 205	Location: CM035919.1	Coordinates: 86809968-86824979	Size [bp]: 15012	Hits (absolute): 1970	Hits (normalized): 527.953835422195	Hits (normalized) per kb: 35.1684596053245	Normalized hits with 1T: 93.3%	Normalized hits with 10A: 2.2%	Normalized hits 25-32 nt: 99.2%	Normalized hits on the main strand(s): 100%	Predicted directionality: mono:minus
Cluster 206	Location: CM035919.1	Coordinates: 86928129-86933768	Size [bp]: 5640	Hits (absolute): 626	Hits (normalized): 110.749382625561	Hits (normalized) per kb: 19.6365794445676	Normalized hits with 1T: 97.6%	Normalized hits with 10A: 4.5%	Normalized hits 25-32 nt: 97.2%	Normalized hits on the main strand(s): 100%	Predicted directionality: mono:plus
Cluster 207	Location: CM035919.1	Coordinates: 86940610-86942505	Size [bp]: 1896	Hits (absolute): 675	Hits (normalized): 206.711278841774	Hits (normalized) per kb: 109.025090308102	Normalized hits with 1T: 98.3%	Normalized hits with 10A: 3.8%	Normalized hits 25-32 nt: 99.9%	Normalized hits on the main strand(s): 99.9%	Predicted directionality: mono:plus
Cluster 208	Location: CM035919.1	Coordinates: 97595418-97601768	Size [bp]: 6351	Hits (absolute): 1256	Hits (normalized): 287.920753291161	Hits (normalized) per kb: 45.334806580007	Normalized hits with 1T: 98.6%	Normalized hits with 10A: 10.6%	Normalized hits 25-32 nt: 99.8%	Normalized hits on the main strand(s): 99.1%	Predicted directionality: mono:plus
Cluster 209	Location: CM035919.1	Coordinates: 101035530-101041972	Size [bp]: 6443	Hits (absolute): 1816	Hits (normalized): 282.580015558654	Hits (normalized) per kb: 43.8587083529666	Normalized hits with 1T: 93.3%	Normalized hits with 10A: 57.9%	Normalized hits 25-32 nt: 99.6%	Normalized hits on the main strand(s): 93.7%	Predicted directionality: mono:minus
Cluster 210	Location: CM035919.1	Coordinates: 124026651-124032770	Size [bp]: 6120	Hits (absolute): 1221	Hits (normalized): 240.405069545125	Hits (normalized) per kb: 39.2815458108797	Normalized hits with 1T: 83.7%	Normalized hits with 10A: 22.9%	Normalized hits 25-32 nt: 99.4%	Normalized hits on the main strand(s): 89.9%	Predicted directionality: bi:minus-plus (split between 124028414 and 124028415)
Cluster 211	Location: CM035919.1	Coordinates: 124136060-124151991	Size [bp]: 15932	Hits (absolute): 1471	Hits (normalized): 355.674346750574	Hits (normalized) per kb: 22.3245878636951	Normalized hits with 1T: 95.5%	Normalized hits with 10A: 14.9%	Normalized hits 25-32 nt: 99.6%	Normalized hits on the main strand(s): 100%	Predicted directionality: mono:plus
Cluster 212	Location: CM035919.1	Coordinates: 124159310-124168020	Size [bp]: 8711	Hits (absolute): 1607	Hits (normalized): 198.267134007625	Hits (normalized) per kb: 22.7607077944116	Normalized hits with 1T: 89.9%	Normalized hits with 10A: 28.6%	Normalized hits 25-32 nt: 99.8%	Normalized hits on the main strand(s): 98.1%	Predicted directionality: mono:plus
Cluster 213	Location: CM035919.1	Coordinates: 124192197-124200027	Size [bp]: 7831	Hits (absolute): 1599	Hits (normalized): 142.412783802642	Hits (normalized) per kb: 18.1856419827608	Normalized hits with 1T: 95.4%	Normalized hits with 10A: 26.5%	Normalized hits 25-32 nt: 99.7%	Normalized hits on the main strand(s): 99.7%	Predicted directionality: mono:plus
Cluster 214	Location: CM035919.1	Coordinates: 127316004-127322226	Size [bp]: 6223	Hits (absolute): 1716	Hits (normalized): 230.192070759643	Hits (normalized) per kb: 36.9905183543274	Normalized hits with 1T: 91.9%	Normalized hits with 10A: 70.2%	Normalized hits 25-32 nt: 99.5%	Normalized hits on the main strand(s): 92.2%	Predicted directionality: mono:plus
Cluster 215	Location: CM035920.1	Coordinates: 9310785-9315882	Size [bp]: 5098	Hits (absolute): 875	Hits (normalized): 110.474534320823	Hits (normalized) per kb: 21.6704079676204	Normalized hits with 1T: 94.2%	Normalized hits with 10A: 14%	Normalized hits 25-32 nt: 99.8%	Normalized hits on the main strand(s): 93.9%	Predicted directionality: mono:plus
Cluster 216	Location: CM035920.1	Coordinates: 9972420-9976932	Size [bp]: 4513	Hits (absolute): 370	Hits (normalized): 146.899178506841	Hits (normalized) per kb: 32.5503422007348	Normalized hits with 1T: 99.1%	Normalized hits with 10A: 4.6%	Normalized hits 25-32 nt: 99.8%	Normalized hits on the main strand(s): 99.9%	Predicted directionality: mono:minus
Cluster 217	Location: CM035920.1	Coordinates: 10305146-10336004	Size [bp]: 30859	Hits (absolute): 6787	Hits (normalized): 1082.61050137871	Hits (normalized) per kb: 35.0824936574429	Normalized hits with 1T: 94.8%	Normalized hits with 10A: 24.3%	Normalized hits 25-32 nt: 99.8%	Normalized hits on the main strand(s): 99.9%	Predicted directionality: mono:minus
Cluster 218	Location: CM035920.1	Coordinates: 10350001-10359587	Size [bp]: 9587	Hits (absolute): 2001	Hits (normalized): 221.257108437031	Hits (normalized) per kb: 23.078711910559	Normalized hits with 1T: 96.6%	Normalized hits with 10A: 15.1%	Normalized hits 25-32 nt: 99.6%	Normalized hits on the main strand(s): 100%	Predicted directionality: mono:minus
Cluster 219	Location: CM035920.1	Coordinates: 10361020-10377925	Size [bp]: 16906	Hits (absolute): 3924	Hits (normalized): 2975.41578054111	Hits (normalized) per kb: 175.997456078896	Normalized hits with 1T: 99.1%	Normalized hits with 10A: 5.8%	Normalized hits 25-32 nt: 99.9%	Normalized hits on the main strand(s): 100%	Predicted directionality: mono:minus
Cluster 220	Location: CM035920.1	Coordinates: 10393215-10413467	Size [bp]: 20253	Hits (absolute): 6593	Hits (normalized): 2491.49214496018	Hits (normalized) per kb: 123.01866923888	Normalized hits with 1T: 96.1%	Normalized hits with 10A: 62.9%	Normalized hits 25-32 nt: 99.9%	Normalized hits on the main strand(s): 99.9%	Predicted directionality: mono:minus
Cluster 221	Location: CM035920.1	Coordinates: 11187064-11223277	Size [bp]: 36214	Hits (absolute): 10982	Hits (normalized): 5704.70163604659	Hits (normalized) per kb: 157.527357666966	Normalized hits with 1T: 97.8%	Normalized hits with 10A: 31.9%	Normalized hits 25-32 nt: 99.9%	Normalized hits on the main strand(s): 100%	Predicted directionality: mono:plus
Cluster 222	Location: CM035920.1	Coordinates: 11227046-11280597	Size [bp]: 53552	Hits (absolute): 14371	Hits (normalized): 2971.23887463618	Hits (normalized) per kb: 55.4829818909096	Normalized hits with 1T: 95.5%	Normalized hits with 10A: 18%	Normalized hits 25-32 nt: 99.7%	Normalized hits on the main strand(s): 99.8%	Predicted directionality: mono:plus
Cluster 223	Location: CM035920.1	Coordinates: 16915756-16922158	Size [bp]: 6403	Hits (absolute): 1257	Hits (normalized): 169.114133706757	Hits (normalized) per kb: 26.4118143938713	Normalized hits with 1T: 79%	Normalized hits with 10A: 26.4%	Normalized hits 25-32 nt: 99.1%	Normalized hits on the main strand(s): 85.9%	Predicted directionality: bi:minus-plus (split between 16918506 and 16918507)
Cluster 224	Location: CM035920.1	Coordinates: 22247237-22253948	Size [bp]: 6712	Hits (absolute): 2036	Hits (normalized): 151.174562935121	Hits (normalized) per kb: 22.5230783449827	Normalized hits with 1T: 92%	Normalized hits with 10A: 24.7%	Normalized hits 25-32 nt: 99.9%	Normalized hits on the main strand(s): 96.1%	Predicted directionality: mono:plus
Cluster 225	Location: CM035920.1	Coordinates: 31252008-31258677	Size [bp]: 6670	Hits (absolute): 1201	Hits (normalized): 188.846668482394	Hits (normalized) per kb: 28.3128499893021	Normalized hits with 1T: 89.7%	Normalized hits with 10A: 26.4%	Normalized hits 25-32 nt: 98%	Normalized hits on the main strand(s): 92.5%	Predicted directionality: bi:minus-plus (split between 31256931 and 31256932)
Cluster 226	Location: CM035920.1	Coordinates: 33037310-33042844	Size [bp]: 5535	Hits (absolute): 847	Hits (normalized): 119.839461881636	Hits (normalized) per kb: 21.6515373936952	Normalized hits with 1T: 92.7%	Normalized hits with 10A: 9.6%	Normalized hits 25-32 nt: 99.8%	Normalized hits on the main strand(s): 98.6%	Predicted directionality: mono:plus
Cluster 227	Location: CM035920.1	Coordinates: 33114087-33136975	Size [bp]: 22889	Hits (absolute): 4934	Hits (normalized): 1288.92104259994	Hits (normalized) per kb: 56.3118893233291	Normalized hits with 1T: 97.3%	Normalized hits with 10A: 15.4%	Normalized hits 25-32 nt: 99.8%	Normalized hits on the main strand(s): 99.8%	Predicted directionality: mono:plus
Cluster 228	Location: CM035920.1	Coordinates: 33144042-33152809	Size [bp]: 8768	Hits (absolute): 1377	Hits (normalized): 155.850874476926	Hits (normalized) per kb: 17.774682817278	Normalized hits with 1T: 93.6%	Normalized hits with 10A: 17%	Normalized hits 25-32 nt: 99.7%	Normalized hits on the main strand(s): 99.9%	Predicted directionality: mono:plus
Cluster 229	Location: CM035920.1	Coordinates: 33167001-33186025	Size [bp]: 19025	Hits (absolute): 4102	Hits (normalized): 751.733067481585	Hits (normalized) per kb: 39.5128850690001	Normalized hits with 1T: 95.9%	Normalized hits with 10A: 16.5%	Normalized hits 25-32 nt: 99.8%	Normalized hits on the main strand(s): 100%	Predicted directionality: mono:plus
Cluster 230	Location: CM035920.1	Coordinates: 35217038-35228960	Size [bp]: 11923	Hits (absolute): 1835	Hits (normalized): 450.387998878999	Hits (normalized) per kb: 37.7746955374427	Normalized hits with 1T: 89.7%	Normalized hits with 10A: 24.2%	Normalized hits 25-32 nt: 99.7%	Normalized hits on the main strand(s): 99.6%	Predicted directionality: mono:minus
Cluster 231	Location: CM035920.1	Coordinates: 35238135-35301026	Size [bp]: 62892	Hits (absolute): 14681	Hits (normalized): 5070.63975724203	Hits (normalized) per kb: 80.6248765506274	Normalized hits with 1T: 97.6%	Normalized hits with 10A: 8%	Normalized hits 25-32 nt: 99.9%	Normalized hits on the main strand(s): 99.9%	Predicted directionality: mono:minus
Cluster 232	Location: CM035920.1	Coordinates: 35306183-35315989	Size [bp]: 9807	Hits (absolute): 1409	Hits (normalized): 290.422714206147	Hits (normalized) per kb: 29.6135217698524	Normalized hits with 1T: 95.9%	Normalized hits with 10A: 48.5%	Normalized hits 25-32 nt: 99.6%	Normalized hits on the main strand(s): 97.7%	Predicted directionality: mono:minus
Cluster 233	Location: CM035920.1	Coordinates: 35336212-35341995	Size [bp]: 5784	Hits (absolute): 1119	Hits (normalized): 120.855073589289	Hits (normalized) per kb: 20.8946177062497	Normalized hits with 1T: 97%	Normalized hits with 10A: 11.4%	Normalized hits 25-32 nt: 99.8%	Normalized hits on the main strand(s): 99.8%	Predicted directionality: mono:minus
Cluster 234	Location: CM035920.1	Coordinates: 37491025-37499662	Size [bp]: 8638	Hits (absolute): 1354	Hits (normalized): 152.773517274338	Hits (normalized) per kb: 17.6859212288149	Normalized hits with 1T: 84.1%	Normalized hits with 10A: 19.3%	Normalized hits 25-32 nt: 99.4%	Normalized hits on the main strand(s): 83.3%	Predicted directionality: mono:plus
Cluster 235	Location: CM035920.1	Coordinates: 43856584-43862931	Size [bp]: 6348	Hits (absolute): 917	Hits (normalized): 114.39216935308	Hits (normalized) per kb: 18.020000278306	Normalized hits with 1T: 94.3%	Normalized hits with 10A: 16%	Normalized hits 25-32 nt: 99.7%	Normalized hits on the main strand(s): 94.4%	Predicted directionality: mono:plus
Cluster 236	Location: CM035920.1	Coordinates: 44058337-44062810	Size [bp]: 4474	Hits (absolute): 833	Hits (normalized): 109.856362696901	Hits (normalized) per kb: 24.554111227454	Normalized hits with 1T: 94.4%	Normalized hits with 10A: 16.2%	Normalized hits 25-32 nt: 99.8%	Normalized hits on the main strand(s): 94.6%	Predicted directionality: mono:plus
Cluster 237	Location: CM035920.1	Coordinates: 44758546-44761458	Size [bp]: 2913	Hits (absolute): 325	Hits (normalized): 91.2233330832528	Hits (normalized) per kb: 31.3160668839956	Normalized hits with 1T: 96.9%	Normalized hits with 10A: 94.6%	Normalized hits 25-32 nt: 99.8%	Normalized hits on the main strand(s): 97.7%	Predicted directionality: mono:minus
Cluster 238	Location: CM035920.1	Coordinates: 45038696-45044298	Size [bp]: 5603	Hits (absolute): 718	Hits (normalized): 151.734436234149	Hits (normalized) per kb: 27.080671402999	Normalized hits with 1T: 96.3%	Normalized hits with 10A: 12.7%	Normalized hits 25-32 nt: 98%	Normalized hits on the main strand(s): 99.8%	Predicted directionality: mono:plus
Cluster 239	Location: CM035920.1	Coordinates: 45137008-45141761	Size [bp]: 4754	Hits (absolute): 1797	Hits (normalized): 91.7742827533453	Hits (normalized) per kb: 19.3045971255126	Normalized hits with 1T: 92.3%	Normalized hits with 10A: 38.3%	Normalized hits 25-32 nt: 98.3%	Normalized hits on the main strand(s): 99.5%	Predicted directionality: mono:minus
Cluster 240	Location: CM035920.1	Coordinates: 51998228-52006529	Size [bp]: 8302	Hits (absolute): 1124	Hits (normalized): 182.601043317449	Hits (normalized) per kb: 21.9947022750762	Normalized hits with 1T: 94%	Normalized hits with 10A: 6.8%	Normalized hits 25-32 nt: 99.8%	Normalized hits on the main strand(s): 99.9%	Predicted directionality: mono:plus
Cluster 241	Location: CM035920.1	Coordinates: 52591870-52597025	Size [bp]: 5156	Hits (absolute): 468	Hits (normalized): 92.1895872880231	Hits (normalized) per kb: 17.8802182492302	Normalized hits with 1T: 96.2%	Normalized hits with 10A: 18.3%	Normalized hits 25-32 nt: 99.8%	Normalized hits on the main strand(s): 99.6%	Predicted directionality: mono:plus
Cluster 242	Location: CM035920.1	Coordinates: 64657274-64663969	Size [bp]: 6696	Hits (absolute): 954	Hits (normalized): 108.925855773311	Hits (normalized) per kb: 16.2671336336956	Normalized hits with 1T: 93.8%	Normalized hits with 10A: 18.1%	Normalized hits 25-32 nt: 99.7%	Normalized hits on the main strand(s): 92.9%	Predicted directionality: mono:plus
Cluster 243	Location: CM035920.1	Coordinates: 64749054-64753727	Size [bp]: 4674	Hits (absolute): 715	Hits (normalized): 94.3782943549345	Hits (normalized) per kb: 20.1922130101439	Normalized hits with 1T: 96.9%	Normalized hits with 10A: 16.1%	Normalized hits 25-32 nt: 99.6%	Normalized hits on the main strand(s): 95.8%	Predicted directionality: mono:minus
Cluster 244	Location: CM035920.1	Coordinates: 66172092-66178854	Size [bp]: 6763	Hits (absolute): 1542	Hits (normalized): 521.085549790162	Hits (normalized) per kb: 77.0492522468686	Normalized hits with 1T: 5.4%	Normalized hits with 10A: 97.7%	Normalized hits 25-32 nt: 99.2%	Normalized hits on the main strand(s): 98.6%	Predicted directionality: mono:minus
Cluster 245	Location: CM035920.1	Coordinates: 91582039-91590521	Size [bp]: 8483	Hits (absolute): 1187	Hits (normalized): 175.235965477784	Hits (normalized) per kb: 20.6569882568209	Normalized hits with 1T: 94.3%	Normalized hits with 10A: 18.3%	Normalized hits 25-32 nt: 99.6%	Normalized hits on the main strand(s): 93.9%	Predicted directionality: mono:minus
Cluster 246	Location: CM035920.1	Coordinates: 96649041-96680000	Size [bp]: 30960	Hits (absolute): 9117	Hits (normalized): 1383.23422034662	Hits (normalized) per kb: 44.6778310433507	Normalized hits with 1T: 95.2%	Normalized hits with 10A: 14.7%	Normalized hits 25-32 nt: 99.7%	Normalized hits on the main strand(s): 100%	Predicted directionality: mono:minus
Cluster 247	Location: CM035920.1	Coordinates: 97536001-97544727	Size [bp]: 8727	Hits (absolute): 1570	Hits (normalized): 190.93026791892	Hits (normalized) per kb: 21.8779842807979	Normalized hits with 1T: 96.4%	Normalized hits with 10A: 8.2%	Normalized hits 25-32 nt: 99.8%	Normalized hits on the main strand(s): 100%	Predicted directionality: mono:plus
Cluster 248	Location: CM035920.1	Coordinates: 103188169-103192153	Size [bp]: 3985	Hits (absolute): 312	Hits (normalized): 145.591386844902	Hits (normalized) per kb: 36.5348289395403	Normalized hits with 1T: 7.9%	Normalized hits with 10A: 92.9%	Normalized hits 25-32 nt: 100%	Normalized hits on the main strand(s): 99.7%	Predicted directionality: mono:plus
Cluster 249	Location: CM035920.1	Coordinates: 105651071-105658888	Size [bp]: 7818	Hits (absolute): 329	Hits (normalized): 110.038511896967	Hits (normalized) per kb: 14.0753514177871	Normalized hits with 1T: 99%	Normalized hits with 10A: 8.3%	Normalized hits 25-32 nt: 99.9%	Normalized hits on the main strand(s): 95.9%	Predicted directionality: mono:plus
Cluster 250	Location: CM035920.1	Coordinates: 106482005-106490787	Size [bp]: 8783	Hits (absolute): 1338	Hits (normalized): 125.128845374956	Hits (normalized) per kb: 14.246584403405	Normalized hits with 1T: 90.5%	Normalized hits with 10A: 18.5%	Normalized hits 25-32 nt: 99.6%	Normalized hits on the main strand(s): 92.8%	Predicted directionality: mono:plus
Cluster 251	Location: CM035920.1	Coordinates: 107178077-107183684	Size [bp]: 5608	Hits (absolute): 506	Hits (normalized): 123.766004898853	Hits (normalized) per kb: 22.0694856606318	Normalized hits with 1T: 97.8%	Normalized hits with 10A: 3.9%	Normalized hits 25-32 nt: 97.6%	Normalized hits on the main strand(s): 99.9%	Predicted directionality: mono:plus
Cluster 252	Location: CM035920.1	Coordinates: 107188877-107190943	Size [bp]: 2067	Hits (absolute): 547	Hits (normalized): 377.202859232131	Hits (normalized) per kb: 182.488234599031	Normalized hits with 1T: 97.8%	Normalized hits with 10A: 62.6%	Normalized hits 25-32 nt: 100%	Normalized hits on the main strand(s): 100%	Predicted directionality: mono:plus
Cluster 253	Location: CM035920.1	Coordinates: 110772137-110778265	Size [bp]: 6129	Hits (absolute): 897	Hits (normalized): 103.152679984974	Hits (normalized) per kb: 16.830455210871	Normalized hits with 1T: 97.7%	Normalized hits with 10A: 13.2%	Normalized hits 25-32 nt: 99.7%	Normalized hits on the main strand(s): 98%	Predicted directionality: mono:minus
Cluster 254	Location: CM035920.1	Coordinates: 113420092-113426543	Size [bp]: 6452	Hits (absolute): 1230	Hits (normalized): 168.546447777636	Hits (normalized) per kb: 26.1231645038298	Normalized hits with 1T: 79%	Normalized hits with 10A: 26.1%	Normalized hits 25-32 nt: 99.1%	Normalized hits on the main strand(s): 85.9%	Predicted directionality: bi:minus-plus (split between 113422271 and 113422271)
Cluster 255	Location: CM035920.1	Coordinates: 115435920-115441452	Size [bp]: 5533	Hits (absolute): 698	Hits (normalized): 149.08985000974	Hits (normalized) per kb: 26.9457817449409	Normalized hits with 1T: 96.2%	Normalized hits with 10A: 13.1%	Normalized hits 25-32 nt: 98%	Normalized hits on the main strand(s): 99.7%	Predicted directionality: mono:plus
Cluster 256	Location: CM035920.1	Coordinates: 116768192-116771384	Size [bp]: 3193	Hits (absolute): 830	Hits (normalized): 101.883288804099	Hits (normalized) per kb: 31.9080437771316	Normalized hits with 1T: 94.1%	Normalized hits with 10A: 16.8%	Normalized hits 25-32 nt: 99.8%	Normalized hits on the main strand(s): 94.2%	Predicted directionality: mono:minus
Cluster 257	Location: CM035921.1	Coordinates: 3079546-3087462	Size [bp]: 7917	Hits (absolute): 610	Hits (normalized): 144.459268238214	Hits (normalized) per kb: 18.2464471654088	Normalized hits with 1T: 31.5%	Normalized hits with 10A: 98.9%	Normalized hits 25-32 nt: 97.9%	Normalized hits on the main strand(s): 99.9%	Predicted directionality: mono:minus
Cluster 258	Location: CM035921.1	Coordinates: 3278481-3283170	Size [bp]: 4690	Hits (absolute): 858	Hits (normalized): 224.461864292094	Hits (normalized) per kb: 47.8599689352612	Normalized hits with 1T: 32.6%	Normalized hits with 10A: 85.9%	Normalized hits 25-32 nt: 98.4%	Normalized hits on the main strand(s): 99.9%	Predicted directionality: mono:minus
Cluster 259	Location: CM035921.1	Coordinates: 5099028-5104130	Size [bp]: 5103	Hits (absolute): 492	Hits (normalized): 92.7298289786952	Hits (normalized) per kb: 18.1716637798533	Normalized hits with 1T: 96.4%	Normalized hits with 10A: 18.3%	Normalized hits 25-32 nt: 99.9%	Normalized hits on the main strand(s): 99.7%	Predicted directionality: mono:minus
Cluster 260	Location: CM035921.1	Coordinates: 7821024-7825866	Size [bp]: 4843	Hits (absolute): 511	Hits (normalized): 229.818479093052	Hits (normalized) per kb: 47.453902140796	Normalized hits with 1T: 13.5%	Normalized hits with 10A: 92%	Normalized hits 25-32 nt: 99.4%	Normalized hits on the main strand(s): 100%	Predicted directionality: mono:minus
Cluster 261	Location: CM035921.1	Coordinates: 8764217-8771686	Size [bp]: 7470	Hits (absolute): 1310	Hits (normalized): 124.706670365712	Hits (normalized) per kb: 16.6941677325221	Normalized hits with 1T: 90.7%	Normalized hits with 10A: 18.7%	Normalized hits 25-32 nt: 99.7%	Normalized hits on the main strand(s): 92.9%	Predicted directionality: mono:plus
Cluster 262	Location: CM035921.1	Coordinates: 9120487-9125259	Size [bp]: 4773	Hits (absolute): 698	Hits (normalized): 227.069965497392	Hits (normalized) per kb: 47.5741146858012	Normalized hits with 1T: 98.4%	Normalized hits with 10A: 4.4%	Normalized hits 25-32 nt: 99.5%	Normalized hits on the main strand(s): 100%	Predicted directionality: mono:minus
Cluster 263	Location: CM035921.1	Coordinates: 9145795-9149377	Size [bp]: 3583	Hits (absolute): 969	Hits (normalized): 147.738281733216	Hits (normalized) per kb: 41.2329029367778	Normalized hits with 1T: 97.1%	Normalized hits with 10A: 11.7%	Normalized hits 25-32 nt: 99.5%	Normalized hits on the main strand(s): 100%	Predicted directionality: mono:minus
Cluster 264	Location: CM035921.1	Coordinates: 9210145-9212645	Size [bp]: 2501	Hits (absolute): 475	Hits (normalized): 201.631416762396	Hits (normalized) per kb: 80.6199841796097	Normalized hits with 1T: 71.8%	Normalized hits with 10A: 87.6%	Normalized hits 25-32 nt: 99.5%	Normalized hits on the main strand(s): 99.9%	Predicted directionality: mono:minus
Cluster 265	Location: CM035921.1	Coordinates: 11696787-11699523	Size [bp]: 2737	Hits (absolute): 328	Hits (normalized): 152.002472967746	Hits (normalized) per kb: 55.5360990619585	Normalized hits with 1T: 3.4%	Normalized hits with 10A: 94.8%	Normalized hits 25-32 nt: 99.7%	Normalized hits on the main strand(s): 99.9%	Predicted directionality: mono:plus
Cluster 266	Location: CM035921.1	Coordinates: 15300081-15308954	Size [bp]: 8874	Hits (absolute): 2338	Hits (normalized): 168.031964669426	Hits (normalized) per kb: 18.9355725687525	Normalized hits with 1T: 77.6%	Normalized hits with 10A: 25.5%	Normalized hits 25-32 nt: 98.8%	Normalized hits on the main strand(s): 76.8%	Predicted directionality: mono:minus
Cluster 267	Location: CM035921.1	Coordinates: 26965202-26967215	Size [bp]: 2014	Hits (absolute): 402	Hits (normalized): 135.916410966162	Hits (normalized) per kb: 67.4860647276481	Normalized hits with 1T: 99.1%	Normalized hits with 10A: 75.7%	Normalized hits 25-32 nt: 99.8%	Normalized hits on the main strand(s): 99.4%	Predicted directionality: mono:plus
Cluster 268	Location: CM035921.1	Coordinates: 36947792-36950880	Size [bp]: 3089	Hits (absolute): 230	Hits (normalized): 107.849090546297	Hits (normalized) per kb: 34.9140563124065	Normalized hits with 1T: 99.1%	Normalized hits with 10A: 47.9%	Normalized hits 25-32 nt: 99.7%	Normalized hits on the main strand(s): 99.9%	Predicted directionality: mono:plus
Cluster 269	Location: CM035921.1	Coordinates: 62633004-62639434	Size [bp]: 6431	Hits (absolute): 1967	Hits (normalized): 102.291086100299	Hits (normalized) per kb: 15.9057970885346	Normalized hits with 1T: 86.3%	Normalized hits with 10A: 32.1%	Normalized hits 25-32 nt: 99.3%	Normalized hits on the main strand(s): 82.8%	Predicted directionality: mono:minus
Cluster 270	Location: CM035921.1	Coordinates: 63095146-63097634	Size [bp]: 2489	Hits (absolute): 603	Hits (normalized): 259.535176767092	Hits (normalized) per kb: 104.27320022967	Normalized hits with 1T: 98.9%	Normalized hits with 10A: 6.2%	Normalized hits 25-32 nt: 100%	Normalized hits on the main strand(s): 98.9%	Predicted directionality: mono:minus
Cluster 271	Location: CM035921.1	Coordinates: 63307624-63319863	Size [bp]: 12240	Hits (absolute): 1402	Hits (normalized): 198.768879420151	Hits (normalized) per kb: 16.2391772278804	Normalized hits with 1T: 67.7%	Normalized hits with 10A: 66.7%	Normalized hits 25-32 nt: 99.5%	Normalized hits on the main strand(s): 80.2%	Predicted directionality: mono:minus
Cluster 272	Location: CM035921.1	Coordinates: 67446925-67449724	Size [bp]: 2800	Hits (absolute): 726	Hits (normalized): 304.555611663914	Hits (normalized) per kb: 108.769988105039	Normalized hits with 1T: 97.8%	Normalized hits with 10A: 77%	Normalized hits 25-32 nt: 99.9%	Normalized hits on the main strand(s): 100%	Predicted directionality: mono:minus
Cluster 273	Location: CM035921.1	Coordinates: 83830003-83838538	Size [bp]: 8536	Hits (absolute): 1281	Hits (normalized): 100.663588908459	Hits (normalized) per kb: 11.7927108829794	Normalized hits with 1T: 86.2%	Normalized hits with 10A: 35.3%	Normalized hits 25-32 nt: 99.3%	Normalized hits on the main strand(s): 84.4%	Predicted directionality: mono:plus
Cluster 274	Location: CM035921.1	Coordinates: 85022026-85030779	Size [bp]: 8754	Hits (absolute): 638	Hits (normalized): 293.158699737058	Hits (normalized) per kb: 33.4882796158334	Normalized hits with 1T: 76.8%	Normalized hits with 10A: 26.3%	Normalized hits 25-32 nt: 99.2%	Normalized hits on the main strand(s): 99.4%	Predicted directionality: mono:minus
Cluster 275	Location: CM035921.1	Coordinates: 85294733-85300642	Size [bp]: 5910	Hits (absolute): 1519	Hits (normalized): 107.256560617617	Hits (normalized) per kb: 18.1485997450557	Normalized hits with 1T: 86.2%	Normalized hits with 10A: 35.4%	Normalized hits 25-32 nt: 99.3%	Normalized hits on the main strand(s): 85%	Predicted directionality: mono:minus
Cluster 276	Location: CM035921.1	Coordinates: 87537043-87541872	Size [bp]: 4830	Hits (absolute): 636	Hits (normalized): 94.9167365001432	Hits (normalized) per kb: 19.6512565576205	Normalized hits with 1T: 94.4%	Normalized hits with 10A: 28%	Normalized hits 25-32 nt: 96.8%	Normalized hits on the main strand(s): 99.8%	Predicted directionality: bi:plus-minus (split between 87540987 and 87540991)
Cluster 277	Location: CM035921.1	Coordinates: 90436009-90442422	Size [bp]: 6414	Hits (absolute): 2710	Hits (normalized): 133.784889113654	Hits (normalized) per kb: 20.85827437869	Normalized hits with 1T: 93.4%	Normalized hits with 10A: 22.1%	Normalized hits 25-32 nt: 99.7%	Normalized hits on the main strand(s): 99.8%	Predicted directionality: mono:minus
Cluster 278	Location: CM035921.1	Coordinates: 92022001-92073267	Size [bp]: 51267	Hits (absolute): 22796	Hits (normalized): 7686.30474060932	Hits (normalized) per kb: 149.92670983597	Normalized hits with 1T: 95.3%	Normalized hits with 10A: 24.8%	Normalized hits 25-32 nt: 99.7%	Normalized hits on the main strand(s): 100%	Predicted directionality: mono:plus
Cluster 279	Location: CM035921.1	Coordinates: 92496405-92500528	Size [bp]: 4124	Hits (absolute): 636	Hits (normalized): 272.959002045506	Hits (normalized) per kb: 66.1881885876793	Normalized hits with 1T: 99.1%	Normalized hits with 10A: 6.5%	Normalized hits 25-32 nt: 99.9%	Normalized hits on the main strand(s): 99.9%	Predicted directionality: mono:plus
Cluster 280	Location: CM035921.1	Coordinates: 96071480-96077698	Size [bp]: 6219	Hits (absolute): 747	Hits (normalized): 219.045390087014	Hits (normalized) per kb: 35.2222756865186	Normalized hits with 1T: 15.2%	Normalized hits with 10A: 87.3%	Normalized hits 25-32 nt: 99.2%	Normalized hits on the main strand(s): 93.5%	Predicted directionality: bi:plus-minus (split between 96076502 and 96076503)
Cluster 281	Location: CM035921.1	Coordinates: 103647872-103653415	Size [bp]: 5544	Hits (absolute): 1094	Hits (normalized): 162.434815062918	Hits (normalized) per kb: 29.2990122044319	Normalized hits with 1T: 80.8%	Normalized hits with 10A: 24.5%	Normalized hits 25-32 nt: 99%	Normalized hits on the main strand(s): 88.1%	Predicted directionality: bi:minus-plus (split between 103649971 and 103649971)
Cluster 282	Location: CM035921.1	Coordinates: 108210205-108217702	Size [bp]: 7498	Hits (absolute): 723	Hits (normalized): 118.414511934633	Hits (normalized) per kb: 15.7925736449832	Normalized hits with 1T: 96.9%	Normalized hits with 10A: 16.4%	Normalized hits 25-32 nt: 99.9%	Normalized hits on the main strand(s): 97%	Predicted directionality: mono:plus
Cluster 283	Location: CM035922.1	Coordinates: 1387107-1389891	Size [bp]: 2785	Hits (absolute): 811	Hits (normalized): 507.321401776961	Hits (normalized) per kb: 182.161843561139	Normalized hits with 1T: 1.8%	Normalized hits with 10A: 95%	Normalized hits 25-32 nt: 99.6%	Normalized hits on the main strand(s): 100%	Predicted directionality: mono:minus
Cluster 284	Location: CM035922.1	Coordinates: 4714001-4722899	Size [bp]: 8899	Hits (absolute): 2007	Hits (normalized): 259.471215763896	Hits (normalized) per kb: 29.1571334449199	Normalized hits with 1T: 84.8%	Normalized hits with 10A: 26.4%	Normalized hits 25-32 nt: 99.6%	Normalized hits on the main strand(s): 99.2%	Predicted directionality: mono:plus
Cluster 285	Location: CM035922.1	Coordinates: 6165084-6172733	Size [bp]: 7650	Hits (absolute): 1646	Hits (normalized): 571.739077525676	Hits (normalized) per kb: 74.7372574859549	Normalized hits with 1T: 13.4%	Normalized hits with 10A: 88.7%	Normalized hits 25-32 nt: 99.5%	Normalized hits on the main strand(s): 86.6%	Predicted directionality: mono:plus
Cluster 286	Location: CM035922.1	Coordinates: 6585359-6590822	Size [bp]: 5464	Hits (absolute): 1651	Hits (normalized): 571.409385800474	Hits (normalized) per kb: 104.57722614291	Normalized hits with 1T: 13.3%	Normalized hits with 10A: 88.8%	Normalized hits 25-32 nt: 99.5%	Normalized hits on the main strand(s): 86.6%	Predicted directionality: mono:plus
Cluster 287	Location: CM035922.1	Coordinates: 7454565-7458189	Size [bp]: 3625	Hits (absolute): 169	Hits (normalized): 90.2010254909092	Hits (normalized) per kb: 24.8832979059275	Normalized hits with 1T: 99.3%	Normalized hits with 10A: 0.8%	Normalized hits 25-32 nt: 99.5%	Normalized hits on the main strand(s): 99.7%	Predicted directionality: mono:minus
Cluster 288	Location: CM035922.1	Coordinates: 11073133-11081861	Size [bp]: 8729	Hits (absolute): 1559	Hits (normalized): 537.225458022172	Hits (normalized) per kb: 61.5446295817815	Normalized hits with 1T: 5.2%	Normalized hits with 10A: 97.8%	Normalized hits 25-32 nt: 99.3%	Normalized hits on the main strand(s): 98.6%	Predicted directionality: mono:plus
Cluster 289	Location: CM035922.1	Coordinates: 19359069-19371536	Size [bp]: 12468	Hits (absolute): 1874	Hits (normalized): 249.315960749322	Hits (normalized) per kb: 19.9965181694377	Normalized hits with 1T: 96.7%	Normalized hits with 10A: 23.4%	Normalized hits 25-32 nt: 99.7%	Normalized hits on the main strand(s): 98.2%	Predicted directionality: mono:minus
Cluster 290	Location: CM035922.1	Coordinates: 19381153-19413947	Size [bp]: 32795	Hits (absolute): 8037	Hits (normalized): 1408.83985029936	Hits (normalized) per kb: 42.9592109958639	Normalized hits with 1T: 84%	Normalized hits with 10A: 15.6%	Normalized hits 25-32 nt: 99.5%	Normalized hits on the main strand(s): 98%	Predicted directionality: mono:minus
Cluster 291	Location: CM035922.1	Coordinates: 19418137-19431087	Size [bp]: 12951	Hits (absolute): 2121	Hits (normalized): 412.300023644953	Hits (normalized) per kb: 31.8353571220121	Normalized hits with 1T: 92.7%	Normalized hits with 10A: 16.2%	Normalized hits 25-32 nt: 99.8%	Normalized hits on the main strand(s): 100%	Predicted directionality: mono:minus
Cluster 292	Location: CM035922.1	Coordinates: 21435080-21443955	Size [bp]: 8876	Hits (absolute): 1412	Hits (normalized): 126.730348050939	Hits (normalized) per kb: 14.278035359947	Normalized hits with 1T: 90.5%	Normalized hits with 10A: 18.5%	Normalized hits 25-32 nt: 99.6%	Normalized hits on the main strand(s): 92.9%	Predicted directionality: mono:plus
Cluster 293	Location: CM035922.1	Coordinates: 21520103-21531962	Size [bp]: 11860	Hits (absolute): 2014	Hits (normalized): 194.575634993197	Hits (normalized) per kb: 16.406216752626	Normalized hits with 1T: 95.7%	Normalized hits with 10A: 16.9%	Normalized hits 25-32 nt: 99.4%	Normalized hits on the main strand(s): 99.9%	Predicted directionality: mono:minus
Cluster 294	Location: CM035922.1	Coordinates: 21551019-21559498	Size [bp]: 8480	Hits (absolute): 1622	Hits (normalized): 194.78174844377	Hits (normalized) per kb: 22.9696819278799	Normalized hits with 1T: 91.5%	Normalized hits with 10A: 24.8%	Normalized hits 25-32 nt: 99.6%	Normalized hits on the main strand(s): 91.7%	Predicted directionality: mono:minus
Cluster 295	Location: CM035922.1	Coordinates: 29444433-29452993	Size [bp]: 8561	Hits (absolute): 1390	Hits (normalized): 152.904013325652	Hits (normalized) per kb: 17.8606487651596	Normalized hits with 1T: 83.8%	Normalized hits with 10A: 19.4%	Normalized hits 25-32 nt: 99.4%	Normalized hits on the main strand(s): 83.1%	Predicted directionality: mono:plus
Cluster 296	Location: CM035922.1	Coordinates: 41815530-41820906	Size [bp]: 5377	Hits (absolute): 1333	Hits (normalized): 972.426999721203	Hits (normalized) per kb: 180.849290308117	Normalized hits with 1T: 92%	Normalized hits with 10A: 51.7%	Normalized hits 25-32 nt: 99.8%	Normalized hits on the main strand(s): 91.9%	Predicted directionality: mono:minus
Cluster 297	Location: CM035922.1	Coordinates: 43554710-43559997	Size [bp]: 5288	Hits (absolute): 686	Hits (normalized): 119.64350433644	Hits (normalized) per kb: 22.6258181363534	Normalized hits with 1T: 95%	Normalized hits with 10A: 18.9%	Normalized hits 25-32 nt: 97.4%	Normalized hits on the main strand(s): 98.8%	Predicted directionality: bi:minus-plus (split between 43555256 and 43555259)
Cluster 298	Location: CM035922.1	Coordinates: 53342836-53346697	Size [bp]: 3862	Hits (absolute): 537	Hits (normalized): 193.959064377238	Hits (normalized) per kb: 50.2222852266421	Normalized hits with 1T: 98.2%	Normalized hits with 10A: 67.8%	Normalized hits 25-32 nt: 99.8%	Normalized hits on the main strand(s): 98.9%	Predicted directionality: mono:minus
Cluster 299	Location: CM035922.1	Coordinates: 58579127-58584397	Size [bp]: 5271	Hits (absolute): 1946	Hits (normalized): 286.680087832843	Hits (normalized) per kb: 54.3884886032462	Normalized hits with 1T: 93.3%	Normalized hits with 10A: 57.7%	Normalized hits 25-32 nt: 99.6%	Normalized hits on the main strand(s): 93.5%	Predicted directionality: mono:plus
Cluster 300	Location: CM035922.1	Coordinates: 58742011-58749676	Size [bp]: 7666	Hits (absolute): 820	Hits (normalized): 126.355964313773	Hits (normalized) per kb: 16.4823979584723	Normalized hits with 1T: 92.7%	Normalized hits with 10A: 54.1%	Normalized hits 25-32 nt: 99.8%	Normalized hits on the main strand(s): 99.7%	Predicted directionality: mono:plus
Cluster 301	Location: CM035922.1	Coordinates: 58862249-58869831	Size [bp]: 7583	Hits (absolute): 790	Hits (normalized): 124.865214477759	Hits (normalized) per kb: 16.4663230251286	Normalized hits with 1T: 92.6%	Normalized hits with 10A: 54.2%	Normalized hits 25-32 nt: 99.9%	Normalized hits on the main strand(s): 99.8%	Predicted directionality: mono:plus
Cluster 302	Location: CM035922.1	Coordinates: 68766063-68770322	Size [bp]: 4260	Hits (absolute): 1416	Hits (normalized): 98.5392326287881	Hits (normalized) per kb: 23.1311301714624	Normalized hits with 1T: 86.1%	Normalized hits with 10A: 37.4%	Normalized hits 25-32 nt: 99.4%	Normalized hits on the main strand(s): 83.7%	Predicted directionality: mono:plus
Cluster 303	Location: CM035922.1	Coordinates: 72209083-72216585	Size [bp]: 7503	Hits (absolute): 1755	Hits (normalized): 231.540895370106	Hits (normalized) per kb: 30.8596785590631	Normalized hits with 1T: 91.9%	Normalized hits with 10A: 70.2%	Normalized hits 25-32 nt: 99.5%	Normalized hits on the main strand(s): 92.2%	Predicted directionality: mono:plus
Cluster 304	Location: CM035922.1	Coordinates: 73234305-73241606	Size [bp]: 7302	Hits (absolute): 2011	Hits (normalized): 155.534613999279	Hits (normalized) per kb: 21.2999855905695	Normalized hits with 1T: 92.7%	Normalized hits with 10A: 26.3%	Normalized hits 25-32 nt: 99.8%	Normalized hits on the main strand(s): 96.9%	Predicted directionality: mono:plus
Cluster 305	Location: CM035922.1	Coordinates: 74608024-74616887	Size [bp]: 8864	Hits (absolute): 2014	Hits (normalized): 102.593648134641	Hits (normalized) per kb: 11.5739520074758	Normalized hits with 1T: 86%	Normalized hits with 10A: 32.1%	Normalized hits 25-32 nt: 99.4%	Normalized hits on the main strand(s): 82.8%	Predicted directionality: mono:minus
Cluster 306	Location: CM035922.1	Coordinates: 79405141-79410920	Size [bp]: 5780	Hits (absolute): 1556	Hits (normalized): 106.811357430456	Hits (normalized) per kb: 18.47918424382	Normalized hits with 1T: 85.6%	Normalized hits with 10A: 36.1%	Normalized hits 25-32 nt: 99.3%	Normalized hits on the main strand(s): 84.9%	Predicted directionality: mono:minus
Cluster 307	Location: CM035922.1	Coordinates: 82585611-82657001	Size [bp]: 71391	Hits (absolute): 13334	Hits (normalized): 4590.65941757325	Hits (normalized) per kb: 64.3032279255923	Normalized hits with 1T: 93.9%	Normalized hits with 10A: 11.5%	Normalized hits 25-32 nt: 99.7%	Normalized hits on the main strand(s): 99.9%	Predicted directionality: mono:minus
Cluster 308	Location: CM035922.1	Coordinates: 82659700-82678951	Size [bp]: 19252	Hits (absolute): 3306	Hits (normalized): 2182.61761256787	Hits (normalized) per kb: 113.370913592068	Normalized hits with 1T: 95.7%	Normalized hits with 10A: 30%	Normalized hits 25-32 nt: 99.8%	Normalized hits on the main strand(s): 99.7%	Predicted directionality: mono:minus
Cluster 309	Location: CM035922.1	Coordinates: 82687058-82694871	Size [bp]: 7814	Hits (absolute): 891	Hits (normalized): 158.485508722291	Hits (normalized) per kb: 20.2823724188978	Normalized hits with 1T: 95.8%	Normalized hits with 10A: 30.5%	Normalized hits 25-32 nt: 99.9%	Normalized hits on the main strand(s): 99.9%	Predicted directionality: mono:minus
Cluster 310	Location: CM035922.1	Coordinates: 90743135-90744527	Size [bp]: 1393	Hits (absolute): 446	Hits (normalized): 226.238284437462	Hits (normalized) per kb: 162.410642852729	Normalized hits with 1T: 98.6%	Normalized hits with 10A: 4.9%	Normalized hits 25-32 nt: 99.9%	Normalized hits on the main strand(s): 99.5%	Predicted directionality: mono:minus
Cluster 311	Location: CM035922.1	Coordinates: 99273777-99278359	Size [bp]: 4583	Hits (absolute): 624	Hits (normalized): 117.30874807867	Hits (normalized) per kb: 25.5961862542141	Normalized hits with 1T: 96.9%	Normalized hits with 10A: 1.3%	Normalized hits 25-32 nt: 98.9%	Normalized hits on the main strand(s): 100%	Predicted directionality: mono:plus
Cluster 312	Location: CM035922.1	Coordinates: 100229003-100238004	Size [bp]: 9002	Hits (absolute): 865	Hits (normalized): 199.202853638075	Hits (normalized) per kb: 22.128893022989	Normalized hits with 1T: 98.2%	Normalized hits with 10A: 4.2%	Normalized hits 25-32 nt: 99.8%	Normalized hits on the main strand(s): 99%	Predicted directionality: mono:minus
Cluster 313	Location: CM035922.1	Coordinates: 100830413-100839022	Size [bp]: 8610	Hits (absolute): 946	Hits (normalized): 206.097434012199	Hits (normalized) per kb: 23.9369735690844	Normalized hits with 1T: 98%	Normalized hits with 10A: 4.7%	Normalized hits 25-32 nt: 99.9%	Normalized hits on the main strand(s): 98.8%	Predicted directionality: mono:plus
Cluster 314	Location: CM035922.1	Coordinates: 100841320-100849288	Size [bp]: 7969	Hits (absolute): 1168	Hits (normalized): 163.737306498395	Hits (normalized) per kb: 20.546560453851	Normalized hits with 1T: 81.4%	Normalized hits with 10A: 24.2%	Normalized hits 25-32 nt: 99.1%	Normalized hits on the main strand(s): 88.5%	Predicted directionality: bi:minus-plus (split between 100845655 and 100845656)
Cluster 315	Location: CM035922.1	Coordinates: 103543213-103574940	Size [bp]: 31728	Hits (absolute): 5579	Hits (normalized): 1977.86338238647	Hits (normalized) per kb: 62.3378925967866	Normalized hits with 1T: 98.7%	Normalized hits with 10A: 6.9%	Normalized hits 25-32 nt: 99.9%	Normalized hits on the main strand(s): 99.8%	Predicted directionality: mono:plus
Cluster 316	Location: CM035922.1	Coordinates: 105005007-105042699	Size [bp]: 37693	Hits (absolute): 7350	Hits (normalized): 2221.01649812306	Hits (normalized) per kb: 58.9237165366103	Normalized hits with 1T: 97.9%	Normalized hits with 10A: 7.9%	Normalized hits 25-32 nt: 99.9%	Normalized hits on the main strand(s): 99.5%	Predicted directionality: mono:plus
Cluster 317	Location: CM035922.1	Coordinates: 105093007-105105885	Size [bp]: 12879	Hits (absolute): 2137	Hits (normalized): 211.044892871888	Hits (normalized) per kb: 16.3866472685554	Normalized hits with 1T: 95%	Normalized hits with 10A: 18%	Normalized hits 25-32 nt: 99.1%	Normalized hits on the main strand(s): 99.9%	Predicted directionality: mono:plus
Cluster 318	Location: CM035922.1	Coordinates: 105404015-105408246	Size [bp]: 4232	Hits (absolute): 376	Hits (normalized): 90.5234457836936	Hits (normalized) per kb: 21.3901449993234	Normalized hits with 1T: 96.4%	Normalized hits with 10A: 18.5%	Normalized hits 25-32 nt: 99.9%	Normalized hits on the main strand(s): 99.5%	Predicted directionality: mono:minus
Cluster 319	Location: CM035923.1	Coordinates: 1229692-1240596	Size [bp]: 10905	Hits (absolute): 2128	Hits (normalized): 518.42076603699	Hits (normalized) per kb: 47.5398680886776	Normalized hits with 1T: 95.4%	Normalized hits with 10A: 10.4%	Normalized hits 25-32 nt: 99.8%	Normalized hits on the main strand(s): 99.5%	Predicted directionality: mono:plus
Cluster 320	Location: CM035923.1	Coordinates: 1250352-1260008	Size [bp]: 9657	Hits (absolute): 1476	Hits (normalized): 493.138853217261	Hits (normalized) per kb: 51.0651708619692	Normalized hits with 1T: 98.8%	Normalized hits with 10A: 32.6%	Normalized hits 25-32 nt: 99.7%	Normalized hits on the main strand(s): 99.4%	Predicted directionality: mono:plus
Cluster 321	Location: CM035923.1	Coordinates: 1282216-1291974	Size [bp]: 9759	Hits (absolute): 2241	Hits (normalized): 272.182220151603	Hits (normalized) per kb: 27.8907082614932	Normalized hits with 1T: 89.2%	Normalized hits with 10A: 47.4%	Normalized hits 25-32 nt: 99.5%	Normalized hits on the main strand(s): 91.5%	Predicted directionality: bi:minus-plus (split between 1284319 and 1284329)
Cluster 322	Location: CM035923.1	Coordinates: 1301308-1313902	Size [bp]: 12595	Hits (absolute): 2111	Hits (normalized): 388.675973326852	Hits (normalized) per kb: 30.8596785590631	Normalized hits with 1T: 89%	Normalized hits with 10A: 15.1%	Normalized hits 25-32 nt: 99.8%	Normalized hits on the main strand(s): 99.8%	Predicted directionality: mono:plus
Cluster 323	Location: CM035923.1	Coordinates: 1317234-1346019	Size [bp]: 28786	Hits (absolute): 6173	Hits (normalized): 804.82980110048	Hits (normalized) per kb: 27.9592014557404	Normalized hits with 1T: 96.4%	Normalized hits with 10A: 18.6%	Normalized hits 25-32 nt: 99.8%	Normalized hits on the main strand(s): 97.6%	Predicted directionality: mono:plus
Cluster 324	Location: CM035923.1	Coordinates: 1347073-1353951	Size [bp]: 6879	Hits (absolute): 1050	Hits (normalized): 112.134833665715	Hits (normalized) per kb: 16.3013802308191	Normalized hits with 1T: 95.3%	Normalized hits with 10A: 18.3%	Normalized hits 25-32 nt: 99.8%	Normalized hits on the main strand(s): 99.9%	Predicted directionality: mono:plus
Cluster 325	Location: CM035923.1	Coordinates: 9558905-9561564	Size [bp]: 2660	Hits (absolute): 597	Hits (normalized): 257.925392394679	Hits (normalized) per kb: 96.9646968394424	Normalized hits with 1T: 99%	Normalized hits with 10A: 6.1%	Normalized hits 25-32 nt: 100%	Normalized hits on the main strand(s): 99.5%	Predicted directionality: mono:minus
Cluster 326	Location: CM035923.1	Coordinates: 11865681-11881433	Size [bp]: 15753	Hits (absolute): 5111	Hits (normalized): 703.976880178116	Hits (normalized) per kb: 44.6883146955314	Normalized hits with 1T: 92.1%	Normalized hits with 10A: 69.8%	Normalized hits 25-32 nt: 99.6%	Normalized hits on the main strand(s): 92.5%	Predicted directionality: mono:minus
Cluster 327	Location: CM035923.1	Coordinates: 23409061-23410236	Size [bp]: 1176	Hits (absolute): 469	Hits (normalized): 225.085553777299	Hits (normalized) per kb: 191.399338952613	Normalized hits with 1T: 99%	Normalized hits with 10A: 4.5%	Normalized hits 25-32 nt: 99.8%	Normalized hits on the main strand(s): 99.9%	Predicted directionality: mono:plus
Cluster 328	Location: CM035923.1	Coordinates: 35433003-35438396	Size [bp]: 5394	Hits (absolute): 1468	Hits (normalized): 105.898891482082	Hits (normalized) per kb: 19.6323859836953	Normalized hits with 1T: 86.5%	Normalized hits with 10A: 35.6%	Normalized hits 25-32 nt: 99.3%	Normalized hits on the main strand(s): 84.9%	Predicted directionality: mono:plus
Cluster 329	Location: CM035923.1	Coordinates: 36527508-36533776	Size [bp]: 6269	Hits (absolute): 520	Hits (normalized): 139.813067604531	Hits (normalized) per kb: 22.302222739043	Normalized hits with 1T: 89.2%	Normalized hits with 10A: 8.4%	Normalized hits 25-32 nt: 92.7%	Normalized hits on the main strand(s): 100%	Predicted directionality: mono:minus
Cluster 330	Location: CM035923.1	Coordinates: 41658100-41666954	Size [bp]: 8855	Hits (absolute): 1475	Hits (normalized): 154.022524428761	Hits (normalized) per kb: 17.3937767880465	Normalized hits with 1T: 83.4%	Normalized hits with 10A: 19.8%	Normalized hits 25-32 nt: 99.4%	Normalized hits on the main strand(s): 82.7%	Predicted directionality: mono:plus
Cluster 331	Location: CM035923.1	Coordinates: 43751270-43755963	Size [bp]: 4694	Hits (absolute): 736	Hits (normalized): 146.896087696204	Hits (normalized) per kb: 31.2944006694888	Normalized hits with 1T: 91.3%	Normalized hits with 10A: 28.7%	Normalized hits 25-32 nt: 98.4%	Normalized hits on the main strand(s): 93.3%	Predicted directionality: bi:minus-plus (split between 43751899 and 43751900)
Cluster 332	Location: CM035923.1	Coordinates: 53165691-53171683	Size [bp]: 5993	Hits (absolute): 618	Hits (normalized): 94.375118265789	Hits (normalized) per kb: 15.747843395679	Normalized hits with 1T: 19.8%	Normalized hits with 10A: 93%	Normalized hits 25-32 nt: 99%	Normalized hits on the main strand(s): 93.8%	Predicted directionality: mono:plus
Cluster 333	Location: CM035923.1	Coordinates: 60313017-60322027	Size [bp]: 9011	Hits (absolute): 1615	Hits (normalized): 188.58908388984	Hits (normalized) per kb: 20.9288643033733	Normalized hits with 1T: 96.3%	Normalized hits with 10A: 28.2%	Normalized hits 25-32 nt: 99.7%	Normalized hits on the main strand(s): 99.1%	Predicted directionality: mono:minus
Cluster 334	Location: CM035923.1	Coordinates: 61579200-61600579	Size [bp]: 21380	Hits (absolute): 4143	Hits (normalized): 1286.84777819545	Hits (normalized) per kb: 60.1894428098916	Normalized hits with 1T: 88.3%	Normalized hits with 10A: 11.7%	Normalized hits 25-32 nt: 99.6%	Normalized hits on the main strand(s): 99.4%	Predicted directionality: mono:plus
Cluster 335	Location: CM035923.1	Coordinates: 61610040-61617771	Size [bp]: 7732	Hits (absolute): 918	Hits (normalized): 164.752197690767	Hits (normalized) per kb: 21.3076736021687	Normalized hits with 1T: 97.7%	Normalized hits with 10A: 41.3%	Normalized hits 25-32 nt: 99.8%	Normalized hits on the main strand(s): 100%	Predicted directionality: mono:plus
Cluster 336	Location: CM035923.1	Coordinates: 67180547-67186644	Size [bp]: 6098	Hits (absolute): 641	Hits (normalized): 115.093064938311	Hits (normalized) per kb: 18.8740684759591	Normalized hits with 1T: 96.8%	Normalized hits with 10A: 16.7%	Normalized hits 25-32 nt: 99.9%	Normalized hits on the main strand(s): 99.6%	Predicted directionality: mono:plus
Cluster 337	Location: CM035923.1	Coordinates: 67418137-67426845	Size [bp]: 8709	Hits (absolute): 3620	Hits (normalized): 2608.87898125049	Hits (normalized) per kb: 299.561275231172	Normalized hits with 1T: 94.9%	Normalized hits with 10A: 8.6%	Normalized hits 25-32 nt: 99.3%	Normalized hits on the main strand(s): 100%	Predicted directionality: mono:minus
Cluster 338	Location: CM035923.1	Coordinates: 68378327-68384976	Size [bp]: 6650	Hits (absolute): 579	Hits (normalized): 114.355965575706	Hits (normalized) per kb: 17.1966841270496	Normalized hits with 1T: 96.8%	Normalized hits with 10A: 16.5%	Normalized hits 25-32 nt: 99.9%	Normalized hits on the main strand(s): 99.5%	Predicted directionality: mono:minus
Cluster 339	Location: CM035923.1	Coordinates: 74786117-74790841	Size [bp]: 4725	Hits (absolute): 1540	Hits (normalized): 108.008430948675	Hits (normalized) per kb: 22.85925412491	Normalized hits with 1T: 85.4%	Normalized hits with 10A: 35.8%	Normalized hits 25-32 nt: 99.2%	Normalized hits on the main strand(s): 85.1%	Predicted directionality: mono:plus
Cluster 340	Location: CM035923.1	Coordinates: 90118239-90123926	Size [bp]: 5688	Hits (absolute): 617	Hits (normalized): 94.5882688010533	Hits (normalized) per kb: 16.6291690890019	Normalized hits with 1T: 97.1%	Normalized hits with 10A: 6.3%	Normalized hits 25-32 nt: 96.7%	Normalized hits on the main strand(s): 99.9%	Predicted directionality: mono:plus
Cluster 341	Location: CM035923.1	Coordinates: 90131402-90135588	Size [bp]: 4187	Hits (absolute): 625	Hits (normalized): 364.302252755905	Hits (normalized) per kb: 87.0080229083735	Normalized hits with 1T: 92.4%	Normalized hits with 10A: 66.3%	Normalized hits 25-32 nt: 100%	Normalized hits on the main strand(s): 100%	Predicted directionality: mono:plus
Cluster 342	Location: CM035923.1	Coordinates: 94220498-94224738	Size [bp]: 4241	Hits (absolute): 315	Hits (normalized): 310.89759678069	Hits (normalized) per kb: 73.3072873285095	Normalized hits with 1T: 5.3%	Normalized hits with 10A: 96.4%	Normalized hits 25-32 nt: 99.7%	Normalized hits on the main strand(s): 99.9%	Predicted directionality: mono:minus
Cluster 343	Location: CM035923.1	Coordinates: 94704412-94712008	Size [bp]: 7597	Hits (absolute): 1823	Hits (normalized): 140.147989461814	Hits (normalized) per kb: 18.4477332872779	Normalized hits with 1T: 96.5%	Normalized hits with 10A: 21.7%	Normalized hits 25-32 nt: 99%	Normalized hits on the main strand(s): 99.8%	Predicted directionality: mono:plus
Cluster 344	Location: CM035923.1	Coordinates: 100763893-100768183	Size [bp]: 4291	Hits (absolute): 1208	Hits (normalized): 128.232893925385	Hits (normalized) per kb: 29.8839999961141	Normalized hits with 1T: 85.1%	Normalized hits with 10A: 21%	Normalized hits 25-32 nt: 99.7%	Normalized hits on the main strand(s): 97.1%	Predicted directionality: mono:plus
Cluster 345	Location: CM035923.1	Coordinates: 106107181-106116027	Size [bp]: 8847	Hits (absolute): 1300	Hits (normalized): 266.791445210766	Hits (normalized) per kb: 30.1558760426665	Normalized hits with 1T: 95.7%	Normalized hits with 10A: 57.4%	Normalized hits 25-32 nt: 99.7%	Normalized hits on the main strand(s): 96%	Predicted directionality: mono:plus
Cluster 346	Location: CM035924.1	Coordinates: 182340-219022	Size [bp]: 36683	Hits (absolute): 6592	Hits (normalized): 1053.0134427139	Hits (normalized) per kb: 28.7056374910051	Normalized hits with 1T: 88.4%	Normalized hits with 10A: 16.7%	Normalized hits 25-32 nt: 99.8%	Normalized hits on the main strand(s): 99.8%	Predicted directionality: mono:plus
Cluster 347	Location: CM035924.1	Coordinates: 492780-497943	Size [bp]: 5164	Hits (absolute): 1044	Hits (normalized): 96.0267778745112	Hits (normalized) per kb: 18.5952033279529	Normalized hits with 1T: 86%	Normalized hits with 10A: 22.5%	Normalized hits 25-32 nt: 99.4%	Normalized hits on the main strand(s): 86.5%	Predicted directionality: mono:minus
Cluster 348	Location: CM035924.1	Coordinates: 1681073-1687568	Size [bp]: 6496	Hits (absolute): 1452	Hits (normalized): 101.488270308123	Hits (normalized) per kb: 15.6234373898015	Normalized hits with 1T: 86.7%	Normalized hits with 10A: 34.3%	Normalized hits 25-32 nt: 99.3%	Normalized hits on the main strand(s): 84.8%	Predicted directionality: mono:minus
Cluster 349	Location: CM035924.1	Coordinates: 3316225-3343987	Size [bp]: 27763	Hits (absolute): 14446	Hits (normalized): 10181.2375206662	Hits (normalized) per kb: 366.719551100637	Normalized hits with 1T: 96.3%	Normalized hits with 10A: 56.8%	Normalized hits 25-32 nt: 99.8%	Normalized hits on the main strand(s): 100%	Predicted directionality: mono:plus
Cluster 350	Location: CM035924.1	Coordinates: 14886006-14891282	Size [bp]: 5277	Hits (absolute): 1071	Hits (normalized): 117.446072584279	Hits (normalized) per kb: 22.256094669448	Normalized hits with 1T: 94.3%	Normalized hits with 10A: 15.9%	Normalized hits 25-32 nt: 99.7%	Normalized hits on the main strand(s): 94.2%	Predicted directionality: mono:minus
Cluster 351	Location: CM035924.1	Coordinates: 15187619-15189178	Size [bp]: 1560	Hits (absolute): 292	Hits (normalized): 123.900294465122	Hits (normalized) per kb: 79.4234500107209	Normalized hits with 1T: 0.4%	Normalized hits with 10A: 99.6%	Normalized hits 25-32 nt: 99%	Normalized hits on the main strand(s): 99.9%	Predicted directionality: mono:minus
Cluster 352	Location: CM035924.1	Coordinates: 17297211-17305025	Size [bp]: 7815	Hits (absolute): 1018	Hits (normalized): 111.743243620166	Hits (normalized) per kb: 14.298303754163	Normalized hits with 1T: 98.2%	Normalized hits with 10A: 10.4%	Normalized hits 25-32 nt: 99.9%	Normalized hits on the main strand(s): 99.4%	Predicted directionality: mono:plus
Cluster 353	Location: CM035924.1	Coordinates: 19512387-19518061	Size [bp]: 5675	Hits (absolute): 697	Hits (normalized): 173.024525290503	Hits (normalized) per kb: 30.4885572718669	Normalized hits with 1T: 97.7%	Normalized hits with 10A: 41.5%	Normalized hits 25-32 nt: 99.8%	Normalized hits on the main strand(s): 97%	Predicted directionality: mono:minus
Cluster 354	Location: CM035924.1	Coordinates: 27661048-27668356	Size [bp]: 7309	Hits (absolute): 1850	Hits (normalized): 157.184180091362	Hits (normalized) per kb: 21.505465173311	Normalized hits with 1T: 93%	Normalized hits with 10A: 29.1%	Normalized hits 25-32 nt: 99.8%	Normalized hits on the main strand(s): 95.9%	Predicted directionality: bi:plus-minus (split between 27667788 and 27667791)
Cluster 355	Location: CM035924.1	Coordinates: 29127905-29132423	Size [bp]: 4519	Hits (absolute): 496	Hits (normalized): 152.548413333967	Hits (normalized) per kb: 33.7573600218043	Normalized hits with 1T: 98%	Normalized hits with 10A: 57.3%	Normalized hits 25-32 nt: 99.7%	Normalized hits on the main strand(s): 98.5%	Predicted directionality: mono:plus
Cluster 356	Location: CM035924.1	Coordinates: 31298090-31303625	Size [bp]: 5536	Hits (absolute): 1505	Hits (normalized): 105.039942706589	Hits (normalized) per kb: 18.9740126267483	Normalized hits with 1T: 85.4%	Normalized hits with 10A: 36%	Normalized hits 25-32 nt: 99.5%	Normalized hits on the main strand(s): 85.1%	Predicted directionality: mono:plus
Cluster 357	Location: CM035924.1	Coordinates: 39048241-39050987	Size [bp]: 2747	Hits (absolute): 438	Hits (normalized): 150.606031468617	Hits (normalized) per kb: 54.825307444108	Normalized hits with 1T: 98%	Normalized hits with 10A: 55.8%	Normalized hits 25-32 nt: 99.7%	Normalized hits on the main strand(s): 97.3%	Predicted directionality: mono:minus
Cluster 358	Location: CM035924.1	Coordinates: 39501431-39505810	Size [bp]: 4380	Hits (absolute): 469	Hits (normalized): 128.340473469757	Hits (normalized) per kb: 29.3018078450134	Normalized hits with 1T: 97.6%	Normalized hits with 10A: 47%	Normalized hits 25-32 nt: 99.6%	Normalized hits on the main strand(s): 98.1%	Predicted directionality: mono:plus
Cluster 359	Location: CM035924.1	Coordinates: 41637338-41642293	Size [bp]: 4956	Hits (absolute): 369	Hits (normalized): 114.19693159077	Hits (normalized) per kb: 23.0423685829993	Normalized hits with 1T: 12.1%	Normalized hits with 10A: 76%	Normalized hits 25-32 nt: 99.2%	Normalized hits on the main strand(s): 99.9%	Predicted directionality: mono:plus
Cluster 360	Location: CM035924.1	Coordinates: 41828002-41833461	Size [bp]: 5460	Hits (absolute): 597	Hits (normalized): 109.620603607467	Hits (normalized) per kb: 20.0768928361563	Normalized hits with 1T: 98%	Normalized hits with 10A: 35.9%	Normalized hits 25-32 nt: 99.7%	Normalized hits on the main strand(s): 97%	Predicted directionality: mono:plus
Cluster 361	Location: CM035924.1	Coordinates: 42079360-42083990	Size [bp]: 4631	Hits (absolute): 1109	Hits (normalized): 265.38627078099	Hits (normalized) per kb: 57.3064384602034	Normalized hits with 1T: 98.5%	Normalized hits with 10A: 10.8%	Normalized hits 25-32 nt: 99.8%	Normalized hits on the main strand(s): 99.1%	Predicted directionality: mono:plus
Cluster 362	Location: CM035924.1	Coordinates: 60342148-60348549	Size [bp]: 6402	Hits (absolute): 586	Hits (normalized): 114.656779312869	Hits (normalized) per kb: 17.9095724753361	Normalized hits with 1T: 97%	Normalized hits with 10A: 16.7%	Normalized hits 25-32 nt: 99.9%	Normalized hits on the main strand(s): 99.6%	Predicted directionality: mono:minus
Cluster 363	Location: CM035924.1	Coordinates: 65104934-65109925	Size [bp]: 4992	Hits (absolute): 1496	Hits (normalized): 99.6813328006108	Hits (normalized) per kb: 19.9685617636226	Normalized hits with 1T: 84.5%	Normalized hits with 10A: 41.1%	Normalized hits 25-32 nt: 98.6%	Normalized hits on the main strand(s): 78.4%	Predicted directionality: mono:minus
Cluster 364	Location: CM035924.1	Coordinates: 65226962-65233652	Size [bp]: 6691	Hits (absolute): 2088	Hits (normalized): 161.00783163834	Hits (normalized) per kb: 24.063476305398	Normalized hits with 1T: 93%	Normalized hits with 10A: 28.6%	Normalized hits 25-32 nt: 99.9%	Normalized hits on the main strand(s): 95.8%	Predicted directionality: bi:plus-minus (split between 65233115 and 65233116)
Cluster 365	Location: CM035924.1	Coordinates: 78178045-78185012	Size [bp]: 6968	Hits (absolute): 587	Hits (normalized): 114.694540136756	Hits (normalized) per kb: 16.4600328338202	Normalized hits with 1T: 96.9%	Normalized hits with 10A: 16.8%	Normalized hits 25-32 nt: 99.9%	Normalized hits on the main strand(s): 99.1%	Predicted directionality: mono:minus
Cluster 366	Location: CM035924.1	Coordinates: 83745701-83749920	Size [bp]: 4220	Hits (absolute): 1474	Hits (normalized): 104.672335493853	Hits (normalized) per kb: 24.8036221493543	Normalized hits with 1T: 86.1%	Normalized hits with 10A: 36%	Normalized hits 25-32 nt: 99.3%	Normalized hits on the main strand(s): 84.8%	Predicted directionality: mono:minus
Cluster 367	Location: CM035924.1	Coordinates: 89211870-89219937	Size [bp]: 8068	Hits (absolute): 1046	Hits (normalized): 167.081276262385	Hits (normalized) per kb: 20.7094065177243	Normalized hits with 1T: 79.4%	Normalized hits with 10A: 25.9%	Normalized hits 25-32 nt: 99.2%	Normalized hits on the main strand(s): 86.1%	Predicted directionality: bi:minus-plus (split between 89215534 and 89215535)
Cluster 368	Location: CM035925.1	Coordinates: 326493-327856	Size [bp]: 1364	Hits (absolute): 259	Hits (normalized): 424.950886200131	Hits (normalized) per kb: 311.547584224421	Normalized hits with 1T: 71.3%	Normalized hits with 10A: 99.5%	Normalized hits 25-32 nt: 99.7%	Normalized hits on the main strand(s): 100%	Predicted directionality: mono:minus
Cluster 369	Location: CM035925.1	Coordinates: 1958657-1960308	Size [bp]: 1652	Hits (absolute): 256	Hits (normalized): 131.674724337136	Hits (normalized) per kb: 79.7065086195994	Normalized hits with 1T: 9.8%	Normalized hits with 10A: 98.8%	Normalized hits 25-32 nt: 99.8%	Normalized hits on the main strand(s): 100%	Predicted directionality: mono:plus
Cluster 370	Location: CM035925.1	Coordinates: 4318089-4324855	Size [bp]: 6767	Hits (absolute): 2046	Hits (normalized): 103.479736429157	Hits (normalized) per kb: 15.2921539808919	Normalized hits with 1T: 86.1%	Normalized hits with 10A: 32.2%	Normalized hits 25-32 nt: 99.5%	Normalized hits on the main strand(s): 82.6%	Predicted directionality: mono:minus
Cluster 371	Location: CM035925.1	Coordinates: 4508533-4511803	Size [bp]: 3271	Hits (absolute): 310	Hits (normalized): 170.81581097607	Hits (normalized) per kb: 52.221168242426	Normalized hits with 1T: 4.6%	Normalized hits with 10A: 95%	Normalized hits 25-32 nt: 99.6%	Normalized hits on the main strand(s): 99.9%	Predicted directionality: mono:plus
Cluster 372	Location: CM035925.1	Coordinates: 6355033-6356316	Size [bp]: 1284	Hits (absolute): 350	Hits (normalized): 539.651369441534	Hits (normalized) per kb: 420.288917013499	Normalized hits with 1T: 98.5%	Normalized hits with 10A: 1%	Normalized hits 25-32 nt: 100%	Normalized hits on the main strand(s): 100%	Predicted directionality: mono:plus
Cluster 373	Location: CM035925.1	Coordinates: 7680899-7689026	Size [bp]: 8128	Hits (absolute): 569	Hits (normalized): 104.16612499797	Hits (normalized) per kb: 12.8159153358142	Normalized hits with 1T: 97.8%	Normalized hits with 10A: 6.4%	Normalized hits 25-32 nt: 99.7%	Normalized hits on the main strand(s): 99.8%	Predicted directionality: mono:minus
Cluster 374	Location: CM035925.1	Coordinates: 7698017-7706471	Size [bp]: 8455	Hits (absolute): 1048	Hits (normalized): 150.80616077058	Hits (normalized) per kb: 17.8361869100714	Normalized hits with 1T: 96.5%	Normalized hits with 10A: 29.4%	Normalized hits 25-32 nt: 99.5%	Normalized hits on the main strand(s): 99.3%	Predicted directionality: mono:minus
Cluster 375	Location: CM035925.1	Coordinates: 7710028-7714878	Size [bp]: 4851	Hits (absolute): 714	Hits (normalized): 101.533447991037	Hits (normalized) per kb: 20.9302621236641	Normalized hits with 1T: 97.2%	Normalized hits with 10A: 22.6%	Normalized hits 25-32 nt: 99.8%	Normalized hits on the main strand(s): 99.7%	Predicted directionality: mono:minus
Cluster 376	Location: CM035925.1	Coordinates: 7725284-7735116	Size [bp]: 9833	Hits (absolute): 1967	Hits (normalized): 887.966636207715	Hits (normalized) per kb: 90.3047820641261	Normalized hits with 1T: 93.3%	Normalized hits with 10A: 7.4%	Normalized hits 25-32 nt: 99.8%	Normalized hits on the main strand(s): 100%	Predicted directionality: mono:minus
Cluster 377	Location: CM035925.1	Coordinates: 7937138-7946024	Size [bp]: 8887	Hits (absolute): 773	Hits (normalized): 238.727440561139	Hits (normalized) per kb: 26.8626114376408	Normalized hits with 1T: 99.4%	Normalized hits with 10A: 60.2%	Normalized hits 25-32 nt: 99.9%	Normalized hits on the main strand(s): 99.9%	Predicted directionality: mono:minus
Cluster 378	Location: CM035925.1	Coordinates: 7955075-7969601	Size [bp]: 14527	Hits (absolute): 1956	Hits (normalized): 639.102968427396	Hits (normalized) per kb: 43.9942969211701	Normalized hits with 1T: 95.2%	Normalized hits with 10A: 25.3%	Normalized hits 25-32 nt: 99.5%	Normalized hits on the main strand(s): 100%	Predicted directionality: mono:minus
Cluster 379	Location: CM035925.1	Coordinates: 8023698-8031902	Size [bp]: 8205	Hits (absolute): 646	Hits (normalized): 115.71212561318	Hits (normalized) per kb: 14.1026089134569	Normalized hits with 1T: 96.4%	Normalized hits with 10A: 14.7%	Normalized hits 25-32 nt: 99.8%	Normalized hits on the main strand(s): 99.4%	Predicted directionality: mono:minus
Cluster 380	Location: CM035925.1	Coordinates: 8035153-8073844	Size [bp]: 38692	Hits (absolute): 9752	Hits (normalized): 1586.08114258184	Hits (normalized) per kb: 40.9924778467674	Normalized hits with 1T: 92.8%	Normalized hits with 10A: 23.8%	Normalized hits 25-32 nt: 99.7%	Normalized hits on the main strand(s): 97.8%	Predicted directionality: mono:minus
Cluster 381	Location: CM035925.1	Coordinates: 8078058-8085021	Size [bp]: 6964	Hits (absolute): 1320	Hits (normalized): 100.959836816668	Hits (normalized) per kb: 14.497493145596	Normalized hits with 1T: 90.3%	Normalized hits with 10A: 19%	Normalized hits 25-32 nt: 99.6%	Normalized hits on the main strand(s): 99.7%	Predicted directionality: mono:minus
Cluster 382	Location: CM035925.1	Coordinates: 8100145-8108938	Size [bp]: 8794	Hits (absolute): 965	Hits (normalized): 140.620943126486	Hits (normalized) per kb: 15.9903652161255	Normalized hits with 1T: 97.1%	Normalized hits with 10A: 31.7%	Normalized hits 25-32 nt: 99.8%	Normalized hits on the main strand(s): 100%	Predicted directionality: mono:minus
Cluster 383	Location: CM035925.1	Coordinates: 9352618-9370022	Size [bp]: 17405	Hits (absolute): 2379	Hits (normalized): 395.177975632443	Hits (normalized) per kb: 22.7047949827813	Normalized hits with 1T: 85.8%	Normalized hits with 10A: 26.6%	Normalized hits 25-32 nt: 99.3%	Normalized hits on the main strand(s): 99.4%	Predicted directionality: mono:minus
Cluster 384	Location: CM035925.1	Coordinates: 9395034-9404961	Size [bp]: 9928	Hits (absolute): 2099	Hits (normalized): 284.05518449136	Hits (normalized) per kb: 28.6112846213789	Normalized hits with 1T: 93.4%	Normalized hits with 10A: 10.6%	Normalized hits 25-32 nt: 99.3%	Normalized hits on the main strand(s): 99.5%	Predicted directionality: mono:minus
Cluster 385	Location: CM035925.1	Coordinates: 9518024-9528023	Size [bp]: 10000	Hits (absolute): 1834	Hits (normalized): 268.076445732842	Hits (normalized) per kb: 26.8073975361558	Normalized hits with 1T: 93.7%	Normalized hits with 10A: 9.7%	Normalized hits 25-32 nt: 99.3%	Normalized hits on the main strand(s): 99.9%	Predicted directionality: mono:plus
Cluster 386	Location: CM035925.1	Coordinates: 9549021-9559923	Size [bp]: 10903	Hits (absolute): 1613	Hits (normalized): 248.753325558882	Hits (normalized) per kb: 22.8152227857511	Normalized hits with 1T: 84.7%	Normalized hits with 10A: 28.5%	Normalized hits 25-32 nt: 98.8%	Normalized hits on the main strand(s): 99.8%	Predicted directionality: mono:plus
Cluster 387	Location: CM035925.1	Coordinates: 12332442-12337588	Size [bp]: 5147	Hits (absolute): 225	Hits (normalized): 101.065168182136	Hits (normalized) per kb: 19.6358805344222	Normalized hits with 1T: 91.2%	Normalized hits with 10A: 8.6%	Normalized hits 25-32 nt: 99.8%	Normalized hits on the main strand(s): 99.5%	Predicted directionality: bi:plus-minus (split between 12334100 and 12334131)
Cluster 388	Location: CM035925.1	Coordinates: 12546260-12551486	Size [bp]: 5227	Hits (absolute): 1509	Hits (normalized): 106.427864435209	Hits (normalized) per kb: 20.3613492653256	Normalized hits with 1T: 86.2%	Normalized hits with 10A: 35.6%	Normalized hits 25-32 nt: 99.3%	Normalized hits on the main strand(s): 84.5%	Predicted directionality: mono:minus
Cluster 389	Location: CM035925.1	Coordinates: 17593667-17603025	Size [bp]: 9359	Hits (absolute): 2869	Hits (normalized): 198.877678086038	Hits (normalized) per kb: 21.2496640601022	Normalized hits with 1T: 91.7%	Normalized hits with 10A: 29.3%	Normalized hits 25-32 nt: 99.2%	Normalized hits on the main strand(s): 99.8%	Predicted directionality: mono:plus
Cluster 390	Location: CM035925.1	Coordinates: 17781027-17789937	Size [bp]: 8911	Hits (absolute): 1340	Hits (normalized): 239.773672473437	Hits (normalized) per kb: 26.907341686945	Normalized hits with 1T: 97.7%	Normalized hits with 10A: 56.4%	Normalized hits 25-32 nt: 99.4%	Normalized hits on the main strand(s): 99.3%	Predicted directionality: mono:minus
Cluster 391	Location: CM035925.1	Coordinates: 19285773-19293880	Size [bp]: 8108	Hits (absolute): 832	Hits (normalized): 167.405658624367	Hits (normalized) per kb: 20.6472035147856	Normalized hits with 1T: 96.7%	Normalized hits with 10A: 13.3%	Normalized hits 25-32 nt: 98%	Normalized hits on the main strand(s): 99.8%	Predicted directionality: mono:plus
Cluster 392	Location: CM035925.1	Coordinates: 27043138-27095560	Size [bp]: 52423	Hits (absolute): 16494	Hits (normalized): 3242.78160297326	Hits (normalized) per kb: 61.8577413269113	Normalized hits with 1T: 88.8%	Normalized hits with 10A: 23.8%	Normalized hits 25-32 nt: 99.5%	Normalized hits on the main strand(s): 100%	Predicted directionality: mono:plus
Cluster 393	Location: CM035925.1	Coordinates: 28258001-28324014	Size [bp]: 66014	Hits (absolute): 27459	Hits (normalized): 9026.18623020108	Hits (normalized) per kb: 136.731286291215	Normalized hits with 1T: 85%	Normalized hits with 10A: 18.9%	Normalized hits 25-32 nt: 99.7%	Normalized hits on the main strand(s): 100%	Predicted directionality: mono:plus
Cluster 394	Location: CM035925.1	Coordinates: 38010336-38017642	Size [bp]: 7307	Hits (absolute): 2004	Hits (normalized): 152.480580638831	Hits (normalized) per kb: 20.8680591207253	Normalized hits with 1T: 96.2%	Normalized hits with 10A: 26.8%	Normalized hits 25-32 nt: 99.9%	Normalized hits on the main strand(s): 95.5%	Predicted directionality: bi:plus-minus (split between 38017053 and 38017095)
Cluster 395	Location: CM035925.1	Coordinates: 43416002-43423981	Size [bp]: 7980	Hits (absolute): 793	Hits (normalized): 97.9127998452791	Hits (normalized) per kb: 12.2700665122732	Normalized hits with 1T: 98.4%	Normalized hits with 10A: 22.4%	Normalized hits 25-32 nt: 100%	Normalized hits on the main strand(s): 98.8%	Predicted directionality: mono:plus
Cluster 396	Location: CM035925.1	Coordinates: 43457232-43487018	Size [bp]: 29787	Hits (absolute): 9183	Hits (normalized): 2593.27085258171	Hits (normalized) per kb: 87.060441169277	Normalized hits with 1T: 96.8%	Normalized hits with 10A: 15.5%	Normalized hits 25-32 nt: 99.6%	Normalized hits on the main strand(s): 98.8%	Predicted directionality: mono:plus
Cluster 397	Location: CM035925.1	Coordinates: 43488056-43496958	Size [bp]: 8903	Hits (absolute): 1674	Hits (normalized): 263.300969523023	Hits (normalized) per kb: 29.5743828017112	Normalized hits with 1T: 97.3%	Normalized hits with 10A: 35.1%	Normalized hits 25-32 nt: 99.7%	Normalized hits on the main strand(s): 99%	Predicted directionality: mono:plus
Cluster 398	Location: CM035925.1	Coordinates: 43500099-43520016	Size [bp]: 19918	Hits (absolute): 6342	Hits (normalized): 938.105311628738	Hits (normalized) per kb: 47.0981568767981	Normalized hits with 1T: 94.5%	Normalized hits with 10A: 21.2%	Normalized hits 25-32 nt: 99.7%	Normalized hits on the main strand(s): 98.7%	Predicted directionality: mono:plus
Cluster 399	Location: CM035925.1	Coordinates: 48364010-48373012	Size [bp]: 9003	Hits (absolute): 1127	Hits (normalized): 166.184073826463	Hits (normalized) per kb: 18.458915849604	Normalized hits with 1T: 97.2%	Normalized hits with 10A: 34.8%	Normalized hits 25-32 nt: 99.8%	Normalized hits on the main strand(s): 99.6%	Predicted directionality: mono:plus
Cluster 400	Location: CM035925.1	Coordinates: 48924009-48929509	Size [bp]: 5501	Hits (absolute): 748	Hits (normalized): 143.807390850259	Hits (normalized) per kb: 26.142035077755	Normalized hits with 1T: 98.1%	Normalized hits with 10A: 37.9%	Normalized hits 25-32 nt: 99.7%	Normalized hits on the main strand(s): 100%	Predicted directionality: mono:minus
Cluster 401	Location: CM035925.1	Coordinates: 52642035-52660014	Size [bp]: 17980	Hits (absolute): 2715	Hits (normalized): 785.111542809892	Hits (normalized) per kb: 43.665809152842	Normalized hits with 1T: 87.4%	Normalized hits with 10A: 20.8%	Normalized hits 25-32 nt: 98.7%	Normalized hits on the main strand(s): 99.2%	Predicted directionality: mono:plus
Cluster 402	Location: CM035925.1	Coordinates: 52672056-52680801	Size [bp]: 8746	Hits (absolute): 1059	Hits (normalized): 227.242591283245	Hits (normalized) per kb: 25.9826835646086	Normalized hits with 1T: 89.2%	Normalized hits with 10A: 38.3%	Normalized hits 25-32 nt: 99.8%	Normalized hits on the main strand(s): 100%	Predicted directionality: mono:plus
Cluster 403	Location: CM035925.1	Coordinates: 52805226-52814005	Size [bp]: 8780	Hits (absolute): 1403	Hits (normalized): 277.718857277254	Hits (normalized) per kb: 31.6305764494161	Normalized hits with 1T: 95.4%	Normalized hits with 10A: 36.9%	Normalized hits 25-32 nt: 96.7%	Normalized hits on the main strand(s): 99.5%	Predicted directionality: mono:plus
Cluster 404	Location: CM035925.1	Coordinates: 52817003-52826029	Size [bp]: 9027	Hits (absolute): 850	Hits (normalized): 146.044680381923	Hits (normalized) per kb: 16.1783720452324	Normalized hits with 1T: 96.6%	Normalized hits with 10A: 8.1%	Normalized hits 25-32 nt: 99.8%	Normalized hits on the main strand(s): 99.7%	Predicted directionality: mono:plus
Cluster 405	Location: CM035925.1	Coordinates: 52842008-52851548	Size [bp]: 9541	Hits (absolute): 1978	Hits (normalized): 297.281838024387	Hits (normalized) per kb: 31.1581131911399	Normalized hits with 1T: 90.6%	Normalized hits with 10A: 35.5%	Normalized hits 25-32 nt: 99.6%	Normalized hits on the main strand(s): 97.7%	Predicted directionality: mono:plus
Cluster 406	Location: CM035925.1	Coordinates: 52860005-52867026	Size [bp]: 7022	Hits (absolute): 1324	Hits (normalized): 114.494761105838	Hits (normalized) per kb: 16.304874781546	Normalized hits with 1T: 97.9%	Normalized hits with 10A: 14.2%	Normalized hits 25-32 nt: 99.6%	Normalized hits on the main strand(s): 99.6%	Predicted directionality: mono:plus
Cluster 407	Location: CM035925.1	Coordinates: 53032073-53040027	Size [bp]: 7955	Hits (absolute): 513	Hits (normalized): 117.394821521237	Hits (normalized) per kb: 14.757487719677	Normalized hits with 1T: 95.3%	Normalized hits with 10A: 30%	Normalized hits 25-32 nt: 99.7%	Normalized hits on the main strand(s): 99.8%	Predicted directionality: mono:plus
Cluster 408	Location: CM035925.1	Coordinates: 53081021-53085542	Size [bp]: 4522	Hits (absolute): 984	Hits (normalized): 99.4048916898798	Hits (normalized) per kb: 21.9828208026048	Normalized hits with 1T: 97.2%	Normalized hits with 10A: 36.9%	Normalized hits 25-32 nt: 98.5%	Normalized hits on the main strand(s): 100%	Predicted directionality: mono:plus
Cluster 409	Location: CM035925.1	Coordinates: 53095053-53104985	Size [bp]: 9933	Hits (absolute): 1107	Hits (normalized): 185.294581213228	Hits (normalized) per kb: 18.6546106903101	Normalized hits with 1T: 93.4%	Normalized hits with 10A: 22.7%	Normalized hits 25-32 nt: 99.9%	Normalized hits on the main strand(s): 99.9%	Predicted directionality: mono:plus
Cluster 410	Location: CM035925.1	Coordinates: 55940063-55993988	Size [bp]: 53926	Hits (absolute): 16014	Hits (normalized): 5850.58288269257	Hits (normalized) per kb: 108.492520777323	Normalized hits with 1T: 95.4%	Normalized hits with 10A: 25.9%	Normalized hits 25-32 nt: 99.8%	Normalized hits on the main strand(s): 99.2%	Predicted directionality: mono:minus
Cluster 411	Location: CM035925.1	Coordinates: 56000082-56013999	Size [bp]: 13918	Hits (absolute): 1391	Hits (normalized): 458.729565036403	Hits (normalized) per kb: 32.9592046357815	Normalized hits with 1T: 97.4%	Normalized hits with 10A: 31.9%	Normalized hits 25-32 nt: 99.9%	Normalized hits on the main strand(s): 100%	Predicted directionality: mono:minus
Cluster 412	Location: CM035925.1	Coordinates: 70625337-70633016	Size [bp]: 7680	Hits (absolute): 1366	Hits (normalized): 149.173957273384	Hits (normalized) per kb: 19.423411850227	Normalized hits with 1T: 83.2%	Normalized hits with 10A: 20.1%	Normalized hits 25-32 nt: 99.4%	Normalized hits on the main strand(s): 82.4%	Predicted directionality: mono:plus
Cluster 413	Location: CM035925.1	Coordinates: 78527515-78530619	Size [bp]: 3105	Hits (absolute): 559	Hits (normalized): 102.62167555614	Hits (normalized) per kb: 33.0507618648262	Normalized hits with 1T: 9.7%	Normalized hits with 10A: 94.5%	Normalized hits 25-32 nt: 99.4%	Normalized hits on the main strand(s): 95.3%	Predicted directionality: mono:minus
Cluster 414	Location: CM035925.1	Coordinates: 81355396-81361651	Size [bp]: 6256	Hits (absolute): 898	Hits (normalized): 107.145993154334	Hits (normalized) per kb: 17.1267931125117	Normalized hits with 1T: 94.5%	Normalized hits with 10A: 17.2%	Normalized hits 25-32 nt: 99.7%	Normalized hits on the main strand(s): 94.3%	Predicted directionality: mono:plus
Cluster 415	Location: CM035925.1	Coordinates: 86730041-86739026	Size [bp]: 8986	Hits (absolute): 1304	Hits (normalized): 131.143711265025	Hits (normalized) per kb: 14.5939427456583	Normalized hits with 1T: 91.6%	Normalized hits with 10A: 14.4%	Normalized hits 25-32 nt: 99.5%	Normalized hits on the main strand(s): 91%	Predicted directionality: mono:plus
Cluster 416	Location: CM035926.1	Coordinates: 127156-131230	Size [bp]: 4075	Hits (absolute): 413	Hits (normalized): 96.4327288722102	Hits (normalized) per kb: 23.6643986123866	Normalized hits with 1T: 92.7%	Normalized hits with 10A: 81.8%	Normalized hits 25-32 nt: 99.5%	Normalized hits on the main strand(s): 97.4%	Predicted directionality: mono:plus
Cluster 417	Location: CM035926.1	Coordinates: 6577489-6584871	Size [bp]: 7383	Hits (absolute): 1217	Hits (normalized): 213.497315334704	Hits (normalized) per kb: 28.9174072650549	Normalized hits with 1T: 94.5%	Normalized hits with 10A: 18.9%	Normalized hits 25-32 nt: 98.3%	Normalized hits on the main strand(s): 98.2%	Predicted directionality: mono:plus
Cluster 418	Location: CM035926.1	Coordinates: 9781101-9789992	Size [bp]: 8892	Hits (absolute): 1804	Hits (normalized): 137.014889406202	Hits (normalized) per kb: 15.4088719751702	Normalized hits with 1T: 89.4%	Normalized hits with 10A: 19.2%	Normalized hits 25-32 nt: 99.3%	Normalized hits on the main strand(s): 91.7%	Predicted directionality: mono:minus
Cluster 419	Location: CM035926.1	Coordinates: 17079036-17085966	Size [bp]: 6931	Hits (absolute): 578	Hits (normalized): 114.447362547889	Hits (normalized) per kb: 16.5124510947236	Normalized hits with 1T: 96.9%	Normalized hits with 10A: 16.6%	Normalized hits 25-32 nt: 99.9%	Normalized hits on the main strand(s): 99.5%	Predicted directionality: mono:minus
Cluster 420	Location: CM035926.1	Coordinates: 17925052-17929750	Size [bp]: 4699	Hits (absolute): 1588	Hits (normalized): 107.057705569028	Hits (normalized) per kb: 22.7830729190637	Normalized hits with 1T: 86%	Normalized hits with 10A: 35.5%	Normalized hits 25-32 nt: 99.3%	Normalized hits on the main strand(s): 84.6%	Predicted directionality: mono:plus
Cluster 421	Location: CM035926.1	Coordinates: 19964694-19971770	Size [bp]: 7077	Hits (absolute): 1238	Hits (normalized): 287.237612813237	Hits (normalized) per kb: 40.587808872593	Normalized hits with 1T: 98.6%	Normalized hits with 10A: 10.6%	Normalized hits 25-32 nt: 99.8%	Normalized hits on the main strand(s): 99.1%	Predicted directionality: mono:plus
Cluster 422	Location: CM035926.1	Coordinates: 36529278-36538022	Size [bp]: 8745	Hits (absolute): 815	Hits (normalized): 151.085153401936	Hits (normalized) per kb: 17.2770587937682	Normalized hits with 1T: 96.8%	Normalized hits with 10A: 5.6%	Normalized hits 25-32 nt: 99.7%	Normalized hits on the main strand(s): 100%	Predicted directionality: mono:minus
Cluster 423	Location: CM035926.1	Coordinates: 53568049-53573024	Size [bp]: 4976	Hits (absolute): 499	Hits (normalized): 96.1375304920023	Hits (normalized) per kb: 19.3199731487109	Normalized hits with 1T: 98.9%	Normalized hits with 10A: 3.6%	Normalized hits 25-32 nt: 99.9%	Normalized hits on the main strand(s): 99.7%	Predicted directionality: mono:minus
Cluster 424	Location: CM035926.1	Coordinates: 56872008-56879634	Size [bp]: 7627	Hits (absolute): 968	Hits (normalized): 136.12499606816	Hits (normalized) per kb: 17.8480683825428	Normalized hits with 1T: 88.9%	Normalized hits with 10A: 27.8%	Normalized hits 25-32 nt: 99.8%	Normalized hits on the main strand(s): 99.3%	Predicted directionality: mono:plus
Cluster 425	Location: CM035926.1	Coordinates: 56896008-56923925	Size [bp]: 27918	Hits (absolute): 3543	Hits (normalized): 481.856713953276	Hits (normalized) per kb: 17.2595860401337	Normalized hits with 1T: 98%	Normalized hits with 10A: 15.1%	Normalized hits 25-32 nt: 99.8%	Normalized hits on the main strand(s): 99.7%	Predicted directionality: mono:plus
Cluster 426	Location: CM035926.1	Coordinates: 56950095-56958958	Size [bp]: 8864	Hits (absolute): 1454	Hits (normalized): 155.867286215944	Hits (normalized) per kb: 17.5845792577349	Normalized hits with 1T: 83.6%	Normalized hits with 10A: 19.8%	Normalized hits 25-32 nt: 99.2%	Normalized hits on the main strand(s): 82.9%	Predicted directionality: mono:plus
Cluster 427	Location: CM035926.1	Coordinates: 56976007-56985024	Size [bp]: 9018	Hits (absolute): 1611	Hits (normalized): 214.167518196556	Hits (normalized) per kb: 23.7489667399774	Normalized hits with 1T: 80.2%	Normalized hits with 10A: 18.7%	Normalized hits 25-32 nt: 99.4%	Normalized hits on the main strand(s): 99.6%	Predicted directionality: mono:plus
Cluster 428	Location: CM035926.1	Coordinates: 56987313-56994392	Size [bp]: 7080	Hits (absolute): 910	Hits (normalized): 135.435875750926	Hits (normalized) per kb: 19.1291706790224	Normalized hits with 1T: 93.2%	Normalized hits with 10A: 11.6%	Normalized hits 25-32 nt: 99.9%	Normalized hits on the main strand(s): 99.8%	Predicted directionality: mono:plus
Cluster 429	Location: CM035926.1	Coordinates: 63970037-63974680	Size [bp]: 4644	Hits (absolute): 444	Hits (normalized): 94.5270817187972	Hits (normalized) per kb: 20.3543601638718	Normalized hits with 1T: 99.1%	Normalized hits with 10A: 3.5%	Normalized hits 25-32 nt: 99.9%	Normalized hits on the main strand(s): 99.8%	Predicted directionality: mono:plus
Cluster 430	Location: CM035926.1	Coordinates: 67036807-67041090	Size [bp]: 4284	Hits (absolute): 1117	Hits (normalized): 119.120123242733	Hits (normalized) per kb: 27.8061401339024	Normalized hits with 1T: 94.2%	Normalized hits with 10A: 16.3%	Normalized hits 25-32 nt: 99.7%	Normalized hits on the main strand(s): 94.1%	Predicted directionality: mono:minus
Cluster 431	Location: CM035926.1	Coordinates: 71558571-71593540	Size [bp]: 34970	Hits (absolute): 19348	Hits (normalized): 7861.75577709463	Hits (normalized) per kb: 224.814233003181	Normalized hits with 1T: 76.7%	Normalized hits with 10A: 17.5%	Normalized hits 25-32 nt: 99.7%	Normalized hits on the main strand(s): 100%	Predicted directionality: mono:minus
Cluster 432	Location: CM035927.1	Coordinates: 24002153-24010831	Size [bp]: 8679	Hits (absolute): 2252	Hits (normalized): 167.286925199254	Hits (normalized) per kb: 19.2752428994067	Normalized hits with 1T: 77.6%	Normalized hits with 10A: 25.5%	Normalized hits 25-32 nt: 98.9%	Normalized hits on the main strand(s): 76.7%	Predicted directionality: mono:minus
Cluster 433	Location: CM035927.1	Coordinates: 34524487-34530963	Size [bp]: 6477	Hits (absolute): 284	Hits (normalized): 108.034655940265	Hits (normalized) per kb: 16.6794906194692	Normalized hits with 1T: 98.8%	Normalized hits with 10A: 88.3%	Normalized hits 25-32 nt: 99.5%	Normalized hits on the main strand(s): 99.6%	Predicted directionality: mono:minus
Cluster 434	Location: CM035927.1	Coordinates: 34659924-34663771	Size [bp]: 3848	Hits (absolute): 259	Hits (normalized): 107.380236332043	Hits (normalized) per kb: 27.9053853745462	Normalized hits with 1T: 98.9%	Normalized hits with 10A: 88.8%	Normalized hits 25-32 nt: 99.7%	Normalized hits on the main strand(s): 99.7%	Predicted directionality: mono:minus
Cluster 435	Location: CM035927.1	Coordinates: 53592363-53596798	Size [bp]: 4436	Hits (absolute): 1506	Hits (normalized): 105.384398888699	Hits (normalized) per kb: 23.7566547515766	Normalized hits with 1T: 86.6%	Normalized hits with 10A: 35.3%	Normalized hits 25-32 nt: 99.3%	Normalized hits on the main strand(s): 84.5%	Predicted directionality: mono:minus
Cluster 436	Location: CM035927.1	Coordinates: 66064664-66071024	Size [bp]: 6361	Hits (absolute): 1182	Hits (normalized): 224.124211707432	Hits (normalized) per kb: 35.2341571589901	Normalized hits with 1T: 93.9%	Normalized hits with 10A: 58.9%	Normalized hits 25-32 nt: 99.7%	Normalized hits on the main strand(s): 94.4%	Predicted directionality: mono:minus
Cluster 437	Location: CM035927.1	Coordinates: 74860076-74864869	Size [bp]: 4794	Hits (absolute): 740	Hits (normalized): 147.069620882852	Hits (normalized) per kb: 30.6779619212646	Normalized hits with 1T: 91.2%	Normalized hits with 10A: 28.6%	Normalized hits 25-32 nt: 98.3%	Normalized hits on the main strand(s): 93.3%	Predicted directionality: bi:minus-plus (split between 74860705 and 74860710)
Cluster 438	Location: CM035928.1	Coordinates: 306881-308647	Size [bp]: 1767	Hits (absolute): 515	Hits (normalized): 180.291507951099	Hits (normalized) per kb: 102.032494303585	Normalized hits with 1T: 99%	Normalized hits with 10A: 48.5%	Normalized hits 25-32 nt: 99.6%	Normalized hits on the main strand(s): 99.1%	Predicted directionality: mono:plus
Cluster 439	Location: CM035928.1	Coordinates: 3860735-3865791	Size [bp]: 5057	Hits (absolute): 1571	Hits (normalized): 108.299922502274	Hits (normalized) per kb: 21.4160046747024	Normalized hits with 1T: 86.4%	Normalized hits with 10A: 35.3%	Normalized hits 25-32 nt: 99.3%	Normalized hits on the main strand(s): 85%	Predicted directionality: mono:minus
Cluster 440	Location: CM035928.1	Coordinates: 11098016-11105625	Size [bp]: 7610	Hits (absolute): 764	Hits (normalized): 432.958534459342	Hits (normalized) per kb: 56.8933825642844	Normalized hits with 1T: 99%	Normalized hits with 10A: 1.1%	Normalized hits 25-32 nt: 100%	Normalized hits on the main strand(s): 99.9%	Predicted directionality: bi:plus-minus (split between 11102331 and 11102675)
Cluster 441	Location: CM035928.1	Coordinates: 14190594-14196996	Size [bp]: 6403	Hits (absolute): 1196	Hits (normalized): 149.967733373382	Hits (normalized) per kb: 23.4211778817947	Normalized hits with 1T: 96%	Normalized hits with 10A: 13.2%	Normalized hits 25-32 nt: 99.8%	Normalized hits on the main strand(s): 97.3%	Predicted directionality: mono:plus
Cluster 442	Location: CM035928.1	Coordinates: 15324537-15332961	Size [bp]: 8425	Hits (absolute): 305	Hits (normalized): 179.55979281582	Hits (normalized) per kb: 21.3125659731864	Normalized hits with 1T: 99.2%	Normalized hits with 10A: 1.1%	Normalized hits 25-32 nt: 99.9%	Normalized hits on the main strand(s): 100%	Predicted directionality: mono:minus
Cluster 443	Location: CM035928.1	Coordinates: 15334049-15342606	Size [bp]: 8558	Hits (absolute): 888	Hits (normalized): 125.197542925335	Hits (normalized) per kb: 14.6295871630726	Normalized hits with 1T: 95.6%	Normalized hits with 10A: 31.5%	Normalized hits 25-32 nt: 99.6%	Normalized hits on the main strand(s): 100%	Predicted directionality: mono:minus
Cluster 444	Location: CM035928.1	Coordinates: 16623549-16627846	Size [bp]: 4298	Hits (absolute): 1624	Hits (normalized): 109.376191590693	Hits (normalized) per kb: 25.4480173033937	Normalized hits with 1T: 85.3%	Normalized hits with 10A: 35.8%	Normalized hits 25-32 nt: 99.3%	Normalized hits on the main strand(s): 84.9%	Predicted directionality: mono:minus
Cluster 445	Location: CM035928.1	Coordinates: 19019582-19027918	Size [bp]: 8337	Hits (absolute): 979	Hits (normalized): 742.449671973265	Hits (normalized) per kb: 89.0544318140432	Normalized hits with 1T: 99.1%	Normalized hits with 10A: 2%	Normalized hits 25-32 nt: 99.9%	Normalized hits on the main strand(s): 99.7%	Predicted directionality: bi:plus-minus (split between 19023698 and 19023706)
Cluster 446	Location: CM035928.1	Coordinates: 28537051-28542859	Size [bp]: 5809	Hits (absolute): 1159	Hits (normalized): 107.58196072387	Hits (normalized) per kb: 18.519721032252	Normalized hits with 1T: 92%	Normalized hits with 10A: 24.9%	Normalized hits 25-32 nt: 99.6%	Normalized hits on the main strand(s): 93.8%	Predicted directionality: mono:minus
Cluster 447	Location: CM035928.1	Coordinates: 31189102-31194960	Size [bp]: 5859	Hits (absolute): 1665	Hits (normalized): 133.775478801383	Hits (normalized) per kb: 22.8326955393856	Normalized hits with 1T: 97.1%	Normalized hits with 10A: 29%	Normalized hits 25-32 nt: 99.8%	Normalized hits on the main strand(s): 98.8%	Predicted directionality: mono:plus
Cluster 448	Location: CM035928.1	Coordinates: 32274488-32287996	Size [bp]: 13509	Hits (absolute): 3141	Hits (normalized): 332.832899756527	Hits (normalized) per kb: 24.6379804448995	Normalized hits with 1T: 79%	Normalized hits with 10A: 25.2%	Normalized hits 25-32 nt: 99%	Normalized hits on the main strand(s): 81.9%	Predicted directionality: bi:minus-plus (split between 32283743 and 32283743)
Cluster 449	Location: CM035928.1	Coordinates: 33890015-33895558	Size [bp]: 5544	Hits (absolute): 997	Hits (normalized): 108.75481448557	Hits (normalized) per kb: 19.617009960497	Normalized hits with 1T: 95.1%	Normalized hits with 10A: 15.8%	Normalized hits 25-32 nt: 99.8%	Normalized hits on the main strand(s): 95%	Predicted directionality: mono:minus
Cluster 450	Location: CM035928.1	Coordinates: 34919619-34921577	Size [bp]: 1959	Hits (absolute): 290	Hits (normalized): 301.845525671793	Hits (normalized) per kb: 154.081730650248	Normalized hits with 1T: 0.3%	Normalized hits with 10A: 99.8%	Normalized hits 25-32 nt: 99.9%	Normalized hits on the main strand(s): 100%	Predicted directionality: mono:minus
Cluster 451	Location: CM035928.1	Coordinates: 43597015-43601509	Size [bp]: 4495	Hits (absolute): 647	Hits (normalized): 96.3982506238843	Hits (normalized) per kb: 21.4453589008084	Normalized hits with 1T: 94.6%	Normalized hits with 10A: 19.3%	Normalized hits 25-32 nt: 96.5%	Normalized hits on the main strand(s): 99.9%	Predicted directionality: mono:minus
Cluster 452	Location: CM035928.1	Coordinates: 44560336-44564513	Size [bp]: 4178	Hits (absolute): 1039	Hits (normalized): 219.032865614177	Hits (normalized) per kb: 52.4252500048766	Normalized hits with 1T: 94.1%	Normalized hits with 10A: 58%	Normalized hits 25-32 nt: 99.7%	Normalized hits on the main strand(s): 94.2%	Predicted directionality: mono:plus
Cluster 453	Location: CM035928.1	Coordinates: 45072870-45076154	Size [bp]: 3285	Hits (absolute): 1058	Hits (normalized): 226.005582673845	Hits (normalized) per kb: 68.7993168908152	Normalized hits with 1T: 95.2%	Normalized hits with 10A: 58.6%	Normalized hits 25-32 nt: 99.7%	Normalized hits on the main strand(s): 95.2%	Predicted directionality: mono:plus
Cluster 454	Location: CM035928.1	Coordinates: 47184020-47193024	Size [bp]: 9005	Hits (absolute): 982	Hits (normalized): 180.658249184486	Hits (normalized) per kb: 20.0622157231034	Normalized hits with 1T: 95.8%	Normalized hits with 10A: 8.1%	Normalized hits 25-32 nt: 99.8%	Normalized hits on the main strand(s): 100%	Predicted directionality: mono:minus
Cluster 455	Location: CM035928.1	Coordinates: 47195116-47204027	Size [bp]: 8912	Hits (absolute): 1480	Hits (normalized): 188.874490060291	Hits (normalized) per kb: 21.1930523383266	Normalized hits with 1T: 97%	Normalized hits with 10A: 21.9%	Normalized hits 25-32 nt: 99.8%	Normalized hits on the main strand(s): 99.7%	Predicted directionality: mono:minus
Cluster 456	Location: CM035928.1	Coordinates: 47938165-47943296	Size [bp]: 5132	Hits (absolute): 1172	Hits (normalized): 566.813785343124	Hits (normalized) per kb: 110.446673543803	Normalized hits with 1T: 12.3%	Normalized hits with 10A: 90.1%	Normalized hits 25-32 nt: 99.8%	Normalized hits on the main strand(s): 90.2%	Predicted directionality: bi:minus-plus (split between 47940626 and 47940627)
Cluster 457	Location: CM035928.1	Coordinates: 48500020-48510028	Size [bp]: 10009	Hits (absolute): 2180	Hits (normalized): 257.576763860585	Hits (normalized) per kb: 25.7345704629991	Normalized hits with 1T: 96.1%	Normalized hits with 10A: 21.9%	Normalized hits 25-32 nt: 99.8%	Normalized hits on the main strand(s): 100%	Predicted directionality: mono:minus
Cluster 458	Location: CM035928.1	Coordinates: 48805020-48813979	Size [bp]: 8960	Hits (absolute): 1794	Hits (normalized): 208.729122943665	Hits (normalized) per kb: 23.2953740556265	Normalized hits with 1T: 95.9%	Normalized hits with 10A: 23.5%	Normalized hits 25-32 nt: 99.8%	Normalized hits on the main strand(s): 100%	Predicted directionality: mono:minus
Cluster 459	Location: CM035928.1	Coordinates: 48815082-48826015	Size [bp]: 10934	Hits (absolute): 3225	Hits (normalized): 669.384380505674	Hits (normalized) per kb: 61.2203352743256	Normalized hits with 1T: 92.5%	Normalized hits with 10A: 18.1%	Normalized hits 25-32 nt: 99.8%	Normalized hits on the main strand(s): 99.8%	Predicted directionality: mono:minus
Cluster 460	Location: CM035928.1	Coordinates: 52718067-52724673	Size [bp]: 6607	Hits (absolute): 1217	Hits (normalized): 168.798672537104	Hits (normalized) per kb: 25.5486603643283	Normalized hits with 1T: 78.9%	Normalized hits with 10A: 26.4%	Normalized hits 25-32 nt: 99%	Normalized hits on the main strand(s): 85.9%	Predicted directionality: bi:minus-plus (split between 52722862 and 52722862)
Cluster 461	Location: CM035928.1	Coordinates: 56906300-56915954	Size [bp]: 9655	Hits (absolute): 1375	Hits (normalized): 212.851056470268	Hits (normalized) per kb: 22.0457227156889	Normalized hits with 1T: 97.5%	Normalized hits with 10A: 24.4%	Normalized hits 25-32 nt: 99.7%	Normalized hits on the main strand(s): 99.5%	Predicted directionality: mono:plus
Cluster 462	Location: CM035928.1	Coordinates: 61851182-61855815	Size [bp]: 4634	Hits (absolute): 689	Hits (normalized): 164.185141895739	Hits (normalized) per kb: 35.4305509098416	Normalized hits with 1T: 96.9%	Normalized hits with 10A: 57.3%	Normalized hits 25-32 nt: 99.8%	Normalized hits on the main strand(s): 97.3%	Predicted directionality: mono:plus
Cluster 463	Location: CM035928.1	Coordinates: 67295151-67299850	Size [bp]: 4700	Hits (absolute): 1044	Hits (normalized): 282.705596881468	Hits (normalized) per kb: 60.1503038417504	Normalized hits with 1T: 98.6%	Normalized hits with 10A: 10.5%	Normalized hits 25-32 nt: 99.8%	Normalized hits on the main strand(s): 99.1%	Predicted directionality: mono:minus
Cluster 464	Location: CM035928.1	Coordinates: 70408064-70422965	Size [bp]: 14902	Hits (absolute): 3323	Hits (normalized): 328.259967932971	Hits (normalized) per kb: 22.0282499620544	Normalized hits with 1T: 95.7%	Normalized hits with 10A: 15.2%	Normalized hits 25-32 nt: 99.7%	Normalized hits on the main strand(s): 100%	Predicted directionality: mono:minus
Cluster 465	Location: CM035929.1	Coordinates: 3735170-3742958	Size [bp]: 7789	Hits (absolute): 1578	Hits (normalized): 106.949902089056	Hits (normalized) per kb: 13.7307887161153	Normalized hits with 1T: 86.3%	Normalized hits with 10A: 35.6%	Normalized hits 25-32 nt: 99.3%	Normalized hits on the main strand(s): 84.5%	Predicted directionality: mono:minus
Cluster 466	Location: CM035929.1	Coordinates: 4578073-4583105	Size [bp]: 5033	Hits (absolute): 1553	Hits (normalized): 107.943270491411	Hits (normalized) per kb: 21.4467567210991	Normalized hits with 1T: 86.2%	Normalized hits with 10A: 35.7%	Normalized hits 25-32 nt: 99.4%	Normalized hits on the main strand(s): 84.8%	Predicted directionality: mono:plus
Cluster 467	Location: CM035929.1	Coordinates: 6907129-6912975	Size [bp]: 5847	Hits (absolute): 702	Hits (normalized): 126.071516822883	Hits (normalized) per kb: 21.5620768950866	Normalized hits with 1T: 97.2%	Normalized hits with 10A: 15.5%	Normalized hits 25-32 nt: 99.9%	Normalized hits on the main strand(s): 99.8%	Predicted directionality: mono:plus
Cluster 468	Location: CM035929.1	Coordinates: 8763419-8771000	Size [bp]: 7582	Hits (absolute): 2183	Hits (normalized): 155.443432250649	Hits (normalized) per kb: 20.5018302045467	Normalized hits with 1T: 95.9%	Normalized hits with 10A: 22.6%	Normalized hits 25-32 nt: 99.8%	Normalized hits on the main strand(s): 99.9%	Predicted directionality: mono:plus
Cluster 469	Location: CM035929.1	Coordinates: 9854043-9860318	Size [bp]: 6276	Hits (absolute): 651	Hits (normalized): 148.22153044413	Hits (normalized) per kb: 23.6168727225008	Normalized hits with 1T: 96.2%	Normalized hits with 10A: 13.1%	Normalized hits 25-32 nt: 98%	Normalized hits on the main strand(s): 99.8%	Predicted directionality: mono:minus
Cluster 470	Location: CM035929.1	Coordinates: 11380038-11393236	Size [bp]: 13199	Hits (absolute): 1184	Hits (normalized): 244.627929637026	Hits (normalized) per kb: 18.5336992351596	Normalized hits with 1T: 97.4%	Normalized hits with 10A: 2.5%	Normalized hits 25-32 nt: 97.5%	Normalized hits on the main strand(s): 99.6%	Predicted directionality: mono:minus
Cluster 471	Location: CM035929.1	Coordinates: 15858226-15864025	Size [bp]: 5800	Hits (absolute): 460	Hits (normalized): 92.6010067835669	Hits (normalized) per kb: 15.9659033610372	Normalized hits with 1T: 81.1%	Normalized hits with 10A: 83.4%	Normalized hits 25-32 nt: 99.4%	Normalized hits on the main strand(s): 87.2%	Predicted directionality: bi:plus-minus (split between 15861461 and 15861470)
Cluster 472	Location: CM035929.1	Coordinates: 18539003-18547672	Size [bp]: 8670	Hits (absolute): 1128	Hits (normalized): 253.186474879742	Hits (normalized) per kb: 29.2025626043696	Normalized hits with 1T: 91.7%	Normalized hits with 10A: 77.8%	Normalized hits 25-32 nt: 99.3%	Normalized hits on the main strand(s): 80%	Predicted directionality: mono:plus
Cluster 473	Location: CM035929.1	Coordinates: 25002742-25007807	Size [bp]: 5066	Hits (absolute): 1023	Hits (normalized): 166.93991713624	Hits (normalized) per kb: 32.9529144444731	Normalized hits with 1T: 79.3%	Normalized hits with 10A: 25.9%	Normalized hits 25-32 nt: 99.3%	Normalized hits on the main strand(s): 86.4%	Predicted directionality: bi:minus-plus (split between 25004526 and 25004526)
Cluster 474	Location: CM035929.1	Coordinates: 26450038-26455973	Size [bp]: 5936	Hits (absolute): 977	Hits (normalized): 160.58430228314	Hits (normalized) per kb: 27.0527149971838	Normalized hits with 1T: 79.5%	Normalized hits with 10A: 23.7%	Normalized hits 25-32 nt: 99.6%	Normalized hits on the main strand(s): 88.7%	Predicted directionality: bi:minus-plus (split between 26454036 and 26454039)
Cluster 475	Location: CM035929.1	Coordinates: 37804010-37806522	Size [bp]: 2513	Hits (absolute): 381	Hits (normalized): 138.065129307268	Hits (normalized) per kb: 54.9406276180956	Normalized hits with 1T: 97.8%	Normalized hits with 10A: 57.1%	Normalized hits 25-32 nt: 99.7%	Normalized hits on the main strand(s): 98.8%	Predicted directionality: mono:minus
Cluster 476	Location: CM035929.1	Coordinates: 49543264-49547960	Size [bp]: 4697	Hits (absolute): 406	Hits (normalized): 90.8941348989801	Hits (normalized) per kb: 19.351424105253	Normalized hits with 1T: 96.3%	Normalized hits with 10A: 18%	Normalized hits 25-32 nt: 99.9%	Normalized hits on the main strand(s): 99.6%	Predicted directionality: mono:plus
Cluster 477	Location: CM035929.1	Coordinates: 57704286-57713024	Size [bp]: 8739	Hits (absolute): 1290	Hits (normalized): 179.38880055998	Hits (normalized) per kb: 20.5276898799258	Normalized hits with 1T: 94.2%	Normalized hits with 10A: 18.2%	Normalized hits 25-32 nt: 99.7%	Normalized hits on the main strand(s): 94.1%	Predicted directionality: mono:plus
Cluster 478	Location: CM035929.1	Coordinates: 60392455-60394571	Size [bp]: 2117	Hits (absolute): 193	Hits (normalized): 527.80633157416	Hits (normalized) per kb: 249.318022700168	Normalized hits with 1T: 99.9%	Normalized hits with 10A: 0.1%	Normalized hits 25-32 nt: 99.9%	Normalized hits on the main strand(s): 100%	Predicted directionality: mono:minus
Cluster 479	Location: CM035929.1	Coordinates: 62730699-62766249	Size [bp]: 35551	Hits (absolute): 11940	Hits (normalized): 5078.29056019728	Hits (normalized) per kb: 142.84535224299	Normalized hits with 1T: 95.5%	Normalized hits with 10A: 29.4%	Normalized hits 25-32 nt: 99.8%	Normalized hits on the main strand(s): 100%	Predicted directionality: mono:minus
Cluster 480	Location: CM035929.1	Coordinates: 63703030-63734760	Size [bp]: 31731	Hits (absolute): 11755	Hits (normalized): 5967.5530403023	Hits (normalized) per kb: 188.066935379446	Normalized hits with 1T: 95.4%	Normalized hits with 10A: 15.8%	Normalized hits 25-32 nt: 99.8%	Normalized hits on the main strand(s): 100%	Predicted directionality: mono:plus
Cluster 481	Location: CM035929.1	Coordinates: 64476869-64482896	Size [bp]: 6028	Hits (absolute): 942	Hits (normalized): 115.328835343643	Hits (normalized) per kb: 19.131966319604	Normalized hits with 1T: 95.1%	Normalized hits with 10A: 14.9%	Normalized hits 25-32 nt: 99.7%	Normalized hits on the main strand(s): 95%	Predicted directionality: mono:plus
Cluster 482	Location: CM035929.1	Coordinates: 65498846-65503770	Size [bp]: 4925	Hits (absolute): 485	Hits (normalized): 1790.82303039389	Hits (normalized) per kb: 363.619185695736	Normalized hits with 1T: 99.8%	Normalized hits with 10A: 0.5%	Normalized hits 25-32 nt: 100%	Normalized hits on the main strand(s): 100%	Predicted directionality: mono:plus
Cluster 483	Location: CM035930.1	Coordinates: 5744109-5750752	Size [bp]: 6644	Hits (absolute): 1215	Hits (normalized): 125.149397874255	Hits (normalized) per kb: 18.8363273281087	Normalized hits with 1T: 95.6%	Normalized hits with 10A: 48.6%	Normalized hits 25-32 nt: 98.5%	Normalized hits on the main strand(s): 99.8%	Predicted directionality: mono:minus
Cluster 484	Location: CM035930.1	Coordinates: 8776201-8781020	Size [bp]: 4820	Hits (absolute): 472	Hits (normalized): 95.4991766001865	Hits (normalized) per kb: 19.8134037113485	Normalized hits with 1T: 99%	Normalized hits with 10A: 3.5%	Normalized hits 25-32 nt: 99.9%	Normalized hits on the main strand(s): 99.7%	Predicted directionality: mono:minus
Cluster 485	Location: CM035930.1	Coordinates: 24730504-24734201	Size [bp]: 3698	Hits (absolute): 143	Hits (normalized): 108.747120242626	Hits (normalized) per kb: 29.4073432769656	Normalized hits with 1T: 99.8%	Normalized hits with 10A: 41.9%	Normalized hits 25-32 nt: 99.9%	Normalized hits on the main strand(s): 99.9%	Predicted directionality: mono:minus
Cluster 486	Location: CM035930.1	Coordinates: 27709067-27717618	Size [bp]: 8552	Hits (absolute): 1367	Hits (normalized): 126.744209768822	Hits (normalized) per kb: 14.8203896327611	Normalized hits with 1T: 90.8%	Normalized hits with 10A: 18.2%	Normalized hits 25-32 nt: 99.6%	Normalized hits on the main strand(s): 93.2%	Predicted directionality: mono:plus
Cluster 487	Location: CM035930.1	Coordinates: 27805041-27813951	Size [bp]: 8911	Hits (absolute): 1596	Hits (normalized): 111.480221925024	Hits (normalized) per kb: 12.5104916022836	Normalized hits with 1T: 85.9%	Normalized hits with 10A: 34.4%	Normalized hits 25-32 nt: 99.3%	Normalized hits on the main strand(s): 85.4%	Predicted directionality: mono:plus
Cluster 488	Location: CM035930.1	Coordinates: 27818001-27826970	Size [bp]: 8970	Hits (absolute): 1737	Hits (normalized): 122.67710178326	Hits (normalized) per kb: 13.6762737247757	Normalized hits with 1T: 86.8%	Normalized hits with 10A: 39.3%	Normalized hits 25-32 nt: 99.4%	Normalized hits on the main strand(s): 86.5%	Predicted directionality: mono:plus
Cluster 489	Location: CM035930.1	Coordinates: 31054810-31059030	Size [bp]: 4221	Hits (absolute): 1514	Hits (normalized): 104.837909997174	Hits (normalized) per kb: 24.8371698363325	Normalized hits with 1T: 86.3%	Normalized hits with 10A: 34.6%	Normalized hits 25-32 nt: 99.3%	Normalized hits on the main strand(s): 84.6%	Predicted directionality: mono:minus
Cluster 490	Location: CM035930.1	Coordinates: 33605097-33609954	Size [bp]: 4858	Hits (absolute): 266	Hits (normalized): 103.56806811986	Hits (normalized) per kb: 21.3188561644948	Normalized hits with 1T: 1.4%	Normalized hits with 10A: 97.8%	Normalized hits 25-32 nt: 99%	Normalized hits on the main strand(s): 99%	Predicted directionality: mono:minus
Cluster 491	Location: CM035930.1	Coordinates: 40524012-40535914	Size [bp]: 11903	Hits (absolute): 2810	Hits (normalized): 325.292433147415	Hits (normalized) per kb: 27.3287845046085	Normalized hits with 1T: 97.4%	Normalized hits with 10A: 15%	Normalized hits 25-32 nt: 99.6%	Normalized hits on the main strand(s): 100%	Predicted directionality: mono:minus
Cluster 492	Location: CM035930.1	Coordinates: 40564036-40579556	Size [bp]: 15521	Hits (absolute): 6051	Hits (normalized): 1090.05122882597	Hits (normalized) per kb: 70.2306848685513	Normalized hits with 1T: 98.4%	Normalized hits with 10A: 7%	Normalized hits 25-32 nt: 99.1%	Normalized hits on the main strand(s): 100%	Predicted directionality: mono:minus
Cluster 493	Location: CM035930.1	Coordinates: 47258129-47270560	Size [bp]: 12432	Hits (absolute): 2480	Hits (normalized): 528.454867508776	Hits (normalized) per kb: 42.507715041949	Normalized hits with 1T: 96.4%	Normalized hits with 10A: 37.4%	Normalized hits 25-32 nt: 99.8%	Normalized hits on the main strand(s): 100%	Predicted directionality: mono:minus
Cluster 494	Location: CM035930.1	Coordinates: 47272002-47277965	Size [bp]: 5964	Hits (absolute): 988	Hits (normalized): 107.515581410459	Hits (normalized) per kb: 18.0276882899052	Normalized hits with 1T: 89.7%	Normalized hits with 10A: 17.3%	Normalized hits 25-32 nt: 99.4%	Normalized hits on the main strand(s): 99.7%	Predicted directionality: mono:minus
Cluster 495	Location: CM035930.1	Coordinates: 47280047-47285015	Size [bp]: 4969	Hits (absolute): 1183	Hits (normalized): 101.378990556175	Hits (normalized) per kb: 20.4025849639029	Normalized hits with 1T: 95.6%	Normalized hits with 10A: 23.7%	Normalized hits 25-32 nt: 99.4%	Normalized hits on the main strand(s): 98.6%	Predicted directionality: mono:minus
Cluster 496	Location: CM035930.1	Coordinates: 47316298-47332746	Size [bp]: 16449	Hits (absolute): 4839	Hits (normalized): 769.561689832678	Hits (normalized) per kb: 46.7850451316683	Normalized hits with 1T: 96.7%	Normalized hits with 10A: 25.6%	Normalized hits 25-32 nt: 99.6%	Normalized hits on the main strand(s): 99.9%	Predicted directionality: mono:minus
Cluster 497	Location: CM035930.1	Coordinates: 47334020-47340463	Size [bp]: 6444	Hits (absolute): 2014	Hits (normalized): 147.885892211463	Hits (normalized) per kb: 22.9494135336639	Normalized hits with 1T: 96.7%	Normalized hits with 10A: 30.2%	Normalized hits 25-32 nt: 99.7%	Normalized hits on the main strand(s): 98.6%	Predicted directionality: mono:minus
Cluster 498	Location: CM035930.1	Coordinates: 47343918-47371948	Size [bp]: 28031	Hits (absolute): 4825	Hits (normalized): 697.542044763474	Hits (normalized) per kb: 24.8846957262183	Normalized hits with 1T: 79.1%	Normalized hits with 10A: 23.5%	Normalized hits 25-32 nt: 99.6%	Normalized hits on the main strand(s): 99.8%	Predicted directionality: mono:minus
Cluster 499	Location: CM035930.1	Coordinates: 47377058-47385873	Size [bp]: 8816	Hits (absolute): 1399	Hits (normalized): 145.282625864847	Hits (normalized) per kb: 16.4796023178908	Normalized hits with 1T: 93.5%	Normalized hits with 10A: 14.9%	Normalized hits 25-32 nt: 99.8%	Normalized hits on the main strand(s): 100%	Predicted directionality: mono:minus
Cluster 500	Location: CM035930.1	Coordinates: 48329230-48334592	Size [bp]: 5363	Hits (absolute): 1878	Hits (normalized): 141.743054458703	Hits (normalized) per kb: 26.4299860576512	Normalized hits with 1T: 96.9%	Normalized hits with 10A: 31%	Normalized hits 25-32 nt: 99.7%	Normalized hits on the main strand(s): 98.5%	Predicted directionality: mono:minus
Cluster 501	Location: JAIWYP010000017.1	Coordinates: 38261-46820	Size [bp]: 8560	Hits (absolute): 1424	Hits (normalized): 153.688654882227	Hits (normalized) per kb: 17.9543027246404	Normalized hits with 1T: 83.6%	Normalized hits with 10A: 19.6%	Normalized hits 25-32 nt: 99.5%	Normalized hits on the main strand(s): 82.9%	Predicted directionality: mono:plus
Cluster 502	Location: JAIWYP010000017.1	Coordinates: 52061-59514	Size [bp]: 7454	Hits (absolute): 743	Hits (normalized): 98.3995782744637	Hits (normalized) per kb: 13.201014825918	Normalized hits with 1T: 97.4%	Normalized hits with 10A: 35.8%	Normalized hits 25-32 nt: 99.8%	Normalized hits on the main strand(s): 99.6%	Predicted directionality: mono:plus
Cluster 503	Location: JAIWYP010000017.1	Coordinates: 88948-96971	Size [bp]: 8024	Hits (absolute): 1515	Hits (normalized): 187.731630643842	Hits (normalized) per kb: 23.3960171165611	Normalized hits with 1T: 95.9%	Normalized hits with 10A: 18.1%	Normalized hits 25-32 nt: 99.9%	Normalized hits on the main strand(s): 100%	Predicted directionality: mono:plus
Cluster 504	Location: JAIWYP010000017.1	Coordinates: 109010-122938	Size [bp]: 13929	Hits (absolute): 2122	Hits (normalized): 220.74007467813	Hits (normalized) per kb: 15.8477875464682	Normalized hits with 1T: 97.5%	Normalized hits with 10A: 8.1%	Normalized hits 25-32 nt: 93.6%	Normalized hits on the main strand(s): 100%	Predicted directionality: mono:plus
Cluster 505	Location: JAIWYP010000017.1	Coordinates: 191189-200005	Size [bp]: 8817	Hits (absolute): 1422	Hits (normalized): 411.855367706297	Hits (normalized) per kb: 46.7116595664036	Normalized hits with 1T: 98.1%	Normalized hits with 10A: 7.9%	Normalized hits 25-32 nt: 99.7%	Normalized hits on the main strand(s): 97.7%	Predicted directionality: mono:plus
Cluster 506	Location: JAIWYP010000017.1	Coordinates: 1226976-1230062	Size [bp]: 3087	Hits (absolute): 523	Hits (normalized): 128.892419602196	Hits (normalized) per kb: 41.7535909950851	Normalized hits with 1T: 98.6%	Normalized hits with 10A: 85.1%	Normalized hits 25-32 nt: 99.7%	Normalized hits on the main strand(s): 97.9%	Predicted directionality: mono:minus
Cluster 507	Location: JAIWYP010000017.1	Coordinates: 1559017-1562615	Size [bp]: 3599	Hits (absolute): 437	Hits (normalized): 124.660279489805	Hits (normalized) per kb: 34.6372878948364	Normalized hits with 1T: 98.7%	Normalized hits with 10A: 87.2%	Normalized hits 25-32 nt: 99.9%	Normalized hits on the main strand(s): 97.9%	Predicted directionality: mono:minus
Cluster 508	Location: JAIWYP010000026.1	Coordinates: 273309-281604	Size [bp]: 8296	Hits (absolute): 2043	Hits (normalized): 159.752803482389	Hits (normalized) per kb: 19.2563723254814	Normalized hits with 1T: 92.6%	Normalized hits with 10A: 28.6%	Normalized hits 25-32 nt: 99.8%	Normalized hits on the main strand(s): 92.4%	Predicted directionality: mono:plus
Cluster 509	Location: JAIWYP010000028.1	Coordinates: 17440-23839	Size [bp]: 6400	Hits (absolute): 727	Hits (normalized): 95.7477369459152	Hits (normalized) per kb: 14.9608705719823	Normalized hits with 1T: 97%	Normalized hits with 10A: 20.6%	Normalized hits 25-32 nt: 99.7%	Normalized hits on the main strand(s): 99.9%	Predicted directionality: mono:plus
Cluster 510	Location: JAIWYP010000040.1	Coordinates: 5722-10892	Size [bp]: 5171	Hits (absolute): 708	Hits (normalized): 150.80712656053	Hits (normalized) per kb: 29.1641225463737	Normalized hits with 1T: 96.4%	Normalized hits with 10A: 13%	Normalized hits 25-32 nt: 98%	Normalized hits on the main strand(s): 99.9%	Predicted directionality: mono:plus
Cluster 511	Location: JAIWYP010000043.1	Coordinates: 296737-301897	Size [bp]: 5161	Hits (absolute): 1436	Hits (normalized): 104.390756160568	Hits (normalized) per kb: 20.2271585174128	Normalized hits with 1T: 86.1%	Normalized hits with 10A: 35.4%	Normalized hits 25-32 nt: 99.5%	Normalized hits on the main strand(s): 84.7%	Predicted directionality: mono:minus
Cluster 512	Location: JAIWYP010000048.1	Coordinates: 145015-150632	Size [bp]: 5618	Hits (absolute): 571	Hits (normalized): 114.843314470396	Hits (normalized) per kb: 20.4417239320442	Normalized hits with 1T: 96.9%	Normalized hits with 10A: 16.5%	Normalized hits 25-32 nt: 99.9%	Normalized hits on the main strand(s): 99.7%	Predicted directionality: mono:minus
Cluster 513	Location: JAIWYP010000074.1	Coordinates: 27764-29724	Size [bp]: 1961	Hits (absolute): 272	Hits (normalized): 301.63123938875	Hits (normalized) per kb: 153.814746974713	Normalized hits with 1T: 0.3%	Normalized hits with 10A: 99.8%	Normalized hits 25-32 nt: 99.9%	Normalized hits on the main strand(s): 100%	Predicted directionality: mono:plus
Cluster 514	Location: JAIWYP010000089.1	Coordinates: 81097-85805	Size [bp]: 4709	Hits (absolute): 469	Hits (normalized): 120.377322239782	Hits (normalized) per kb: 25.5633374773813	Normalized hits with 1T: 99.2%	Normalized hits with 10A: 8%	Normalized hits 25-32 nt: 99.9%	Normalized hits on the main strand(s): 99.8%	Predicted directionality: bi:plus-minus (split between 83196 and 83201)
Cluster 515	Location: JAIWYP010000103.1	Coordinates: 105000-111991	Size [bp]: 6992	Hits (absolute): 889	Hits (normalized): 99.3297980062835	Hits (normalized) per kb: 14.206047614973	Normalized hits with 1T: 92.2%	Normalized hits with 10A: 6.4%	Normalized hits 25-32 nt: 98%	Normalized hits on the main strand(s): 99.9%	Predicted directionality: mono:plus
Cluster 516	Location: JAIWYP010000147.1	Coordinates: 39162-45598	Size [bp]: 6437	Hits (absolute): 1020	Hits (normalized): 166.605980494503	Hits (normalized) per kb: 25.8827394138195	Normalized hits with 1T: 79.3%	Normalized hits with 10A: 25.9%	Normalized hits 25-32 nt: 99.1%	Normalized hits on the main strand(s): 86.2%	Predicted directionality: bi:minus-plus (split between 43390 and 43390)

Total size of 516 predicted piRNA clusters: 5232895 bp (0.291%)
Non identical sequences that can be assigned to clusters: 747952 (25.761%)
Sequence reads that can be assigned to clusters: 48968340 (34.224%)


                                              /\
                _______________________/\___ /  \_______
               I                      /  \  /    \      I
               I     pro             /    \/      \     I
               I        TRAC        /               \   I
               I   ________________/_________________\_ I
               I   \              /                     I
               I    \            /                      I
               I     \  /\      /       V.2.4.2         I
               I      \/  \    /                        I
               I___________\  /_________________________I
                            \/


================================= proTRAC ====================================
VERSION: .......... 2.4.2
LAST MODIFIED: .... 11. May 2018

Please cite:
Rosenkranz D, Zischler H. proTRAC - a software for probabilistic piRNA cluster
detection, visualization and analysis. 2012. BMC Bioinformatics 13:5.


Contact:
David Rosenkranz
Institute of Organismic and Molecular Evolutionary Biology
Dept. Anthropology, small RNA group
Johannes Gutenberg University Mainz


You can find the latest proTRAC version at:
http://sourceforge.net/projects/protrac/files
http://www.smallRNAgroup-mainz.de/software
==============================================================================

PARAMETERS:
Map file: .............../Volumes/My_Book/SEQUENCING_DATA/sRNA/sRNA_DETECTION/original_25.sam
Genome file: ............/Volumes/My_Book/SEQUENCING_DATA/sRNA/miRDeep2/Bowtiew_genome_index/GCA_020536995.1_UMN_Dpol_1.0_genomic.fasta
RepeatMasker annotation: n.a.
GeneSet:................./Volumes/My_Book/SEQUENCING_DATA/sRNA/miRDeep2/Bowtiew_genome_index/genomic.gff

Significant (p<=0.01) hit density will be calculated based
on observed hit distribution.

Sliding window size: ........................................ 5000 bp
Sliding window increament: .................................. 1000 bp
Normalize each hit by number of genomic hits: ............... yes
Normalize each hit by number of sequence reads: ............. yes
Normalize values (-> per million mapped reads): ............. yes
Min. fraction of hits with 1T(U) or 10A: .................... 0.75
Alternatively: Min. fraction of hits with 1T(U) and 10A: .... 0.5
Min. fraction of hits with typical piRNA length: ............ 0.75
Typical piRNA length: ....................................... 25-32 nt
Min. size of a piRNA cluster: ............................... 1000 bp.
Min. number of hits (absolute): ............................. 0
Min. number of hits (normalized): ........................... 0
Min. fraction of hits on the mainstrand: .................... 0.75
Top fraction of mapped sequences (in terms of read counts): . 1%
Top fraction accounts for max. n% of sequence reads: ........ 90%
Min. fraction of hits on each arm of a bidirectional cluster: 0.05
Output html file for each cluster: .......................... no
Output a summary table: ..................................... yes
Output a FASTA file for each cluster (piRNA sequences): ..... yes
Output a FASTA file comprising cluster sequences: ........... yes
Output a GTF file for predicted piRNA clusters: ..............yes
Search DNA motifs in clusters: .............................. yes
Output flanking sequences: +/- .............................. 0 bp
Output ~.pTi file: .......................................... no
==============================================================================


Cluster 1	Location: CM035915.1	Coordinates: 19049135-19057792	Size [bp]: 8658	Hits (absolute): 1130	Hits (normalized): 163.344376979855	Hits (normalized) per kb: 18.8661240619997	Normalized hits with 1T: 96.8%	Normalized hits with 10A: 73.1%	Normalized hits 25-32 nt: 99.6%	Normalized hits on the main strand(s): 99.5%	Predicted directionality: mono:minus
Cluster 2	Location: CM035915.1	Coordinates: 19844270-19853024	Size [bp]: 8755	Hits (absolute): 968	Hits (normalized): 156.013171408756	Hits (normalized) per kb: 17.819742715228	Normalized hits with 1T: 96.9%	Normalized hits with 10A: 74.9%	Normalized hits 25-32 nt: 99.5%	Normalized hits on the main strand(s): 99.5%	Predicted directionality: mono:plus
Cluster 3	Location: CM035915.1	Coordinates: 19871056-19879925	Size [bp]: 8870	Hits (absolute): 1484	Hits (normalized): 192.022958762704	Hits (normalized) per kb: 21.6482480516083	Normalized hits with 1T: 92.1%	Normalized hits with 10A: 21.7%	Normalized hits 25-32 nt: 98.3%	Normalized hits on the main strand(s): 99.7%	Predicted directionality: mono:plus
Cluster 4	Location: CM035915.1	Coordinates: 19924058-19931809	Size [bp]: 7752	Hits (absolute): 356	Hits (normalized): 144.285182266933	Hits (normalized) per kb: 18.6127549937563	Normalized hits with 1T: 97.1%	Normalized hits with 10A: 2.9%	Normalized hits 25-32 nt: 99.8%	Normalized hits on the main strand(s): 99.4%	Predicted directionality: mono:minus
Cluster 5	Location: CM035915.1	Coordinates: 19991034-19998987	Size [bp]: 7954	Hits (absolute): 396	Hits (normalized): 145.141081176152	Hits (normalized) per kb: 18.2475086746001	Normalized hits with 1T: 96.9%	Normalized hits with 10A: 2.9%	Normalized hits 25-32 nt: 99.8%	Normalized hits on the main strand(s): 99.3%	Predicted directionality: mono:minus
Cluster 6	Location: CM035915.1	Coordinates: 22387083-22391922	Size [bp]: 4840	Hits (absolute): 2154	Hits (normalized): 124.425017083996	Hits (normalized) per kb: 25.7079115314186	Normalized hits with 1T: 94.3%	Normalized hits with 10A: 43.1%	Normalized hits 25-32 nt: 98.4%	Normalized hits on the main strand(s): 99.6%	Predicted directionality: mono:minus
Cluster 7	Location: CM035915.1	Coordinates: 25447003-25457972	Size [bp]: 10970	Hits (absolute): 1534	Hits (normalized): 492.971755825058	Hits (normalized) per kb: 44.9384592857296	Normalized hits with 1T: 98.9%	Normalized hits with 10A: 8.5%	Normalized hits 25-32 nt: 99.9%	Normalized hits on the main strand(s): 98.4%	Predicted directionality: mono:minus
Cluster 8	Location: CM035915.1	Coordinates: 39073593-39081487	Size [bp]: 7895	Hits (absolute): 1978	Hits (normalized): 599.322363623291	Hits (normalized) per kb: 75.9120052516498	Normalized hits with 1T: 15.7%	Normalized hits with 10A: 85.4%	Normalized hits 25-32 nt: 99%	Normalized hits on the main strand(s): 91.7%	Predicted directionality: bi:plus-minus (split between 39078499 and 39078499)
Cluster 9	Location: CM035915.1	Coordinates: 50015780-50026020	Size [bp]: 10241	Hits (absolute): 1490	Hits (normalized): 738.569472806289	Hits (normalized) per kb: 72.1188728696021	Normalized hits with 1T: 98.9%	Normalized hits with 10A: 42.2%	Normalized hits 25-32 nt: 99.4%	Normalized hits on the main strand(s): 100%	Predicted directionality: mono:minus
Cluster 10	Location: CM035915.1	Coordinates: 50036103-50073603	Size [bp]: 37501	Hits (absolute): 11980	Hits (normalized): 5826.36366080154	Hits (normalized) per kb: 155.365419070794	Normalized hits with 1T: 87.9%	Normalized hits with 10A: 27%	Normalized hits 25-32 nt: 99.6%	Normalized hits on the main strand(s): 100%	Predicted directionality: mono:minus
Cluster 11	Location: CM035915.1	Coordinates: 57679142-57686827	Size [bp]: 7686	Hits (absolute): 1820	Hits (normalized): 340.296869575804	Hits (normalized) per kb: 44.2745994218579	Normalized hits with 1T: 89.9%	Normalized hits with 10A: 17.5%	Normalized hits 25-32 nt: 98.6%	Normalized hits on the main strand(s): 93.2%	Predicted directionality: mono:plus
Cluster 12	Location: CM035915.1	Coordinates: 62087199-62093400	Size [bp]: 6202	Hits (absolute): 705	Hits (normalized): 256.700364204325	Hits (normalized) per kb: 41.389647076631	Normalized hits with 1T: 95.6%	Normalized hits with 10A: 10.9%	Normalized hits 25-32 nt: 99.6%	Normalized hits on the main strand(s): 98.3%	Predicted directionality: mono:minus
Cluster 13	Location: CM035915.1	Coordinates: 62132571-62137248	Size [bp]: 4678	Hits (absolute): 1346	Hits (normalized): 133.493251087504	Hits (normalized) per kb: 28.5361026243442	Normalized hits with 1T: 87.2%	Normalized hits with 10A: 20.2%	Normalized hits 25-32 nt: 99.4%	Normalized hits on the main strand(s): 93.9%	Predicted directionality: mono:plus
Cluster 14	Location: CM035915.1	Coordinates: 62784151-62790958	Size [bp]: 6808	Hits (absolute): 610	Hits (normalized): 188.716069851458	Hits (normalized) per kb: 27.7200567941574	Normalized hits with 1T: 99%	Normalized hits with 10A: 4.5%	Normalized hits 25-32 nt: 99.9%	Normalized hits on the main strand(s): 99.3%	Predicted directionality: mono:plus
Cluster 15	Location: CM035915.1	Coordinates: 79762457-79772017	Size [bp]: 9561	Hits (absolute): 2582	Hits (normalized): 303.770652114809	Hits (normalized) per kb: 31.7714940055182	Normalized hits with 1T: 77.7%	Normalized hits with 10A: 28.5%	Normalized hits 25-32 nt: 99.5%	Normalized hits on the main strand(s): 100%	Predicted directionality: mono:minus
Cluster 16	Location: CM035915.1	Coordinates: 79773001-79777994	Size [bp]: 4994	Hits (absolute): 1294	Hits (normalized): 123.384281392148	Hits (normalized) per kb: 24.7067746611189	Normalized hits with 1T: 90%	Normalized hits with 10A: 37.4%	Normalized hits 25-32 nt: 99.2%	Normalized hits on the main strand(s): 99.9%	Predicted directionality: mono:minus
Cluster 17	Location: CM035915.1	Coordinates: 82992351-82999028	Size [bp]: 6678	Hits (absolute): 1542	Hits (normalized): 135.382453871418	Hits (normalized) per kb: 20.272815966858	Normalized hits with 1T: 91.9%	Normalized hits with 10A: 39.9%	Normalized hits 25-32 nt: 98.6%	Normalized hits on the main strand(s): 99.2%	Predicted directionality: mono:plus
Cluster 18	Location: CM035915.1	Coordinates: 84299856-84318001	Size [bp]: 18146	Hits (absolute): 5143	Hits (normalized): 697.989234844417	Hits (normalized) per kb: 38.4652086428467	Normalized hits with 1T: 92.9%	Normalized hits with 10A: 23.2%	Normalized hits 25-32 nt: 99.5%	Normalized hits on the main strand(s): 99.5%	Predicted directionality: mono:plus
Cluster 19	Location: CM035915.1	Coordinates: 84340070-84346856	Size [bp]: 6787	Hits (absolute): 2696	Hits (normalized): 1714.75752888399	Hits (normalized) per kb: 252.653382888374	Normalized hits with 1T: 97.6%	Normalized hits with 10A: 2.6%	Normalized hits 25-32 nt: 99.6%	Normalized hits on the main strand(s): 100%	Predicted directionality: mono:plus
Cluster 20	Location: CM035915.1	Coordinates: 89519286-89525067	Size [bp]: 5782	Hits (absolute): 194	Hits (normalized): 124.473882275435	Hits (normalized) per kb: 21.5281445322462	Normalized hits with 1T: 0.8%	Normalized hits with 10A: 99.5%	Normalized hits 25-32 nt: 99.7%	Normalized hits on the main strand(s): 99.9%	Predicted directionality: mono:plus
Cluster 21	Location: CM035915.1	Coordinates: 91509738-91513056	Size [bp]: 3319	Hits (absolute): 965	Hits (normalized): 721.168300129372	Hits (normalized) per kb: 217.284541689906	Normalized hits with 1T: 90.9%	Normalized hits with 10A: 25.8%	Normalized hits 25-32 nt: 99.8%	Normalized hits on the main strand(s): 91.3%	Predicted directionality: mono:plus
Cluster 22	Location: CM035915.1	Coordinates: 105213122-105217406	Size [bp]: 4285	Hits (absolute): 650	Hits (normalized): 154.865839788575	Hits (normalized) per kb: 36.1412878058732	Normalized hits with 1T: 98%	Normalized hits with 10A: 7.3%	Normalized hits 25-32 nt: 99.4%	Normalized hits on the main strand(s): 96.7%	Predicted directionality: mono:plus
Cluster 23	Location: CM035915.1	Coordinates: 108739309-108743933	Size [bp]: 4625	Hits (absolute): 589	Hits (normalized): 123.446260393598	Hits (normalized) per kb: 26.6909506111295	Normalized hits with 1T: 94.6%	Normalized hits with 10A: 33.3%	Normalized hits 25-32 nt: 99.8%	Normalized hits on the main strand(s): 99.5%	Predicted directionality: mono:plus
Cluster 24	Location: CM035915.1	Coordinates: 116999018-117005126	Size [bp]: 6109	Hits (absolute): 864	Hits (normalized): 154.070504003512	Hits (normalized) per kb: 25.2200938123654	Normalized hits with 1T: 88%	Normalized hits with 10A: 10.3%	Normalized hits 25-32 nt: 99.7%	Normalized hits on the main strand(s): 99.8%	Predicted directionality: mono:minus
Cluster 25	Location: CM035915.1	Coordinates: 117041001-117049397	Size [bp]: 8397	Hits (absolute): 546	Hits (normalized): 491.897436190542	Hits (normalized) per kb: 58.580080255475	Normalized hits with 1T: 99.5%	Normalized hits with 10A: 62.9%	Normalized hits 25-32 nt: 100%	Normalized hits on the main strand(s): 99.9%	Predicted directionality: mono:minus
Cluster 26	Location: CM035915.1	Coordinates: 119301075-119306843	Size [bp]: 5769	Hits (absolute): 788	Hits (normalized): 137.711687191688	Hits (normalized) per kb: 23.8709857866534	Normalized hits with 1T: 94.4%	Normalized hits with 10A: 52.1%	Normalized hits 25-32 nt: 99.7%	Normalized hits on the main strand(s): 99.5%	Predicted directionality: mono:plus
Cluster 27	Location: CM035915.1	Coordinates: 119312021-119321029	Size [bp]: 9009	Hits (absolute): 2521	Hits (normalized): 232.582148859351	Hits (normalized) per kb: 25.8164982749515	Normalized hits with 1T: 87.2%	Normalized hits with 10A: 34.9%	Normalized hits 25-32 nt: 99.7%	Normalized hits on the main strand(s): 99.9%	Predicted directionality: mono:plus
Cluster 28	Location: CM035915.1	Coordinates: 119340003-119355321	Size [bp]: 15319	Hits (absolute): 3308	Hits (normalized): 438.661791268893	Hits (normalized) per kb: 28.6348178457377	Normalized hits with 1T: 96.4%	Normalized hits with 10A: 28.5%	Normalized hits 25-32 nt: 98.4%	Normalized hits on the main strand(s): 99.7%	Predicted directionality: mono:plus
Cluster 29	Location: CM035915.1	Coordinates: 129808500-129813729	Size [bp]: 5230	Hits (absolute): 1427	Hits (normalized): 172.645684284263	Hits (normalized) per kb: 33.0103700340074	Normalized hits with 1T: 96.2%	Normalized hits with 10A: 52.2%	Normalized hits 25-32 nt: 99.7%	Normalized hits on the main strand(s): 97.3%	Predicted directionality: mono:minus
Cluster 30	Location: CM035915.1	Coordinates: 133352017-133361026	Size [bp]: 9010	Hits (absolute): 2035	Hits (normalized): 1521.94557934617	Hits (normalized) per kb: 168.91737371444	Normalized hits with 1T: 87.3%	Normalized hits with 10A: 11%	Normalized hits 25-32 nt: 99.8%	Normalized hits on the main strand(s): 99.9%	Predicted directionality: mono:minus
Cluster 31	Location: CM035915.1	Coordinates: 133362009-133399878	Size [bp]: 37870	Hits (absolute): 12813	Hits (normalized): 6272.47637032519	Hits (normalized) per kb: 165.631802095724	Normalized hits with 1T: 80.8%	Normalized hits with 10A: 16.9%	Normalized hits 25-32 nt: 99.6%	Normalized hits on the main strand(s): 99.9%	Predicted directionality: mono:minus
Cluster 32	Location: CM035915.1	Coordinates: 133401500-133422991	Size [bp]: 21492	Hits (absolute): 5844	Hits (normalized): 1142.18111374374	Hits (normalized) per kb: 53.1441620640695	Normalized hits with 1T: 85.4%	Normalized hits with 10A: 14.7%	Normalized hits 25-32 nt: 99.3%	Normalized hits on the main strand(s): 99.9%	Predicted directionality: mono:minus
Cluster 33	Location: CM035915.1	Coordinates: 133425171-133450651	Size [bp]: 25481	Hits (absolute): 5513	Hits (normalized): 1269.14568808328	Hits (normalized) per kb: 49.807587580967	Normalized hits with 1T: 92.5%	Normalized hits with 10A: 19.2%	Normalized hits 25-32 nt: 99.6%	Normalized hits on the main strand(s): 99.9%	Predicted directionality: mono:minus
Cluster 34	Location: CM035915.1	Coordinates: 133712082-133717898	Size [bp]: 5817	Hits (absolute): 996	Hits (normalized): 144.877866793717	Hits (normalized) per kb: 24.9058503575959	Normalized hits with 1T: 94.5%	Normalized hits with 10A: 18.3%	Normalized hits 25-32 nt: 99.7%	Normalized hits on the main strand(s): 100%	Predicted directionality: mono:plus
Cluster 35	Location: CM035915.1	Coordinates: 133755150-133768927	Size [bp]: 13778	Hits (absolute): 2325	Hits (normalized): 571.422561049859	Hits (normalized) per kb: 41.4735550148155	Normalized hits with 1T: 76.7%	Normalized hits with 10A: 16.5%	Normalized hits 25-32 nt: 99.5%	Normalized hits on the main strand(s): 100%	Predicted directionality: mono:plus
Cluster 36	Location: CM035915.1	Coordinates: 133779308-133785395	Size [bp]: 6088	Hits (absolute): 1133	Hits (normalized): 176.542606992728	Hits (normalized) per kb: 28.998418911204	Normalized hits with 1T: 90.8%	Normalized hits with 10A: 26%	Normalized hits 25-32 nt: 99.4%	Normalized hits on the main strand(s): 100%	Predicted directionality: mono:plus
Cluster 37	Location: CM035915.1	Coordinates: 133794042-133803025	Size [bp]: 8984	Hits (absolute): 1137	Hits (normalized): 365.003053011962	Hits (normalized) per kb: 40.6278946182107	Normalized hits with 1T: 98.4%	Normalized hits with 10A: 15%	Normalized hits 25-32 nt: 99.5%	Normalized hits on the main strand(s): 99.9%	Predicted directionality: mono:plus
Cluster 38	Location: CM035915.1	Coordinates: 133813011-133827990	Size [bp]: 14980	Hits (absolute): 2197	Hits (normalized): 363.510399121783	Hits (normalized) per kb: 24.2666692990726	Normalized hits with 1T: 91.1%	Normalized hits with 10A: 9.9%	Normalized hits 25-32 nt: 99.7%	Normalized hits on the main strand(s): 99.7%	Predicted directionality: mono:plus
Cluster 39	Location: CM035915.1	Coordinates: 133837027-133846986	Size [bp]: 9960	Hits (absolute): 1855	Hits (normalized): 1504.99081235194	Hits (normalized) per kb: 151.103389387127	Normalized hits with 1T: 87.4%	Normalized hits with 10A: 11%	Normalized hits 25-32 nt: 99.8%	Normalized hits on the main strand(s): 100%	Predicted directionality: mono:plus
Cluster 40	Location: CM035915.1	Coordinates: 139692044-139713018	Size [bp]: 20975	Hits (absolute): 10131	Hits (normalized): 3740.21361891542	Hits (normalized) per kb: 178.317530671642	Normalized hits with 1T: 88.8%	Normalized hits with 10A: 12.8%	Normalized hits 25-32 nt: 99.2%	Normalized hits on the main strand(s): 100%	Predicted directionality: mono:minus
Cluster 41	Location: CM035915.1	Coordinates: 139714401-139740959	Size [bp]: 26559	Hits (absolute): 9127	Hits (normalized): 2919.54127216922	Hits (normalized) per kb: 109.926802663337	Normalized hits with 1T: 87.4%	Normalized hits with 10A: 14.1%	Normalized hits 25-32 nt: 99.7%	Normalized hits on the main strand(s): 99.5%	Predicted directionality: mono:minus
Cluster 42	Location: CM035915.1	Coordinates: 140106166-140111022	Size [bp]: 4857	Hits (absolute): 1683	Hits (normalized): 146.913731279461	Hits (normalized) per kb: 30.2479890886775	Normalized hits with 1T: 95.4%	Normalized hits with 10A: 27%	Normalized hits 25-32 nt: 99.6%	Normalized hits on the main strand(s): 99.8%	Predicted directionality: mono:minus
Cluster 43	Location: CM035915.1	Coordinates: 140117627-140122024	Size [bp]: 4398	Hits (absolute): 1458	Hits (normalized): 121.290258155519	Hits (normalized) per kb: 27.5785649768266	Normalized hits with 1T: 96.4%	Normalized hits with 10A: 9.1%	Normalized hits 25-32 nt: 99.7%	Normalized hits on the main strand(s): 100%	Predicted directionality: mono:minus
Cluster 44	Location: CM035915.1	Coordinates: 140140455-140146851	Size [bp]: 6397	Hits (absolute): 1096	Hits (normalized): 149.840382971186	Hits (normalized) per kb: 23.4234767830026	Normalized hits with 1T: 95.9%	Normalized hits with 10A: 24.4%	Normalized hits 25-32 nt: 99.6%	Normalized hits on the main strand(s): 99.3%	Predicted directionality: mono:minus
Cluster 45	Location: CM035915.1	Coordinates: 140154867-140172024	Size [bp]: 17158	Hits (absolute): 6656	Hits (normalized): 1573.04618981742	Hits (normalized) per kb: 91.6801166155815	Normalized hits with 1T: 92.6%	Normalized hits with 10A: 41.5%	Normalized hits 25-32 nt: 99.4%	Normalized hits on the main strand(s): 100%	Predicted directionality: mono:minus
Cluster 46	Location: CM035915.1	Coordinates: 140178015-140182997	Size [bp]: 4983	Hits (absolute): 919	Hits (normalized): 125.513730464669	Hits (normalized) per kb: 25.1880113654125	Normalized hits with 1T: 90.8%	Normalized hits with 10A: 52%	Normalized hits 25-32 nt: 98.9%	Normalized hits on the main strand(s): 100%	Predicted directionality: mono:minus
Cluster 47	Location: CM035915.1	Coordinates: 179074838-179080981	Size [bp]: 6144	Hits (absolute): 1454	Hits (normalized): 125.774652027312	Hits (normalized) per kb: 20.4710690364901	Normalized hits with 1T: 87.7%	Normalized hits with 10A: 21.5%	Normalized hits 25-32 nt: 99.6%	Normalized hits on the main strand(s): 99.6%	Predicted directionality: mono:minus
Cluster 48	Location: CM035915.1	Coordinates: 194367022-194374403	Size [bp]: 7382	Hits (absolute): 1425	Hits (normalized): 157.599928833578	Hits (normalized) per kb: 21.3488118800478	Normalized hits with 1T: 92.9%	Normalized hits with 10A: 20.9%	Normalized hits 25-32 nt: 98.8%	Normalized hits on the main strand(s): 92%	Predicted directionality: bi:plus-minus (split between 194369814 and 194369847)
Cluster 49	Location: CM035915.1	Coordinates: 194386259-194391517	Size [bp]: 5259	Hits (absolute): 544	Hits (normalized): 156.286212719237	Hits (normalized) per kb: 29.7182174005321	Normalized hits with 1T: 96.8%	Normalized hits with 10A: 2.6%	Normalized hits 25-32 nt: 99.8%	Normalized hits on the main strand(s): 99.8%	Predicted directionality: mono:minus
Cluster 50	Location: CM035915.1	Coordinates: 199152030-199158939	Size [bp]: 6910	Hits (absolute): 219	Hits (normalized): 265.558442734638	Hits (normalized) per kb: 38.4306583153589	Normalized hits with 1T: 99.7%	Normalized hits with 10A: 99.6%	Normalized hits 25-32 nt: 99.7%	Normalized hits on the main strand(s): 100%	Predicted directionality: mono:minus
Cluster 51	Location: CM035915.1	Coordinates: 206177015-206183022	Size [bp]: 6008	Hits (absolute): 1347	Hits (normalized): 141.673234297418	Hits (normalized) per kb: 23.5805985103873	Normalized hits with 1T: 92.9%	Normalized hits with 10A: 52.9%	Normalized hits 25-32 nt: 97.8%	Normalized hits on the main strand(s): 99.6%	Predicted directionality: mono:plus
Cluster 52	Location: CM035916.1	Coordinates: 17650779-17655969	Size [bp]: 5191	Hits (absolute): 659	Hits (normalized): 246.332404698414	Hits (normalized) per kb: 47.4540521775755	Normalized hits with 1T: 98.9%	Normalized hits with 10A: 2%	Normalized hits 25-32 nt: 99.6%	Normalized hits on the main strand(s): 99.7%	Predicted directionality: mono:minus
Cluster 53	Location: CM035916.1	Coordinates: 22466938-22470961	Size [bp]: 4024	Hits (absolute): 1106	Hits (normalized): 585.981265454756	Hits (normalized) per kb: 145.621404092404	Normalized hits with 1T: 92.8%	Normalized hits with 10A: 7.4%	Normalized hits 25-32 nt: 99.7%	Normalized hits on the main strand(s): 92.1%	Predicted directionality: mono:minus
Cluster 54	Location: CM035916.1	Coordinates: 42401554-42408891	Size [bp]: 7338	Hits (absolute): 1298	Hits (normalized): 300.146831938435	Hits (normalized) per kb: 40.9034746112677	Normalized hits with 1T: 96.7%	Normalized hits with 10A: 5.6%	Normalized hits 25-32 nt: 99.8%	Normalized hits on the main strand(s): 96.9%	Predicted directionality: mono:plus
Cluster 55	Location: CM035916.1	Coordinates: 44320453-44328096	Size [bp]: 7644	Hits (absolute): 1760	Hits (normalized): 374.631405080133	Hits (normalized) per kb: 49.0096395413691	Normalized hits with 1T: 85.5%	Normalized hits with 10A: 14.7%	Normalized hits 25-32 nt: 99.7%	Normalized hits on the main strand(s): 97.7%	Predicted directionality: mono:minus
Cluster 56	Location: CM035916.1	Coordinates: 46081003-46089849	Size [bp]: 8847	Hits (absolute): 1280	Hits (normalized): 729.144385516162	Hits (normalized) per kb: 82.4173383414856	Normalized hits with 1T: 90.7%	Normalized hits with 10A: 26.3%	Normalized hits 25-32 nt: 99.8%	Normalized hits on the main strand(s): 90.9%	Predicted directionality: mono:plus
Cluster 57	Location: CM035916.1	Coordinates: 52183701-52188458	Size [bp]: 4758	Hits (absolute): 471	Hits (normalized): 149.227066969765	Hits (normalized) per kb: 31.3634710904248	Normalized hits with 1T: 95.2%	Normalized hits with 10A: 39.9%	Normalized hits 25-32 nt: 99.7%	Normalized hits on the main strand(s): 96.9%	Predicted directionality: mono:plus
Cluster 58	Location: CM035916.1	Coordinates: 53504516-53509029	Size [bp]: 4514	Hits (absolute): 366	Hits (normalized): 173.107381546587	Hits (normalized) per kb: 38.3492182577093	Normalized hits with 1T: 1.3%	Normalized hits with 10A: 98.3%	Normalized hits 25-32 nt: 99.5%	Normalized hits on the main strand(s): 99.7%	Predicted directionality: mono:plus
Cluster 59	Location: CM035916.1	Coordinates: 55577113-55583898	Size [bp]: 6786	Hits (absolute): 490	Hits (normalized): 180.417392421543	Hits (normalized) per kb: 26.5864770018213	Normalized hits with 1T: 1.6%	Normalized hits with 10A: 97.7%	Normalized hits 25-32 nt: 99.1%	Normalized hits on the main strand(s): 99.7%	Predicted directionality: mono:minus
Cluster 60	Location: CM035916.1	Coordinates: 65382120-65389945	Size [bp]: 7826	Hits (absolute): 242	Hits (normalized): 222.692446318278	Hits (normalized) per kb: 28.4554851935394	Normalized hits with 1T: 97%	Normalized hits with 10A: 0.7%	Normalized hits 25-32 nt: 99.4%	Normalized hits on the main strand(s): 99.3%	Predicted directionality: mono:minus
Cluster 61	Location: CM035916.1	Coordinates: 75198014-75207022	Size [bp]: 9009	Hits (absolute): 1229	Hits (normalized): 204.184525021686	Hits (normalized) per kb: 22.6641922051171	Normalized hits with 1T: 97.5%	Normalized hits with 10A: 14.4%	Normalized hits 25-32 nt: 99.6%	Normalized hits on the main strand(s): 99.9%	Predicted directionality: mono:plus
Cluster 62	Location: CM035916.1	Coordinates: 75219001-75226020	Size [bp]: 7020	Hits (absolute): 1644	Hits (normalized): 142.172173152558	Hits (normalized) per kb: 20.2522502957344	Normalized hits with 1T: 92.3%	Normalized hits with 10A: 24.2%	Normalized hits 25-32 nt: 99.2%	Normalized hits on the main strand(s): 99.4%	Predicted directionality: mono:plus
Cluster 63	Location: CM035916.1	Coordinates: 75318977-75342092	Size [bp]: 23116	Hits (absolute): 6778	Hits (normalized): 1467.34627787598	Hits (normalized) per kb: 63.4771778634407	Normalized hits with 1T: 86.7%	Normalized hits with 10A: 12.7%	Normalized hits 25-32 nt: 99.6%	Normalized hits on the main strand(s): 97.9%	Predicted directionality: mono:minus
Cluster 64	Location: CM035916.1	Coordinates: 75865439-75868907	Size [bp]: 3469	Hits (absolute): 436	Hits (normalized): 299.385586282621	Hits (normalized) per kb: 86.3034275570122	Normalized hits with 1T: 99.2%	Normalized hits with 10A: 3%	Normalized hits 25-32 nt: 99.8%	Normalized hits on the main strand(s): 99.9%	Predicted directionality: mono:minus
Cluster 65	Location: CM035916.1	Coordinates: 101413674-101419815	Size [bp]: 6142	Hits (absolute): 364	Hits (normalized): 122.127351363365	Hits (normalized) per kb: 19.8837134691984	Normalized hits with 1T: 4.3%	Normalized hits with 10A: 89.5%	Normalized hits 25-32 nt: 99.6%	Normalized hits on the main strand(s): 99.8%	Predicted directionality: mono:minus
Cluster 66	Location: CM035916.1	Coordinates: 102106009-102107069	Size [bp]: 1061	Hits (absolute): 258	Hits (normalized): 590.020192153428	Hits (normalized) per kb: 556.098215064272	Normalized hits with 1T: 0.1%	Normalized hits with 10A: 99.7%	Normalized hits 25-32 nt: 99.8%	Normalized hits on the main strand(s): 100%	Predicted directionality: mono:minus
Cluster 67	Location: CM035916.1	Coordinates: 117112203-117116761	Size [bp]: 4559	Hits (absolute): 307	Hits (normalized): 129.524683776294	Hits (normalized) per kb: 28.4110633439123	Normalized hits with 1T: 94.2%	Normalized hits with 10A: 4.9%	Normalized hits 25-32 nt: 97.3%	Normalized hits on the main strand(s): 99.9%	Predicted directionality: mono:plus
Cluster 68	Location: CM035916.1	Coordinates: 118290869-118297021	Size [bp]: 6153	Hits (absolute): 2097	Hits (normalized): 164.448053548953	Hits (normalized) per kb: 26.7263235654622	Normalized hits with 1T: 94.6%	Normalized hits with 10A: 34.3%	Normalized hits 25-32 nt: 99%	Normalized hits on the main strand(s): 99.4%	Predicted directionality: mono:plus
Cluster 69	Location: CM035916.1	Coordinates: 121064001-121073002	Size [bp]: 9002	Hits (absolute): 1606	Hits (normalized): 227.727534584477	Hits (normalized) per kb: 25.2974207357904	Normalized hits with 1T: 93.7%	Normalized hits with 10A: 7.2%	Normalized hits 25-32 nt: 99.5%	Normalized hits on the main strand(s): 100%	Predicted directionality: mono:minus
Cluster 70	Location: CM035916.1	Coordinates: 121084098-121090572	Size [bp]: 6475	Hits (absolute): 1294	Hits (normalized): 133.63305112532	Hits (normalized) per kb: 20.6380622860142	Normalized hits with 1T: 94.3%	Normalized hits with 10A: 18.9%	Normalized hits 25-32 nt: 99.4%	Normalized hits on the main strand(s): 99.9%	Predicted directionality: mono:minus
Cluster 71	Location: CM035916.1	Coordinates: 121098025-121118016	Size [bp]: 19992	Hits (absolute): 8191	Hits (normalized): 1549.27951032477	Hits (normalized) per kb: 77.4947393013266	Normalized hits with 1T: 89.5%	Normalized hits with 10A: 27.1%	Normalized hits 25-32 nt: 98.5%	Normalized hits on the main strand(s): 100%	Predicted directionality: mono:minus
Cluster 72	Location: CM035916.1	Coordinates: 128959186-128961191	Size [bp]: 2006	Hits (absolute): 378	Hits (normalized): 217.306407097063	Hits (normalized) per kb: 108.328438703606	Normalized hits with 1T: 98.5%	Normalized hits with 10A: 98.5%	Normalized hits 25-32 nt: 98.4%	Normalized hits on the main strand(s): 100%	Predicted directionality: mono:minus
Cluster 73	Location: CM035916.1	Coordinates: 134007326-134014023	Size [bp]: 6698	Hits (absolute): 1207	Hits (normalized): 125.171989888778	Hits (normalized) per kb: 18.6876140366464	Normalized hits with 1T: 87.7%	Normalized hits with 10A: 20.2%	Normalized hits 25-32 nt: 99.4%	Normalized hits on the main strand(s): 94.9%	Predicted directionality: mono:minus
Cluster 74	Location: CM035916.1	Coordinates: 141456109-141461793	Size [bp]: 5685	Hits (absolute): 835	Hits (normalized): 217.312425720619	Hits (normalized) per kb: 38.2258242309673	Normalized hits with 1T: 66%	Normalized hits with 10A: 95%	Normalized hits 25-32 nt: 99.3%	Normalized hits on the main strand(s): 99.8%	Predicted directionality: mono:minus
Cluster 75	Location: CM035916.1	Coordinates: 151400049-151407013	Size [bp]: 6965	Hits (absolute): 782	Hits (normalized): 405.852664683157	Hits (normalized) per kb: 58.2699499349302	Normalized hits with 1T: 4.4%	Normalized hits with 10A: 95.7%	Normalized hits 25-32 nt: 99.7%	Normalized hits on the main strand(s): 99.7%	Predicted directionality: mono:minus
Cluster 76	Location: CM035917.1	Coordinates: 6285156-6291984	Size [bp]: 6829	Hits (absolute): 812	Hits (normalized): 129.24049219639	Hits (normalized) per kb: 18.9253531948359	Normalized hits with 1T: 83.6%	Normalized hits with 10A: 59.4%	Normalized hits 25-32 nt: 96.4%	Normalized hits on the main strand(s): 99.9%	Predicted directionality: mono:plus
Cluster 77	Location: CM035917.1	Coordinates: 6293058-6308028	Size [bp]: 14971	Hits (absolute): 3370	Hits (normalized): 485.118831122065	Hits (normalized) per kb: 32.4040940492819	Normalized hits with 1T: 92.2%	Normalized hits with 10A: 30%	Normalized hits 25-32 nt: 99.3%	Normalized hits on the main strand(s): 99.8%	Predicted directionality: mono:plus
Cluster 78	Location: CM035917.1	Coordinates: 6344051-6356001	Size [bp]: 11951	Hits (absolute): 2357	Hits (normalized): 276.427173331264	Hits (normalized) per kb: 23.1297989993567	Normalized hits with 1T: 88.9%	Normalized hits with 10A: 23.5%	Normalized hits 25-32 nt: 99.8%	Normalized hits on the main strand(s): 100%	Predicted directionality: mono:plus
Cluster 79	Location: CM035917.1	Coordinates: 9948033-9955018	Size [bp]: 6986	Hits (absolute): 941	Hits (normalized): 135.977668164893	Hits (normalized) per kb: 19.4641737782757	Normalized hits with 1T: 98.3%	Normalized hits with 10A: 55.3%	Normalized hits 25-32 nt: 99.7%	Normalized hits on the main strand(s): 99.9%	Predicted directionality: mono:minus
Cluster 80	Location: CM035917.1	Coordinates: 9979114-9987761	Size [bp]: 8648	Hits (absolute): 1622	Hits (normalized): 206.847352432441	Hits (normalized) per kb: 23.9186981436603	Normalized hits with 1T: 88.3%	Normalized hits with 10A: 14.8%	Normalized hits 25-32 nt: 99.7%	Normalized hits on the main strand(s): 100%	Predicted directionality: mono:minus
Cluster 81	Location: CM035917.1	Coordinates: 10520015-10525008	Size [bp]: 4994	Hits (absolute): 719	Hits (normalized): 123.201816568628	Hits (normalized) per kb: 24.6697564530963	Normalized hits with 1T: 98.4%	Normalized hits with 10A: 59.6%	Normalized hits 25-32 nt: 99.8%	Normalized hits on the main strand(s): 99.9%	Predicted directionality: mono:plus
Cluster 82	Location: CM035917.1	Coordinates: 28530945-28537019	Size [bp]: 6075	Hits (absolute): 2361	Hits (normalized): 171.203930310274	Hits (normalized) per kb: 28.1815504541723	Normalized hits with 1T: 91.1%	Normalized hits with 10A: 38.9%	Normalized hits 25-32 nt: 98.9%	Normalized hits on the main strand(s): 99.6%	Predicted directionality: mono:plus
Cluster 83	Location: CM035917.1	Coordinates: 32821019-32825823	Size [bp]: 4805	Hits (absolute): 1272	Hits (normalized): 120.433451486052	Hits (normalized) per kb: 25.0637947118256	Normalized hits with 1T: 92.5%	Normalized hits with 10A: 10.9%	Normalized hits 25-32 nt: 99.4%	Normalized hits on the main strand(s): 98.8%	Predicted directionality: mono:plus
Cluster 84	Location: CM035917.1	Coordinates: 32828000-32836945	Size [bp]: 8946	Hits (absolute): 2959	Hits (normalized): 324.314176551182	Hits (normalized) per kb: 36.252342429941	Normalized hits with 1T: 88.5%	Normalized hits with 10A: 11.9%	Normalized hits 25-32 nt: 98.1%	Normalized hits on the main strand(s): 99.6%	Predicted directionality: mono:minus
Cluster 85	Location: CM035917.1	Coordinates: 35134813-35139756	Size [bp]: 4944	Hits (absolute): 1782	Hits (normalized): 572.284524453915	Hits (normalized) per kb: 115.753468606092	Normalized hits with 1T: 12.9%	Normalized hits with 10A: 88.1%	Normalized hits 25-32 nt: 99%	Normalized hits on the main strand(s): 93%	Predicted directionality: bi:plus-minus (split between 35138341 and 35138342)
Cluster 86	Location: CM035917.1	Coordinates: 38349232-38355524	Size [bp]: 6293	Hits (absolute): 1674	Hits (normalized): 192.241640918045	Hits (normalized) per kb: 30.548247887083	Normalized hits with 1T: 89.2%	Normalized hits with 10A: 28.8%	Normalized hits 25-32 nt: 99.4%	Normalized hits on the main strand(s): 89%	Predicted directionality: mono:minus
Cluster 87	Location: CM035917.1	Coordinates: 45972059-45984910	Size [bp]: 12852	Hits (absolute): 2491	Hits (normalized): 439.874712061775	Hits (normalized) per kb: 34.2262125108381	Normalized hits with 1T: 86.9%	Normalized hits with 10A: 12.9%	Normalized hits 25-32 nt: 97.2%	Normalized hits on the main strand(s): 99.4%	Predicted directionality: mono:plus
Cluster 88	Location: CM035917.1	Coordinates: 46555006-46562400	Size [bp]: 7395	Hits (absolute): 1401	Hits (normalized): 170.811715927413	Hits (normalized) per kb: 23.0985391792487	Normalized hits with 1T: 96.3%	Normalized hits with 10A: 52.1%	Normalized hits 25-32 nt: 99.7%	Normalized hits on the main strand(s): 97.3%	Predicted directionality: mono:plus
Cluster 89	Location: CM035917.1	Coordinates: 48288048-48295986	Size [bp]: 7939	Hits (absolute): 886	Hits (normalized): 155.885632054966	Hits (normalized) per kb: 19.6352801620246	Normalized hits with 1T: 86.9%	Normalized hits with 10A: 26.9%	Normalized hits 25-32 nt: 99.7%	Normalized hits on the main strand(s): 88.8%	Predicted directionality: mono:minus
Cluster 90	Location: CM035917.1	Coordinates: 55830054-55837500	Size [bp]: 7447	Hits (absolute): 550	Hits (normalized): 166.989234321653	Hits (normalized) per kb: 22.4239851663927	Normalized hits with 1T: 83.1%	Normalized hits with 10A: 84.1%	Normalized hits 25-32 nt: 99.8%	Normalized hits on the main strand(s): 98%	Predicted directionality: bi:minus-plus (split between 55834123 and 55834136)
Cluster 91	Location: CM035917.1	Coordinates: 56321095-56327799	Size [bp]: 6705	Hits (absolute): 908	Hits (normalized): 133.95470076095	Hits (normalized) per kb: 19.9783155563672	Normalized hits with 1T: 87.4%	Normalized hits with 10A: 27.2%	Normalized hits 25-32 nt: 99.7%	Normalized hits on the main strand(s): 86%	Predicted directionality: mono:minus
Cluster 92	Location: CM035917.1	Coordinates: 56350047-56383023	Size [bp]: 32977	Hits (absolute): 7952	Hits (normalized): 1995.10532747658	Hits (normalized) per kb: 60.5000913115798	Normalized hits with 1T: 89.9%	Normalized hits with 10A: 14.9%	Normalized hits 25-32 nt: 99.8%	Normalized hits on the main strand(s): 99.9%	Predicted directionality: mono:minus
Cluster 93	Location: CM035917.1	Coordinates: 56417229-56426023	Size [bp]: 8795	Hits (absolute): 1316	Hits (normalized): 306.502769872431	Hits (normalized) per kb: 34.8497636593074	Normalized hits with 1T: 95.6%	Normalized hits with 10A: 30.4%	Normalized hits 25-32 nt: 99.6%	Normalized hits on the main strand(s): 99.8%	Predicted directionality: mono:minus
Cluster 94	Location: CM035917.1	Coordinates: 56427129-56436014	Size [bp]: 8886	Hits (absolute): 1132	Hits (normalized): 293.124861116204	Hits (normalized) per kb: 32.9873364823489	Normalized hits with 1T: 98.3%	Normalized hits with 10A: 19%	Normalized hits 25-32 nt: 99.8%	Normalized hits on the main strand(s): 99.9%	Predicted directionality: mono:minus
Cluster 95	Location: CM035917.1	Coordinates: 56445070-56488020	Size [bp]: 42951	Hits (absolute): 11172	Hits (normalized): 2128.73008869937	Hits (normalized) per kb: 49.5616221543281	Normalized hits with 1T: 87.7%	Normalized hits with 10A: 20.9%	Normalized hits 25-32 nt: 99.6%	Normalized hits on the main strand(s): 99.7%	Predicted directionality: mono:minus
Cluster 96	Location: CM035917.1	Coordinates: 67206075-67210763	Size [bp]: 4689	Hits (absolute): 1299	Hits (normalized): 167.830952221658	Hits (normalized) per kb: 35.792494023616	Normalized hits with 1T: 96.4%	Normalized hits with 10A: 52.6%	Normalized hits 25-32 nt: 99.7%	Normalized hits on the main strand(s): 97.4%	Predicted directionality: mono:plus
Cluster 97	Location: CM035917.1	Coordinates: 67390470-67395706	Size [bp]: 5237	Hits (absolute): 1323	Hits (normalized): 169.426878007113	Hits (normalized) per kb: 32.3522685580503	Normalized hits with 1T: 96.6%	Normalized hits with 10A: 52.3%	Normalized hits 25-32 nt: 99.7%	Normalized hits on the main strand(s): 97.5%	Predicted directionality: mono:minus
Cluster 98	Location: CM035917.1	Coordinates: 73531009-73538955	Size [bp]: 7947	Hits (absolute): 1626	Hits (normalized): 212.792932388092	Hits (normalized) per kb: 26.7765038030039	Normalized hits with 1T: 91.7%	Normalized hits with 10A: 12.8%	Normalized hits 25-32 nt: 99.7%	Normalized hits on the main strand(s): 98.2%	Predicted directionality: mono:minus
Cluster 99	Location: CM035917.1	Coordinates: 73552110-73558981	Size [bp]: 6872	Hits (absolute): 1700	Hits (normalized): 487.882411146442	Hits (normalized) per kb: 70.9959872262503	Normalized hits with 1T: 98%	Normalized hits with 10A: 20%	Normalized hits 25-32 nt: 99.2%	Normalized hits on the main strand(s): 99.9%	Predicted directionality: mono:minus
Cluster 100	Location: CM035917.1	Coordinates: 73651155-73659995	Size [bp]: 8841	Hits (absolute): 1306	Hits (normalized): 284.454205961917	Hits (normalized) per kb: 32.1745811595419	Normalized hits with 1T: 88.3%	Normalized hits with 10A: 4.6%	Normalized hits 25-32 nt: 99.7%	Normalized hits on the main strand(s): 99.8%	Predicted directionality: mono:minus
Cluster 101	Location: CM035917.1	Coordinates: 73764004-73776002	Size [bp]: 11999	Hits (absolute): 1944	Hits (normalized): 347.752619041482	Hits (normalized) per kb: 28.9819663743051	Normalized hits with 1T: 94.3%	Normalized hits with 10A: 22.1%	Normalized hits 25-32 nt: 99.4%	Normalized hits on the main strand(s): 98.9%	Predicted directionality: mono:minus
Cluster 102	Location: CM035917.1	Coordinates: 74268108-74276928	Size [bp]: 8821	Hits (absolute): 1220	Hits (normalized): 155.378708650706	Hits (normalized) per kb: 17.6149086308363	Normalized hits with 1T: 93.9%	Normalized hits with 10A: 8.1%	Normalized hits 25-32 nt: 99.7%	Normalized hits on the main strand(s): 99.8%	Predicted directionality: mono:minus
Cluster 103	Location: CM035917.1	Coordinates: 81885229-81888899	Size [bp]: 3671	Hits (absolute): 341	Hits (normalized): 123.995786806578	Hits (normalized) per kb: 33.7770582534974	Normalized hits with 1T: 99.1%	Normalized hits with 10A: 41.9%	Normalized hits 25-32 nt: 99.8%	Normalized hits on the main strand(s): 100%	Predicted directionality: mono:minus
Cluster 104	Location: CM035917.1	Coordinates: 87968032-87975017	Size [bp]: 6986	Hits (absolute): 2232	Hits (normalized): 724.289926071832	Hits (normalized) per kb: 103.677306522279	Normalized hits with 1T: 97.2%	Normalized hits with 10A: 13%	Normalized hits 25-32 nt: 99.5%	Normalized hits on the main strand(s): 100%	Predicted directionality: mono:plus
Cluster 105	Location: CM035917.1	Coordinates: 87979367-87991895	Size [bp]: 12529	Hits (absolute): 6013	Hits (normalized): 2719.65581965853	Hits (normalized) per kb: 217.06901345653	Normalized hits with 1T: 98.7%	Normalized hits with 10A: 29.8%	Normalized hits 25-32 nt: 99.4%	Normalized hits on the main strand(s): 100%	Predicted directionality: mono:plus
Cluster 106	Location: CM035917.1	Coordinates: 103477201-103485734	Size [bp]: 8534	Hits (absolute): 869	Hits (normalized): 154.243002985839	Hits (normalized) per kb: 18.0739344103164	Normalized hits with 1T: 92.6%	Normalized hits with 10A: 10.9%	Normalized hits 25-32 nt: 99.6%	Normalized hits on the main strand(s): 92.9%	Predicted directionality: mono:minus
Cluster 107	Location: CM035917.1	Coordinates: 111049122-111063015	Size [bp]: 13894	Hits (absolute): 2414	Hits (normalized): 993.924988519726	Hits (normalized) per kb: 71.5364530633801	Normalized hits with 1T: 99.1%	Normalized hits with 10A: 11.1%	Normalized hits 25-32 nt: 99.9%	Normalized hits on the main strand(s): 99.9%	Predicted directionality: mono:minus
Cluster 108	Location: CM035917.1	Coordinates: 111094109-111116316	Size [bp]: 22208	Hits (absolute): 11552	Hits (normalized): 2991.58592405366	Hits (normalized) per kb: 134.707613740501	Normalized hits with 1T: 88.8%	Normalized hits with 10A: 26.6%	Normalized hits 25-32 nt: 99.4%	Normalized hits on the main strand(s): 98.4%	Predicted directionality: mono:minus
Cluster 109	Location: CM035917.1	Coordinates: 111151091-111158022	Size [bp]: 6932	Hits (absolute): 1141	Hits (normalized): 197.452820700135	Hits (normalized) per kb: 28.4842771331125	Normalized hits with 1T: 95.2%	Normalized hits with 10A: 16.8%	Normalized hits 25-32 nt: 99.3%	Normalized hits on the main strand(s): 98.2%	Predicted directionality: mono:minus
Cluster 110	Location: CM035917.1	Coordinates: 111705053-111712910	Size [bp]: 7858	Hits (absolute): 1137	Hits (normalized): 192.140994916837	Hits (normalized) per kb: 24.4517603391855	Normalized hits with 1T: 95%	Normalized hits with 10A: 16.6%	Normalized hits 25-32 nt: 99.3%	Normalized hits on the main strand(s): 98.1%	Predicted directionality: mono:plus
Cluster 111	Location: CM035917.1	Coordinates: 111765006-111778993	Size [bp]: 13988	Hits (absolute): 2488	Hits (normalized): 901.033145761817	Hits (normalized) per kb: 64.4149724666795	Normalized hits with 1T: 96.5%	Normalized hits with 10A: 11%	Normalized hits 25-32 nt: 99.8%	Normalized hits on the main strand(s): 99.9%	Predicted directionality: mono:plus
Cluster 112	Location: CM035917.1	Coordinates: 111787112-111796024	Size [bp]: 8913	Hits (absolute): 2078	Hits (normalized): 179.910228006204	Hits (normalized) per kb: 20.1847948944487	Normalized hits with 1T: 95.6%	Normalized hits with 10A: 27.1%	Normalized hits 25-32 nt: 99.8%	Normalized hits on the main strand(s): 93.5%	Predicted directionality: mono:plus
Cluster 113	Location: CM035917.1	Coordinates: 113771526-113783417	Size [bp]: 11892	Hits (absolute): 10914	Hits (normalized): 13225.8132626093	Hits (normalized) per kb: 1112.16023454737	Normalized hits with 1T: 86.1%	Normalized hits with 10A: 69.6%	Normalized hits 25-32 nt: 99.9%	Normalized hits on the main strand(s): 100%	Predicted directionality: mono:plus
Cluster 114	Location: CM035917.1	Coordinates: 119910005-119916534	Size [bp]: 6530	Hits (absolute): 1592	Hits (normalized): 491.874712768128	Hits (normalized) per kb: 75.325472311203	Normalized hits with 1T: 98.8%	Normalized hits with 10A: 8.7%	Normalized hits 25-32 nt: 99.8%	Normalized hits on the main strand(s): 99.1%	Predicted directionality: mono:plus
Cluster 115	Location: CM035917.1	Coordinates: 121408176-121421919	Size [bp]: 13744	Hits (absolute): 3443	Hits (normalized): 1353.17392408584	Hits (normalized) per kb: 98.4552713105597	Normalized hits with 1T: 93.2%	Normalized hits with 10A: 20.3%	Normalized hits 25-32 nt: 99.7%	Normalized hits on the main strand(s): 100%	Predicted directionality: mono:plus
Cluster 116	Location: CM035917.1	Coordinates: 121424310-121437958	Size [bp]: 13649	Hits (absolute): 2639	Hits (normalized): 641.976851227274	Hits (normalized) per kb: 47.0345124866529	Normalized hits with 1T: 95.6%	Normalized hits with 10A: 34.9%	Normalized hits 25-32 nt: 99.7%	Normalized hits on the main strand(s): 100%	Predicted directionality: mono:plus
Cluster 117	Location: CM035917.1	Coordinates: 127665025-127688003	Size [bp]: 22979	Hits (absolute): 3715	Hits (normalized): 1202.18583562977	Hits (normalized) per kb: 52.3165994580534	Normalized hits with 1T: 97.3%	Normalized hits with 10A: 27.5%	Normalized hits 25-32 nt: 99.3%	Normalized hits on the main strand(s): 99.1%	Predicted directionality: mono:plus
Cluster 118	Location: CM035917.1	Coordinates: 127774524-127779886	Size [bp]: 5363	Hits (absolute): 861	Hits (normalized): 330.643271581165	Hits (normalized) per kb: 61.6525915213496	Normalized hits with 1T: 90%	Normalized hits with 10A: 40.9%	Normalized hits 25-32 nt: 99.8%	Normalized hits on the main strand(s): 91.5%	Predicted directionality: mono:plus
Cluster 119	Location: CM035917.1	Coordinates: 127793044-127800975	Size [bp]: 7932	Hits (absolute): 781	Hits (normalized): 184.856573861275	Hits (normalized) per kb: 23.3050185173303	Normalized hits with 1T: 86.9%	Normalized hits with 10A: 17.5%	Normalized hits 25-32 nt: 99.5%	Normalized hits on the main strand(s): 86.1%	Predicted directionality: mono:plus
Cluster 120	Location: CM035917.1	Coordinates: 127814161-127823018	Size [bp]: 8858	Hits (absolute): 1523	Hits (normalized): 547.438307820975	Hits (normalized) per kb: 61.8014869802849	Normalized hits with 1T: 99.1%	Normalized hits with 10A: 15.1%	Normalized hits 25-32 nt: 98.9%	Normalized hits on the main strand(s): 99.5%	Predicted directionality: mono:plus
Cluster 121	Location: CM035917.1	Coordinates: 141732036-141740563	Size [bp]: 8528	Hits (absolute): 2014	Hits (normalized): 201.186947246091	Hits (normalized) per kb: 23.5912926593716	Normalized hits with 1T: 85.5%	Normalized hits with 10A: 31%	Normalized hits 25-32 nt: 99.2%	Normalized hits on the main strand(s): 98.4%	Predicted directionality: mono:minus
Cluster 122	Location: CM035917.1	Coordinates: 149200098-149208709	Size [bp]: 8612	Hits (absolute): 1213	Hits (normalized): 213.942071714415	Hits (normalized) per kb: 24.842508090535	Normalized hits with 1T: 82.9%	Normalized hits with 10A: 17.3%	Normalized hits 25-32 nt: 99.7%	Normalized hits on the main strand(s): 99.7%	Predicted directionality: mono:minus
Cluster 123	Location: CM035917.1	Coordinates: 151653590-151660904	Size [bp]: 7315	Hits (absolute): 1479	Hits (normalized): 318.771246021256	Hits (normalized) per kb: 43.5778344841883	Normalized hits with 1T: 96.7%	Normalized hits with 10A: 6.4%	Normalized hits 25-32 nt: 99.8%	Normalized hits on the main strand(s): 96.3%	Predicted directionality: mono:plus
Cluster 124	Location: CM035918.1	Coordinates: 11881922-11889026	Size [bp]: 7105	Hits (absolute): 1660	Hits (normalized): 166.028719548411	Hits (normalized) per kb: 23.3675381575462	Normalized hits with 1T: 91.5%	Normalized hits with 10A: 36.4%	Normalized hits 25-32 nt: 99.4%	Normalized hits on the main strand(s): 97.5%	Predicted directionality: mono:plus
Cluster 125	Location: CM035918.1	Coordinates: 20662002-20708997	Size [bp]: 46996	Hits (absolute): 14975	Hits (normalized): 2547.18182667545	Hits (normalized) per kb: 54.1995923061356	Normalized hits with 1T: 86.5%	Normalized hits with 10A: 14.4%	Normalized hits 25-32 nt: 99.6%	Normalized hits on the main strand(s): 99.5%	Predicted directionality: mono:plus
Cluster 126	Location: CM035918.1	Coordinates: 20727001-20748910	Size [bp]: 21910	Hits (absolute): 5589	Hits (normalized): 984.530417157957	Hits (normalized) per kb: 44.9351687783498	Normalized hits with 1T: 86.2%	Normalized hits with 10A: 31.6%	Normalized hits 25-32 nt: 99.6%	Normalized hits on the main strand(s): 93.3%	Predicted directionality: mono:plus
Cluster 127	Location: CM035918.1	Coordinates: 31759041-31767022	Size [bp]: 7982	Hits (absolute): 880	Hits (normalized): 149.679752677044	Hits (normalized) per kb: 18.7517789305522	Normalized hits with 1T: 88.7%	Normalized hits with 10A: 27.5%	Normalized hits 25-32 nt: 99.6%	Normalized hits on the main strand(s): 98.4%	Predicted directionality: mono:plus
Cluster 128	Location: CM035918.1	Coordinates: 32460015-32465691	Size [bp]: 5677	Hits (absolute): 2261	Hits (normalized): 175.937424067137	Hits (normalized) per kb: 30.991643756509	Normalized hits with 1T: 95.7%	Normalized hits with 10A: 28%	Normalized hits 25-32 nt: 99.4%	Normalized hits on the main strand(s): 94.8%	Predicted directionality: bi:plus-minus (split between 32460534 and 32460535)
Cluster 129	Location: CM035918.1	Coordinates: 32613168-32622992	Size [bp]: 9825	Hits (absolute): 1525	Hits (normalized): 253.603026068837	Hits (normalized) per kb: 25.8123851407268	Normalized hits with 1T: 96%	Normalized hits with 10A: 11%	Normalized hits 25-32 nt: 99.9%	Normalized hits on the main strand(s): 99.9%	Predicted directionality: mono:minus
Cluster 130	Location: CM035918.1	Coordinates: 34279094-34283559	Size [bp]: 4466	Hits (absolute): 287	Hits (normalized): 197.50206060811	Hits (normalized) per kb: 44.2235965574712	Normalized hits with 1T: 85.4%	Normalized hits with 10A: 0.9%	Normalized hits 25-32 nt: 97.9%	Normalized hits on the main strand(s): 99.8%	Predicted directionality: mono:minus
Cluster 131	Location: CM035918.1	Coordinates: 34381272-34384233	Size [bp]: 2962	Hits (absolute): 274	Hits (normalized): 197.305158996866	Hits (normalized) per kb: 66.6122087695312	Normalized hits with 1T: 85.4%	Normalized hits with 10A: 0.9%	Normalized hits 25-32 nt: 97.9%	Normalized hits on the main strand(s): 99.9%	Predicted directionality: mono:minus
Cluster 132	Location: CM035918.1	Coordinates: 53359024-53372013	Size [bp]: 12990	Hits (absolute): 4440	Hits (normalized): 877.514409224909	Hits (normalized) per kb: 67.5532938801499	Normalized hits with 1T: 94.2%	Normalized hits with 10A: 33.6%	Normalized hits 25-32 nt: 99.3%	Normalized hits on the main strand(s): 99.6%	Predicted directionality: mono:plus
Cluster 133	Location: CM035918.1	Coordinates: 53382044-53397889	Size [bp]: 15846	Hits (absolute): 6154	Hits (normalized): 3317.08127528548	Hits (normalized) per kb: 209.332207979809	Normalized hits with 1T: 97.7%	Normalized hits with 10A: 28.8%	Normalized hits 25-32 nt: 99.4%	Normalized hits on the main strand(s): 100%	Predicted directionality: mono:plus
Cluster 134	Location: CM035918.1	Coordinates: 53399054-53405990	Size [bp]: 6937	Hits (absolute): 946	Hits (normalized): 223.349169009925	Hits (normalized) per kb: 32.1967920843555	Normalized hits with 1T: 94.8%	Normalized hits with 10A: 30.6%	Normalized hits 25-32 nt: 99.6%	Normalized hits on the main strand(s): 100%	Predicted directionality: mono:plus
Cluster 135	Location: CM035918.1	Coordinates: 53412182-53420937	Size [bp]: 8756	Hits (absolute): 869	Hits (normalized): 163.093832708177	Hits (normalized) per kb: 18.6267396501203	Normalized hits with 1T: 95.3%	Normalized hits with 10A: 6.5%	Normalized hits 25-32 nt: 99.6%	Normalized hits on the main strand(s): 98.6%	Predicted directionality: mono:plus
Cluster 136	Location: CM035918.1	Coordinates: 60552058-60560841	Size [bp]: 8784	Hits (absolute): 1358	Hits (normalized): 318.671824631154	Hits (normalized) per kb: 36.2786664889793	Normalized hits with 1T: 99.3%	Normalized hits with 10A: 2.3%	Normalized hits 25-32 nt: 85%	Normalized hits on the main strand(s): 99.9%	Predicted directionality: mono:minus
Cluster 137	Location: CM035918.1	Coordinates: 68128030-68139023	Size [bp]: 10994	Hits (absolute): 1768	Hits (normalized): 1369.9073479608	Hits (normalized) per kb: 124.604933457715	Normalized hits with 1T: 97.3%	Normalized hits with 10A: 8.7%	Normalized hits 25-32 nt: 99.8%	Normalized hits on the main strand(s): 100%	Predicted directionality: mono:minus
Cluster 138	Location: CM035918.1	Coordinates: 68147140-68159947	Size [bp]: 12808	Hits (absolute): 4450	Hits (normalized): 4190.90791879089	Hits (normalized) per kb: 327.210521726398	Normalized hits with 1T: 98.6%	Normalized hits with 10A: 6.2%	Normalized hits 25-32 nt: 98.2%	Normalized hits on the main strand(s): 100%	Predicted directionality: mono:minus
Cluster 139	Location: CM035918.1	Coordinates: 68190122-68209997	Size [bp]: 19876	Hits (absolute): 6368	Hits (normalized): 2741.03457446044	Hits (normalized) per kb: 137.906809540497	Normalized hits with 1T: 91.8%	Normalized hits with 10A: 16.8%	Normalized hits 25-32 nt: 99.6%	Normalized hits on the main strand(s): 99.9%	Predicted directionality: mono:minus
Cluster 140	Location: CM035918.1	Coordinates: 68218001-68237029	Size [bp]: 19029	Hits (absolute): 6552	Hits (normalized): 3254.04116317296	Hits (normalized) per kb: 171.004378020069	Normalized hits with 1T: 86.3%	Normalized hits with 10A: 15.9%	Normalized hits 25-32 nt: 99.7%	Normalized hits on the main strand(s): 99.9%	Predicted directionality: mono:minus
Cluster 141	Location: CM035918.1	Coordinates: 83126397-83135008	Size [bp]: 8612	Hits (absolute): 935	Hits (normalized): 259.056778801674	Hits (normalized) per kb: 30.0809958391534	Normalized hits with 1T: 97.1%	Normalized hits with 10A: 15.3%	Normalized hits 25-32 nt: 99.4%	Normalized hits on the main strand(s): 99.8%	Predicted directionality: mono:plus
Cluster 142	Location: CM035918.1	Coordinates: 89575003-89597008	Size [bp]: 22006	Hits (absolute): 3724	Hits (normalized): 1127.53886347262	Hits (normalized) per kb: 51.2381356643288	Normalized hits with 1T: 98.8%	Normalized hits with 10A: 28.8%	Normalized hits 25-32 nt: 99.5%	Normalized hits on the main strand(s): 99.8%	Predicted directionality: mono:plus
Cluster 143	Location: CM035918.1	Coordinates: 92929231-92934847	Size [bp]: 5617	Hits (absolute): 2215	Hits (normalized): 162.678571722367	Hits (normalized) per kb: 28.9622233300264	Normalized hits with 1T: 94.9%	Normalized hits with 10A: 38.5%	Normalized hits 25-32 nt: 99.6%	Normalized hits on the main strand(s): 97.7%	Predicted directionality: mono:plus
Cluster 144	Location: CM035918.1	Coordinates: 103317452-103322923	Size [bp]: 5472	Hits (absolute): 1129	Hits (normalized): 120.342249891171	Hits (normalized) per kb: 21.9921060727959	Normalized hits with 1T: 96%	Normalized hits with 10A: 10.9%	Normalized hits 25-32 nt: 99.2%	Normalized hits on the main strand(s): 99.8%	Predicted directionality: mono:plus
Cluster 145	Location: CM035918.1	Coordinates: 103687033-103696016	Size [bp]: 8984	Hits (absolute): 636	Hits (normalized): 288.631444065523	Hits (normalized) per kb: 32.126868802535	Normalized hits with 1T: 97.7%	Normalized hits with 10A: 7.4%	Normalized hits 25-32 nt: 100%	Normalized hits on the main strand(s): 100%	Predicted directionality: mono:plus
Cluster 146	Location: CM035918.1	Coordinates: 103712001-103722742	Size [bp]: 10742	Hits (absolute): 3030	Hits (normalized): 765.254571326944	Hits (normalized) per kb: 71.2394847723545	Normalized hits with 1T: 86%	Normalized hits with 10A: 17.6%	Normalized hits 25-32 nt: 99.6%	Normalized hits on the main strand(s): 99.8%	Predicted directionality: mono:plus
Cluster 147	Location: CM035918.1	Coordinates: 111214236-111221601	Size [bp]: 7366	Hits (absolute): 2165	Hits (normalized): 182.742701917579	Hits (normalized) per kb: 24.8087803898922	Normalized hits with 1T: 95.2%	Normalized hits with 10A: 25.6%	Normalized hits 25-32 nt: 99.8%	Normalized hits on the main strand(s): 93.2%	Predicted directionality: mono:plus
Cluster 148	Location: CM035918.1	Coordinates: 116572025-116577937	Size [bp]: 5913	Hits (absolute): 2664	Hits (normalized): 207.284451960201	Hits (normalized) per kb: 35.055420370544	Normalized hits with 1T: 91.4%	Normalized hits with 10A: 25.4%	Normalized hits 25-32 nt: 99.4%	Normalized hits on the main strand(s): 91.6%	Predicted directionality: mono:minus
Cluster 149	Location: CM035918.1	Coordinates: 116621338-116628986	Size [bp]: 7649	Hits (absolute): 2583	Hits (normalized): 220.759432250187	Hits (normalized) per kb: 28.861040228098	Normalized hits with 1T: 92.2%	Normalized hits with 10A: 23.2%	Normalized hits 25-32 nt: 99.4%	Normalized hits on the main strand(s): 92.1%	Predicted directionality: mono:plus
Cluster 150	Location: CM035918.1	Coordinates: 116738078-116746013	Size [bp]: 7936	Hits (absolute): 1989	Hits (normalized): 187.62879462601	Hits (normalized) per kb: 23.6431181506032	Normalized hits with 1T: 93.5%	Normalized hits with 10A: 20.1%	Normalized hits 25-32 nt: 99.5%	Normalized hits on the main strand(s): 93.8%	Predicted directionality: mono:plus
Cluster 151	Location: CM035918.1	Coordinates: 117002029-117008974	Size [bp]: 6946	Hits (absolute): 1479	Hits (normalized): 139.641408958809	Hits (normalized) per kb: 20.104177463644	Normalized hits with 1T: 94.8%	Normalized hits with 10A: 19.7%	Normalized hits 25-32 nt: 99.6%	Normalized hits on the main strand(s): 95.2%	Predicted directionality: mono:minus
Cluster 152	Location: CM035918.1	Coordinates: 121522201-121528003	Size [bp]: 5803	Hits (absolute): 1034	Hits (normalized): 134.740856083365	Hits (normalized) per kb: 23.2194653254559	Normalized hits with 1T: 79.7%	Normalized hits with 10A: 13.1%	Normalized hits 25-32 nt: 99.6%	Normalized hits on the main strand(s): 100%	Predicted directionality: mono:minus
Cluster 153	Location: CM035918.1	Coordinates: 121544112-121552780	Size [bp]: 8669	Hits (absolute): 1702	Hits (normalized): 199.62481542662	Hits (normalized) per kb: 23.0277932705834	Normalized hits with 1T: 91.5%	Normalized hits with 10A: 13.6%	Normalized hits 25-32 nt: 99.5%	Normalized hits on the main strand(s): 100%	Predicted directionality: mono:minus
Cluster 154	Location: CM035918.1	Coordinates: 132560003-132567896	Size [bp]: 7894	Hits (absolute): 706	Hits (normalized): 142.192666792573	Hits (normalized) per kb: 18.0130600237904	Normalized hits with 1T: 97%	Normalized hits with 10A: 74%	Normalized hits 25-32 nt: 99.3%	Normalized hits on the main strand(s): 98.4%	Predicted directionality: mono:minus
Cluster 155	Location: CM035918.1	Coordinates: 137035727-137043846	Size [bp]: 8120	Hits (absolute): 3136	Hits (normalized): 212.849427440429	Hits (normalized) per kb: 26.2130044142156	Normalized hits with 1T: 94.9%	Normalized hits with 10A: 31.8%	Normalized hits 25-32 nt: 99.4%	Normalized hits on the main strand(s): 94.6%	Predicted directionality: bi:plus-minus (split between 137037296 and 137037306)
Cluster 156	Location: CM035918.1	Coordinates: 138162086-138174879	Size [bp]: 12794	Hits (absolute): 1361	Hits (normalized): 488.502565250024	Hits (normalized) per kb: 38.1822250081851	Normalized hits with 1T: 98.9%	Normalized hits with 10A: 8.5%	Normalized hits 25-32 nt: 99.8%	Normalized hits on the main strand(s): 99.2%	Predicted directionality: mono:minus
Cluster 157	Location: CM035919.1	Coordinates: 4709164-4713369	Size [bp]: 4206	Hits (absolute): 988	Hits (normalized): 133.859759672309	Hits (normalized) per kb: 31.8257873772846	Normalized hits with 1T: 19.4%	Normalized hits with 10A: 85.8%	Normalized hits 25-32 nt: 98.3%	Normalized hits on the main strand(s): 90.3%	Predicted directionality: bi:plus-minus (split between 4710551 and 4710555)
Cluster 158	Location: CM035919.1	Coordinates: 15793029-15800156	Size [bp]: 7128	Hits (absolute): 1617	Hits (normalized): 549.207481297579	Hits (normalized) per kb: 77.0496981782106	Normalized hits with 1T: 73.6%	Normalized hits with 10A: 84%	Normalized hits 25-32 nt: 99.4%	Normalized hits on the main strand(s): 94.5%	Predicted directionality: bi:plus-minus (split between 15798903 and 15798905)
Cluster 159	Location: CM035919.1	Coordinates: 16697216-16705750	Size [bp]: 8535	Hits (absolute): 963	Hits (normalized): 354.110165435967	Hits (normalized) per kb: 41.4891849248695	Normalized hits with 1T: 98.7%	Normalized hits with 10A: 60.4%	Normalized hits 25-32 nt: 99.7%	Normalized hits on the main strand(s): 99.5%	Predicted directionality: mono:minus
Cluster 160	Location: CM035919.1	Coordinates: 16709219-16719560	Size [bp]: 10342	Hits (absolute): 3133	Hits (normalized): 1683.99966400237	Hits (normalized) per kb: 162.830757688682	Normalized hits with 1T: 97.9%	Normalized hits with 10A: 63.8%	Normalized hits 25-32 nt: 99.7%	Normalized hits on the main strand(s): 98.8%	Predicted directionality: mono:minus
Cluster 161	Location: CM035919.1	Coordinates: 16736061-16745993	Size [bp]: 9933	Hits (absolute): 2013	Hits (normalized): 351.38369584043	Hits (normalized) per kb: 35.3754222132282	Normalized hits with 1T: 90.3%	Normalized hits with 10A: 15.7%	Normalized hits 25-32 nt: 99.5%	Normalized hits on the main strand(s): 99.9%	Predicted directionality: mono:minus
Cluster 162	Location: CM035919.1	Coordinates: 17417003-17424995	Size [bp]: 7993	Hits (absolute): 1704	Hits (normalized): 225.008582575998	Hits (normalized) per kb: 28.1511132609093	Normalized hits with 1T: 86.2%	Normalized hits with 10A: 19.7%	Normalized hits 25-32 nt: 99.4%	Normalized hits on the main strand(s): 99.8%	Predicted directionality: mono:plus
Cluster 163	Location: CM035919.1	Coordinates: 17457040-17476891	Size [bp]: 19852	Hits (absolute): 3498	Hits (normalized): 1842.93163183356	Hits (normalized) per kb: 92.8334394521963	Normalized hits with 1T: 97.3%	Normalized hits with 10A: 70.4%	Normalized hits 25-32 nt: 99.7%	Normalized hits on the main strand(s): 99.8%	Predicted directionality: mono:plus
Cluster 164	Location: CM035919.1	Coordinates: 23738369-23793948	Size [bp]: 55580	Hits (absolute): 15628	Hits (normalized): 5664.03133673306	Hits (normalized) per kb: 101.9078361788	Normalized hits with 1T: 97.1%	Normalized hits with 10A: 44%	Normalized hits 25-32 nt: 99.8%	Normalized hits on the main strand(s): 99.9%	Predicted directionality: mono:minus
Cluster 165	Location: CM035919.1	Coordinates: 23807215-23822016	Size [bp]: 14802	Hits (absolute): 2786	Hits (normalized): 492.35427293408	Hits (normalized) per kb: 33.2629164754059	Normalized hits with 1T: 88.6%	Normalized hits with 10A: 24.4%	Normalized hits 25-32 nt: 99.7%	Normalized hits on the main strand(s): 100%	Predicted directionality: mono:minus
Cluster 166	Location: CM035919.1	Coordinates: 36846020-36862951	Size [bp]: 16932	Hits (absolute): 2926	Hits (normalized): 380.794788284394	Hits (normalized) per kb: 22.4897953139884	Normalized hits with 1T: 89.9%	Normalized hits with 10A: 12.8%	Normalized hits 25-32 nt: 99.6%	Normalized hits on the main strand(s): 100%	Predicted directionality: mono:plus
Cluster 167	Location: CM035919.1	Coordinates: 36874056-36879025	Size [bp]: 4970	Hits (absolute): 890	Hits (normalized): 147.48118794001	Hits (normalized) per kb: 29.6746181777499	Normalized hits with 1T: 97.4%	Normalized hits with 10A: 14.9%	Normalized hits 25-32 nt: 99.5%	Normalized hits on the main strand(s): 100%	Predicted directionality: mono:plus
Cluster 168	Location: CM035919.1	Coordinates: 36881282-36889685	Size [bp]: 8404	Hits (absolute): 854	Hits (normalized): 183.634497227777	Hits (normalized) per kb: 21.8506142554651	Normalized hits with 1T: 96%	Normalized hits with 10A: 15.5%	Normalized hits 25-32 nt: 99.8%	Normalized hits on the main strand(s): 100%	Predicted directionality: mono:plus
Cluster 169	Location: CM035919.1	Coordinates: 36908043-36916591	Size [bp]: 8549	Hits (absolute): 754	Hits (normalized): 154.839988147211	Hits (normalized) per kb: 18.1117752451839	Normalized hits with 1T: 96.2%	Normalized hits with 10A: 12.4%	Normalized hits 25-32 nt: 99.9%	Normalized hits on the main strand(s): 100%	Predicted directionality: mono:plus
Cluster 170	Location: CM035919.1	Coordinates: 41910047-41919021	Size [bp]: 8975	Hits (absolute): 1323	Hits (normalized): 250.757731760034	Hits (normalized) per kb: 27.9396981617581	Normalized hits with 1T: 81.1%	Normalized hits with 10A: 10.2%	Normalized hits 25-32 nt: 99.6%	Normalized hits on the main strand(s): 100%	Predicted directionality: mono:minus
Cluster 171	Location: CM035919.1	Coordinates: 41924009-41934933	Size [bp]: 10925	Hits (absolute): 2433	Hits (normalized): 438.83041203236	Hits (normalized) per kb: 40.1672235850407	Normalized hits with 1T: 95.6%	Normalized hits with 10A: 30.1%	Normalized hits 25-32 nt: 99.7%	Normalized hits on the main strand(s): 99.9%	Predicted directionality: mono:minus
Cluster 172	Location: CM035919.1	Coordinates: 41953094-41979859	Size [bp]: 26766	Hits (absolute): 6556	Hits (normalized): 1043.54593336791	Hits (normalized) per kb: 38.9875766893876	Normalized hits with 1T: 92.9%	Normalized hits with 10A: 18.5%	Normalized hits 25-32 nt: 99.1%	Normalized hits on the main strand(s): 100%	Predicted directionality: mono:minus
Cluster 173	Location: CM035919.1	Coordinates: 43759146-43766031	Size [bp]: 6886	Hits (absolute): 2504	Hits (normalized): 194.083391818876	Hits (normalized) per kb: 28.1848409615521	Normalized hits with 1T: 91.1%	Normalized hits with 10A: 23.7%	Normalized hits 25-32 nt: 99.3%	Normalized hits on the main strand(s): 90.4%	Predicted directionality: mono:minus
Cluster 174	Location: CM035919.1	Coordinates: 47845172-47847707	Size [bp]: 2536	Hits (absolute): 328	Hits (normalized): 221.23428290514	Hits (normalized) per kb: 87.2371090260263	Normalized hits with 1T: 75.7%	Normalized hits with 10A: 0.3%	Normalized hits 25-32 nt: 96.5%	Normalized hits on the main strand(s): 100%	Predicted directionality: mono:plus
Cluster 175	Location: CM035919.1	Coordinates: 50601172-50619926	Size [bp]: 18755	Hits (absolute): 6926	Hits (normalized): 1981.51173528411	Hits (normalized) per kb: 105.652433576995	Normalized hits with 1T: 94.2%	Normalized hits with 10A: 30.7%	Normalized hits 25-32 nt: 99.7%	Normalized hits on the main strand(s): 99.9%	Predicted directionality: mono:minus
Cluster 176	Location: CM035919.1	Coordinates: 50623102-50630026	Size [bp]: 6925	Hits (absolute): 1195	Hits (normalized): 155.872007861335	Hits (normalized) per kb: 22.5087157314222	Normalized hits with 1T: 97.6%	Normalized hits with 10A: 26.7%	Normalized hits 25-32 nt: 99.7%	Normalized hits on the main strand(s): 100%	Predicted directionality: mono:minus
Cluster 177	Location: CM035919.1	Coordinates: 54647039-54653118	Size [bp]: 6080	Hits (absolute): 1038	Hits (normalized): 603.040736813336	Hits (normalized) per kb: 99.1841186951822	Normalized hits with 1T: 88.6%	Normalized hits with 10A: 11.3%	Normalized hits 25-32 nt: 99.8%	Normalized hits on the main strand(s): 89.2%	Predicted directionality: mono:plus
Cluster 178	Location: CM035919.1	Coordinates: 68792879-68798852	Size [bp]: 5974	Hits (absolute): 1214	Hits (normalized): 197.28863758814	Hits (normalized) per kb: 33.0243546903715	Normalized hits with 1T: 97.6%	Normalized hits with 10A: 4.2%	Normalized hits 25-32 nt: 99.6%	Normalized hits on the main strand(s): 100%	Predicted directionality: mono:plus
Cluster 179	Location: CM035919.1	Coordinates: 82095345-82101766	Size [bp]: 6422	Hits (absolute): 1666	Hits (normalized): 495.78897318072	Hits (normalized) per kb: 77.2018841445256	Normalized hits with 1T: 96.1%	Normalized hits with 10A: 18.6%	Normalized hits 25-32 nt: 99.4%	Normalized hits on the main strand(s): 96.8%	Predicted directionality: mono:minus
Cluster 180	Location: CM035919.1	Coordinates: 83819594-83824694	Size [bp]: 5101	Hits (absolute): 665	Hits (normalized): 200.416578234281	Hits (normalized) per kb: 39.2894807414829	Normalized hits with 1T: 99.1%	Normalized hits with 10A: 2.2%	Normalized hits 25-32 nt: 99.9%	Normalized hits on the main strand(s): 99.8%	Predicted directionality: mono:plus
Cluster 181	Location: CM035919.1	Coordinates: 94346016-94351903	Size [bp]: 5888	Hits (absolute): 1383	Hits (normalized): 129.19272337198	Hits (normalized) per kb: 21.9419258352542	Normalized hits with 1T: 85.3%	Normalized hits with 10A: 31.1%	Normalized hits 25-32 nt: 98.5%	Normalized hits on the main strand(s): 95.8%	Predicted directionality: mono:plus
Cluster 182	Location: CM035919.1	Coordinates: 97595719-97601964	Size [bp]: 6246	Hits (absolute): 1541	Hits (normalized): 491.57299660229	Hits (normalized) per kb: 78.7023555097078	Normalized hits with 1T: 98.9%	Normalized hits with 10A: 8.7%	Normalized hits 25-32 nt: 99.8%	Normalized hits on the main strand(s): 99.1%	Predicted directionality: mono:plus
Cluster 183	Location: CM035919.1	Coordinates: 101035291-101041951	Size [bp]: 6661	Hits (absolute): 1482	Hits (normalized): 226.788354136742	Hits (normalized) per kb: 34.0468798586398	Normalized hits with 1T: 97.2%	Normalized hits with 10A: 39.5%	Normalized hits 25-32 nt: 99.7%	Normalized hits on the main strand(s): 98%	Predicted directionality: mono:minus
Cluster 184	Location: CM035919.1	Coordinates: 124133020-124141826	Size [bp]: 8807	Hits (absolute): 1671	Hits (normalized): 201.402152151813	Hits (normalized) per kb: 22.8682036626638	Normalized hits with 1T: 95.4%	Normalized hits with 10A: 17.8%	Normalized hits 25-32 nt: 99.7%	Normalized hits on the main strand(s): 99.9%	Predicted directionality: mono:plus
Cluster 185	Location: CM035919.1	Coordinates: 124161190-124166972	Size [bp]: 5783	Hits (absolute): 1385	Hits (normalized): 123.340859585511	Hits (normalized) per kb: 21.3282462089242	Normalized hits with 1T: 87.3%	Normalized hits with 10A: 19.4%	Normalized hits 25-32 nt: 99.7%	Normalized hits on the main strand(s): 99.8%	Predicted directionality: mono:plus
Cluster 186	Location: CM035919.1	Coordinates: 124191109-124202987	Size [bp]: 11879	Hits (absolute): 3220	Hits (normalized): 376.621763209126	Hits (normalized) per kb: 31.7048612310775	Normalized hits with 1T: 92.8%	Normalized hits with 10A: 38.6%	Normalized hits 25-32 nt: 99.7%	Normalized hits on the main strand(s): 99.5%	Predicted directionality: mono:plus
Cluster 187	Location: CM035919.1	Coordinates: 124596124-124603924	Size [bp]: 7801	Hits (absolute): 597	Hits (normalized): 153.447328664379	Hits (normalized) per kb: 19.6698304895123	Normalized hits with 1T: 85.2%	Normalized hits with 10A: 16.2%	Normalized hits 25-32 nt: 99.8%	Normalized hits on the main strand(s): 95.6%	Predicted directionality: mono:minus
Cluster 188	Location: CM035919.1	Coordinates: 127316004-127323794	Size [bp]: 7791	Hits (absolute): 1411	Hits (normalized): 171.714929132424	Hits (normalized) per kb: 22.0398184298028	Normalized hits with 1T: 96.4%	Normalized hits with 10A: 51.3%	Normalized hits 25-32 nt: 99.7%	Normalized hits on the main strand(s): 97.2%	Predicted directionality: mono:plus
Cluster 189	Location: CM035920.1	Coordinates: 6673018-6677504	Size [bp]: 4487	Hits (absolute): 754	Hits (normalized): 192.377457392043	Hits (normalized) per kb: 42.8744885317591	Normalized hits with 1T: 91.4%	Normalized hits with 10A: 56.2%	Normalized hits 25-32 nt: 99.4%	Normalized hits on the main strand(s): 89.1%	Predicted directionality: mono:plus
Cluster 190	Location: CM035920.1	Coordinates: 9308218-9315900	Size [bp]: 7683	Hits (absolute): 1209	Hits (normalized): 289.533748891569	Hits (normalized) per kb: 37.6853583938375	Normalized hits with 1T: 96.7%	Normalized hits with 10A: 6.3%	Normalized hits 25-32 nt: 99.8%	Normalized hits on the main strand(s): 96.9%	Predicted directionality: mono:plus
Cluster 191	Location: CM035920.1	Coordinates: 10306033-10320684	Size [bp]: 14652	Hits (absolute): 3476	Hits (normalized): 681.115871796086	Hits (normalized) per kb: 46.4858203810737	Normalized hits with 1T: 87.8%	Normalized hits with 10A: 19.9%	Normalized hits 25-32 nt: 99.8%	Normalized hits on the main strand(s): 100%	Predicted directionality: mono:minus
Cluster 192	Location: CM035920.1	Coordinates: 10351056-10356970	Size [bp]: 5915	Hits (absolute): 1177	Hits (normalized): 158.340445344848	Hits (normalized) per kb: 26.7691001613994	Normalized hits with 1T: 95.6%	Normalized hits with 10A: 10.6%	Normalized hits 25-32 nt: 99.7%	Normalized hits on the main strand(s): 100%	Predicted directionality: mono:minus
Cluster 193	Location: CM035920.1	Coordinates: 10369116-10376851	Size [bp]: 7736	Hits (absolute): 1007	Hits (normalized): 221.886123445868	Hits (normalized) per kb: 28.6825302027446	Normalized hits with 1T: 97.7%	Normalized hits with 10A: 23.4%	Normalized hits 25-32 nt: 99.8%	Normalized hits on the main strand(s): 100%	Predicted directionality: mono:minus
Cluster 194	Location: CM035920.1	Coordinates: 10393167-10412696	Size [bp]: 19530	Hits (absolute): 6780	Hits (normalized): 3494.72129782051	Hits (normalized) per kb: 178.941081820111	Normalized hits with 1T: 94.5%	Normalized hits with 10A: 65.1%	Normalized hits 25-32 nt: 99.8%	Normalized hits on the main strand(s): 97%	Predicted directionality: mono:minus
Cluster 195	Location: CM035920.1	Coordinates: 11187066-11211019	Size [bp]: 23954	Hits (absolute): 6123	Hits (normalized): 3351.13818042426	Hits (normalized) per kb: 139.899211758957	Normalized hits with 1T: 97.6%	Normalized hits with 10A: 63.9%	Normalized hits 25-32 nt: 99.9%	Normalized hits on the main strand(s): 99.8%	Predicted directionality: mono:plus
Cluster 196	Location: CM035920.1	Coordinates: 11214057-11223964	Size [bp]: 9908	Hits (absolute): 3124	Hits (normalized): 601.55483134011	Hits (normalized) per kb: 60.7139742912658	Normalized hits with 1T: 92.3%	Normalized hits with 10A: 21.7%	Normalized hits 25-32 nt: 99.8%	Normalized hits on the main strand(s): 100%	Predicted directionality: mono:plus
Cluster 197	Location: CM035920.1	Coordinates: 11227059-11246865	Size [bp]: 19807	Hits (absolute): 7152	Hits (normalized): 1453.20073737158	Hits (normalized) per kb: 73.3684430470756	Normalized hits with 1T: 96.1%	Normalized hits with 10A: 13.2%	Normalized hits 25-32 nt: 99.6%	Normalized hits on the main strand(s): 100%	Predicted directionality: mono:plus
Cluster 198	Location: CM035920.1	Coordinates: 11252013-11279930	Size [bp]: 27918	Hits (absolute): 6882	Hits (normalized): 1371.47373965377	Hits (normalized) per kb: 49.1248072996616	Normalized hits with 1T: 88.8%	Normalized hits with 10A: 21.2%	Normalized hits 25-32 nt: 99.8%	Normalized hits on the main strand(s): 100%	Predicted directionality: mono:plus
Cluster 199	Location: CM035920.1	Coordinates: 19734770-19736654	Size [bp]: 1885	Hits (absolute): 115	Hits (normalized): 160.123193604929	Hits (normalized) per kb: 84.9460932628507	Normalized hits with 1T: 0.2%	Normalized hits with 10A: 98.7%	Normalized hits 25-32 nt: 99.9%	Normalized hits on the main strand(s): 100%	Predicted directionality: mono:plus
Cluster 200	Location: CM035920.1	Coordinates: 20024376-20032602	Size [bp]: 8227	Hits (absolute): 365	Hits (normalized): 311.650178986465	Hits (normalized) per kb: 37.8811435829348	Normalized hits with 1T: 99.6%	Normalized hits with 10A: 0.5%	Normalized hits 25-32 nt: 99.9%	Normalized hits on the main strand(s): 99.4%	Predicted directionality: mono:plus
Cluster 201	Location: CM035920.1	Coordinates: 22247237-22253912	Size [bp]: 6676	Hits (absolute): 2178	Hits (normalized): 173.040797375568	Hits (normalized) per kb: 25.9201492574147	Normalized hits with 1T: 94.9%	Normalized hits with 10A: 21.5%	Normalized hits 25-32 nt: 99.8%	Normalized hits on the main strand(s): 96.7%	Predicted directionality: mono:plus
Cluster 202	Location: CM035920.1	Coordinates: 33103003-33113014	Size [bp]: 10012	Hits (absolute): 2299	Hits (normalized): 458.462573814236	Hits (normalized) per kb: 45.7915233239389	Normalized hits with 1T: 97.7%	Normalized hits with 10A: 12.1%	Normalized hits 25-32 nt: 99.7%	Normalized hits on the main strand(s): 99.7%	Predicted directionality: mono:plus
Cluster 203	Location: CM035920.1	Coordinates: 33114615-33136884	Size [bp]: 22270	Hits (absolute): 5705	Hits (normalized): 1195.24751973298	Hits (normalized) per kb: 53.6706432448351	Normalized hits with 1T: 96.6%	Normalized hits with 10A: 16.4%	Normalized hits 25-32 nt: 99.7%	Normalized hits on the main strand(s): 99.8%	Predicted directionality: mono:plus
Cluster 204	Location: CM035920.1	Coordinates: 33168002-33186025	Size [bp]: 18024	Hits (absolute): 3914	Hits (normalized): 611.027668325993	Hits (normalized) per kb: 33.9004522802394	Normalized hits with 1T: 95.6%	Normalized hits with 10A: 12.2%	Normalized hits 25-32 nt: 99.4%	Normalized hits on the main strand(s): 99.9%	Predicted directionality: mono:plus
Cluster 205	Location: CM035920.1	Coordinates: 35220089-35229003	Size [bp]: 8915	Hits (absolute): 1154	Hits (normalized): 214.476383363293	Hits (normalized) per kb: 24.0577220804562	Normalized hits with 1T: 97.7%	Normalized hits with 10A: 13.5%	Normalized hits 25-32 nt: 99.1%	Normalized hits on the main strand(s): 99.7%	Predicted directionality: mono:minus
Cluster 206	Location: CM035920.1	Coordinates: 35248044-35262940	Size [bp]: 14897	Hits (absolute): 3645	Hits (normalized): 423.677722036401	Hits (normalized) per kb: 28.4406779103304	Normalized hits with 1T: 85.7%	Normalized hits with 10A: 21.4%	Normalized hits 25-32 nt: 99.1%	Normalized hits on the main strand(s): 99.8%	Predicted directionality: mono:minus
Cluster 207	Location: CM035920.1	Coordinates: 35267359-35275866	Size [bp]: 8508	Hits (absolute): 1782	Hits (normalized): 247.495973914136	Hits (normalized) per kb: 29.0897304909931	Normalized hits with 1T: 93%	Normalized hits with 10A: 13.2%	Normalized hits 25-32 nt: 99.6%	Normalized hits on the main strand(s): 100%	Predicted directionality: mono:minus
Cluster 208	Location: CM035920.1	Coordinates: 35277091-35294940	Size [bp]: 17850	Hits (absolute): 5638	Hits (normalized): 1199.16198726729	Hits (normalized) per kb: 67.1798212925442	Normalized hits with 1T: 94.6%	Normalized hits with 10A: 9.3%	Normalized hits 25-32 nt: 99.6%	Normalized hits on the main strand(s): 99.7%	Predicted directionality: mono:minus
Cluster 209	Location: CM035920.1	Coordinates: 35333007-35344000	Size [bp]: 10994	Hits (absolute): 1699	Hits (normalized): 679.109849449779	Hits (normalized) per kb: 61.7710497870219	Normalized hits with 1T: 99%	Normalized hits with 10A: 3.3%	Normalized hits 25-32 nt: 99.9%	Normalized hits on the main strand(s): 99.9%	Predicted directionality: mono:minus
Cluster 210	Location: CM035920.1	Coordinates: 43855365-43862941	Size [bp]: 7577	Hits (absolute): 1249	Hits (normalized): 299.8079336757	Hits (normalized) per kb: 39.5683512419197	Normalized hits with 1T: 96.8%	Normalized hits with 10A: 6.4%	Normalized hits 25-32 nt: 99.8%	Normalized hits on the main strand(s): 97.1%	Predicted directionality: mono:plus
Cluster 211	Location: CM035920.1	Coordinates: 44056633-44062809	Size [bp]: 6177	Hits (absolute): 1124	Hits (normalized): 294.136345154495	Hits (normalized) per kb: 47.6177549197198	Normalized hits with 1T: 97.1%	Normalized hits with 10A: 6%	Normalized hits 25-32 nt: 99.9%	Normalized hits on the main strand(s): 97.5%	Predicted directionality: mono:plus
Cluster 212	Location: CM035920.1	Coordinates: 45136005-45143898	Size [bp]: 7894	Hits (absolute): 2190	Hits (normalized): 148.855508049822	Hits (normalized) per kb: 18.8570751667053	Normalized hits with 1T: 91.5%	Normalized hits with 10A: 42.7%	Normalized hits 25-32 nt: 98.5%	Normalized hits on the main strand(s): 99.3%	Predicted directionality: mono:minus
Cluster 213	Location: CM035920.1	Coordinates: 46213076-46218916	Size [bp]: 5841	Hits (absolute): 573	Hits (normalized): 192.360662093958	Hits (normalized) per kb: 32.9330431105824	Normalized hits with 1T: 94.2%	Normalized hits with 10A: 66.2%	Normalized hits 25-32 nt: 99.4%	Normalized hits on the main strand(s): 99.6%	Predicted directionality: mono:plus
Cluster 214	Location: CM035920.1	Coordinates: 51998276-52006917	Size [bp]: 8642	Hits (absolute): 1345	Hits (normalized): 222.425649516566	Hits (normalized) per kb: 25.7375260978367	Normalized hits with 1T: 89.6%	Normalized hits with 10A: 15.6%	Normalized hits 25-32 nt: 99.6%	Normalized hits on the main strand(s): 99.8%	Predicted directionality: mono:plus
Cluster 215	Location: CM035920.1	Coordinates: 64656810-64664007	Size [bp]: 7198	Hits (absolute): 1197	Hits (normalized): 244.575945278295	Hits (normalized) per kb: 33.9786018305093	Normalized hits with 1T: 96.2%	Normalized hits with 10A: 8.4%	Normalized hits 25-32 nt: 99.8%	Normalized hits on the main strand(s): 95.8%	Predicted directionality: mono:plus
Cluster 216	Location: CM035920.1	Coordinates: 64749001-64754046	Size [bp]: 5046	Hits (absolute): 940	Hits (normalized): 219.512685127062	Hits (normalized) per kb: 43.5021528144532	Normalized hits with 1T: 98%	Normalized hits with 10A: 5.6%	Normalized hits 25-32 nt: 99.8%	Normalized hits on the main strand(s): 97.9%	Predicted directionality: mono:minus
Cluster 217	Location: CM035920.1	Coordinates: 78276076-78281935	Size [bp]: 5860	Hits (absolute): 728	Hits (normalized): 129.172373150745	Hits (normalized) per kb: 22.0431089371826	Normalized hits with 1T: 81.2%	Normalized hits with 10A: 66.2%	Normalized hits 25-32 nt: 99.6%	Normalized hits on the main strand(s): 99.4%	Predicted directionality: mono:minus
Cluster 218	Location: CM035920.1	Coordinates: 80057250-80062850	Size [bp]: 5601	Hits (absolute): 353	Hits (normalized): 124.098743739124	Hits (normalized) per kb: 22.1566314417852	Normalized hits with 1T: 2%	Normalized hits with 10A: 98.8%	Normalized hits 25-32 nt: 99.1%	Normalized hits on the main strand(s): 99.6%	Predicted directionality: mono:minus
Cluster 219	Location: CM035920.1	Coordinates: 91581002-91590516	Size [bp]: 9515	Hits (absolute): 1832	Hits (normalized): 449.383773951853	Hits (normalized) per kb: 47.2286524220602	Normalized hits with 1T: 96.5%	Normalized hits with 10A: 17.6%	Normalized hits 25-32 nt: 99.6%	Normalized hits on the main strand(s): 96%	Predicted directionality: mono:minus
Cluster 220	Location: CM035920.1	Coordinates: 96649040-96680001	Size [bp]: 30962	Hits (absolute): 12412	Hits (normalized): 3450.08352678643	Hits (normalized) per kb: 111.429741909054	Normalized hits with 1T: 95.6%	Normalized hits with 10A: 9.9%	Normalized hits 25-32 nt: 99.8%	Normalized hits on the main strand(s): 100%	Predicted directionality: mono:minus
Cluster 221	Location: CM035920.1	Coordinates: 97533002-97544727	Size [bp]: 11726	Hits (absolute): 2500	Hits (normalized): 465.204392607985	Hits (normalized) per kb: 39.6728248512279	Normalized hits with 1T: 95.9%	Normalized hits with 10A: 4.4%	Normalized hits 25-32 nt: 99.8%	Normalized hits on the main strand(s): 100%	Predicted directionality: mono:plus
Cluster 222	Location: CM035920.1	Coordinates: 110773625-110779205	Size [bp]: 5581	Hits (absolute): 1365	Hits (normalized): 272.452820904012	Hits (normalized) per kb: 48.8179674864966	Normalized hits with 1T: 98.3%	Normalized hits with 10A: 4.7%	Normalized hits 25-32 nt: 99.8%	Normalized hits on the main strand(s): 98.7%	Predicted directionality: mono:minus
Cluster 223	Location: CM035920.1	Coordinates: 116768125-116772332	Size [bp]: 4208	Hits (absolute): 1198	Hits (normalized): 270.622635698853	Hits (normalized) per kb: 64.3113214842163	Normalized hits with 1T: 96.8%	Normalized hits with 10A: 6.1%	Normalized hits 25-32 nt: 99.8%	Normalized hits on the main strand(s): 97.1%	Predicted directionality: mono:minus
Cluster 224	Location: CM035921.1	Coordinates: 3035454-3039954	Size [bp]: 4501	Hits (absolute): 570	Hits (normalized): 192.560355885144	Hits (normalized) per kb: 42.7815316982802	Normalized hits with 1T: 35.8%	Normalized hits with 10A: 93.9%	Normalized hits 25-32 nt: 96.5%	Normalized hits on the main strand(s): 99.8%	Predicted directionality: mono:minus
Cluster 225	Location: CM035921.1	Coordinates: 3079336-3084907	Size [bp]: 5572	Hits (absolute): 590	Hits (normalized): 219.447891832551	Hits (normalized) per kb: 39.3840828286518	Normalized hits with 1T: 44%	Normalized hits with 10A: 99.1%	Normalized hits 25-32 nt: 98.3%	Normalized hits on the main strand(s): 99.5%	Predicted directionality: mono:minus
Cluster 226	Location: CM035921.1	Coordinates: 3163259-3169077	Size [bp]: 5819	Hits (absolute): 394	Hits (normalized): 148.448863367941	Hits (normalized) per kb: 25.5113037154764	Normalized hits with 1T: 39.6%	Normalized hits with 10A: 98.3%	Normalized hits 25-32 nt: 99%	Normalized hits on the main strand(s): 99.9%	Predicted directionality: mono:minus
Cluster 227	Location: CM035921.1	Coordinates: 3278484-3280911	Size [bp]: 2428	Hits (absolute): 714	Hits (normalized): 265.412729444604	Hits (normalized) per kb: 109.313123037007	Normalized hits with 1T: 39%	Normalized hits with 10A: 95.5%	Normalized hits 25-32 nt: 98.5%	Normalized hits on the main strand(s): 99.9%	Predicted directionality: mono:minus
Cluster 228	Location: CM035921.1	Coordinates: 9145795-9151963	Size [bp]: 6169	Hits (absolute): 1031	Hits (normalized): 165.595202323434	Hits (normalized) per kb: 26.8431365774446	Normalized hits with 1T: 97%	Normalized hits with 10A: 6.6%	Normalized hits 25-32 nt: 98.5%	Normalized hits on the main strand(s): 100%	Predicted directionality: mono:minus
Cluster 229	Location: CM035921.1	Coordinates: 26965204-26969908	Size [bp]: 4705	Hits (absolute): 450	Hits (normalized): 170.035695554786	Hits (normalized) per kb: 36.1396425521833	Normalized hits with 1T: 98.6%	Normalized hits with 10A: 54%	Normalized hits 25-32 nt: 99.6%	Normalized hits on the main strand(s): 99.8%	Predicted directionality: mono:plus
Cluster 230	Location: CM035921.1	Coordinates: 63094086-63097627	Size [bp]: 3542	Hits (absolute): 783	Hits (normalized): 465.954176040338	Hits (normalized) per kb: 131.551194536442	Normalized hits with 1T: 99%	Normalized hits with 10A: 6.8%	Normalized hits 25-32 nt: 99.9%	Normalized hits on the main strand(s): 98.9%	Predicted directionality: mono:minus
Cluster 231	Location: CM035921.1	Coordinates: 75332047-75342003	Size [bp]: 9957	Hits (absolute): 1545	Hits (normalized): 270.307643637433	Hits (normalized) per kb: 27.1475085100747	Normalized hits with 1T: 97.3%	Normalized hits with 10A: 15.6%	Normalized hits 25-32 nt: 99.6%	Normalized hits on the main strand(s): 98.9%	Predicted directionality: mono:plus
Cluster 232	Location: CM035921.1	Coordinates: 80071148-80077764	Size [bp]: 6617	Hits (absolute): 277	Hits (normalized): 217.387623184304	Hits (normalized) per kb: 32.8532483066226	Normalized hits with 1T: 97.4%	Normalized hits with 10A: 99%	Normalized hits 25-32 nt: 99.9%	Normalized hits on the main strand(s): 99.5%	Predicted directionality: mono:plus
Cluster 233	Location: CM035921.1	Coordinates: 85326383-85335020	Size [bp]: 8638	Hits (absolute): 411	Hits (normalized): 150.499004337053	Hits (normalized) per kb: 17.4232365759638	Normalized hits with 1T: 1.9%	Normalized hits with 10A: 98.4%	Normalized hits 25-32 nt: 96%	Normalized hits on the main strand(s): 99.6%	Predicted directionality: mono:minus
Cluster 234	Location: CM035921.1	Coordinates: 90437004-90442896	Size [bp]: 5893	Hits (absolute): 2678	Hits (normalized): 153.391946377712	Hits (normalized) per kb: 26.0295586277926	Normalized hits with 1T: 89.6%	Normalized hits with 10A: 28.7%	Normalized hits 25-32 nt: 99.6%	Normalized hits on the main strand(s): 99.5%	Predicted directionality: mono:minus
Cluster 235	Location: CM035921.1	Coordinates: 92495248-92503996	Size [bp]: 8749	Hits (absolute): 959	Hits (normalized): 311.952506043373	Hits (normalized) per kb: 35.6559379673549	Normalized hits with 1T: 97.7%	Normalized hits with 10A: 7.1%	Normalized hits 25-32 nt: 99.8%	Normalized hits on the main strand(s): 99.8%	Predicted directionality: mono:plus
Cluster 236	Location: CM035921.1	Coordinates: 96071530-96077864	Size [bp]: 6335	Hits (absolute): 1715	Hits (normalized): 583.54624801267	Hits (normalized) per kb: 92.1144635897132	Normalized hits with 1T: 14.9%	Normalized hits with 10A: 85.2%	Normalized hits 25-32 nt: 99%	Normalized hits on the main strand(s): 93.8%	Predicted directionality: bi:plus-minus (split between 96076504 and 96076504)
Cluster 237	Location: CM035921.1	Coordinates: 97191044-97199856	Size [bp]: 8813	Hits (absolute): 704	Hits (normalized): 167.978821554445	Hits (normalized) per kb: 19.0602639974071	Normalized hits with 1T: 98.3%	Normalized hits with 10A: 9.1%	Normalized hits 25-32 nt: 99.1%	Normalized hits on the main strand(s): 100%	Predicted directionality: mono:plus
Cluster 238	Location: CM035922.1	Coordinates: 4714008-4723011	Size [bp]: 9004	Hits (absolute): 2706	Hits (normalized): 292.644654210048	Hits (normalized) per kb: 32.5019866438305	Normalized hits with 1T: 89.5%	Normalized hits with 10A: 23.6%	Normalized hits 25-32 nt: 99.3%	Normalized hits on the main strand(s): 99%	Predicted directionality: mono:plus
Cluster 239	Location: CM035922.1	Coordinates: 17174000-17179028	Size [bp]: 5029	Hits (absolute): 898	Hits (normalized): 125.973020151711	Hits (normalized) per kb: 25.0489874286166	Normalized hits with 1T: 86.2%	Normalized hits with 10A: 18.9%	Normalized hits 25-32 nt: 99.3%	Normalized hits on the main strand(s): 99.6%	Predicted directionality: mono:plus
Cluster 240	Location: CM035922.1	Coordinates: 19365143-19373026	Size [bp]: 7884	Hits (absolute): 1284	Hits (normalized): 232.753061411811	Hits (normalized) per kb: 29.5224322114348	Normalized hits with 1T: 96.7%	Normalized hits with 10A: 48.1%	Normalized hits 25-32 nt: 99.8%	Normalized hits on the main strand(s): 99.1%	Predicted directionality: mono:minus
Cluster 241	Location: CM035922.1	Coordinates: 19418469-19427024	Size [bp]: 8556	Hits (absolute): 1519	Hits (normalized): 232.890217692076	Hits (normalized) per kb: 27.21989967243	Normalized hits with 1T: 96.7%	Normalized hits with 10A: 44%	Normalized hits 25-32 nt: 99.7%	Normalized hits on the main strand(s): 100%	Predicted directionality: mono:minus
Cluster 242	Location: CM035922.1	Coordinates: 21362736-21374864	Size [bp]: 12129	Hits (absolute): 2654	Hits (normalized): 444.986422023792	Hits (normalized) per kb: 36.6875120309176	Normalized hits with 1T: 96%	Normalized hits with 10A: 22.5%	Normalized hits 25-32 nt: 99.7%	Normalized hits on the main strand(s): 97.5%	Predicted directionality: mono:plus
Cluster 243	Location: CM035922.1	Coordinates: 21381112-21387899	Size [bp]: 6788	Hits (absolute): 995	Hits (normalized): 147.905185951289	Hits (normalized) per kb: 21.7889172420941	Normalized hits with 1T: 93.9%	Normalized hits with 10A: 30.5%	Normalized hits 25-32 nt: 99.6%	Normalized hits on the main strand(s): 100%	Predicted directionality: mono:plus
Cluster 244	Location: CM035922.1	Coordinates: 21397098-21403945	Size [bp]: 6848	Hits (absolute): 711	Hits (normalized): 121.086408581196	Hits (normalized) per kb: 17.6823640321219	Normalized hits with 1T: 97.8%	Normalized hits with 10A: 53.5%	Normalized hits 25-32 nt: 98.9%	Normalized hits on the main strand(s): 99.9%	Predicted directionality: mono:plus
Cluster 245	Location: CM035922.1	Coordinates: 21520093-21532884	Size [bp]: 12792	Hits (absolute): 2510	Hits (normalized): 338.384921865428	Hits (normalized) per kb: 26.45321145294	Normalized hits with 1T: 95.4%	Normalized hits with 10A: 15%	Normalized hits 25-32 nt: 99.5%	Normalized hits on the main strand(s): 99.9%	Predicted directionality: mono:minus
Cluster 246	Location: CM035922.1	Coordinates: 28769063-28777571	Size [bp]: 8509	Hits (absolute): 887	Hits (normalized): 177.145587753094	Hits (normalized) per kb: 20.8182175650574	Normalized hits with 1T: 97.1%	Normalized hits with 10A: 12.2%	Normalized hits 25-32 nt: 99.1%	Normalized hits on the main strand(s): 99.9%	Predicted directionality: mono:minus
Cluster 247	Location: CM035922.1	Coordinates: 28799138-28814969	Size [bp]: 15832	Hits (absolute): 2423	Hits (normalized): 361.88633150379	Hits (normalized) per kb: 22.8575095136795	Normalized hits with 1T: 92.8%	Normalized hits with 10A: 26.6%	Normalized hits 25-32 nt: 99.6%	Normalized hits on the main strand(s): 83.8%	Predicted directionality: mono:minus
Cluster 248	Location: CM035922.1	Coordinates: 36495435-36500025	Size [bp]: 4591	Hits (absolute): 1100	Hits (normalized): 141.79303693341	Hits (normalized) per kb: 30.884702266666	Normalized hits with 1T: 92.2%	Normalized hits with 10A: 17%	Normalized hits 25-32 nt: 99.6%	Normalized hits on the main strand(s): 99.7%	Predicted directionality: mono:plus
Cluster 249	Location: CM035922.1	Coordinates: 37577243-37582016	Size [bp]: 4774	Hits (absolute): 803	Hits (normalized): 160.306068213028	Hits (normalized) per kb: 33.5788051838653	Normalized hits with 1T: 94.9%	Normalized hits with 10A: 39.5%	Normalized hits 25-32 nt: 99.6%	Normalized hits on the main strand(s): 96.8%	Predicted directionality: mono:minus
Cluster 250	Location: CM035922.1	Coordinates: 41814029-41820990	Size [bp]: 6962	Hits (absolute): 1433	Hits (normalized): 1265.96309002361	Hits (normalized) per kb: 181.839196194857	Normalized hits with 1T: 91.8%	Normalized hits with 10A: 9.3%	Normalized hits 25-32 nt: 99.8%	Normalized hits on the main strand(s): 95%	Predicted directionality: mono:minus
Cluster 251	Location: CM035922.1	Coordinates: 44337089-44343449	Size [bp]: 6361	Hits (absolute): 1191	Hits (normalized): 162.517450975069	Hits (normalized) per kb: 25.5491445503439	Normalized hits with 1T: 86.8%	Normalized hits with 10A: 29.5%	Normalized hits 25-32 nt: 99.5%	Normalized hits on the main strand(s): 99.8%	Predicted directionality: mono:minus
Cluster 252	Location: CM035922.1	Coordinates: 58481132-58497019	Size [bp]: 15888	Hits (absolute): 2477	Hits (normalized): 1010.62713296874	Hits (normalized) per kb: 63.609620785477	Normalized hits with 1T: 91.6%	Normalized hits with 10A: 10.7%	Normalized hits 25-32 nt: 99.5%	Normalized hits on the main strand(s): 100%	Predicted directionality: mono:plus
Cluster 253	Location: CM035922.1	Coordinates: 58579140-58587700	Size [bp]: 8561	Hits (absolute): 1587	Hits (normalized): 230.237489075706	Hits (normalized) per kb: 26.8941394418312	Normalized hits with 1T: 97.2%	Normalized hits with 10A: 39.3%	Normalized hits 25-32 nt: 99.7%	Normalized hits on the main strand(s): 97.8%	Predicted directionality: mono:plus
Cluster 254	Location: CM035922.1	Coordinates: 72209083-72215909	Size [bp]: 6827	Hits (absolute): 1409	Hits (normalized): 171.865807360268	Hits (normalized) per kb: 25.1740267090484	Normalized hits with 1T: 96.5%	Normalized hits with 10A: 51.5%	Normalized hits 25-32 nt: 99.7%	Normalized hits on the main strand(s): 97.2%	Predicted directionality: mono:plus
Cluster 255	Location: CM035922.1	Coordinates: 73234276-73241606	Size [bp]: 7331	Hits (absolute): 2216	Hits (normalized): 178.696786789988	Hits (normalized) per kb: 24.3752560426055	Normalized hits with 1T: 95.3%	Normalized hits with 10A: 24.3%	Normalized hits 25-32 nt: 99.8%	Normalized hits on the main strand(s): 97.3%	Predicted directionality: mono:plus
Cluster 256	Location: CM035922.1	Coordinates: 82585281-82641023	Size [bp]: 55743	Hits (absolute): 10968	Hits (normalized): 2800.41659090628	Hits (normalized) per kb: 50.237821420874	Normalized hits with 1T: 95.9%	Normalized hits with 10A: 20.3%	Normalized hits 25-32 nt: 99.4%	Normalized hits on the main strand(s): 99.9%	Predicted directionality: mono:minus
Cluster 257	Location: CM035922.1	Coordinates: 82650015-82679027	Size [bp]: 29013	Hits (absolute): 6079	Hits (normalized): 4380.26251504475	Hits (normalized) per kb: 150.97588222616	Normalized hits with 1T: 97.6%	Normalized hits with 10A: 13.9%	Normalized hits 25-32 nt: 99.8%	Normalized hits on the main strand(s): 99.9%	Predicted directionality: mono:minus
Cluster 258	Location: CM035922.1	Coordinates: 90741755-90744530	Size [bp]: 2776	Hits (absolute): 420	Hits (normalized): 299.918441091079	Hits (normalized) per kb: 108.03969668103	Normalized hits with 1T: 99.1%	Normalized hits with 10A: 3.1%	Normalized hits 25-32 nt: 99.8%	Normalized hits on the main strand(s): 99.8%	Predicted directionality: mono:minus
Cluster 259	Location: CM035922.1	Coordinates: 97650039-97655862	Size [bp]: 5824	Hits (absolute): 1571	Hits (normalized): 181.492263116178	Hits (normalized) per kb: 31.1627501402579	Normalized hits with 1T: 91%	Normalized hits with 10A: 22.1%	Normalized hits 25-32 nt: 99.6%	Normalized hits on the main strand(s): 90.6%	Predicted directionality: mono:minus
Cluster 260	Location: CM035922.1	Coordinates: 99199600-99204676	Size [bp]: 5077	Hits (absolute): 1200	Hits (normalized): 198.494801735499	Hits (normalized) per kb: 39.0969860597655	Normalized hits with 1T: 96.1%	Normalized hits with 10A: 3.1%	Normalized hits 25-32 nt: 97.2%	Normalized hits on the main strand(s): 100%	Predicted directionality: mono:plus
Cluster 261	Location: CM035922.1	Coordinates: 99271105-99279087	Size [bp]: 7983	Hits (absolute): 1478	Hits (normalized): 661.510226152655	Hits (normalized) per kb: 82.8648473451364	Normalized hits with 1T: 98.1%	Normalized hits with 10A: 0.7%	Normalized hits 25-32 nt: 98.4%	Normalized hits on the main strand(s): 99.9%	Predicted directionality: mono:plus
Cluster 262	Location: CM035922.1	Coordinates: 100229028-100237771	Size [bp]: 8744	Hits (absolute): 983	Hits (normalized): 209.480324316861	Hits (normalized) per kb: 23.9573616053727	Normalized hits with 1T: 97.6%	Normalized hits with 10A: 7%	Normalized hits 25-32 nt: 99.6%	Normalized hits on the main strand(s): 98.5%	Predicted directionality: mono:minus
Cluster 263	Location: CM035922.1	Coordinates: 100830329-100839022	Size [bp]: 8694	Hits (absolute): 1246	Hits (normalized): 230.874936143716	Hits (normalized) per kb: 26.5560398085583	Normalized hits with 1T: 97.1%	Normalized hits with 10A: 9.3%	Normalized hits 25-32 nt: 99.5%	Normalized hits on the main strand(s): 98.3%	Predicted directionality: mono:plus
Cluster 264	Location: CM035922.1	Coordinates: 103558032-103564013	Size [bp]: 5982	Hits (absolute): 1045	Hits (normalized): 134.421097439908	Hits (normalized) per kb: 22.4708748965547	Normalized hits with 1T: 94.6%	Normalized hits with 10A: 15.7%	Normalized hits 25-32 nt: 98.7%	Normalized hits on the main strand(s): 99.8%	Predicted directionality: mono:plus
Cluster 265	Location: CM035922.1	Coordinates: 103566520-103575012	Size [bp]: 8493	Hits (absolute): 1921	Hits (normalized): 182.465096565215	Hits (normalized) per kb: 21.484545309464	Normalized hits with 1T: 81.9%	Normalized hits with 10A: 21.7%	Normalized hits 25-32 nt: 99.7%	Normalized hits on the main strand(s): 99.9%	Predicted directionality: mono:plus
Cluster 266	Location: CM035922.1	Coordinates: 105025554-105035018	Size [bp]: 9465	Hits (absolute): 1465	Hits (normalized): 275.158964803245	Hits (normalized) per kb: 29.0708100735593	Normalized hits with 1T: 96.7%	Normalized hits with 10A: 11.4%	Normalized hits 25-32 nt: 99.4%	Normalized hits on the main strand(s): 97.1%	Predicted directionality: mono:plus
Cluster 267	Location: CM035923.1	Coordinates: 1273007-1281869	Size [bp]: 8863	Hits (absolute): 936	Hits (normalized): 161.047588362636	Hits (normalized) per kb: 18.1710043780201	Normalized hits with 1T: 96.3%	Normalized hits with 10A: 14.4%	Normalized hits 25-32 nt: 99.8%	Normalized hits on the main strand(s): 96.6%	Predicted directionality: mono:plus
Cluster 268	Location: CM035923.1	Coordinates: 1283000-1291980	Size [bp]: 8981	Hits (absolute): 1750	Hits (normalized): 218.048108694658	Hits (normalized) per kb: 24.2790087017468	Normalized hits with 1T: 94.4%	Normalized hits with 10A: 18.9%	Normalized hits 25-32 nt: 99.7%	Normalized hits on the main strand(s): 95.1%	Predicted directionality: bi:minus-plus (split between 1285862 and 1285862)
Cluster 269	Location: CM035923.1	Coordinates: 1301308-1309970	Size [bp]: 8663	Hits (absolute): 1932	Hits (normalized): 474.142764384008	Hits (normalized) per kb: 54.7318318748159	Normalized hits with 1T: 94.6%	Normalized hits with 10A: 13.2%	Normalized hits 25-32 nt: 99.6%	Normalized hits on the main strand(s): 99.8%	Predicted directionality: mono:plus
Cluster 270	Location: CM035923.1	Coordinates: 1319026-1326025	Size [bp]: 7000	Hits (absolute): 1974	Hits (normalized): 164.51189965662	Hits (normalized) per kb: 23.5016263332725	Normalized hits with 1T: 88.1%	Normalized hits with 10A: 20%	Normalized hits 25-32 nt: 99.6%	Normalized hits on the main strand(s): 99.9%	Predicted directionality: mono:plus
Cluster 271	Location: CM035923.1	Coordinates: 1328017-1352805	Size [bp]: 24789	Hits (absolute): 6197	Hits (normalized): 765.168765700325	Hits (normalized) per kb: 30.8674271029221	Normalized hits with 1T: 94.5%	Normalized hits with 10A: 11.8%	Normalized hits 25-32 nt: 99.5%	Normalized hits on the main strand(s): 98.3%	Predicted directionality: mono:plus
Cluster 272	Location: CM035923.1	Coordinates: 9558410-9564024	Size [bp]: 5615	Hits (absolute): 760	Hits (normalized): 463.009995157594	Hits (normalized) per kb: 82.4592923105778	Normalized hits with 1T: 99.1%	Normalized hits with 10A: 6.6%	Normalized hits 25-32 nt: 99.9%	Normalized hits on the main strand(s): 99.3%	Predicted directionality: mono:minus
Cluster 273	Location: CM035923.1	Coordinates: 11865677-11881015	Size [bp]: 15339	Hits (absolute): 4109	Hits (normalized): 524.338102774973	Hits (normalized) per kb: 34.1834359149009	Normalized hits with 1T: 96.5%	Normalized hits with 10A: 51%	Normalized hits 25-32 nt: 99.7%	Normalized hits on the main strand(s): 97.4%	Predicted directionality: mono:minus
Cluster 274	Location: CM035923.1	Coordinates: 23405454-23413895	Size [bp]: 8442	Hits (absolute): 602	Hits (normalized): 304.897709523415	Hits (normalized) per kb: 36.1166090005248	Normalized hits with 1T: 99.1%	Normalized hits with 10A: 3.3%	Normalized hits 25-32 nt: 99.8%	Normalized hits on the main strand(s): 98.1%	Predicted directionality: mono:plus
Cluster 275	Location: CM035923.1	Coordinates: 24251007-24259145	Size [bp]: 8139	Hits (absolute): 322	Hits (normalized): 160.902637782469	Hits (normalized) per kb: 19.7693683377508	Normalized hits with 1T: 99.3%	Normalized hits with 10A: 1%	Normalized hits 25-32 nt: 99.8%	Normalized hits on the main strand(s): 99.6%	Predicted directionality: mono:minus
Cluster 276	Location: CM035923.1	Coordinates: 36527472-36535852	Size [bp]: 8381	Hits (absolute): 747	Hits (normalized): 220.470092523105	Hits (normalized) per kb: 26.3059612476946	Normalized hits with 1T: 89.9%	Normalized hits with 10A: 8%	Normalized hits 25-32 nt: 92.5%	Normalized hits on the main strand(s): 100%	Predicted directionality: mono:minus
Cluster 277	Location: CM035923.1	Coordinates: 49544098-49546975	Size [bp]: 2878	Hits (absolute): 337	Hits (normalized): 212.043008263326	Hits (normalized) per kb: 73.6769281139305	Normalized hits with 1T: 14.9%	Normalized hits with 10A: 85.5%	Normalized hits 25-32 nt: 98.1%	Normalized hits on the main strand(s): 99.6%	Predicted directionality: bi:plus-minus (split between 49545791 and 49545795)
Cluster 278	Location: CM035923.1	Coordinates: 53172410-53178222	Size [bp]: 5813	Hits (absolute): 1296	Hits (normalized): 310.498833066428	Hits (normalized) per kb: 53.4148062960568	Normalized hits with 1T: 93.3%	Normalized hits with 10A: 14%	Normalized hits 25-32 nt: 98.6%	Normalized hits on the main strand(s): 96%	Predicted directionality: mono:plus
Cluster 279	Location: CM035923.1	Coordinates: 53863127-53871993	Size [bp]: 8867	Hits (absolute): 333	Hits (normalized): 148.806878218569	Hits (normalized) per kb: 16.7824102637506	Normalized hits with 1T: 1%	Normalized hits with 10A: 99%	Normalized hits 25-32 nt: 96%	Normalized hits on the main strand(s): 99.7%	Predicted directionality: mono:minus
Cluster 280	Location: CM035923.1	Coordinates: 60317017-60322677	Size [bp]: 5661	Hits (absolute): 1505	Hits (normalized): 185.356542927137	Hits (normalized) per kb: 32.7430163093998	Normalized hits with 1T: 81.1%	Normalized hits with 10A: 34%	Normalized hits 25-32 nt: 98.9%	Normalized hits on the main strand(s): 100%	Predicted directionality: mono:minus
Cluster 281	Location: CM035923.1	Coordinates: 61580460-61598913	Size [bp]: 18454	Hits (absolute): 3892	Hits (normalized): 849.565034665304	Hits (normalized) per kb: 46.0366661237329	Normalized hits with 1T: 90.9%	Normalized hits with 10A: 23.8%	Normalized hits 25-32 nt: 98.9%	Normalized hits on the main strand(s): 99.6%	Predicted directionality: mono:plus
Cluster 282	Location: CM035923.1	Coordinates: 61612013-61620525	Size [bp]: 8513	Hits (absolute): 1293	Hits (normalized): 383.694091766036	Hits (normalized) per kb: 45.0717248346109	Normalized hits with 1T: 97.7%	Normalized hits with 10A: 15.8%	Normalized hits 25-32 nt: 99.8%	Normalized hits on the main strand(s): 99.8%	Predicted directionality: mono:plus
Cluster 283	Location: CM035923.1	Coordinates: 67417261-67426853	Size [bp]: 9593	Hits (absolute): 5004	Hits (normalized): 4746.40802031762	Hits (normalized) per kb: 494.77796478828	Normalized hits with 1T: 87.3%	Normalized hits with 10A: 6.7%	Normalized hits 25-32 nt: 99%	Normalized hits on the main strand(s): 100%	Predicted directionality: mono:minus
Cluster 284	Location: CM035923.1	Coordinates: 94704411-94710022	Size [bp]: 5612	Hits (absolute): 1703	Hits (normalized): 132.737094491559	Hits (normalized) per kb: 23.6521670458976	Normalized hits with 1T: 92.9%	Normalized hits with 10A: 41.5%	Normalized hits 25-32 nt: 98.7%	Normalized hits on the main strand(s): 99.5%	Predicted directionality: mono:plus
Cluster 285	Location: CM035923.1	Coordinates: 94749712-94755480	Size [bp]: 5769	Hits (absolute): 1969	Hits (normalized): 256.854185631153	Hits (normalized) per kb: 44.5230327290317	Normalized hits with 1T: 89.5%	Normalized hits with 10A: 57.6%	Normalized hits 25-32 nt: 99.5%	Normalized hits on the main strand(s): 91.9%	Predicted directionality: mono:minus
Cluster 286	Location: CM035923.1	Coordinates: 106085100-106102942	Size [bp]: 17843	Hits (absolute): 2641	Hits (normalized): 545.60517528906	Hits (normalized) per kb: 30.577862453501	Normalized hits with 1T: 80.9%	Normalized hits with 10A: 13.4%	Normalized hits 25-32 nt: 98.3%	Normalized hits on the main strand(s): 100%	Predicted directionality: mono:plus
Cluster 287	Location: CM035923.1	Coordinates: 106107491-106112746	Size [bp]: 5256	Hits (absolute): 1100	Hits (normalized): 215.524251102901	Hits (normalized) per kb: 41.005480340041	Normalized hits with 1T: 98%	Normalized hits with 10A: 39.3%	Normalized hits 25-32 nt: 99.8%	Normalized hits on the main strand(s): 98.7%	Predicted directionality: mono:plus
Cluster 288	Location: CM035924.1	Coordinates: 184226-190854	Size [bp]: 6629	Hits (absolute): 1522	Hits (normalized): 168.796502558778	Hits (normalized) per kb: 25.4635913584695	Normalized hits with 1T: 91.2%	Normalized hits with 10A: 24%	Normalized hits 25-32 nt: 99.3%	Normalized hits on the main strand(s): 99.9%	Predicted directionality: mono:plus
Cluster 289	Location: CM035924.1	Coordinates: 3316209-3330026	Size [bp]: 13818	Hits (absolute): 5521	Hits (normalized): 3470.09631618656	Hits (normalized) per kb: 251.129055344689	Normalized hits with 1T: 94%	Normalized hits with 10A: 20.6%	Normalized hits 25-32 nt: 99.8%	Normalized hits on the main strand(s): 100%	Predicted directionality: mono:plus
Cluster 290	Location: CM035924.1	Coordinates: 3331051-3343965	Size [bp]: 12915	Hits (absolute): 14175	Hits (normalized): 15363.5014040261	Hits (normalized) per kb: 1189.58587319372	Normalized hits with 1T: 92.8%	Normalized hits with 10A: 68.1%	Normalized hits 25-32 nt: 99.8%	Normalized hits on the main strand(s): 100%	Predicted directionality: mono:plus
Cluster 291	Location: CM035924.1	Coordinates: 14886589-14891639	Size [bp]: 5051	Hits (absolute): 1541	Hits (normalized): 314.910104744074	Hits (normalized) per kb: 62.3460659516394	Normalized hits with 1T: 96.9%	Normalized hits with 10A: 6.1%	Normalized hits 25-32 nt: 99.8%	Normalized hits on the main strand(s): 97.1%	Predicted directionality: mono:minus
Cluster 292	Location: CM035924.1	Coordinates: 15187328-15189276	Size [bp]: 1949	Hits (absolute): 649	Hits (normalized): 266.34880757105	Hits (normalized) per kb: 136.658884616713	Normalized hits with 1T: 1.1%	Normalized hits with 10A: 98.9%	Normalized hits 25-32 nt: 99%	Normalized hits on the main strand(s): 99.3%	Predicted directionality: mono:minus
Cluster 293	Location: CM035924.1	Coordinates: 27661007-27668356	Size [bp]: 7350	Hits (absolute): 1973	Hits (normalized): 179.922126896146	Hits (normalized) per kb: 24.4789070250687	Normalized hits with 1T: 95.9%	Normalized hits with 10A: 26.6%	Normalized hits 25-32 nt: 99.8%	Normalized hits on the main strand(s): 93.7%	Predicted directionality: mono:plus
Cluster 294	Location: CM035924.1	Coordinates: 33147294-33148840	Size [bp]: 1547	Hits (absolute): 181	Hits (normalized): 839.478419232697	Hits (normalized) per kb: 542.649088776244	Normalized hits with 1T: 99.9%	Normalized hits with 10A: 2.6%	Normalized hits 25-32 nt: 100%	Normalized hits on the main strand(s): 100%	Predicted directionality: mono:plus
Cluster 295	Location: CM035924.1	Coordinates: 42075327-42084003	Size [bp]: 8677	Hits (absolute): 1351	Hits (normalized): 372.906204218805	Hits (normalized) per kb: 42.9764942605325	Normalized hits with 1T: 98.7%	Normalized hits with 10A: 10.6%	Normalized hits 25-32 nt: 99.8%	Normalized hits on the main strand(s): 99%	Predicted directionality: mono:plus
Cluster 296	Location: CM035924.1	Coordinates: 46497004-46503967	Size [bp]: 6964	Hits (absolute): 404	Hits (normalized): 149.679449746396	Hits (normalized) per kb: 21.4935942047584	Normalized hits with 1T: 1%	Normalized hits with 10A: 99%	Normalized hits 25-32 nt: 95.7%	Normalized hits on the main strand(s): 99.7%	Predicted directionality: mono:plus
Cluster 297	Location: CM035924.1	Coordinates: 54400901-54405766	Size [bp]: 4866	Hits (absolute): 562	Hits (normalized): 172.565312458355	Hits (normalized) per kb: 35.4634432856374	Normalized hits with 1T: 94.7%	Normalized hits with 10A: 6.1%	Normalized hits 25-32 nt: 99.2%	Normalized hits on the main strand(s): 97.7%	Predicted directionality: mono:minus
Cluster 298	Location: CM035924.1	Coordinates: 65226309-65234113	Size [bp]: 7805	Hits (absolute): 2249	Hits (normalized): 186.586627813985	Hits (normalized) per kb: 23.9063587409861	Normalized hits with 1T: 95.2%	Normalized hits with 10A: 26.3%	Normalized hits 25-32 nt: 99.7%	Normalized hits on the main strand(s): 93%	Predicted directionality: mono:plus
Cluster 299	Location: CM035925.1	Coordinates: 662750-670018	Size [bp]: 7269	Hits (absolute): 512	Hits (normalized): 153.533253380615	Hits (normalized) per kb: 21.1217668708426	Normalized hits with 1T: 94.5%	Normalized hits with 10A: 89.3%	Normalized hits 25-32 nt: 99.8%	Normalized hits on the main strand(s): 99.5%	Predicted directionality: mono:minus
Cluster 300	Location: CM035925.1	Coordinates: 6355024-6357086	Size [bp]: 2063	Hits (absolute): 507	Hits (normalized): 971.97547834228	Hits (normalized) per kb: 471.146363413507	Normalized hits with 1T: 99.4%	Normalized hits with 10A: 0.3%	Normalized hits 25-32 nt: 100%	Normalized hits on the main strand(s): 99.9%	Predicted directionality: mono:plus
Cluster 301	Location: CM035925.1	Coordinates: 7737068-7744979	Size [bp]: 7912	Hits (absolute): 1143	Hits (normalized): 156.868700429698	Hits (normalized) per kb: 19.8269522168971	Normalized hits with 1T: 97.9%	Normalized hits with 10A: 38%	Normalized hits 25-32 nt: 99.6%	Normalized hits on the main strand(s): 99.5%	Predicted directionality: mono:minus
Cluster 302	Location: CM035925.1	Coordinates: 7937006-7946025	Size [bp]: 9020	Hits (absolute): 1046	Hits (normalized): 266.739128906807	Hits (normalized) per kb: 29.5717898221316	Normalized hits with 1T: 99.3%	Normalized hits with 10A: 56.2%	Normalized hits 25-32 nt: 99.8%	Normalized hits on the main strand(s): 99.9%	Predicted directionality: mono:minus
Cluster 303	Location: CM035925.1	Coordinates: 7956959-7969027	Size [bp]: 12069	Hits (absolute): 2729	Hits (normalized): 1151.4313784965	Hits (normalized) per kb: 95.4041483426537	Normalized hits with 1T: 95%	Normalized hits with 10A: 12.2%	Normalized hits 25-32 nt: 99%	Normalized hits on the main strand(s): 100%	Predicted directionality: mono:minus
Cluster 304	Location: CM035925.1	Coordinates: 7970286-7977879	Size [bp]: 7594	Hits (absolute): 684	Hits (normalized): 155.374207834774	Hits (normalized) per kb: 20.4603748875058	Normalized hits with 1T: 94.4%	Normalized hits with 10A: 9.6%	Normalized hits 25-32 nt: 99.8%	Normalized hits on the main strand(s): 100%	Predicted directionality: mono:minus
Cluster 305	Location: CM035925.1	Coordinates: 8023073-8060027	Size [bp]: 36955	Hits (absolute): 10431	Hits (normalized): 2136.04178275263	Hits (normalized) per kb: 57.8010526333108	Normalized hits with 1T: 80.9%	Normalized hits with 10A: 31.4%	Normalized hits 25-32 nt: 99.7%	Normalized hits on the main strand(s): 99.7%	Predicted directionality: mono:minus
Cluster 306	Location: CM035925.1	Coordinates: 8064010-8074001	Size [bp]: 9992	Hits (absolute): 2569	Hits (normalized): 728.257068579034	Hits (normalized) per kb: 72.8839158354022	Normalized hits with 1T: 96.6%	Normalized hits with 10A: 15.7%	Normalized hits 25-32 nt: 99.1%	Normalized hits on the main strand(s): 95.9%	Predicted directionality: mono:minus
Cluster 307	Location: CM035925.1	Coordinates: 9352611-9370005	Size [bp]: 17395	Hits (absolute): 2508	Hits (normalized): 913.203664759889	Hits (normalized) per kb: 52.4983999907866	Normalized hits with 1T: 85.2%	Normalized hits with 10A: 19.4%	Normalized hits 25-32 nt: 99.3%	Normalized hits on the main strand(s): 99.7%	Predicted directionality: mono:minus
Cluster 308	Location: CM035925.1	Coordinates: 9391065-9405022	Size [bp]: 13958	Hits (absolute): 3007	Hits (normalized): 441.666236351654	Hits (normalized) per kb: 31.6423415908616	Normalized hits with 1T: 92.5%	Normalized hits with 10A: 17.3%	Normalized hits 25-32 nt: 98.7%	Normalized hits on the main strand(s): 99.8%	Predicted directionality: mono:minus
Cluster 309	Location: CM035925.1	Coordinates: 9518024-9530953	Size [bp]: 12930	Hits (absolute): 2536	Hits (normalized): 403.563486422595	Hits (normalized) per kb: 31.2112851241097	Normalized hits with 1T: 92.4%	Normalized hits with 10A: 16.2%	Normalized hits 25-32 nt: 98.7%	Normalized hits on the main strand(s): 99.8%	Predicted directionality: mono:plus
Cluster 310	Location: CM035925.1	Coordinates: 9549026-9556925	Size [bp]: 7900	Hits (absolute): 1208	Hits (normalized): 196.660079681298	Hits (normalized) per kb: 24.8935109549217	Normalized hits with 1T: 97%	Normalized hits with 10A: 21.8%	Normalized hits 25-32 nt: 99.2%	Normalized hits on the main strand(s): 99.7%	Predicted directionality: mono:plus
Cluster 311	Location: CM035925.1	Coordinates: 16636003-16643777	Size [bp]: 7775	Hits (absolute): 1376	Hits (normalized): 163.976460402943	Hits (normalized) per kb: 21.0905070507347	Normalized hits with 1T: 90.8%	Normalized hits with 10A: 47.5%	Normalized hits 25-32 nt: 99.3%	Normalized hits on the main strand(s): 90.3%	Predicted directionality: mono:minus
Cluster 312	Location: CM035925.1	Coordinates: 17593235-17601990	Size [bp]: 8756	Hits (absolute): 2910	Hits (normalized): 248.344942695073	Hits (normalized) per kb: 28.3625283600605	Normalized hits with 1T: 87.1%	Normalized hits with 10A: 36.2%	Normalized hits 25-32 nt: 99.2%	Normalized hits on the main strand(s): 99.8%	Predicted directionality: mono:plus
Cluster 313	Location: CM035925.1	Coordinates: 17781026-17789947	Size [bp]: 8922	Hits (absolute): 2456	Hits (normalized): 697.43455592135	Hits (normalized) per kb: 78.1701159410275	Normalized hits with 1T: 96.9%	Normalized hits with 10A: 55.2%	Normalized hits 25-32 nt: 98.8%	Normalized hits on the main strand(s): 99.5%	Predicted directionality: mono:minus
Cluster 314	Location: CM035925.1	Coordinates: 27042365-27052837	Size [bp]: 10473	Hits (absolute): 4672	Hits (normalized): 1371.8208376473	Hits (normalized) per kb: 130.986049893963	Normalized hits with 1T: 92.3%	Normalized hits with 10A: 16.6%	Normalized hits 25-32 nt: 99.4%	Normalized hits on the main strand(s): 99.9%	Predicted directionality: mono:plus
Cluster 315	Location: CM035925.1	Coordinates: 27055053-27096018	Size [bp]: 40966	Hits (absolute): 10802	Hits (normalized): 2242.93552283995	Hits (normalized) per kb: 54.7507522922497	Normalized hits with 1T: 93.6%	Normalized hits with 10A: 13.8%	Normalized hits 25-32 nt: 99.5%	Normalized hits on the main strand(s): 100%	Predicted directionality: mono:plus
Cluster 316	Location: CM035925.1	Coordinates: 28258012-28313617	Size [bp]: 55606	Hits (absolute): 26355	Hits (normalized): 10128.4778177344	Hits (normalized) per kb: 182.146858634867	Normalized hits with 1T: 87.7%	Normalized hits with 10A: 15.8%	Normalized hits 25-32 nt: 99.6%	Normalized hits on the main strand(s): 100%	Predicted directionality: mono:plus
Cluster 317	Location: CM035925.1	Coordinates: 28316278-28325530	Size [bp]: 9253	Hits (absolute): 2194	Hits (normalized): 420.33860284824	Hits (normalized) per kb: 45.4270996316277	Normalized hits with 1T: 95.1%	Normalized hits with 10A: 14.7%	Normalized hits 25-32 nt: 99.8%	Normalized hits on the main strand(s): 99.9%	Predicted directionality: mono:plus
Cluster 318	Location: CM035925.1	Coordinates: 38010083-38017612	Size [bp]: 7530	Hits (absolute): 2152	Hits (normalized): 180.15266485742	Hits (normalized) per kb: 23.9244565315749	Normalized hits with 1T: 95.9%	Normalized hits with 10A: 26.7%	Normalized hits 25-32 nt: 99.8%	Normalized hits on the main strand(s): 93.4%	Predicted directionality: mono:plus
Cluster 319	Location: CM035925.1	Coordinates: 43457006-43484803	Size [bp]: 27798	Hits (absolute): 7865	Hits (normalized): 1727.36437259212	Hits (normalized) per kb: 62.1395866135579	Normalized hits with 1T: 96%	Normalized hits with 10A: 13.9%	Normalized hits 25-32 nt: 99.6%	Normalized hits on the main strand(s): 98.9%	Predicted directionality: mono:plus
Cluster 320	Location: CM035925.1	Coordinates: 43490232-43496013	Size [bp]: 5782	Hits (absolute): 1438	Hits (normalized): 132.619077316214	Hits (normalized) per kb: 22.9364816907943	Normalized hits with 1T: 93.2%	Normalized hits with 10A: 32.8%	Normalized hits 25-32 nt: 99.7%	Normalized hits on the main strand(s): 99.9%	Predicted directionality: mono:plus
Cluster 321	Location: CM035925.1	Coordinates: 43500071-43520011	Size [bp]: 19941	Hits (absolute): 7239	Hits (normalized): 972.335391666769	Hits (normalized) per kb: 48.7603836073503	Normalized hits with 1T: 94%	Normalized hits with 10A: 21.6%	Normalized hits 25-32 nt: 99.7%	Normalized hits on the main strand(s): 98.7%	Predicted directionality: mono:plus
Cluster 322	Location: CM035925.1	Coordinates: 44869097-44880027	Size [bp]: 10931	Hits (absolute): 3411	Hits (normalized): 414.441208388938	Hits (normalized) per kb: 37.9140486567326	Normalized hits with 1T: 90.3%	Normalized hits with 10A: 45.8%	Normalized hits 25-32 nt: 99.5%	Normalized hits on the main strand(s): 95.6%	Predicted directionality: mono:plus
Cluster 323	Location: CM035925.1	Coordinates: 52631199-52637548	Size [bp]: 6350	Hits (absolute): 379	Hits (normalized): 138.073585591364	Hits (normalized) per kb: 21.7436727656221	Normalized hits with 1T: 99.4%	Normalized hits with 10A: 34.4%	Normalized hits 25-32 nt: 99.8%	Normalized hits on the main strand(s): 99.9%	Predicted directionality: mono:plus
Cluster 324	Location: CM035925.1	Coordinates: 52643002-52651666	Size [bp]: 8665	Hits (absolute): 1706	Hits (normalized): 371.961677967103	Hits (normalized) per kb: 42.9271366498357	Normalized hits with 1T: 93.6%	Normalized hits with 10A: 37.1%	Normalized hits 25-32 nt: 96.7%	Normalized hits on the main strand(s): 98.2%	Predicted directionality: mono:plus
Cluster 325	Location: CM035925.1	Coordinates: 52702028-52710722	Size [bp]: 8695	Hits (absolute): 1935	Hits (normalized): 402.074444351846	Hits (normalized) per kb: 46.2423228349695	Normalized hits with 1T: 81.5%	Normalized hits with 10A: 29.7%	Normalized hits 25-32 nt: 99.7%	Normalized hits on the main strand(s): 99.7%	Predicted directionality: mono:plus
Cluster 326	Location: CM035925.1	Coordinates: 52788008-52797006	Size [bp]: 8999	Hits (absolute): 585	Hits (normalized): 142.545778304574	Hits (normalized) per kb: 15.840502526287	Normalized hits with 1T: 98.6%	Normalized hits with 10A: 34.2%	Normalized hits 25-32 nt: 99.7%	Normalized hits on the main strand(s): 99%	Predicted directionality: mono:plus
Cluster 327	Location: CM035925.1	Coordinates: 52805242-52813997	Size [bp]: 8756	Hits (absolute): 1721	Hits (normalized): 375.9832174967	Hits (normalized) per kb: 42.9402986793549	Normalized hits with 1T: 95.2%	Normalized hits with 10A: 35.3%	Normalized hits 25-32 nt: 96.7%	Normalized hits on the main strand(s): 99.9%	Predicted directionality: mono:plus
Cluster 328	Location: CM035925.1	Coordinates: 52842004-52849082	Size [bp]: 7079	Hits (absolute): 2010	Hits (normalized): 202.698760421287	Hits (normalized) per kb: 28.6339952188928	Normalized hits with 1T: 88.8%	Normalized hits with 10A: 29%	Normalized hits 25-32 nt: 99.3%	Normalized hits on the main strand(s): 96.9%	Predicted directionality: mono:plus
Cluster 329	Location: CM035925.1	Coordinates: 52862031-52867027	Size [bp]: 4997	Hits (absolute): 1573	Hits (normalized): 123.612845482709	Hits (normalized) per kb: 24.7372118543819	Normalized hits with 1T: 97.3%	Normalized hits with 10A: 22.5%	Normalized hits 25-32 nt: 99.6%	Normalized hits on the main strand(s): 99.8%	Predicted directionality: mono:plus
Cluster 330	Location: CM035925.1	Coordinates: 52872001-52880691	Size [bp]: 8691	Hits (absolute): 2228	Hits (normalized): 212.024511675728	Hits (normalized) per kb: 24.3958217137291	Normalized hits with 1T: 94.5%	Normalized hits with 10A: 8.6%	Normalized hits 25-32 nt: 99.4%	Normalized hits on the main strand(s): 98.8%	Predicted directionality: mono:plus
Cluster 331	Location: CM035925.1	Coordinates: 52927224-52935126	Size [bp]: 7903	Hits (absolute): 723	Hits (normalized): 294.674520115407	Hits (normalized) per kb: 37.2863843740386	Normalized hits with 1T: 97.2%	Normalized hits with 10A: 89.1%	Normalized hits 25-32 nt: 100%	Normalized hits on the main strand(s): 99.9%	Predicted directionality: mono:plus
Cluster 332	Location: CM035925.1	Coordinates: 53077077-53087949	Size [bp]: 10873	Hits (absolute): 3485	Hits (normalized): 813.26073433699	Hits (normalized) per kb: 74.7965232499025	Normalized hits with 1T: 93%	Normalized hits with 10A: 22.9%	Normalized hits 25-32 nt: 98.7%	Normalized hits on the main strand(s): 99.9%	Predicted directionality: mono:plus
Cluster 333	Location: CM035925.1	Coordinates: 53094050-53104989	Size [bp]: 10940	Hits (absolute): 2224	Hits (normalized): 348.031537765895	Hits (normalized) per kb: 31.8126253477655	Normalized hits with 1T: 84.9%	Normalized hits with 10A: 31.8%	Normalized hits 25-32 nt: 99.6%	Normalized hits on the main strand(s): 99.9%	Predicted directionality: mono:plus
Cluster 334	Location: CM035925.1	Coordinates: 53111004-53119751	Size [bp]: 8748	Hits (absolute): 1958	Hits (normalized): 382.19806537105	Hits (normalized) per kb: 43.689711735101	Normalized hits with 1T: 80.6%	Normalized hits with 10A: 28.6%	Normalized hits 25-32 nt: 99.6%	Normalized hits on the main strand(s): 99.6%	Predicted directionality: mono:plus
Cluster 335	Location: CM035925.1	Coordinates: 55953015-55991010	Size [bp]: 37996	Hits (absolute): 9200	Hits (normalized): 3233.22844824186	Hits (normalized) per kb: 85.094166094941	Normalized hits with 1T: 93.5%	Normalized hits with 10A: 47.1%	Normalized hits 25-32 nt: 99.8%	Normalized hits on the main strand(s): 100%	Predicted directionality: mono:minus
Cluster 336	Location: CM035925.1	Coordinates: 56005006-56011704	Size [bp]: 6699	Hits (absolute): 617	Hits (normalized): 202.763733118053	Hits (normalized) per kb: 30.2677321329562	Normalized hits with 1T: 94.5%	Normalized hits with 10A: 36.2%	Normalized hits 25-32 nt: 99.8%	Normalized hits on the main strand(s): 99.9%	Predicted directionality: mono:minus
Cluster 337	Location: CM035925.1	Coordinates: 65854059-65857907	Size [bp]: 3849	Hits (absolute): 526	Hits (normalized): 153.616591892271	Hits (normalized) per kb: 39.9105640094174	Normalized hits with 1T: 95.5%	Normalized hits with 10A: 39.4%	Normalized hits 25-32 nt: 99.7%	Normalized hits on the main strand(s): 97.2%	Predicted directionality: mono:plus
Cluster 338	Location: CM035925.1	Coordinates: 78519916-78524597	Size [bp]: 4682	Hits (absolute): 988	Hits (normalized): 166.699658332764	Hits (normalized) per kb: 35.6041124761233	Normalized hits with 1T: 87.3%	Normalized hits with 10A: 19.4%	Normalized hits 25-32 nt: 98.1%	Normalized hits on the main strand(s): 91.2%	Predicted directionality: mono:minus
Cluster 339	Location: CM035925.1	Coordinates: 81355396-81361857	Size [bp]: 6462	Hits (absolute): 1272	Hits (normalized): 257.220604847832	Hits (normalized) per kb: 39.8052677732643	Normalized hits with 1T: 96.5%	Normalized hits with 10A: 7.6%	Normalized hits 25-32 nt: 99.8%	Normalized hits on the main strand(s): 96.8%	Predicted directionality: mono:plus
Cluster 340	Location: CM035925.1	Coordinates: 86730030-86740829	Size [bp]: 10800	Hits (absolute): 2460	Hits (normalized): 375.292341396842	Hits (normalized) per kb: 34.749403184224	Normalized hits with 1T: 94.4%	Normalized hits with 10A: 14.6%	Normalized hits 25-32 nt: 99.3%	Normalized hits on the main strand(s): 94.8%	Predicted directionality: mono:plus
Cluster 341	Location: CM035926.1	Coordinates: 5702393-5707012	Size [bp]: 4620	Hits (absolute): 1122	Hits (normalized): 123.730165306835	Hits (normalized) per kb: 26.7814395640736	Normalized hits with 1T: 88.4%	Normalized hits with 10A: 18.5%	Normalized hits 25-32 nt: 99.5%	Normalized hits on the main strand(s): 95.6%	Predicted directionality: mono:minus
Cluster 342	Location: CM035926.1	Coordinates: 19963493-19976021	Size [bp]: 12529	Hits (absolute): 1777	Hits (normalized): 517.310596622509	Hits (normalized) per kb: 41.2892866015475	Normalized hits with 1T: 98.8%	Normalized hits with 10A: 8.9%	Normalized hits 25-32 nt: 99.8%	Normalized hits on the main strand(s): 99.1%	Predicted directionality: mono:plus
Cluster 343	Location: CM035926.1	Coordinates: 56889164-56895023	Size [bp]: 5860	Hits (absolute): 1026	Hits (normalized): 131.738428039746	Hits (normalized) per kb: 22.480746418694	Normalized hits with 1T: 96.8%	Normalized hits with 10A: 15.7%	Normalized hits 25-32 nt: 99.7%	Normalized hits on the main strand(s): 100%	Predicted directionality: mono:plus
Cluster 344	Location: CM035926.1	Coordinates: 56971001-56984922	Size [bp]: 13922	Hits (absolute): 2580	Hits (normalized): 549.829857831109	Hits (normalized) per kb: 39.4934921990296	Normalized hits with 1T: 76.1%	Normalized hits with 10A: 13.5%	Normalized hits 25-32 nt: 99.3%	Normalized hits on the main strand(s): 99.5%	Predicted directionality: mono:plus
Cluster 345	Location: CM035926.1	Coordinates: 67034906-67041933	Size [bp]: 7028	Hits (absolute): 1598	Hits (normalized): 320.414153266982	Hits (normalized) per kb: 45.590802373772	Normalized hits with 1T: 96.7%	Normalized hits with 10A: 6.4%	Normalized hits 25-32 nt: 99.8%	Normalized hits on the main strand(s): 97%	Predicted directionality: mono:minus
Cluster 346	Location: CM035926.1	Coordinates: 67047015-67054981	Size [bp]: 7967	Hits (absolute): 1076	Hits (normalized): 289.584029821095	Hits (normalized) per kb: 36.3477671439548	Normalized hits with 1T: 21%	Normalized hits with 10A: 80.6%	Normalized hits 25-32 nt: 98.5%	Normalized hits on the main strand(s): 95.1%	Predicted directionality: bi:plus-minus (split between 67050782 and 67050782)
Cluster 347	Location: CM035927.1	Coordinates: 7638632-7643336	Size [bp]: 4705	Hits (absolute): 974	Hits (normalized): 2381.96590129465	Hits (normalized) per kb: 506.262658170577	Normalized hits with 1T: 97.5%	Normalized hits with 10A: 9.6%	Normalized hits 25-32 nt: 99.9%	Normalized hits on the main strand(s): 97.5%	Predicted directionality: mono:minus
Cluster 348	Location: CM035927.1	Coordinates: 34524487-34529826	Size [bp]: 5340	Hits (absolute): 268	Hits (normalized): 120.091406310227	Hits (normalized) per kb: 22.4889726871435	Normalized hits with 1T: 97.8%	Normalized hits with 10A: 94.4%	Normalized hits 25-32 nt: 99.6%	Normalized hits on the main strand(s): 99.4%	Predicted directionality: mono:minus
Cluster 349	Location: CM035927.1	Coordinates: 34659896-34664529	Size [bp]: 4634	Hits (absolute): 242	Hits (normalized): 119.809731816208	Hits (normalized) per kb: 25.854339109819	Normalized hits with 1T: 97.9%	Normalized hits with 10A: 94.6%	Normalized hits 25-32 nt: 99.7%	Normalized hits on the main strand(s): 99.4%	Predicted directionality: mono:minus
Cluster 350	Location: CM035927.1	Coordinates: 34877667-34885875	Size [bp]: 8209	Hits (absolute): 370	Hits (normalized): 177.608046646067	Hits (normalized) per kb: 21.6359086489341	Normalized hits with 1T: 1.2%	Normalized hits with 10A: 90.7%	Normalized hits 25-32 nt: 99.6%	Normalized hits on the main strand(s): 99.4%	Predicted directionality: mono:minus
Cluster 351	Location: CM035927.1	Coordinates: 35114499-35120270	Size [bp]: 5772	Hits (absolute): 293	Hits (normalized): 164.354410686964	Hits (normalized) per kb: 28.4744056109732	Normalized hits with 1T: 1.5%	Normalized hits with 10A: 98.4%	Normalized hits 25-32 nt: 99.8%	Normalized hits on the main strand(s): 98.1%	Predicted directionality: mono:minus
Cluster 352	Location: CM035927.1	Coordinates: 49941209-49945977	Size [bp]: 4769	Hits (absolute): 114	Hits (normalized): 137.231474631089	Hits (normalized) per kb: 28.7754870362236	Normalized hits with 1T: 99.8%	Normalized hits with 10A: 99.5%	Normalized hits 25-32 nt: 100%	Normalized hits on the main strand(s): 99.8%	Predicted directionality: mono:minus
Cluster 353	Location: CM035927.1	Coordinates: 50679159-50687772	Size [bp]: 8614	Hits (absolute): 2124	Hits (normalized): 947.495877350114	Hits (normalized) per kb: 109.995080691467	Normalized hits with 1T: 93.7%	Normalized hits with 10A: 7.8%	Normalized hits 25-32 nt: 99.7%	Normalized hits on the main strand(s): 99.9%	Predicted directionality: mono:plus
Cluster 354	Location: CM035927.1	Coordinates: 66064324-66071017	Size [bp]: 6694	Hits (absolute): 955	Hits (normalized): 172.865047211694	Hits (normalized) per kb: 25.823901916556	Normalized hits with 1T: 97.8%	Normalized hits with 10A: 38.3%	Normalized hits 25-32 nt: 99.7%	Normalized hits on the main strand(s): 98.3%	Predicted directionality: mono:minus
Cluster 355	Location: CM035927.1	Coordinates: 67022040-67028813	Size [bp]: 6774	Hits (absolute): 1364	Hits (normalized): 148.79382243892	Hits (normalized) per kb: 21.9657820137576	Normalized hits with 1T: 95.7%	Normalized hits with 10A: 22.7%	Normalized hits 25-32 nt: 99.5%	Normalized hits on the main strand(s): 100%	Predicted directionality: mono:minus
Cluster 356	Location: CM035927.1	Coordinates: 67037136-67049015	Size [bp]: 11880	Hits (absolute): 2736	Hits (normalized): 536.55395465961	Hits (normalized) per kb: 45.1646816680898	Normalized hits with 1T: 88.4%	Normalized hits with 10A: 14.2%	Normalized hits 25-32 nt: 99.6%	Normalized hits on the main strand(s): 99.3%	Predicted directionality: mono:minus
Cluster 357	Location: CM035928.1	Coordinates: 304200-308646	Size [bp]: 4447	Hits (absolute): 533	Hits (normalized): 422.843017499843	Hits (normalized) per kb: 95.0849691268145	Normalized hits with 1T: 99.6%	Normalized hits with 10A: 45.5%	Normalized hits 25-32 nt: 99.7%	Normalized hits on the main strand(s): 99.9%	Predicted directionality: mono:plus
Cluster 358	Location: CM035928.1	Coordinates: 1648098-1654099	Size [bp]: 6002	Hits (absolute): 1171	Hits (normalized): 124.198623755762	Hits (normalized) per kb: 20.6931782846256	Normalized hits with 1T: 86.9%	Normalized hits with 10A: 20.9%	Normalized hits 25-32 nt: 99.4%	Normalized hits on the main strand(s): 94%	Predicted directionality: mono:minus
Cluster 359	Location: CM035928.1	Coordinates: 11097416-11106614	Size [bp]: 9199	Hits (absolute): 796	Hits (normalized): 595.006357541378	Hits (normalized) per kb: 64.6815035644421	Normalized hits with 1T: 99.2%	Normalized hits with 10A: 0.8%	Normalized hits 25-32 nt: 100%	Normalized hits on the main strand(s): 99.9%	Predicted directionality: bi:plus-minus (split between 11103456 and 11103481)
Cluster 360	Location: CM035928.1	Coordinates: 29201270-29204827	Size [bp]: 3558	Hits (absolute): 751	Hits (normalized): 316.736523354885	Hits (normalized) per kb: 89.02056402587	Normalized hits with 1T: 97.6%	Normalized hits with 10A: 2.3%	Normalized hits 25-32 nt: 99.5%	Normalized hits on the main strand(s): 96.9%	Predicted directionality: mono:plus
Cluster 361	Location: CM035928.1	Coordinates: 31189107-31194960	Size [bp]: 5854	Hits (absolute): 2051	Hits (normalized): 153.036928084682	Hits (normalized) per kb: 26.1422585055503	Normalized hits with 1T: 95.8%	Normalized hits with 10A: 34.2%	Normalized hits 25-32 nt: 99.7%	Normalized hits on the main strand(s): 98.1%	Predicted directionality: mono:plus
Cluster 362	Location: CM035928.1	Coordinates: 33890052-33896387	Size [bp]: 6336	Hits (absolute): 1447	Hits (normalized): 279.37781156096	Hits (normalized) per kb: 44.0936215159697	Normalized hits with 1T: 96.9%	Normalized hits with 10A: 6.1%	Normalized hits 25-32 nt: 99.8%	Normalized hits on the main strand(s): 97.3%	Predicted directionality: mono:minus
Cluster 363	Location: CM035928.1	Coordinates: 44560417-44564472	Size [bp]: 4056	Hits (absolute): 818	Hits (normalized): 170.087044702882	Hits (normalized) per kb: 41.9350486748304	Normalized hits with 1T: 97.7%	Normalized hits with 10A: 36.3%	Normalized hits 25-32 nt: 99.8%	Normalized hits on the main strand(s): 98.3%	Predicted directionality: mono:plus
Cluster 364	Location: CM035928.1	Coordinates: 44843147-44850286	Size [bp]: 7140	Hits (absolute): 1117	Hits (normalized): 162.027779762458	Hits (normalized) per kb: 22.6929841446902	Normalized hits with 1T: 94.4%	Normalized hits with 10A: 10.2%	Normalized hits 25-32 nt: 99.7%	Normalized hits on the main strand(s): 94.4%	Predicted directionality: mono:minus
Cluster 365	Location: CM035928.1	Coordinates: 45072870-45076166	Size [bp]: 3297	Hits (absolute): 899	Hits (normalized): 176.763239114883	Hits (normalized) per kb: 53.6130593656889	Normalized hits with 1T: 98%	Normalized hits with 10A: 38.2%	Normalized hits 25-32 nt: 99.8%	Normalized hits on the main strand(s): 98.6%	Predicted directionality: mono:plus
Cluster 366	Location: CM035928.1	Coordinates: 48501001-48509975	Size [bp]: 8975	Hits (absolute): 1984	Hits (normalized): 242.741422187804	Hits (normalized) per kb: 27.0463254081463	Normalized hits with 1T: 93.5%	Normalized hits with 10A: 27.8%	Normalized hits 25-32 nt: 99.6%	Normalized hits on the main strand(s): 100%	Predicted directionality: mono:minus
Cluster 367	Location: CM035928.1	Coordinates: 48805024-48813913	Size [bp]: 8890	Hits (absolute): 1750	Hits (normalized): 224.015403730293	Hits (normalized) per kb: 25.1987055143968	Normalized hits with 1T: 92.6%	Normalized hits with 10A: 28.3%	Normalized hits 25-32 nt: 99.7%	Normalized hits on the main strand(s): 100%	Predicted directionality: mono:minus
Cluster 368	Location: CM035928.1	Coordinates: 48816003-48825996	Size [bp]: 9994	Hits (absolute): 2548	Hits (normalized): 355.716496728754	Hits (normalized) per kb: 35.592595700294	Normalized hits with 1T: 94.9%	Normalized hits with 10A: 25.5%	Normalized hits 25-32 nt: 99.7%	Normalized hits on the main strand(s): 99.6%	Predicted directionality: mono:minus
Cluster 369	Location: CM035928.1	Coordinates: 48833001-48856988	Size [bp]: 23988	Hits (absolute): 11209	Hits (normalized): 4954.20338587881	Hits (normalized) per kb: 206.528695692232	Normalized hits with 1T: 79.1%	Normalized hits with 10A: 8.3%	Normalized hits 25-32 nt: 99.5%	Normalized hits on the main strand(s): 99.8%	Predicted directionality: mono:minus
Cluster 370	Location: CM035928.1	Coordinates: 52237229-52242026	Size [bp]: 4798	Hits (absolute): 619	Hits (normalized): 119.397275959211	Hits (normalized) per kb: 24.8844620596273	Normalized hits with 1T: 96.8%	Normalized hits with 10A: 69.3%	Normalized hits 25-32 nt: 99.9%	Normalized hits on the main strand(s): 99.1%	Predicted directionality: mono:minus
Cluster 371	Location: CM035928.1	Coordinates: 56853184-56861809	Size [bp]: 8626	Hits (absolute): 465	Hits (normalized): 129.888528968538	Hits (normalized) per kb: 15.0581843967431	Normalized hits with 1T: 93.3%	Normalized hits with 10A: 24.9%	Normalized hits 25-32 nt: 99.8%	Normalized hits on the main strand(s): 100%	Predicted directionality: mono:plus
Cluster 372	Location: CM035928.1	Coordinates: 56906300-56911939	Size [bp]: 5640	Hits (absolute): 872	Hits (normalized): 148.00800505915	Hits (normalized) per kb: 26.2426189806337	Normalized hits with 1T: 98.1%	Normalized hits with 10A: 28.9%	Normalized hits 25-32 nt: 99.6%	Normalized hits on the main strand(s): 99.5%	Predicted directionality: mono:plus
Cluster 373	Location: CM035928.1	Coordinates: 57626628-57642015	Size [bp]: 15388	Hits (absolute): 3543	Hits (normalized): 499.575155736017	Hits (normalized) per kb: 32.464968435808	Normalized hits with 1T: 94%	Normalized hits with 10A: 17%	Normalized hits 25-32 nt: 98.8%	Normalized hits on the main strand(s): 99.6%	Predicted directionality: mono:plus
Cluster 374	Location: CM035928.1	Coordinates: 67293639-67301622	Size [bp]: 7984	Hits (absolute): 1305	Hits (normalized): 487.657268975109	Hits (normalized) per kb: 61.079220610422	Normalized hits with 1T: 98.9%	Normalized hits with 10A: 8.4%	Normalized hits 25-32 nt: 99.8%	Normalized hits on the main strand(s): 99.2%	Predicted directionality: mono:minus
Cluster 375	Location: CM035929.1	Coordinates: 8762360-8769025	Size [bp]: 6666	Hits (absolute): 2120	Hits (normalized): 159.856294632181	Hits (normalized) per kb: 23.9812177838762	Normalized hits with 1T: 91.1%	Normalized hits with 10A: 35.1%	Normalized hits 25-32 nt: 99.7%	Normalized hits on the main strand(s): 99.7%	Predicted directionality: mono:plus
Cluster 376	Location: CM035929.1	Coordinates: 57702053-57714021	Size [bp]: 11969	Hits (absolute): 2217	Hits (normalized): 504.292962029565	Hits (normalized) per kb: 42.1333017444625	Normalized hits with 1T: 95.8%	Normalized hits with 10A: 15.8%	Normalized hits 25-32 nt: 99.4%	Normalized hits on the main strand(s): 96.4%	Predicted directionality: mono:plus
Cluster 377	Location: CM035929.1	Coordinates: 60394186-60395324	Size [bp]: 1139	Hits (absolute): 379	Hits (normalized): 2317.95489118256	Hits (normalized) per kb: 2035.07927654905	Normalized hits with 1T: 99.9%	Normalized hits with 10A: 0%	Normalized hits 25-32 nt: 99.9%	Normalized hits on the main strand(s): 100%	Predicted directionality: mono:minus
Cluster 378	Location: CM035929.1	Coordinates: 62732407-62766016	Size [bp]: 33610	Hits (absolute): 13265	Hits (normalized): 6544.91587720652	Hits (normalized) per kb: 194.731404108857	Normalized hits with 1T: 96.1%	Normalized hits with 10A: 26.8%	Normalized hits 25-32 nt: 99.7%	Normalized hits on the main strand(s): 99.9%	Predicted directionality: mono:minus
Cluster 379	Location: CM035929.1	Coordinates: 63703024-63723027	Size [bp]: 20004	Hits (absolute): 7865	Hits (normalized): 5054.47125015342	Hits (normalized) per kb: 252.673125932653	Normalized hits with 1T: 95.9%	Normalized hits with 10A: 13.8%	Normalized hits 25-32 nt: 99.8%	Normalized hits on the main strand(s): 100%	Predicted directionality: mono:plus
Cluster 380	Location: CM035929.1	Coordinates: 63726595-63733847	Size [bp]: 7253	Hits (absolute): 2452	Hits (normalized): 2307.36831195087	Hits (normalized) per kb: 318.126253477655	Normalized hits with 1T: 98.7%	Normalized hits with 10A: 3.9%	Normalized hits 25-32 nt: 99.9%	Normalized hits on the main strand(s): 100%	Predicted directionality: mono:plus
Cluster 381	Location: CM035929.1	Coordinates: 64474214-64483881	Size [bp]: 9668	Hits (absolute): 1461	Hits (normalized): 316.372892159254	Hits (normalized) per kb: 32.7240958919661	Normalized hits with 1T: 97.1%	Normalized hits with 10A: 6%	Normalized hits 25-32 nt: 99.9%	Normalized hits on the main strand(s): 96.8%	Predicted directionality: mono:plus
Cluster 382	Location: CM035930.1	Coordinates: 5830032-5836980	Size [bp]: 6949	Hits (absolute): 1543	Hits (normalized): 138.478521556069	Hits (normalized) per kb: 19.9281353188255	Normalized hits with 1T: 88.1%	Normalized hits with 10A: 37.3%	Normalized hits 25-32 nt: 98.7%	Normalized hits on the main strand(s): 99.3%	Predicted directionality: mono:plus
Cluster 383	Location: CM035930.1	Coordinates: 9935498-9942791	Size [bp]: 7294	Hits (absolute): 2097	Hits (normalized): 225.455254544393	Hits (normalized) per kb: 30.9093810720144	Normalized hits with 1T: 87.1%	Normalized hits with 10A: 21.5%	Normalized hits 25-32 nt: 99.4%	Normalized hits on the main strand(s): 94.7%	Predicted directionality: bi:plus-minus (split between 9938729 and 9938795)
Cluster 384	Location: CM035930.1	Coordinates: 18226204-18230630	Size [bp]: 4427	Hits (absolute): 127	Hits (normalized): 173.28634488795	Hits (normalized) per kb: 39.1430531630825	Normalized hits with 1T: 0.1%	Normalized hits with 10A: 99.7%	Normalized hits 25-32 nt: 99.8%	Normalized hits on the main strand(s): 100%	Predicted directionality: mono:plus
Cluster 385	Location: CM035930.1	Coordinates: 24732262-24734216	Size [bp]: 1955	Hits (absolute): 374	Hits (normalized): 891.937715973885	Hits (normalized) per kb: 456.233783968319	Normalized hits with 1T: 99.9%	Normalized hits with 10A: 46.2%	Normalized hits 25-32 nt: 99.9%	Normalized hits on the main strand(s): 100%	Predicted directionality: mono:minus
Cluster 386	Location: CM035930.1	Coordinates: 27810011-27819028	Size [bp]: 9018	Hits (absolute): 920	Hits (normalized): 190.105087301529	Hits (normalized) per kb: 21.0806355285953	Normalized hits with 1T: 89.3%	Normalized hits with 10A: 29.3%	Normalized hits 25-32 nt: 96.1%	Normalized hits on the main strand(s): 97.4%	Predicted directionality: mono:plus
Cluster 387	Location: CM035930.1	Coordinates: 40523176-40539828	Size [bp]: 16653	Hits (absolute): 5587	Hits (normalized): 738.402570530961	Hits (normalized) per kb: 44.3404095694536	Normalized hits with 1T: 95.5%	Normalized hits with 10A: 12%	Normalized hits 25-32 nt: 99.7%	Normalized hits on the main strand(s): 99.8%	Predicted directionality: mono:minus
Cluster 388	Location: CM035930.1	Coordinates: 40563130-40579626	Size [bp]: 16497	Hits (absolute): 7029	Hits (normalized): 1222.92035942218	Hits (normalized) per kb: 74.130195505496	Normalized hits with 1T: 97.4%	Normalized hits with 10A: 7.8%	Normalized hits 25-32 nt: 99.2%	Normalized hits on the main strand(s): 100%	Predicted directionality: mono:minus
Cluster 389	Location: CM035930.1	Coordinates: 47261235-47284910	Size [bp]: 23676	Hits (absolute): 4622	Hits (normalized): 680.674859607474	Hits (normalized) per kb: 28.7491629771853	Normalized hits with 1T: 79.2%	Normalized hits with 10A: 22.9%	Normalized hits 25-32 nt: 99.2%	Normalized hits on the main strand(s): 99.5%	Predicted directionality: mono:minus
Cluster 390	Location: CM035930.1	Coordinates: 47318001-47328991	Size [bp]: 10991	Hits (absolute): 2513	Hits (normalized): 313.02969967353	Hits (normalized) per kb: 28.4801639988878	Normalized hits with 1T: 95.3%	Normalized hits with 10A: 29.3%	Normalized hits 25-32 nt: 98.9%	Normalized hits on the main strand(s): 99.9%	Predicted directionality: mono:minus
Cluster 391	Location: CM035930.1	Coordinates: 47335025-47340888	Size [bp]: 5864	Hits (absolute): 2374	Hits (normalized): 169.593435876512	Hits (normalized) per kb: 28.9210919877791	Normalized hits with 1T: 95%	Normalized hits with 10A: 37.2%	Normalized hits 25-32 nt: 99.6%	Normalized hits on the main strand(s): 97.7%	Predicted directionality: mono:minus
Cluster 392	Location: CM035930.1	Coordinates: 47488051-47494830	Size [bp]: 6780	Hits (absolute): 1270	Hits (normalized): 131.878792380093	Hits (normalized) per kb: 19.4510117487566	Normalized hits with 1T: 87.4%	Normalized hits with 10A: 20.1%	Normalized hits 25-32 nt: 99.4%	Normalized hits on the main strand(s): 94.1%	Predicted directionality: mono:minus
Cluster 393	Location: CM035930.1	Coordinates: 48329230-48334517	Size [bp]: 5288	Hits (absolute): 2375	Hits (normalized): 169.353503046736	Hits (normalized) per kb: 32.0256857006066	Normalized hits with 1T: 94.9%	Normalized hits with 10A: 37.6%	Normalized hits 25-32 nt: 99.6%	Normalized hits on the main strand(s): 97.6%	Predicted directionality: mono:minus
Cluster 394	Location: JAIWYP010000017.1	Coordinates: 75533-84986	Size [bp]: 9454	Hits (absolute): 2344	Hits (normalized): 307.768989988169	Hits (normalized) per kb: 32.5546347619071	Normalized hits with 1T: 87.8%	Normalized hits with 10A: 22%	Normalized hits 25-32 nt: 99.4%	Normalized hits on the main strand(s): 96.9%	Predicted directionality: mono:plus
Cluster 395	Location: JAIWYP010000017.1	Coordinates: 88068-101814	Size [bp]: 13747	Hits (absolute): 3382	Hits (normalized): 408.76158047033	Hits (normalized) per kb: 29.734669937431	Normalized hits with 1T: 93.3%	Normalized hits with 10A: 16.4%	Normalized hits 25-32 nt: 99.7%	Normalized hits on the main strand(s): 99.8%	Predicted directionality: mono:plus
Cluster 396	Location: JAIWYP010000017.1	Coordinates: 107000-129018	Size [bp]: 22019	Hits (absolute): 5111	Hits (normalized): 692.092635486018	Hits (normalized) per kb: 31.4317491185553	Normalized hits with 1T: 84.9%	Normalized hits with 10A: 8.4%	Normalized hits 25-32 nt: 95.6%	Normalized hits on the main strand(s): 99.6%	Predicted directionality: mono:plus
Cluster 397	Location: JAIWYP010000017.1	Coordinates: 191048-199961	Size [bp]: 8914	Hits (absolute): 1905	Hits (normalized): 369.802686144868	Hits (normalized) per kb: 41.4858944174897	Normalized hits with 1T: 95.3%	Normalized hits with 10A: 18.3%	Normalized hits 25-32 nt: 99.7%	Normalized hits on the main strand(s): 96.5%	Predicted directionality: mono:plus
Cluster 398	Location: JAIWYP010000022.1	Coordinates: 293199-294966	Size [bp]: 1768	Hits (absolute): 257	Hits (normalized): 300.5713966065	Hits (normalized) per kb: 170.006531657149	Normalized hits with 1T: 98.8%	Normalized hits with 10A: 0.4%	Normalized hits 25-32 nt: 99.7%	Normalized hits on the main strand(s): 100%	Predicted directionality: mono:plus
Cluster 399	Location: JAIWYP010000026.1	Coordinates: 273191-280278	Size [bp]: 7088	Hits (absolute): 2222	Hits (normalized): 184.928732325344	Hits (normalized) per kb: 26.0904330143186	Normalized hits with 1T: 95.3%	Normalized hits with 10A: 26.5%	Normalized hits 25-32 nt: 99.8%	Normalized hits on the main strand(s): 93.4%	Predicted directionality: mono:plus
Cluster 400	Location: JAIWYP010000031.1	Coordinates: 291818-296593	Size [bp]: 4776	Hits (absolute): 552	Hits (normalized): 154.067094201677	Hits (normalized) per kb: 32.2584890977264	Normalized hits with 1T: 95.2%	Normalized hits with 10A: 39.5%	Normalized hits 25-32 nt: 99.7%	Normalized hits on the main strand(s): 97%	Predicted directionality: mono:plus
Cluster 401	Location: JAIWYP010000041.1	Coordinates: 205035-210931	Size [bp]: 5897	Hits (absolute): 1557	Hits (normalized): 128.368126523386	Hits (normalized) per kb: 21.7683515709705	Normalized hits with 1T: 93.1%	Normalized hits with 10A: 10.8%	Normalized hits 25-32 nt: 99.3%	Normalized hits on the main strand(s): 99%	Predicted directionality: mono:minus
Cluster 402	Location: JAIWYP010000089.1	Coordinates: 79299-87023	Size [bp]: 7725	Hits (absolute): 470	Hits (normalized): 133.781157500984	Hits (normalized) per kb: 17.3179403398107	Normalized hits with 1T: 99%	Normalized hits with 10A: 6.6%	Normalized hits 25-32 nt: 99.9%	Normalized hits on the main strand(s): 96.6%	Predicted directionality: mono:plus

Total size of 402 predicted piRNA clusters: 4039661 bp (0.225%)
Non identical sequences that can be assigned to clusters: 723627 (12.596%)
Sequence reads that can be assigned to clusters: 42576481 (35.025%)
